# Integrative analysis of 10,000 epigenomic maps across 800 samples for regulatory genomics and disease dissection

**DOI:** 10.1101/810291

**Authors:** Carles B. Adsera, Yongjin P. Park, Wouter Meuleman, Manolis Kellis

**Affiliations:** Computer Science and Artificial Intelligence Laboratory, Massachusetts Institute of Technology, Cambridge, MA, USA; Broad Institute of MIT and Harvard, Cambridge, MA, USA; Computational and Systems Biology Program, Massachusetts Institute of Technology, Cambridge, MA, USA; Altius Institute for Biomedical Sciences, United States

## Abstract

To help elucidate genetic variants underlying complex traits, we develop EpiMap, a compendium of 833 reference epigenomes across 18 uniformly-processed and computationally-completed assays. We define chromatin states, high-resolution enhancers, activity patterns, enhancer modules, upstream regulators, and downstream target gene functions. We annotate 30,247 genetic variants associated with 534 traits, recognize principal and partner tissues underlying each trait, infer trait-tissue, tissue-tissue and trait-trait relationships, and partition multifactorial traits into their tissue-specific contributing factors. Our results demonstrate the importance of dense, rich, and high-resolution epigenomic annotations for complex trait dissection, and yield numerous new insights for understanding the molecular basis of human disease.

## Introduction

Genome-wide association studies (GWAS) have been extremely successful in discovering more than 100,000 genomic loci containing common single-nucleotide polymorphisms (SNPs) associated with complex traits and disease-related phenotypes, providing a very important starting point for the systematic dissection of the molecular mechanism of human disease^1,2^. However, the vast majority of these genetic associations remain devoid of any mechanistic hypothesis underlying their molecular and cellular functions, as more than 90% of them lie outside protein-coding exons and likely play non-coding roles in gene-regulatory regions whose annotation remains incomplete^3–6^.

To help annotate non-coding regions of the genome, large-scale experimental mapping of epigenomic modifications associated with diverse classes of gene-regulatory elements has been undertaken by several large consortia, including the ENCyclopedia of DNA Elements (ENCODE), Roadmap Epigenomics, and the Genomics of Gene Regulation (GGR)^7–9^. Integration of these datasets, and in particular histone modification marks and DNA accessibility maps, has helped infer chromatin states and annotate diverse classes of gene-regulatory elements, including distal-acting and tissue-specific enhancer regions and proximal-acting and mostly-constitutive promoter regions^10–12^ across 16 cell types by ENCODE in 2012^7,8^, and 111 cell and tissue types by Roadmap Epigenomics in 2015^7,8^, These maps have been widely used to gain insights into the molecular basis of complex traits by recognizing the preferential localization (enrichment) of genetic variants associated with the same traits within gene-regulatory elements active in the same tissue or cell type^3,8,13–16^. These enrichments can help gain insights into the tissues and cell types that may underlie complex disorders, and within which the molecular effects of these genetic variants may first manifest^17–19^. They can also help fine-map likely causal genetic variants in regions of linkage disequilibrium^20,21^ by prioritizing those variants that lie within enriched annotations^16,22–24^.

However, these maps also have great limitations. First, they remain highly incomplete, covering only a small fraction of human tissue diversity, and missing many tissues of great relevance to human disease. Second, the quality of reference epigenomes is highly variable, as each reference epigenome typically relies on only 1-2 replicates, making them more prone to experimental noise and even variation between experimental protocols, experimentalists, labs, antibody lots, or reagent batches.

Third, in addition to experimental noise, differences in the computational processing pipelines of different consortia and across different versions of processing software and integration pipelines can lead to dramatic differences in the regions annotated in different chromatin states, or within accessible regions called by different peak-calling algorithms. Fourth, maps that rely on multiple epigenomic marks are inherently limited in their dimensionality, as they need to exclude samples that do not contain all marks incorporated, or marks that are not present in all the samples. Lastly, epigenomic reference profiling projects must balance exploring biological space with exhaustively performing assays in each sample, leading to incomplete epigenomic annotation matrices with few marks across many samples, and few samples with many marks.

Here, we overcome these limitations and present a new reference of the human epigenome, EpiMap, by incorporation of 1698 new datasets across three consortia, joint uniform re-processing of a total of 3030 reference datasets, and computational completion of 14,952 epigenomic maps across 859 tissues and 18 marks. We rely on epigenomic imputation^25–27^, which maximizes the consistency and quality of an ensemble of epigenomic maps by leveraging correlation patterns between related assays and between related cell types to infer missing datasets, and to generate high-quality predictions of experimentally-profiled datasets using all experimentally-profiled datasets.

We show that EpiMap greatly surpasses previous reference maps in scope, scale, and coverage of biological space. We combine single-mark maps into a multi-mark chromatin state reference for each of 833 high-quality epigenomes, combine multiple enhancer chromatin states with DNA accessibility to infer a high-resolution enhancer map for each cell type, group enhancers into modules based on their common activity patterns across cell types, and use these modules to infer candidate upstream regulators based on motif enrichments, and downstream gene functions based on gene ontology enrichments, providing an important reference for studies of gene regulation both for disease dissection and for basic biological studies of human tissues.

Lastly, we use EpiMap to increase our understanding of the molecular processes underlying complex traits and human disease by systematic integration of genetic variants associated with 926 traits and high-resolution enhancer annotations across 833 tissues. Compared to the Roadmap Epigenomics resource analysis^8^, we achieve a nearly 10-fold increase in the number of traits showing significant epigenomic enrichments (N=534) and a dramatic increase in the number of loci with putative causal variants within enriched enhancer annotations (N=30k). We also exploit the greatly-increased number of samples by developing a new hierarchical approach for narrowing down the tissue-level resolution of GWAS enrichments thus resulting in many new enriched traits and loci, to distinguish multifactorial and polyfactorial traits from unifactorial traits, to learn principal-partner tissue pairs that cooperate to explain multifactorial traits, to partition multifactorial-trait SNPs by tissue thus revealing the distinct biological processes and explaining trait comorbidity patterns, and to learn a trait-trait relationship network that helps guide the interpretation of unifactorial and multifactorial traits.

These results indicate that EpiMap is a valuable new reference for both gene-regulatory studies and disease studies seeking to elucidate the molecular basis of complex disorders.

## Results

### Generation and validation of EpiMap reference epigenome compendium

We generated EpiMap, the largest integrated compendium of epigenomics maps to-date, spanning 859 epigenomes across 18 epigenomic assays (**Fig. 1a, Supp. Fig. S1**) by aggregation, uniform processing, and computational completion of three major resources: Roadmap Epigenomics^8^ (425 samples, of which 241 were generated since the 2015 data freeze), ENCODE^7^ (434 samples of which 381 were generated since the 2012 data release), and Genomics of Gene Regulation (GGR) consortium^28^ (25 samples). We uniformly processed a total of 3,030 datasets, including 1994 ChIP-seq experiments for 33 histone marks, 701 experiments for DNA accessibility (DNase-seq, ATAC-seq), and 335 experiments for chromatin-associated general factors. We used each unique combination of biosample, donor, sex, age, and lifestage, and removed samples with genetic perturbations (Supplementary Table S1).

**Figure 1.**
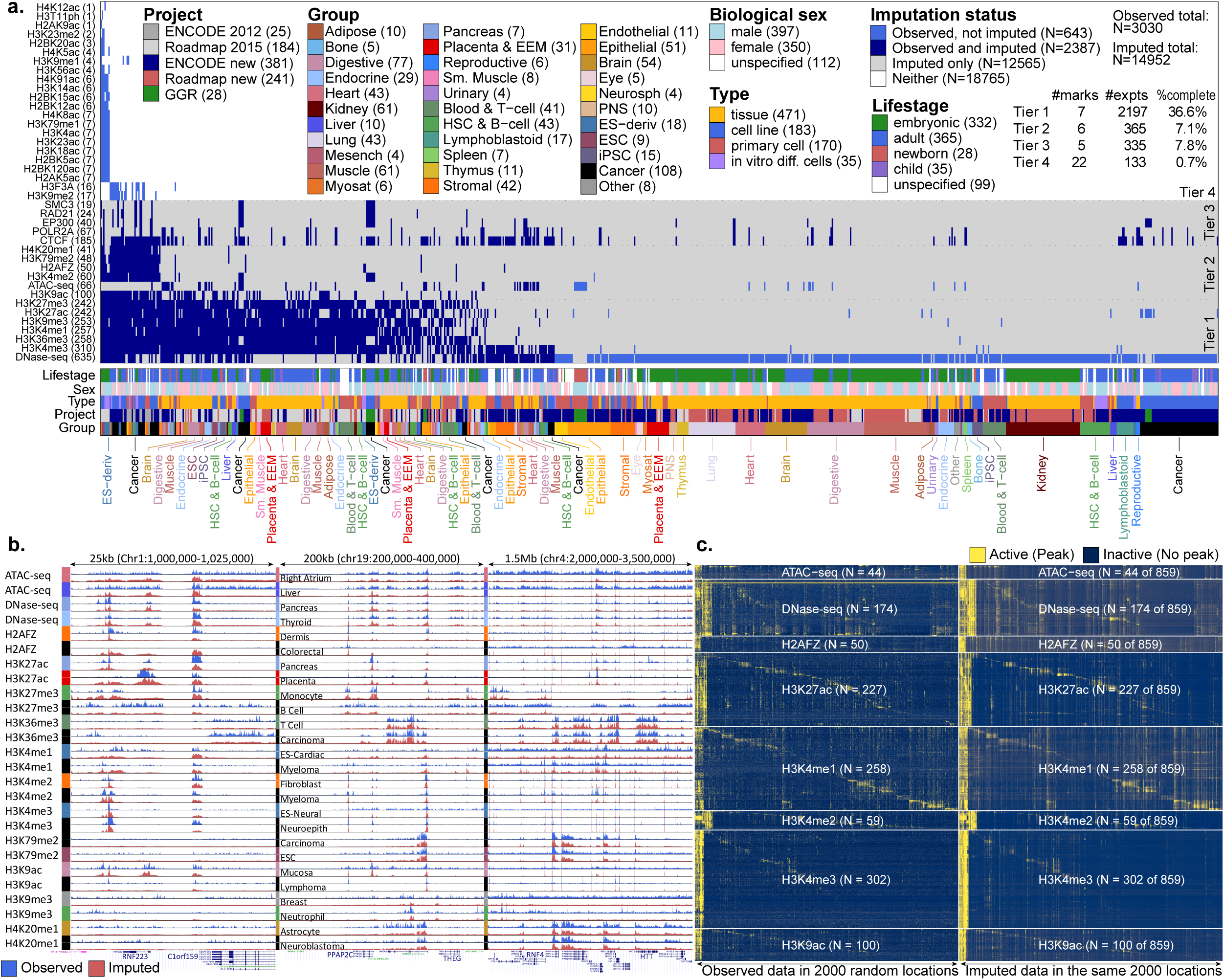
EpiMap resource overview. **a**. EpiMap data matrix across 859 samples (columns) and 35 assays (rows), ordered by number of experiments (parentheses) and colored by metadata. **b**. Paired observed (blue) and imputed (red) tracks for all Tier 1 and Tier 2 assays in three regions at different resolutions for randomly-selected samples. Full tracks at https://epigenome.wustl.edu/epimap. **c**. Heatmap of paired observed and imputed signal intensity across all punctate Tier 1 and Tier 2 assays across 2000 highest-max-signal bins among 5000 randomly-selected 25bp bins. Samples (rows) and bins (columns) are clustered and diagonalized using maximum imputed signal intensity, with broadly-active regions shown first.

We classified these 3,030 assays into multiple tiers, according to their completeness. Tier 1 assays (7 marks, 2197 total experiments, 36% complete on average) include DNase-seq (accessible chromatin), H3K4me3 (promoters), H3K36me3 (transcribed), H3K4me1 (poised enhancers), H3K9me3 (heterochromatin), H3K27ac (active enhancers/promoters), and H3K27me3 (polycomb repression). Tier 2 assays (6 marks, 365 experiments, 7% complete on average) include H3K9ac, ATAC-seq, H3K4me2, H2AFZ, H3K79me2, and H4K20me1, in order of abundance. Tier 3 assays consist of general factors associated with key biological processes (5 factors, 335 total experiments, 8% complete on average), including POLR2A (transcription initiation), EP300 (enhancer activation), CTCF (insulation, chromatin looping), SMC3 (cohesin), and RAD21 (cohesin component, double-stranded-break repair). Lastly, Tier 4 assays (22 marks, 133 experiments, 0.7% complete on average) consist of 16 acetylation marks, 4 methylation marks (H3K9me2, H3K79me1, H3K9me1, H3K23me2), H3F3A, and H3T11ph.

We generated a total of 14,952 imputed datasets covering 18 marks and assays across 859 samples using ChromImpute^25^. We constructed genome-wise browser visualization tracks available from the WashU Epigenome Browser^29^. Imputed tracks showed strong agreement with high-quality observed tracks, individual enhancer elements, precise boundaries in histone modification peaks, and finer-grained features of epigenomic assays, such as dips in H3K27ac peaks indicative of nucleosome displacement (**Fig. 1b**). They also captured the density, location, intensity, and cell-type-specificity of both active and repressed marks across thousands of regions and all 859 samples (**Fig. 1c**). We found 85% peak recovery and 75% average genome-wide correlation for puncate marks representing 59% of tracks (**Supp. Fig. S2**).

Disagreement between imputed and observed tracks helped flag 138 potentially problematic datasets which showed markedly lower QC scores (**Supp. Fig. S2, S3**), and revealed potential sample or antibody swaps (**Supp. Fig S4, S5**) some of which were independently flagged by the data producers. We also used the difference between observed and imputed datasets to recognize 15 experiments with potential antibody cross-reactivity or secondary specificity (**Supp. Fig S6-S8**). We removed from subsequent analyses the 138 flagged datasets and 442 tracks based solely on ATAC-seq or low-quality DNase-seq data as they showed lower correlations, resulting in 2,850 observed and 14,510 imputed marks across 833 samples used in the remainder of this work.

### Sample relationships, chromatin states, and high-resolution enhancer mapping

The resulting compendium of 833 high-quality epigenomes represents a major increase in biological space coverage, with 75% of epigenomes (624 of 833) corresponding to new biological specimens across 33 tissue groups (categories), providing the opportunity to study their relationships systematically, providing insights into the primary determinants of the epigenomic landscape. Both hierarchical and two-dimensional embedding^30^ clustering of the genome-wide correlation patterns of multiple marks (**Fig. 2a,b, Supp. Fig. S9-S11**) grouped these samples firstly by lifestage (adult vs. embryonic) and sample type (complex tissues vs. primary cells vs. cell lines), and secondly by distinct groups of brain, blood, immune, stem-cell, epithelial, stromal, and endothelial samples within them. By contrast, donor sex was not a primary factor in sample grouping.

**Figure 2.**
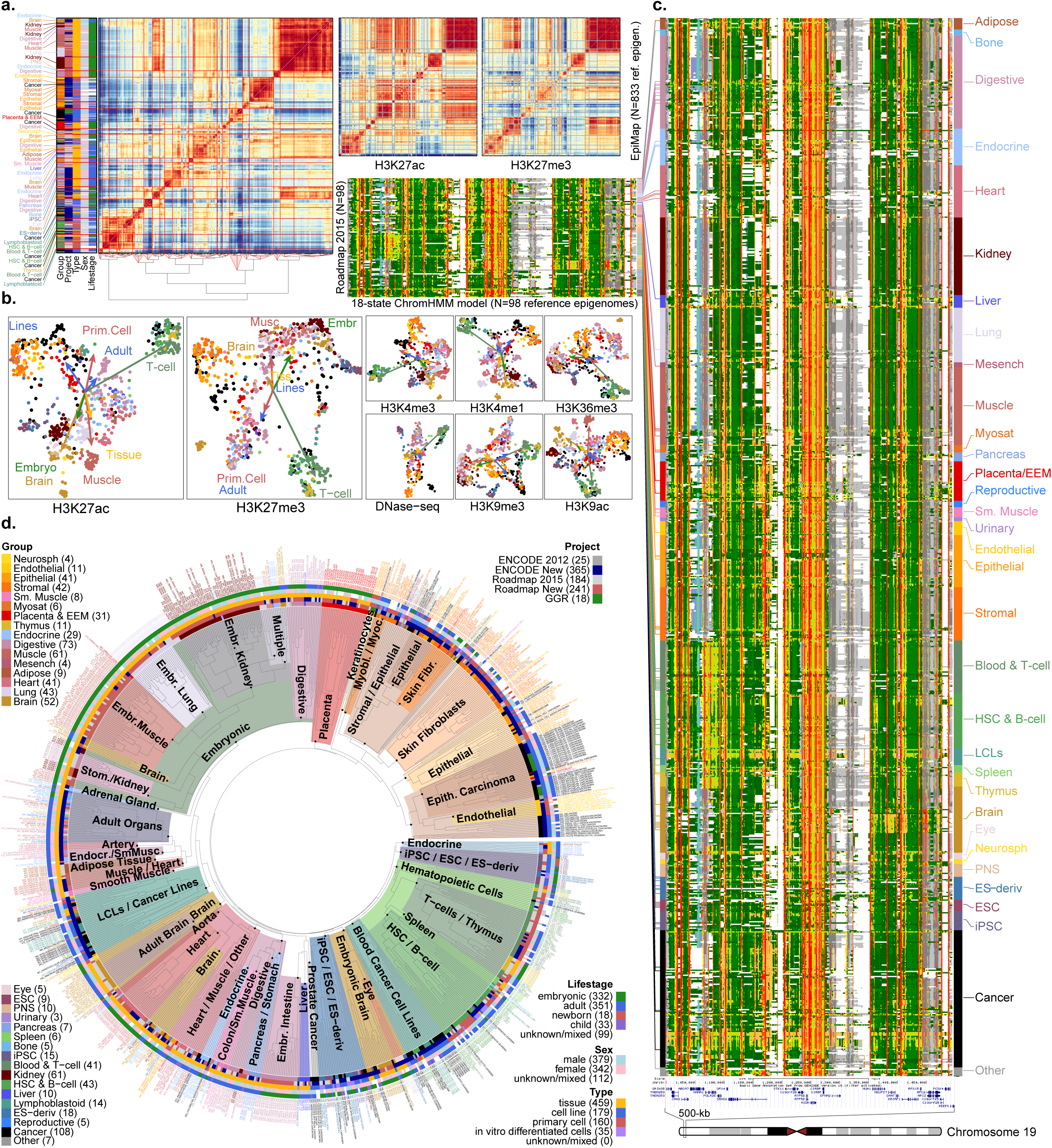
Reference epigenome relationships. **a**. Genome-wide correlation across all 833 samples using all Tier 1 marks, hierarchically clustered (left), only H3K27ac (middle), and only H3K27me3 (right). **b**. Two-dimensional embeddings of Tier 1 and 2 marks colored by tissue group, using Spearman correlation within matched chromatin states. Arrows point to average of specified groups. **c**. Chromatin state annotations for 98 Roadmap (left) and 833 EpiMap (right) epigenomes using an 18-state ChromHMM model^8^ based on all Tier 1 histone marks, with colored lines connecting matched epigenomes. **d**. Hierarchical clustering of 833 reference epigenomes based on enhancer activity distances (**Supp. Fig. S19**). Subtrees enriched for specific sample types are highlighted and labeled (colors). Samples labeled with reduced sample names and colored by metadata (**Supp. Table S1**).

We clustered samples using each individual mark (**Fig. 2a,b**) evaluated within mark-specific relevant genomic regions. We found that active marks (H3K27ac, H3K4me1, H3K9ac) primarily group samples by differentiation lineage, leading to separate blood, immune, spleen, thymus, epithelial, stromal, and endothelial clusters. These marks also grouped lung, kidney, heart, muscle, and brain samples each into distinct clusters, regardless of whether they were adult or embryonic. By contrast, repressive marks (H3K27me3, H3K9me3) better captured the stage of differentiation, grouping together pluripotent, iPSC-derived, and embryonic samples, which were separated into different groups by active marks. In each case, observed and imputed data co-clustered, but imputed datasets better captured the continuity between different sample types, and more clearly revealed the grouping of sample types, likely driven both by their cleaner signal tracks and their sheer number.

We generated a reference epigenomic annotations for each for the 833 samples, using combinations of histone modification maps to map genome-wide locations of 18 chromatin states^7,8,10^ (**Fig. 2c**), including multiple types of enhancer, promoter, transcribed, bivalent, and repressed regions (**Supp. Fig. S12**). We applied this model on a scaled mixture of observed and imputed data, excluding the 138 flagged observed datasets (see Methods). We found broad consistency in state coverage (**Supp. Fig. S13, S14**) and state definitions (**Supp. Fig. S15**) across cell types.

We also generated a high-resolution annotation of active enhancer regions by intersecting 3.5M accessible DNA regions from 733 DNase-seq experiments (Meuleman et al., in preparation) with five active enhancer states, resulting in 2.1M high-resolution enhancer regions (57% of all DNase regions), covering 0.8% of the genome in each sample on average, and together capturing 13% of the human genome (**Fig. 3a, Supp. Fig. S18**) a 31% increase from previous maps (**Supp. Fig S19**), and constructed an ‘enhancer-sharing tree’ (**Fig. 2d, Supp. Fig. S16, S17**) relating all 833 samples in a nested set of binary groupings based on their number of shared active enhancers.

**Figure 3.**
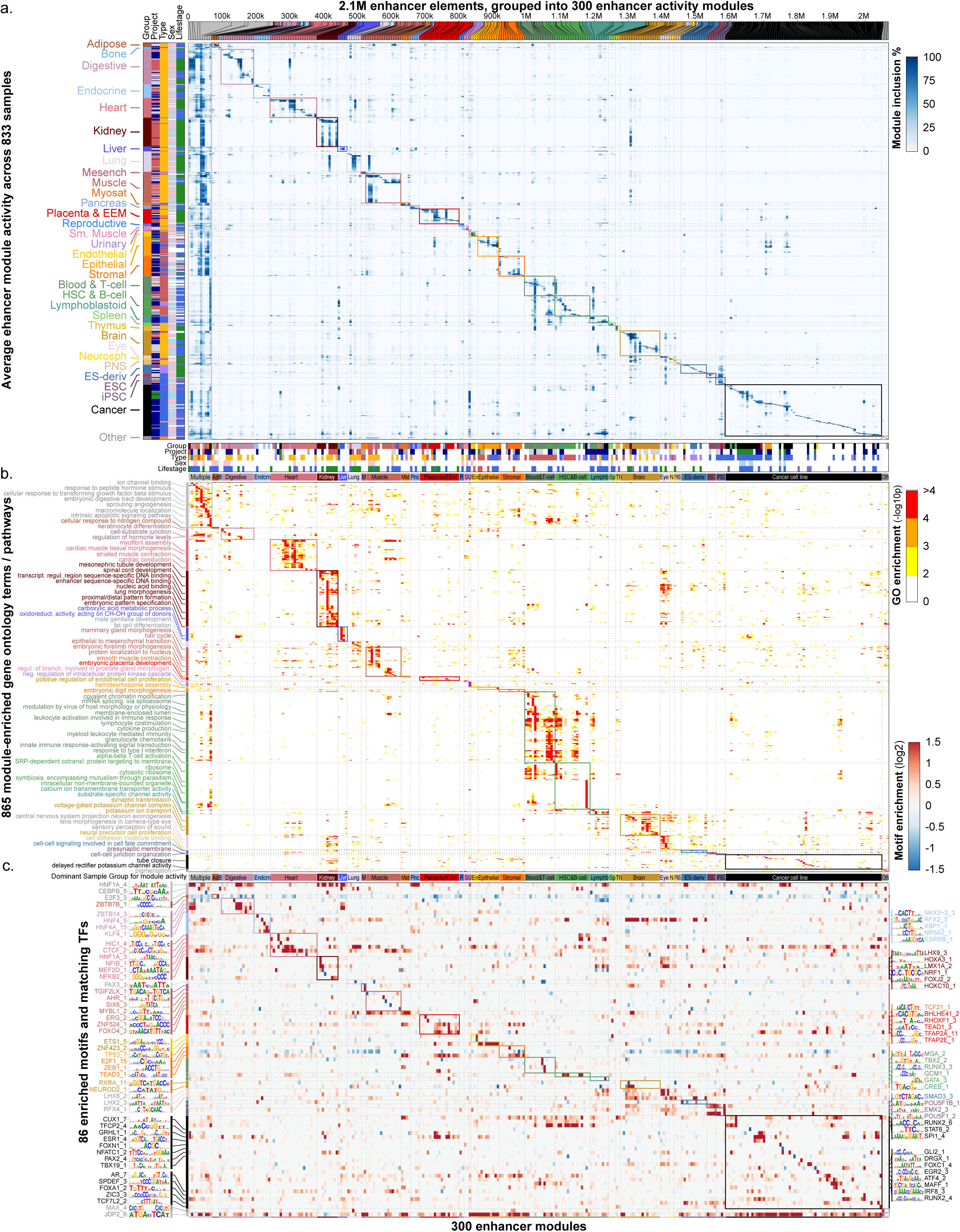
Enhancer module circuitry. **a**. Clustering of 2.1M enhancer elements (top) into 300 modules (columns) using enhancer activity levels (heatmap) across 833 samples (rows), quantified by H3K27ac levels within accessible enhancer chromatin states. Bottom panel shows enrichment of each module for each metadata annotation, highlighting 34 groups of modules (separated by dotted lines): 33 sample-type-specific (colored boxes) and 1 multiply-enriched (left-most). **b**. Gene ontology^31,59^ (GO) enrichments (heatmap) for each module (columns) across 865 terms (rows) with P<e-4. GO terms colored by maximal enrichment group. Only 72 representative terms are shown, chosen by a bag-of-words approach within each tissue group. **c**. Motif enrichment (heatmap) for each module (columns) across 86 motif clusters (rows) with enrichment log_2_FC>1.5. Motifs colored by module of maximal enrichment.

### Regulatory genomics of enhancer modules, target biological processes, and upstream regulators

For each high-resolution enhancer region, we defined its activation state in each of the 833 epigenomes using flanking nucleosome H3K27ac levels, and used it to group enhancers with coordinated activity into 300 enhancer modules (**Fig. 3a and Supp. Fig S20-S22**). We distinguished 290 tissue-specific modules (1.8M enhancers, 88%) with activity in only 2% of samples on average, and 10 broadly-active modules (251k enhancers, 12%) showing activity across 77% of sample categories on average. Embryonic modules combined multiple tissue types, including heart, lung, kidney, muscle, and digestive system, while adult modules separated internal organs.

We predicted candidate gene-regulatory roles of each enhancer module based on highly-significant gene ontology enrichment of its target genes^31^, revealing fine-grained tissue-specific biological processes (**Fig. 3b and Supp. Fig S23**), including: ion channels (for brain modules), camera-type eye development (eye modules), neural precursor cell proliferation (neurosphere modules), endothelial proliferation, hemidesmosomes, and digit morphogenesis (endothelial, stromal, and epithelial modules), and organ development and morphogenesis (embryonic modules). These enrichments can help guide the elucidation of target genes and biological processes controlled by individual enhancers within each module. Endocrine, mesenchymal, pancreas, and reproductive modules lacked any gene ontology enrichments, possibly due to small counts or still underexplored areas of biology.

We found significant motif enrichments for 202 modules (67% of 300) across 86 motifs (**Fig. 3c**), both suggesting potential new regulators, and confirming known roles for well-studied factors, including: GATA and SPI1 for blood and immune groups^32^; NEUROD2 and RFX4 for brain and nervous system^33,34^; POU5F1 for iPSCs^35^; KLF4 for digestive tissues^36^; MEF2D for skeletal and digestive muscle^37^; TEAD3 for placenta, myosatellite and epithelial cells^38^; and HIC1 in heart, HSC, and B-cells^39^. Several motifs distinguished developmental vs. adult tissues, including: NEUROD2 for embryonic-only brain; NFIB for embryonic-only heart, lung, and muscle; MEF2D for adult-only heart, lung, and muscle; and AR for adult-only adipose, heart, liver, and muscle. Motifs were generally highly specific, with 95% (N=82) enriched in only 3% of modules on average, but 4 “promiscuous” motifs were enriched across 79 modules on average, including: RFX1-5 (44 modules) in brain and testis^40^; GRHL1 (52 modules) in trophoblast and epithelial differentiation^41,42^; HNF1A/B (67 modules) in liver, kidney, and pancreas^43^; and JDP2/AP-1 (152 modules) in immune and bone development, cancer, and response to diverse stimuli^44,45^.

### New enhancer annotations help interpret trait-associated SNPs from genome-wide associations studies

We next used our 2.1M enhancer annotations and their tissue specificity to interpret genetic variants associated with complex traits, which are increasingly recognized to play gene-regulatory roles^3,8,13^. We assembled a compendium of 926 well-powered genome-wide association studies (GWAS) from the NHGRI/Ensembl GWAS catalog^1^ that have at least 20,000 cases in the initial study sample. These capture 66,801 associations in 33,417 regions, corresponding to 59% of the 113,655 pruned associations in the GWAS catalog^46^ (even though these studies only account for 17% of the 5,454 GWAS publications to date), providing an important benchmark for epigenomic enrichment studies.

For each trait, we calculated the enrichment of trait-associated SNPs in active enhancers from each of our 833 epigenomes, resulting in 30,844 significant trait-tissue pairs, implicating 534 unique traits and all 833 samples (hypergeometric statistic, FDR<1%) (**Fig. 4a and Supp. Fig. S24**). These capture 30,247 SNPs in enriched annotations (45% of all loci 66,801 loci), providing invaluable insights for epigenomics-based fine-mapping of SNPs within regions of extended linkage disequilibrium. The number of enriched traits represents a nearly 10-fold increase from 58 traits^8^ enriched in H3K4me1 and 54 traits^25^ enriched in H3K27ac reported by the Roadmap Epigenomics project, due to methodological improvements, increases in GWAS results and power, and more available epigenomes. Quantifying the latter component, the new reference epigenomes capture the strongest GWAS enrichment in 75% of cases (402 of 534 traits) (**Fig. 4b)**, and provide the only significant enrichment in 22% of cases (116 traits) (**Fig. 4c**).

**Figure 4.**
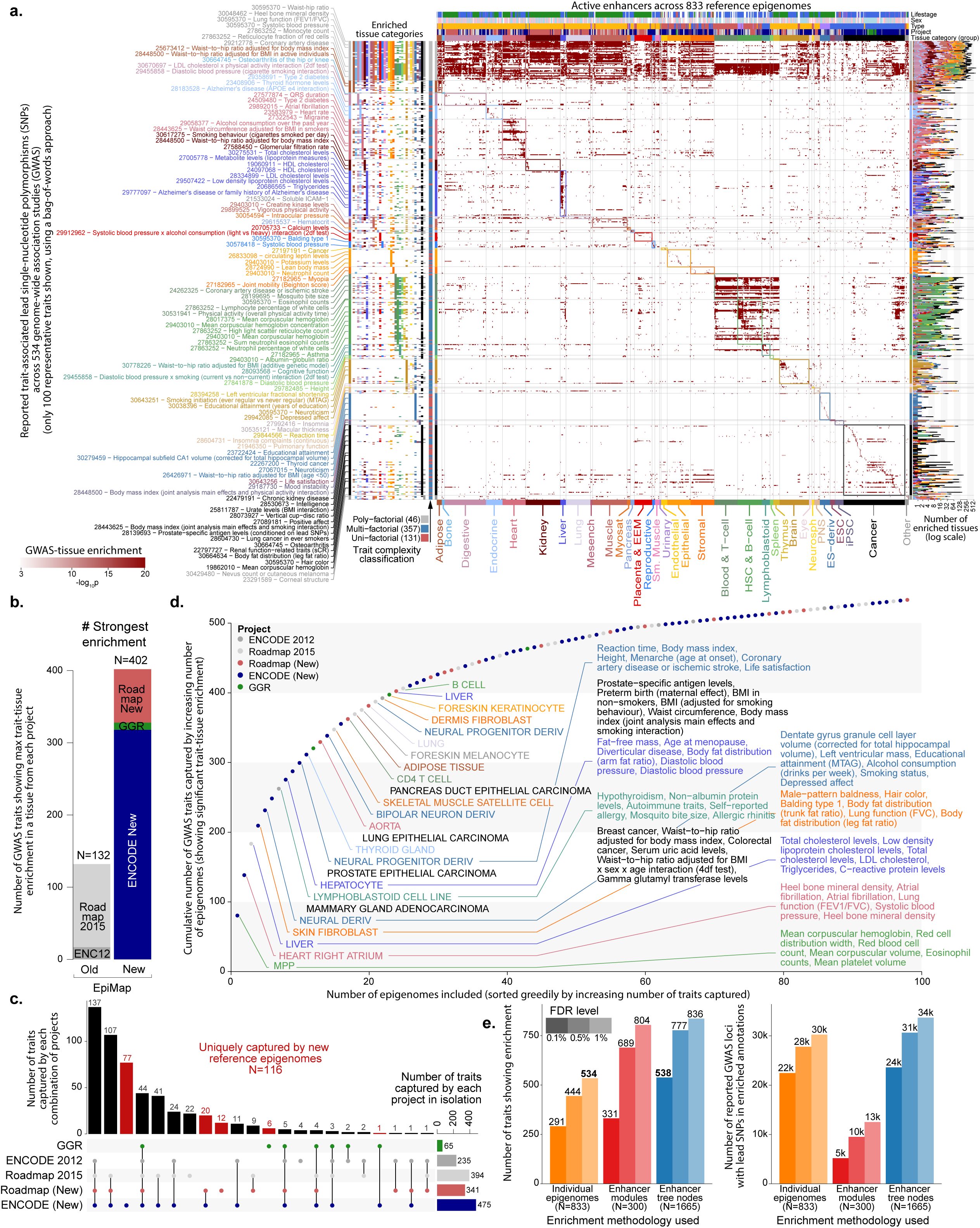
GWAS tissue-prioritization. **a**. Trait-tissue enrichment (center, heatmap) between reported lead single-nucleotide polymorphisms (SNPs) from 534 genome-wide association studies (rows) and accessible active enhancers across 833 epigenomes (columns) (FDR<1%). Enriched tissue groups (left) and number of enriched epigenomes (right) shown for each trait. Only 100 representative traits labeled, using a bag-of-words approach (full list of traits in Supplementary Fig. S30). Traits colored by sample with maximal trait-tissue enrichment. **b**. Contribution of each project to the maximum GWAS trait-tissue enrichment for the 534 traits with significant enrichments. **c**. Number of traits (y-axis) with significant GWAS trait-tissue enrichments for each combination (column) of projects (rows). **d**. Increase in the cumulative number of GWAS traits (y-axis) with significant trait-tissue enrichments with increasing numbers of epigenomes (x-axis), ordered to maximize the number of novel trait annotations captured with each new epigenome. Top 25 samples labeled and colored by tissue group, with top 6 GWAS traits shown for the first 8 samples. Points colored by project. All 534 traits are captured after inclusion of 98 samples. **e**. Comparison of GWAS enrichments found (y-axis, left) and number of lead SNPs in significantly-enriched annotations (y-axis, right) using different methodologies (x-axis) for three FDR cutoffs (shades).

We also studied the number of traits for which enrichments are found with increasing numbers of epigenomes (**Fig. 4d**). New epigenomes represent 71 of the top 98 epigenomes that together capture at least one significant GWAS enrichment, and 174 of the top 238 reference epigenomes that together capture all maximal GWAS enrichments (**Supp. Fig. S25**). The tissue categories represented in top contributing samples are very diverse, including blood (which captures several blood traits), heart right atrium (atrial fibrillation, PR interval), liver (cholesterol, metabolites), skin fibroblasts (baldness, hair color), neural cells (educational attainment), adenocarcinoma cells (breast cancer), and LCLs (immune traits). To capture the full space of enriched GWAS traits, nearly the entire tree of sample types was needed, highlighting the importance of broadly sampling cells and tissues to understand the molecular basis of complex disorders.

### Clustering and hierarchical analysis of GWAS enrichments and tissue sharing

Given the block-like structure of the trait-tissue enrichment matrix (**Fig. 4a**), with many biologically-similar traits showing enrichment in samples of common tissue groups, we also calculated trait enrichments for enhancer modules, seeking to capture the sharing between epigenomes (**Supp. Fig. S26, S27**). This resulted in a 51% increase in the number of traits showing epigenomic enrichments (from 534 to 804 at 1% FDR) (**Fig. 4e and Supp. Fig. S28**), indicating that the common biological processes captured by enhancer modules are also relevant to complex trait interpretation.

Beyond the single-resolution grouping of our modules, we also carried out a multi-resolution analysis of GWAS enrichments using our enhancer-sharing tree (**Fig. 2d**) and reporting significant internal node enrichments relative to parent nodes, thus pinpointing the likely biological specificity at which genetic variants may act (see Methods). This further increased the number of traits showing epigenomic enrichments to 836 (51% relative to individual epigenomes, 4% relative to modules), and substantially increased the number of lead SNPs falling in enriched annotations to 33,706 (11% relative to individual epigenomes, 169% relative to modules) at the same FDR (1%), indicating that multi-resolution approaches can capture SNP-enhancer intersections missed by other approaches (**Fig. 4e, Supp. Fig. S28**).

We next studied the fraction of GWAS traits for which a tissue showed maximal enrichment to distinguish ‘principal’ tissues (e.g. immune cells, liver, heart, brain, adipose) that typically showed the strongest enrichments in GWAS traits they are enriched in, suggesting they more frequently play driver roles (**Fig. 5a**). By contrast, ‘partner’ tissues (e.g. digestive, lung, muscle, epithelial) were enriched in many traits but rarely showed the maximal enrichment, suggesting they may play auxiliary roles. Several tissue pairs showed common trait enrichments significantly more frequently than expected by chance (**Fig. 5b**) in the context of distinct and biologically-meaningful traits (**Fig. 5c**), including liver and adipose (in the context of cholesterol and triglyceride traits), liver and digestive (metabolite traits), liver and immune cells (eating disorders), adipose and endothelial (sleep duration, waist-to-hip ratio), adipose and heart (atrial fibrillation), and adipose and muscle (gestational age), providing a guide for understanding multifactorial traits.

**Figure 5.**
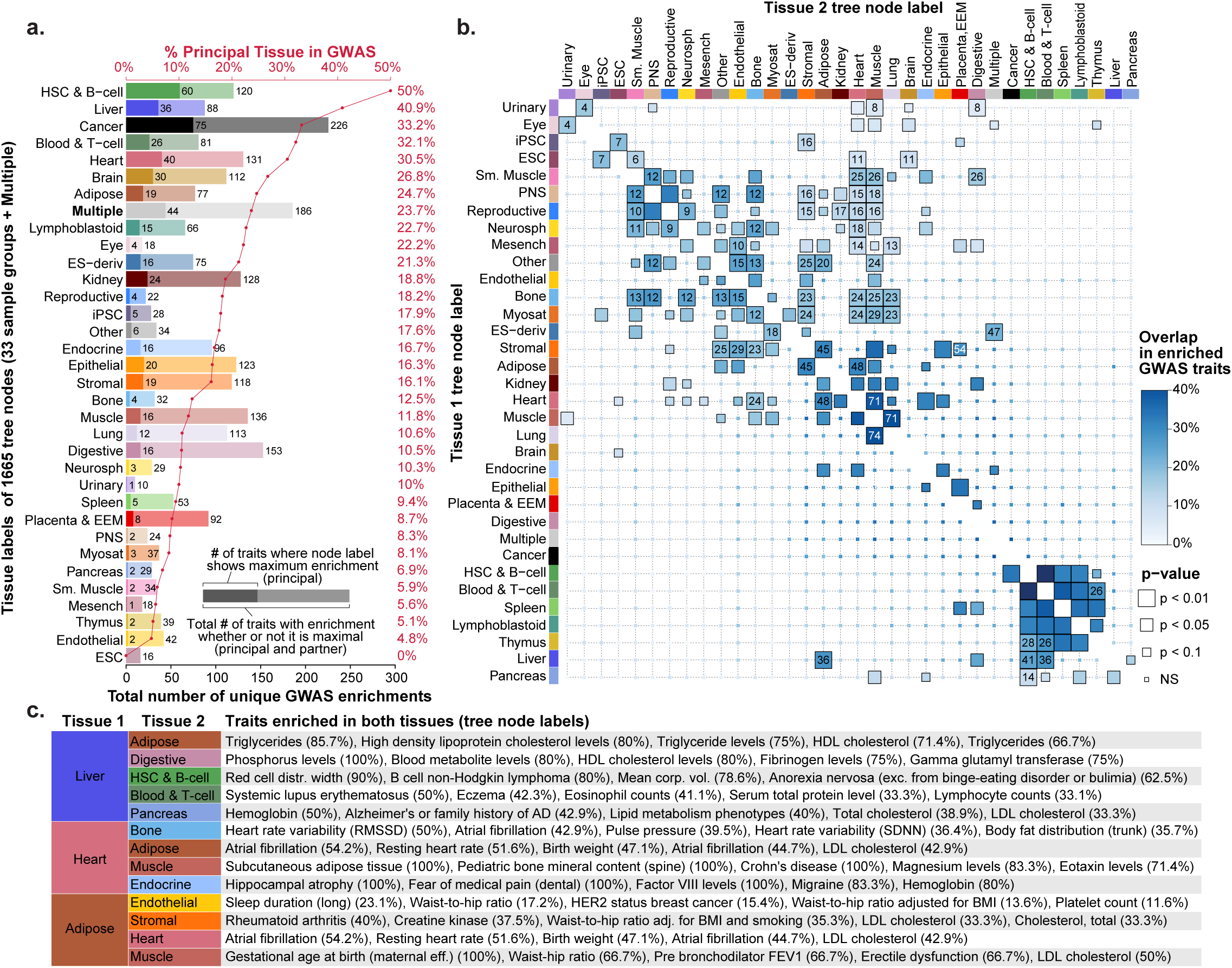
Principal and partner tissue enrichments. **a**. For each tree node label (rows), the number of GWAS traits (black x-axis, bottom) showing maximum enrichment in that tree node (dark bars, principal tissue) or any enrichment in that tree node (light bars, partner tissue), and the percentage of tissue-enriched traits for which the tissue shows the maximal enrichment (red x-axis, top) across 538 traits. **b**. Overlap in enriched GWAS traits between pairs of tissues with maximal enrichment in the trait (principal tissue, rows) and lower enrichment in the same trait (partner tissue, columns), using tree node labels. **c**. Top traits in significant interactions for selected tissue pairs (Liver, Endocrine, Muscle, Heart, Adipose, PNS). For each pair of co-enriched tissue groups we reported the top 5 GWAS by their percent of significant enrichments coming from either group.

### Partitioning multifactorial traits and trait combinations into their tissues and pathways of action

We used the number of distinct tissue categories enriched in each trait (**Fig. 4a; Supp. Data S1**) to distinguish 303 ‘unifactorial’ traits (56%) with most enriched nodes in only one tissue group (e.g. QT interval in heart, educational attainment in brain, hypothyroidism in immune cells), indicating a more constrained set of biological processes involved (**Fig. 6a**). Another 146 ‘multifactorial’ traits (27%) were enriched on average in 5 different tissue categories indicating multiple modes of action, including: Alzheimer’s disease (AD) in both immune and brain tissues^47,48^; waist-to-hip ratio (adjusted for BMI)^49^ in adipose, muscle, kidney, and digestive tissues; and healthspan in ES, T cells, adipose, and digestive tissues. A subset of 92 ‘polyfactorial’ traits (17%) implicated an average of 14 tissue categories each (**Fig. 6c**), including coronary artery disease (CAD)^50^ with 19 different tissue groups, including liver, heart, adipose, muscle, and endocrine samples.

**Figure 6.**
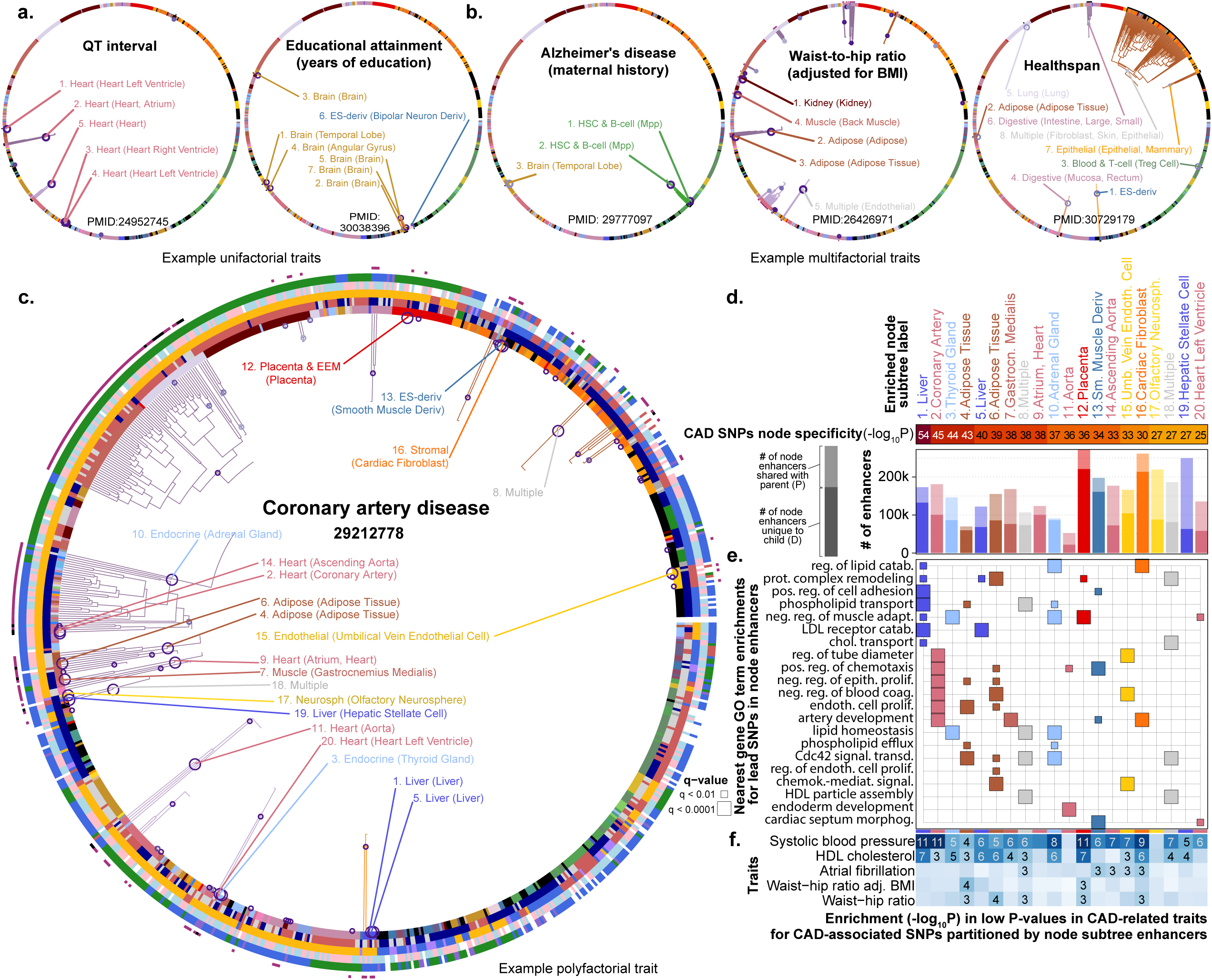
Examples of trait epigenomic enrichments on enhancer sharing tree, including. **a**. single-tissue (QT, Macular) and **b**. multi-tissue (AD, WHR) traits, highlighting nodes with FDR < 0.1% (labels = top 5 by −log_10_p). **c**. Epigenomic enrichments for Coronary Artery Disease (CAD, PubMedID 29212778) on enhancer activity tree. Nodes passing FDR < 0.1% are labeled by rank, tissue group, and top components, and their subtrees are drawn with solid lines (large circles = top 20 nodes by −log_10_p). Leaves annotated by metadata and number of enriched parent nodes (outer, red=1, black=2). **d**. Top 20 enriched nodes for CAD with nominal p-values (heatmap) and shared enhancer set sizes (barplot) with number at the subtree (full bar) and number of differential enhancers between the node and its parent (tested set, dark bar). **e**. GO enrichments of node enhancers with lead SNPs (nearest expressed genes), colored by each node’s tissue group and diagonalized (over-representation test). **f**. Enrichment for significant loci in overlap of CAD loci with loci from five related traits, within enriched enhancers in each node (heatmap, −log_10_p of one-tailed Mann-Whitney test against each trait’s loci in enhancer annotations).

We next used the enriched tissues of multifactorial traits to partition their associated SNPs into (potentially-overlapping) sub-groups, which were enriched in distinct biological pathways, thus revealing distinct processes through which multifactorial traits may act (**Fig. 6d, Supp. Fig. S29**). For example, 339 CAD-associated SNPs in enriched enhancers partitioned into: 212 SNPs in heart enhancers that preferentially localized near artery, cardiac, and vessel morphogenesis genes; 121 SNPs in endocrine enhancers, which enriched in lipid homeostasis; 122 SNPs in adipose enhancers, which enriched in axon guidance/extension and focal adhesion, consistent with adipose tissue innervation processes; 169 SNPs in liver enhancers, which enriched in cholesterol/lipid metabolism and transport; and 112 SNPs in ES-derived muscle cells, which enriched in septum morphogenesis, cardiac chamber and aorta development.

This partitioning of genetic loci into tissues also helped inform the shared genetic risk between pairs of co-enriched traits, by revealing the tissues that may underlie their common biological basis (**Fig. 6d**). For example, the same partitioning of CAD loci showed that CAD loci in heart, muscle, and endothelial enhancers were preferentially also associated with high blood pressure and atrial fibrillation risk loci. However, CAD loci in liver and endocrine enhancers were instead associated with systolic blood pressure^51^. Similarly CAD loci also associated with waist-to-hip ratio^49,51,52^ overlapped adipose but not liver, endocrine, or heart enhancers, and CAD loci associated with HDL cholesterol^53^ overlapped liver, adipose, and endocrine enhancer but not heart tissues.

### Tissue enrichment and co-enrichment patterns paint network of complex trait associations

We next classified each GWAS trait according to its enriched tissues, and linked traits showing similar enrichment patterns into a trait-trait co-enrichment network (**Fig. 7, and Supp. Fig. S30, S31**). Unifactorial traits formed the cores of highly-connected communities, including: cognitive and psychiatric traits in brain and neurons; heart beat intervals in heart; cholesterol measures in liver; filtration rate in kidney; immune traits in T-cells; and blood cell counts in hematopoietic cells.

**Figure 7.**
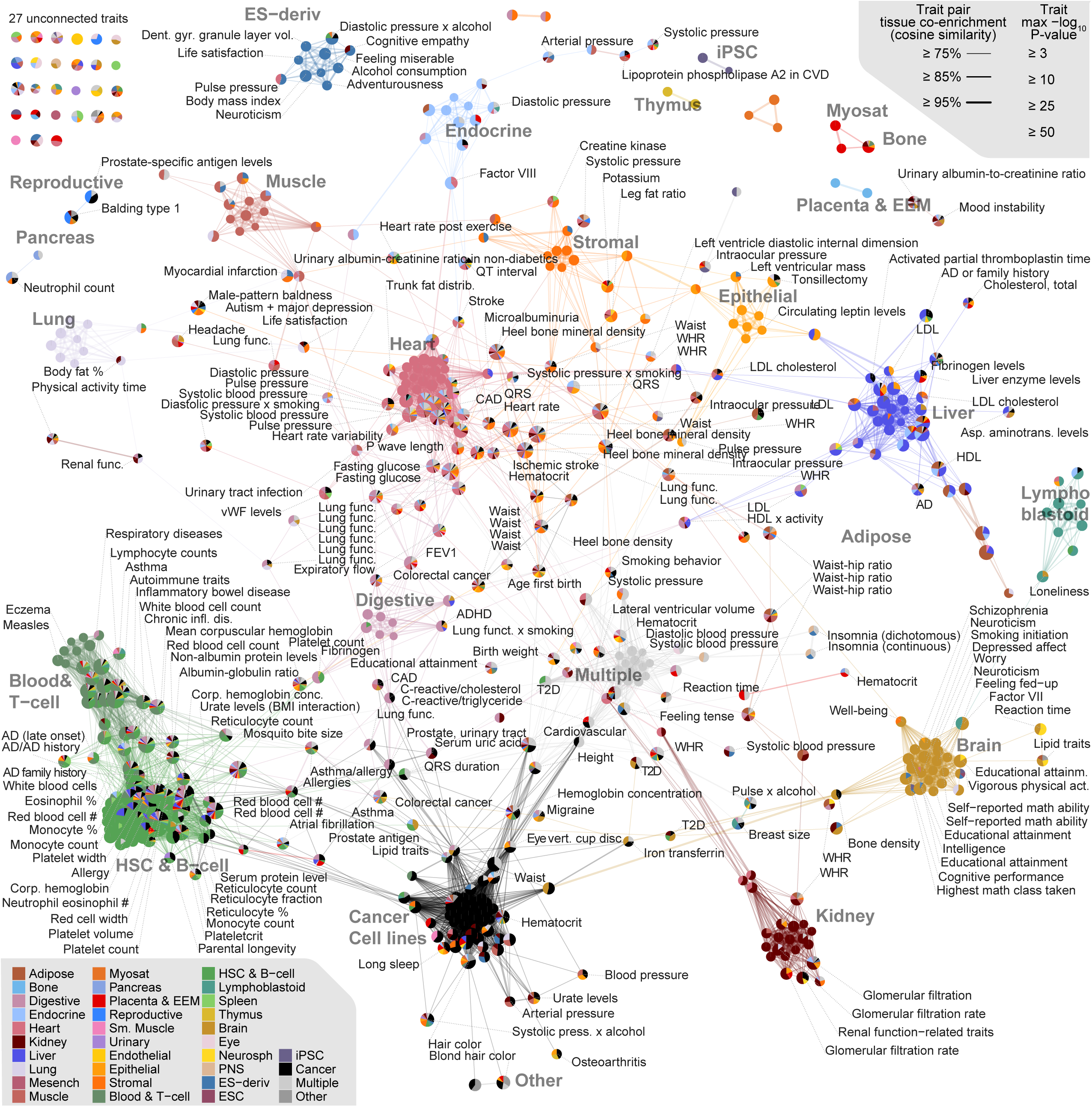
Trait-trait network across 538 traits by similarity of epigenetic enrichments (cosine sim. >= 0.75), laid out using the Fruchterman-Reingold algorithm. Traits (nodes) are colored by contributing groups (pie chart by fraction of −log_10_p, size by maximal −log_10_p) and interactions (edges) by the group with maximal dot product of enrichments between two traits.

Multifactorial traits connected these communities, including: CAD linking heart, endocrine, and liver; HDL and triglycerides levels linking liver and adipose; lung function-related traits linking lung, heart, and digestive tissues; blood pressure measurements surrounding heart and linking to endocrine, endothelial, and liver; cell count fraction traits implicated principally blood often partnered with liver, digestive, and other tissues. Polyfactorial traits (e.g. waist-to-hip ratio and heel bone mineral density) were centrally located linking diverse categories.

We found that this trait-trait co-enrichment network captures many biologically-meaningful relationships that are missed by genetic information alone (**Supp Fig. S32, S33, S34**). A genetic overlap network connecting any two traits that share even 5% of their loci (Jaccard index, 10kb resolution) results in only 934 edges, and only captures 5% of the edges captured by our epigenomics-centered analysis (N=283 of 5547), highlighting the importance of our epigenomics lens in capturing the common biological basis of complex traits.

## Discussion

In this work, we presented EpiMap, the most complete and comprehensive map of the human epigenome, encompassing ~15k uniformly-processed, QC-metric pruned, and computationally-completed datasets. Our resource encompasses 833 distinct biological samples, each with 18 epigenomic marks, painting a rich epigenomic landscape. For each reference epigenome, we generate a chromatin state annotation based on multiple chromatin marks and distinguish diverse classes of enhancer, promoter, transcribed, repressed, repetitive, heterochromatic, and quiescent states. We also provide a high-resolution enhancer annotation track, that combines multiple active enhancer states and DNase-accessible regions, covering only 0.8% of each reference epigenome, and collectively 13% of the genome across all 833 tissues.

EpiMap greatly expands the biological space covered by previous reference epigenome maps, by incorporating samples of the endocrine system, placenta, extraembryonic membranes, reproductive system, stromal and endothelial primary cells, and a large collection of widely-used cancer cell lines. We also greatly increase the coverage of embryonic and adult brain, heart, muscle, kidney, lung, and liver tissues, as well as blood, immune, lymphoblastoid, and epithelial cells. This broader biological space has important implications both in capturing gene-regulatory elements and upstream regulators of an increased set of tissue-specific biological pathways, and in annotating gene-regulatory variants across a broader biological spectrum that now capture many more traits and disease phenotypes that were previously uncaptured.

The integration and uniform processing of these samples enabled us to paint a detailed picture of biological sample relationships, both outlining the relationships between our 33 broad tissue categories, and detailing the relationships of individual samples within these categories. For example, we found that lifestage played an important role in establishing high-level organization of sample similarities, that primary cells clustered separately from their tissue of origin, that adult samples separated by tissue but embryonic samples clustered together. We also recognized differences in the epigenomic relationships painted by different marks, with repressive marks better distinguishing developmental stages, and active marks better distinguishing different tissues and lineages. These relationships can help guide the prioritization of new biological samples to profile for gene-regulatory or disease studies, by studying gene-regulatory motif enrichments, downstream-gene pathway enrichments, and GWAS trait enrichments in the context of the coverage of biological space by closely- and distantly-related samples.

Our analysis enabled us to elucidate important gene-regulatory relationships between enhancers, their gene-regulatory targets, and their upstream regulators. By recognizing modules of common enhancer activity, we partitioned 2.1 million gene-regulatory elements into 300 co-regulated sets, distinguishing broadly-active enhancer modules vs. tissue-specific modules, and their distinct gene-regulatory circuitry and biological roles. Gene-regulatory sequence motifs enriched in modules of common activity patterns enabled us to recognize upstream regulators of these modules, including many newly-profiled tissues such as TEAD3 in placenta and epithelial cells and NFIB in various embryonic organs, and distinguishing tissue-specific vs. promiscuous gene-regulatory motifs such as RFX1-5, GRHL1, HNF1A/B, and AP-1. Similarly, common gene ontology enrichments of genes proximal to these modules enabled us to pinpoint the common biological pathways they likely control in tissue-specific, lineage-specific, and broadly-active biological roles.

Our work also provided the most comprehensive analysis to date of the gene-regulatory underpinnings of complex traits and human disease. We found statistically-significant enrichments between 534 traits and all 833 tissues, shedding light on 30,247 loci containing SNPs within enriched annotations, and thus providing meaningful insights into their potential mechanisms of action. These enrichments helped distinguish unifactorial, multifactorial, and polyfactorial traits, based on the number of distinct tissue types they implicate, and revealed principal vs. partner tissues that play likely driver vs. auxiliary roles across traits. The multiple enriched tissues in multifactorial traits allowed us to dissect their complexity, by partitioning SNPs by tissues, which showed distinct gene pathway enrichments and shared genetic risks with different related traits. Finally, we used these tissue-trait enrichment and co-enrichment patterns to reveal the shared biological basis of hundreds of complex traits through the lens of their enriched tissues, providing an important basis for studying the common and distinct components of disease comorbidity relationships.

We expect EpiMap to enable many new methodological developments in the study of gene regulation and disease. For example, the much more densely-populated tree of samples enabled us to develop a new hierarchical approach to GWAS trait-tissue enrichment analysis that directly compares the enrichment of parent-child tree node pairs, thus finding the appropriate level of tissue specificity where genetic traits may act. The approach itself is general, and likely also applicable to hierarchical analyses of motif and gene pathway enrichments across the tree. We expect that many additional multi-resolution approaches will be applicable to these datasets, enabling us to recognize variability between and within group rigorously and systematically. Similarly, the large number of traits and tissues enabled us to infer networks of tissue-trait, trait-trait, and tissue-tissue relationships, and we expect that many additional graph algorithms and analysis approaches will be applicable to the study of these networks.

EpiMap also has several limitations that we hope will be overcome with continued technological improvements. First, outside cancer lines and isolated primary cells, our tissue samples are primarily from bulk dissections, which combine multiple underlying cell types, thus hiding the individual contributions and distinct biology of each cell type, and sometimes missing altogether the contribution of lower-abundance cell types. As single-cell ATAC-seq^55^ and single-cell ChIP-seq^56,57^ approaches mature, we expect that new cell-type-specific maps will become possible, enabling us to partition the enhancers and chromatin states discovered here into their constituent cell types, and also to discover new enhancers and gene-regulatory elements from lower-abundance cell types that are currently not detectable in bulk samples. The presence of such single-cell maps should also allow systematic methods for epigenomic deconvolution^58^. In addition, our current approach for uniformly processing and imputing missing marks does not take into consideration the genotype of different individuals, which contains important information for inferring the activity pattern of genes or gene-regulatory regions in a new individual based on their genotype, but also their phenotypic variables. Lastly, while EpiMap represents a substantial increase relative to previous maps, we are still missing many tissues, environmental stimulation conditions and developmental stages that may active enhancers and other gene-regulatory elements that may not be currently visible in our compendium.

## Acknowledgements

We thank the ENCODE, Roadmap, and GGR consortia for generating high-quality public datasets and rapidly disseminating their results to the broader community. We thank Daofeng Li and Ting Wang, and Idan Gabdank and J. Seth Strattan for making our observed and imputed genome-wide tracks and chromatin state annotations available through the WashU Epigenome Browser and the ENCODE portal respectively. We thank Jason Ernst for advice, guidance, and for developing the ChromImpute methodology and code base. We thank Pouya Kheradpour for help with motif enrichment analysis software. We thank Luke Ward for incorporating our annotations in HaploReg. We thank the Charles Epstein, Xihong Lin, Hufeng Zhou, Corbin Quick, Jacob Schreiber, Bill Noble, Martin Hirst, Zhiping Weng, Mark Gerstein, ENCODE, IHEC, Roadmap, GENCODE, GTEx, GSP, and TopMed consortia for feedback on early versions of this work. We thank Irwin Jungreis, Lei Hou, Leandro Agudelo, Shahin Mohammadi, Xinchen Wang, Azim Amirabad, Max Wolf, Alvin Shi, Khoi Nguyen for feedback on the work and the resource. This work was supported by National Institutes of Health grants HG008155, HG009446, HG009088, HG007234, HG007610, GM113708, MH109978, MH119509, and AG058002 (M.K.) and NIH training grant GM087237 (C.B.).

## Methods

### Epigenomic datasets and processing

#### Primary data sources and metadata information

We analyzed 3,030 datasets, including 2329 epigenomic ChIP-seq datasets, 635 DNase-seq datasets, and 66 ATAC-seq datasets from ENCODE at https://www.encodeproject.org/, released as of Sept. 24th 2018. These marks include: Tier 1 assays: DNase-seq, H3K4me1, H3K4me3, H3K27ac, H3K36me3, H3K9me3, and H3K27me3; Tier 2 assays: ATAC−seq, H3K9ac, H3K4me2, H2AFZ, H3K79me2, and H4K20me1; Tier 3 assays: POLR2A, EP300, CTCF, SMC3, RAD21; and Tier 4 histone marks: 16 non-imputed histone acetylation marks, 4 methylation marks (H3K9me2, H3K79me1, H3K9me1, H3K23me2), H3F3A, and H3T11ph. We assigned unique sample IDs to each unique combination of: extended biosample summary, donor, sex, age, and lifestage, wherever each attribute was available. We removed samples with genetic perturbations, and kept only samples with appropriately matched ChIP-seq controls. We provide a metadata matrix including the mapping between ENCODE accessions and our unique sample IDs (Supp. Table S1, also at compbio.mit.edu/epimap). We mapped the 111 Roadmap epigenomes and 16 ENCODE 2012 epigenomes to any of our samples with overlapping dataset accessions if the accessions were used in the flagship Roadmap epigenomics analysis. This mapping assigned 25 samples to ENCODE 2012 and 184 samples to Roadmap 2015, some of which were merged multi-donor samples in Roadmap, out of the final 833 samples that passed QC. These were merged into 16 and 111 tissue types respectively in the Roadmap 2015 publication.

#### Uniform data processing

We downloaded one alignment file per replicate, prioritizing filtered alignments in hg19 whenever possible. We uniformly processed ChIP-seq and DNase-seq datasets according to the processing pipelines established by the Roadmap Epigenomics Consortium^8^. Briefly, we filtered out improperly paired and non-uniquely mapped reads, truncated reads to 36 base pairs, filtered out a blacklist of low complexity and artifact regions (ENCODE accession ENCSR636HFF), and filtered reads against a mappability track of uniquely mappable regions for 36bp reads^60^. We converted bam files to tagAlign, used liftOver^61^ to map GRCh38 alignments to hg19, and pooled all experiments within each ID and assay combination. We subsampled pooled ChIP-seq data sets to a maximum of 30 million reads and DNase-seq and ATAC-seq data sets to a maximum of 50 million reads. We used SPP to estimate fragment length. In cases with extremely low fragment length in ATAC-seq and DNase-seq datasets we used the average fragment length (73) from the average of the rest of the tracks. We generated −log10 p-value signal tracks against matched whole cell extract (WCE) for both ChIP-seq and accessibility data sets using the MACS2^62^ and the SPP^63^ peak caller and cross correlation analysis to identify the proper fragment length as in the Roadmap analysis.

### Epigenomic Imputation

#### Imputation

We carried out epigenomic imputation on 859 unique cell types using ChromImpute^25^ for a total of 10,778 imputed datasets over thirteen Tier 1 and Tier 2 assays using predictors trained on all 35 epigenomic assays across 859 samples. We additionally imputed 4,345 datasets for the five DNA associated factors, using only the 35 epigenomic assays as features to train predictors with ChromImpute. We provide all imputed and processed observed tracks along with tracksets for the 833 QCed samples at https://epigenome.wustl.edu/epimap^29^.

#### Quality control

For imputation quality control (QC) and validation, we compare observed tracks to imputed tracks when both were available (i.e. when at least two original observed datasets were available for that cell type). We calculated all imputation QC metrics from the original ChromImpute publication^25^, including genome-wide correlation, % imputed and observed peak recovery, and AUC for all pairs of imputed and observed tracks. In addition to quantitative metrics, we visually inspected epigenomic predictions as part of our quality control. We show (Fig. 1B) three dense and varied regions of different resolutions (25kb, 200kb, 1.5Mb) for each of two randomly chosen samples containing both observed and imputed tracks for each assay. We calculated epigenomic profile quality metrics NSC, RSC, and read depth for all datasets and compared these to the imputation QC metrics (tables in Supp. Table S1). We flagged low-quality tracks by detecting the elbow in the ranked correlation metrics, which we calculated as the point where the change in correlation exceeded 5% of the correlation.

#### Sample and antibody swap detection

To systematically identify both potential sample or antibody (Ab) swaps and poor quality experiments, we computed the correlation of each observed experiment against all 10,734 imputed tracks for histone marks and assays (all imputed tracks before removing samples by QC). We then calculated the average correlation among the top 10 most similar tracks to each observed track. We flagged potential Ab swaps by comparing the average correlation against samples of the putative mark against those computed for other marks. We fit an OLS model to each mark comparison, flagged datasets with residuals greater than 3 standard deviations of the average correlation, and visually confirmed 7 Ab swaps (6 low-quality tracks). Similarly, we flagged potential sample swaps by comparing the correlation between imputed and observed tracks against the average correlation in the top 10 tracks in the same mark. We fit an OLS model and flagged datasets with residuals greater than 3 standard deviations of the residuals distribution. We report 19 potentially swapped samples, of which 5 were also flagged as low-quality tracks (**Supp. Fig. S8**).

#### Secondary reactivities

In addition to genome-wide QC of imputed tracks, we also focused on the specific differences between observed and imputed tracks. For each observed mark, we generated a genome-wide ‘delta’ track, computed as the difference in signal intensity between observed and imputed data, re-scaling imputed tracks to match signal intensity properties of the observed tracks, as observed tracks showed a general bias for higher intensity. Some of these ‘delta’ tracks showed surprisingly high correlations with ‘primary’ tracks of non-putative marks, indicating potential secondary antibody reactivities. In order to flag these reactivities, we compared the average correlation of each of the delta tracks to the top 10 closest imputed tracks for each mark. As with antibody swaps, we fit an OLS model in each mark combination to flag outliers. We flagged 19 tracks and report 15 after visual inspection as potential secondary reactivities or single replicate swaps (e.g. in the case of DNase-seq) (**Supp. Fig. S7 and S8**). We noted that some cases showed clear difference tracks that don’t match available antibodies, suggesting that the secondary reactivity is not a common mark in our compendium.

#### Biological space coverage

To evaluate the similarity of imputed and observed tracks across samples, we calculated the pairwise genomic correlations between all pairs of imputed and observed signal tracks. We hierarchically clustered each individual mark’s imputed or observed correlation matrix using Ward’s method. We averaged all imputed matrices for the six main marks (H3K27ac, H3K4me1, H3K4me3, H3K36me3, H3K27me3, H3K9me3) to create a fused correlation matrix, which we similarly clustered. We plotted the hierarchically clustered tree for the fused matrix alongside the metadata information for each epigenome using the circlize R package^64^.

Additionally, we calculated mark-specific spearman correlations restricted to relevant features within all observed and imputed tracks per mark. We mapped each of 13 marks to its top state by emission probability in the ChromHMM 25 state model and any other states with emission probability over 80%. For ATAC-seq, we use the same region list as DNase-seq. For each mark, we averaged and reduced each 25bp signal track to any 200bp regions that were labeled as one of the states associated with the mark in any of the 127 imputed Roadmap epigenomes under the 25-state model^8,25^. We calculated the spearman correlation between sets of these region-restricted mark signal tracks and generate similarity matrices across all datasets for a mark. Using these spearman correlation matrices on all observed and imputed signal tracks, we computed UMAP dimensionality reductions for each mark and assay using with the uwot R package^65^ with the default parameters except for n_neighbors=250, min_dist=0.25, and repulsion_strength=0.25.

### Epigenomic annotations

#### Chromatin state annotations

We computed epigenomic annotations on 3,533 imputed and 1,465 observed datasets for six marks on 833 samples using ChromHMM with the fixed 18-state model from Roadmap^8^ with the same mnemonics and colors. We use observed data wherever possible, except in cases with no observed data or where observed data was removed in QC. The table of signal tracks used to calculate annotations is available as Supplementary Table S2. Observed data was binarized from signal tracks with a −log10 p-value signal cutoff of 2. In order to binarize imputed data and facilitate the comparison with observed data, we established mark-specific binarization cutoffs. We first separately calculated the overall probability distributions of all imputed and observed tracks for each mark. Then for each mark, we set the imputed binarization cutoff value to the value of the quantile matching the quantile in observed data for the −log10 p-value > 2 cutoff. We used liftOver^61^ to map all 833 (after QC) ChromHMM annotations to GRCh38 and provide these alongside hg19 annotations and binarized imputed and observed datasets and as tracksets at https://epigenome.wustl.edu/epimap.

#### Defining active enhancers

We define active enhancers as the intersection of DHS (DNase I Hypersensitive Sites) regions with enhancer annotations and high H3K27ac signal (average signal > 2 in the region containing the DHS +/− 100bp). We define DHS regions from an index list of 3,568,912 DHS consensus locations determined from 733 DNase-seq experiments. We intersect these regions with the 833 imputed enhancer annotations (states 7,8,9,10,11, and 15 in the 18-state model). This results in 2,842,995 regions with at least one enhancer annotation in any epigenome. Finally, we intersect this matrix with the H3K27ac signal in the +/−100bp region encompassing each DHS from the same tissue-specific imputed and observed datasets used to calculate the ChromHMM annotations. This procedure results in 2,356,914 active enhancer regions. We created an equivalent promoter element region using the promoter annotations (states 1,2,3,4, and 14 in the 18-state model). We noticed that a number of regions share both enhancer and promoter annotations. As a conservative cutoff, we assign all regions to either enhancers or promoters if 75% or more of its active occurrences are labeled as that type of element (**Supp. Fig. S18**). This final thresholding procedure yields 2,069,090 enhancers, 204,104 promoters, and 122,358 dyadic elements (neither specifically promoter or enhancer). Matrices and enhancer locations are available through compbio.mit.edu/epimap.

For all images using tissue group order, including ChromHMM tracks and modules heatmaps, groups are ordered alphabetically within six major groups: Tissue/Organs (Adipose, Bone, Digestive, Endocrine, Heart, Kidney, Liver, Lung, Mesench, Muscle, Myosat, Pancreas, Placenta & EEM, Reproductive, Sm. Muscle, and Urinary), Other Primary Cells (Endothelial, Epithelial, and Stromal), Blood + Immune (Blood & T-cell, HSC & B-cell, Lymphoblastoid, Spleen, and Thymus), Nervous system (Brain, Eye, Neurosph, and PNS), Stem (ES-deriv, ESC, iPSC), and Other (Cancer, Other).

#### Defining enhancer modules

In order to define enhancer modules, we clustered the binary enhancer matrix defined by intersecting enhancer annotations with DHS regions and with average centered and flanking (+/− 100 bp) H3K27ac signal above a −log10 pval of 2 using the k-centroids algorithm with the Jaccard distance with the number of clusters set to k=300. The average module contains 6,897 enhancers, and the largest module (enumerating constitutive elements) contains 93,554 enhancer regions. In all heatmap plots of module centers (and associated enrichment figures), we diagonalize the matrix by ordering each column in the heatmap (module centers) by the epigenome which contributes the maximal signal. All columns which have signal over 25% in more than 50% of rows are shown first. We use this diagonalization procedure for all diagonalized heatmaps. We colored each module by the tissue group which contains its maximal signal. Modules highlight sample groupings and organize according to cell type and tissue. Major groups are ordered alphabetically within six major groups and samples are ordered within groups according to Ward method’s clustering of the Jaccard distance of of the modules centers matrix. We performed enrichment on the module centers against the metadata of included samples (signal over 25%) by the hypergeometric test, and show enrichments with −log_10_p > 2.

#### Gene ontology enrichment

We performed gene ontology (GO) enrichments on each enhancer module using GREAT v3.0.0 for the Biological Process, Cellular Component, and Molecular Function ontologies^31^. We analyzed and visualized the results in the same manner as in the Roadmap core paper^8^. We only considered enrichments of 2 or greater with a multiple testing-corrected p-value less than 0.01. For Figure 4C, we reduced the enrichments by modules matrix to terms with a maximal −log_10_p-value > 4 that were enriched in less than 10% of modules. The full enrichment matrix is shown as **Supp. Figure S27**. As in the case of the diagonalized module centers, we labeled each term according to the module containing its maximal signal. We used a bag of words approach (as described in Roadmap^8^) to pick 72 representative terms out of 865 total terms for **Figure 3B** such that each tissue group has at least one term and the rest are representatively allocated across groups.

#### Motif enrichment

We performed motif enrichment analysis across enhancer modules as described in the Roadmap paper. Briefly, we measured enrichments for 1,772 motifs against a background of all enhancers. We report the motif with the highest enrichment in any module for each of 300 previously identified motif clusters^8,66^. We only report motifs with a maximal log_2_ enrichment ratio of at least 1.5, resulting in 86 motifs, which we show with their PWM logos against all 300 modules as **Figure 3C**.

### GWAS enrichment analysis

We pruned the NHGRI-EBI GWAS catalog^1^ (downloaded from https://www.ebi.ac.uk/gwas/docs/file-downloads on May 3rd, 2019) using a greedy approach: within each trait + PMID combination, we ranked associations by their significance (p-value), and added SNPs iteratively if they were not within 5kb of previously added SNPs. We also removed all associations in the HLA locus (for hg19: chr6:29,691,116-33,054,976). This reduced the catalog from 121k to 113k associations. Finally, we reduced the catalog to 926 unique GWAS (from 5454 GWAS) with an initial sample size of at least 20,000 cases or individuals (wherever cases and controls were not annotated). This resulted in 66,801 lead SNPs, which landed in 33,417 unique genome intervals when we split the genome into 10,000bp intervals.

#### Flat epigenome enrichments and module epigenome enrichments

We performed the hyper-geometric test to evaluate GWAS enrichments on full epigenomes and on modules. For these flat enrichments, we compare each number of SNP-enhancer intersections for each enhancer set (epigenome or module) to the full set of intersections in all M enhancers. As above, we correct for multiple testing for each GWAS and enhancer set combination by computing and correcting with null association p-values for epigenomes and modules using the null catalogs generated for the tree enrichment. Rarefaction curves were calculated on the epigenome enrichments by iteratively adding the sample that was either (**Fig. 4D**) significantly enriched or (**Supp. Fig. S29**) the maximal enrichment for the most remaining GWAS until all GWAS were accounted for.

#### Tree epigenome enrichments

We constructed a tree by hierarchically clustering the Jaccard similarity of the binary enhancer by epigenomes matrix using complete-linkage clustering. Then, for each node in the tree, we calculated its consensus epigenomic set, defined as the set of all enhancers present in all leaves of the subtree, such that each node’s set was a superset of that of its parent. For each GWAS, we asked whether the novel consensus enhancers at a node were significantly enriched for lead SNPs by comparing the enrichment between each node and its parent as measured by the likelihood ratio test between two logistic regressions.

Briefly, for each GWAS catalog unique trait and PubMedID, we find all intersections of its pruned SNPs with M=2,069,090 enhancers. Then **Y** is an indicator vector of size M which shows the intersected enhancers. We find all consensus enhancers (intersection of epigenomes in the subtree) in the node of interest (vector **X**_**N**_) and in its parent (**X**_**P**_). All vectors are 1×M. We calculate **X**_**D**_ = **X**_**N**_ − **X**_**P**_(specific enhancers), which is also in {0,1}^(1×M)^ as each node contains a superset of its parent’s enhancers.

We then calculate following two logistic regressions:

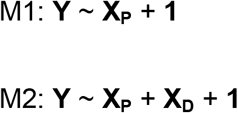

We calculated the log-likelihood difference and apply the likelihood ratio test to test whether adding the specific enhancers (M2) is significantly different from the parent model (M1). To correct for multiple testing on a per GWAS and node basis, we generated 1000 null GWAS catalogs by shuffling the associations across GWAS. We used these catalogs to compute the null association p-values for each permuted GWAS and used the 1st, 5th, and 10th smallest p-values for each GWAS and node combination as their 0.1%, 0.5%, and 1% FDR cutoff.

For the CAD example, GO terms^67^ were calculated using the nearest gene of each enhancer hit by a lead SNP. We pruned genes to expressed genes by calculating average RNA-seq profiles for each tissue group and excluded genes that had log_2_FPKM of less than 2 in the average RNA-seq of each sample’s group. 341 of 833 samples have matched RNA-seq, which we list in addition to releasing the processed data at compbio.mit.edu/epimap. We kept only the GO terms that were significant in 25% or less of nodes, report the top 2 GO terms per node in **Figure 6C**, and all GO terms in **Supp. Fig. S37**.

#### Tissue similarity

We assigned each internal node in the tree to a unique tissue if its subtree’s leaves are more than 50% of that tissue and as “Multiple” if the subtree is not majority one tissue. We assigned tissue labels to 641 of 832 (77%) internal nodes where the majority of leaves corresponded to a single group. Using these assignments, we created a tissue by GWAS matrix by adding the −log10 p-values for each tissue node set from all of the GWAS enrichments on the tree. We binarized this matrix and computed the jaccard similarity across tissues to calculate a tissue similarity matrix. To assess significance of tissue overlap, we compared each overlap value against the overlaps from 10,000 permuted enrichments. We collapsed each permuted matrix into a tissue by GWAS matrix to compute the overlaps under the null. We performed the permutations for each tissue against other tissues by shuffling the enrichment p-values on the node by GWAS matrix. Specifically, we (a) binarized the enrichment matrix, (b) fixed the column of the group of interest and (c) permuted the remainder of the matrix keeping its row and column marginals the same and then (d) calculated the cosine distance between the permuted and the original matrix of enrichments.

#### Cross-GWAS Network

To evaluate the cross-GWAS similarity, we normalized the tissue by GWAS matrix for each GWAS to obtain the proportion of significance attributed to each tissue for each GWAS (**Supp. Fig S38**). We reduced the matrix to 538 significant GWAS with at least 20,000 cases (or individuals when no cases were specified). We created a GWAS-GWAS network using the cosine distance matrix as an adjacency matrix, keeping 5,547 links with a cosine distance of 0.25 or less. We used the Fruchterman-Reingold algorithm to lay out the graph^68^. We use the tissue by GWAS matrix to color links according to the maximum tissue in the product between each pair of nodes and to color nodes according to the maximal tissue for each node (**Supp Fig. S39**).

In order to compare the epigenetic network to trait genetic similarity, we binned snps in the GWAS catalog into 10kb windows starting from the beginning of each chromosome. We counted the number of intersecting bins between two traits and keep any trait pairs with jaccard similarity of at least 1%. To compare this to the epigenetic network, we plotted only links in the epigenetic network that coincide with any SNP-sharing GWAS pairs. Additionally, we plotted the heatmaps of the tree enrichments distance matrix and the genetic similarity matrix side by side, first organized by hierarchically clustering the enrichments matrix and then by clustering the genetic similarity matrix (**Supp. Fig. S41 and S42**).

