## Supplemental Figures for "Integrative analysis of 10,000 epigenomic maps across 800 samples for regulatory genomics and disease dissection"

| | Group | BSSID | Extended Info |
|---|---|---|---|
| 1 | Adipose | BSS00038 | ADIPOCYTE |
| 2 | Adipose | BSS00043 | ADIPOSE TISSUE |
| 3 | Adipose | BSS01671 | ADIPOSE TISSUE |
| 4 | Adipose | BSS01665 | ADIPOSE TISSUE |
| 5 | Adipose | BSS01666 | ADIPOSE TISSUE |
| 6 | Adipose | BSS01668 | ADIPOSE TISSUE |
| 7 | Adipose | BSS01669 | ADIPOSE TISSUE |
| 8 | Adipose | BSS01667 | ADIPOSE TISSUE |
| 9 | Adipose | BSS01394 | OMENTAL FAT PAD |
| 10 | Adipose | BSS01393 | OMENTAL FAT PAD |
| 11 | Blood & T-cell | BSS00188 | CD4 T CELL |
| 12 | Blood & T-cell | BSS00185 | CD4 T CELL |
| 13 | Blood & T-cell | BSS00189 | CD4 T CELL |
| 14 | Blood & T-cell | BSS00190 | CD4 T CELL |
| 15 | Blood & T-cell | BSS00183 | CD4 T CELL |
| 16 | Blood & T-cell | BSS00186 | CD4 T CELL |
| 17 | Blood & T-cell | BSS00191 | CD4 T CELL |
| 18 | Blood & T-cell | BSS00192 | CD4 T CELL |
| 19 | Blood & T-cell | BSS00274 | CD4 T CELL |
| 20 | Blood & T-cell | BSS00195 | CD8 T CELL |
| 21 | Blood & T-cell | BSS00196 | CD8 T CELL |
| 22 | Blood & T-cell | BSS00198 | CD8 T CELL |
| 23 | Blood & T-cell | BSS00200 | CD8 T CELL |
| 24 | Blood & T-cell | BSS00193 | CD8 T CELL |
| 25 | Blood & T-cell | BSS00194 | CD8 T CELL |
| 26 | Blood & T-cell | BSS00197 | CD8 T CELL |
| 27 | Blood & T-cell | BSS01420 | MONONUCLEAR CELL |
| 28 | Blood & T-cell | BSS01421 | MONONUCLEAR CELL |
| 29 | Blood & T-cell | BSS01279 | MONONUCLEAR CELL |
| 30 | Blood & T-cell | BSS01419 | MONONUCLEAR CELL |
| 31 | Blood & T-cell | BSS01423 | MONONUCLEAR CELL |
| 32 | Blood & T-cell | BSS01424 | MONONUCLEAR CELL |
| 33 | Blood & T-cell | BSS01347 | NAIVE T CELL |
| 34 | Blood & T-cell | BSS01348 | NAIVE T CELL |
| 35 | Blood & T-cell | BSS01346 | NAIVE T CELL |
| 36 | Blood & T-cell | BSS01688 | T CELL |
| 37 | Blood & T-cell | BSS01687 | T CELL |
| 38 | Blood & T-cell | BSS01689 | T CELL |
| 39 | Blood & T-cell | BSS01684 | T CELL |
| 40 | Blood & T-cell | BSS01691 | T1 CELL |
| 41 | Blood & T-cell | BSS01692 | T1 CELL |
| 42 | Blood & T-cell | BSS01690 | T1 CELL |
| 43 | Blood & T-cell | BSS01693 | T17 CELL |
| 44 | Blood & T-cell | BSS01694 | T17 CELL |
| 45 | Blood & T-cell | BSS01695 | T17 CELL |
| 46 | Blood & T-cell | BSS01697 | T2 CELL |
| 47 | Blood & T-cell | BSS01698 | T2 CELL |
| 48 | Blood & T-cell | BSS01696 | T2 CELL |
| 49 | Blood & T-cell | BSS01478 | TREG CELL |
| 50 | Blood & T-cell | BSS01479 | TREG CELL |
| 51 | Blood & T-cell | BSS01480 | TREG CELL |
| 52 | Bone | BSS00084 | BONE ARM |
| 53 | Bone | BSS00330 | BONE FEMUR |
| 54 | Bone | BSS01154 | BONE LEG |
| 55 | Bone | BSS00705 | BONE MARROW STROMA |
| 56 | Bone | BSS01397 | OSTEOBLAST |
| 57 | Brain | BSS00071 | AMMONS HORN |
| 58 | Brain | BSS00077 | ANGULAR GYRUS |
| 59 | Brain | BSS00078 | ANGULAR GYRUS |
| 60 | Brain | BSS00089 | ASTROCYTE |
| 61 | Brain | BSS00090 | ASTROCYTE CEREBELLUM |
| 62 | Brain | BSS00091 | ASTROCYTE HIPPOCAMPUS |
| 63 | Brain | BSS00092 | ASTROCYTE SPINAL CORD |
| 64 | Brain | BSS00135 | BRAIN |
| 65 | Brain | BSS00136 | BRAIN |
| 66 | Brain | BSS00129 | BRAIN |
| 67 | Brain | BSS00130 | BRAIN |
| 68 | Brain | BSS00131 | BRAIN |
| 69 | Brain | BSS00133 | BRAIN |
| 70 | Brain | BSS00138 | BRAIN |
| 71 | Brain | BSS00139 | BRAIN |
| 72 | Brain | BSS00140 | BRAIN |
| 73 | Brain | BSS00142 | BRAIN |
| 74 | Brain | BSS00126 | BRAIN |
| 75 | Brain | BSS00127 | BRAIN |
| 76 | Brain | BSS00125 | BRAIN |
| 77 | Brain | BSS00132 | BRAIN |
| 78 | Brain | BSS00134 | BRAIN |
| 79 | Brain | BSS00141 | BRAIN |
| 80 | Brain | BSS00174 | CAUDATE NUCLEUS |
| 81 | Brain | BSS00175 | CAUDATE NUCLEUS |
| 82 | Brain | BSS00173 | CAUDATE NUCLEUS |
| 83 | Brain | BSS00201 | CEREBELLAR CORTEX |
| 84 | Brain | BSS00205 | CEREBELLUM |
| 85 | Brain | BSS00207 | CEREBELLUM |
| 288 | Digestive | BSS01654 | STOMACH |
| 289 | Digestive | BSS01651 | STOMACH |
| 290 | Digestive | BSS01639 | STOMACH |
| 291 | Digestive | BSS01284 | STOMACH MUCOSA |
| 292 | Digestive | BSS01848 | TRANSVERSE COLON |
| 293 | Digestive | BSS01851 | TRANSVERSE COLON |
| 294 | Digestive | BSS01849 | TRANSVERSE COLON |
| 295 | Digestive | BSS01850 | TRANSVERSE COLON |
| 296 | Endocrine | BSS00282 | ENDOCRINE PANCREAS |
| 297 | Endocrine | BSS00283 | ENDOCRINE PANCREAS |
| 298 | Endocrine | BSS00284 | ENDOCRINE PANCREAS |
| 299 | Endocrine | BSS00281 | ENDOCRINE PANCREAS |
| 300 | Endocrine | BSS01403 | OVARY |
| 301 | Endocrine | BSS01401 | OVARY |
| 302 | Endocrine | BSS01402 | OVARY |
| 303 | Endocrine | BSS01399 | OVARY |
| 304 | Endocrine | BSS01719 | TESTIS |
| 305 | Endocrine | BSS01718 | TESTIS |
| 306 | Endocrine | BSS01715 | TESTIS |
| 307 | Endocrine | BSS00052 | ADRENAL GLAND |
| 308 | Endocrine | BSS00050 | ADRENAL GLAND |
| 309 | Endocrine | BSS00051 | ADRENAL GLAND |
| 310 | Endocrine | BSS00059 | ADRENAL GLAND |
| 311 | Endocrine | BSS00057 | ADRENAL GLAND |
| 312 | Endocrine | BSS00058 | ADRENAL GLAND |
| 313 | Endocrine | BSS00045 | ADRENAL GLAND |
| 314 | Endocrine | BSS00046 | ADRENAL GLAND |
| 315 | Endocrine | BSS00047 | ADRENAL GLAND |
| 316 | Endocrine | BSS00048 | ADRENAL GLAND |
| 317 | Endocrine | BSS00054 | ADRENAL GLAND |
| 318 | Endocrine | BSS00060 | ADRENAL GLAND |
| 319 | Endocrine | BSS00055 | ADRENAL GLAND |
| 320 | Endocrine | BSS00056 | ADRENAL GLAND |
| 321 | Endocrine | BSS01831 | THYROID GLAND |
| 322 | Endocrine | BSS01834 | THYROID GLAND |
| 323 | Endocrine | BSS01832 | THYROID GLAND |
| 324 | Endocrine | BSS01835 | THYROID GLAND |
| 325 | Endothelial | BSS00143 | BRAIN MICROVASCULAR ENDOTHELIAL CELL |
| 326 | Endothelial | BSS00387 | GLOMERULUS ENDOTHELIAL CELL |
| 327 | Endothelial | BSS01077 | KIDNEY CAPILLARY ENDOTHELIAL CELL |
| 328 | Endothelial | BSS01206 | LUNG MICROVASCULAR ENDOTHELIAL CELL |
| 329 | Endothelial | BSS01465 | PULMONARY ARTERY ENDOTHELIAL CELL |
| 330 | Endothelial | BSS00298 | UMBILICAL VEIN ENDOTHELIAL CELL |
| 331 | Endothelial | BSS00296 | UMBILICAL VEIN ENDOTHELIAL CELL |
| 332 | Endothelial | BSS00258 | DERMIS BLOOD VESSEL ENDOTHELIAL CELL |
| 333 | Endothelial | BSS00260 | DERMIS BLOOD VESSEL ENDOTHELIAL CELL |
| 334 | Endothelial | BSS00262 | DERMIS LYMPHATIC VESSEL ENDOTHELIAL CELL |
| 335 | Endothelial | BSS00264 | DERMIS LYMPHATIC VESSEL ENDOTHELIAL CELL |
| 336 | Epithelial | BSS00704 | BONE MARROW EPITHELIAL CELL |
| 337 | Epithelial | BSS00218 | CHOROID PLEXUS EPITHELIAL CELL |
| 338 | Epithelial | BSS00223 | COLON EPITHELIAL CELL |
| 339 | Epithelial | BSS00307 | ESOPHAGUS EPITHELIAL CELL |
| 340 | Epithelial | BSS00743 | IRIS PIGMENT EPITHELIAL CELL |
| 341 | Epithelial | BSS01385 | NON-PIGMENTED CILIARY EPITHELIAL CELL |
| 342 | Epithelial | BSS01092 | GLOMERULUS EPITHELIAL CELL |
| 343 | Epithelial | BSS00389 | GLOMERULUS VISCERAL EPITHELIAL CELL |
| 344 | Epithelial | BSS01080 | KIDNEY EPITHELIAL CELL |
| 345 | Epithelial | BSS00310 | PROXIMAL TUBULE EPITHELIAL CELL |
| 346 | Epithelial | BSS00701 | PROXIMAL TUBULE EPITHELIAL CELL |
| 347 | Epithelial | BSS01491 | RENAL CORTICAL EPITHELIAL CELL |
| 348 | Epithelial | BSS01505 | RETINAL EPITHELIAL CELL |
| 349 | Epithelial | BSS01103 | TUBULE CELL |
| 350 | Epithelial | BSS01102 | TUBULE CELL |
| 351 | Epithelial | BSS00153 | BRONCHIAL EPITHELIAL CELL |
| 352 | Epithelial | BSS00150 | BRONCHIAL EPITHELIAL CELL |
| 353 | Epithelial | BSS00703 | PANCREATIC DUCT EPITHELIAL CELL |
| 354 | Epithelial | BSS00075 | AMNION EPITHELIAL CELL |
| 355 | Epithelial | BSS00308 | PROSTATE EPITHELIAL CELL |
| 356 | Epithelial | BSS00309 | PROSTATE EPITHELIAL CELL |
| 357 | Epithelial | BSS01539 | PROSTATE EPITHELIAL CELL |
| 358 | Epithelial | BSS01538 | PROSTATE EPITHELIAL CELL |
| 359 | Epithelial | BSS01217 | BREAST EPITHELIAL CELL |
| 360 | Epithelial | BSS01224 | BREAST EPITHELIAL CELL |
| 361 | Epithelial | BSS01225 | BREAST EPITHELIAL CELL |
| 362 | Epithelial | BSS00356 | FORESKIN KERATINOCYTE |
| 363 | Epithelial | BSS00357 | FORESKIN KERATINOCYTE |
| 364 | Epithelial | BSS00358 | FORESKIN KERATINOCYTE |
| 365 | Epithelial | BSS00359 | FORESKIN KERATINOCYTE |
| 366 | Epithelial | BSS00360 | FORESKIN KERATINOCYTE |
| 367 | Epithelial | BSS00362 | FORESKIN KERATINOCYTE |
| 368 | Epithelial | BSS00363 | FORESKIN KERATINOCYTE |
| 369 | Epithelial | BSS00364 | FORESKIN KERATINOCYTE |
| 370 | Epithelial | BSS00365 | FORESKIN KERATINOCYTE |
| 371 | Epithelial | BSS00366 | FORESKIN KERATINOCYTE |
| 372 | Epithelial | BSS00355 | FORESKIN KERATINOCYTE |
| 575 | Kidney | BSS01533 | RENAL PELVIS |
| 576 | Kidney | BSS01497 | RENAL PELVIS |
| 577 | Kidney | BSS01498 | RENAL PELVIS |
| 578 | Kidney | BSS01153 | RENAL PELVIS |
| 579 | Kidney | BSS01534 | RENAL PELVIS |
| 580 | Kidney | BSS01499 | RENAL PELVIS |
| 581 | Liver | BSS00553 | HEPATIC STELLATE CELL |
| 582 | Liver | BSS00554 | HEPATOCYTE |
| 583 | Liver | BSS00511 | LIVER |
| 584 | Liver | BSS01164 | LIVER |
| 585 | Liver | BSS01170 | LIVER |
| 586 | Liver | BSS01169 | LIVER |
| 587 | Liver | BSS01159 | LIVER |
| 588 | Liver | BSS01168 | LIVER |
| 589 | Liver | BSS01519 | LIVER |
| 590 | Liver | BSS01158 | LIVER |
| 591 | Lung | BSS01195 | LUNG |
| 592 | Lung | BSS01142 | LUNG |
| 593 | Lung | BSS01525 | LUNG |
| 594 | Lung | BSS01143 | LUNG |
| 595 | Lung | BSS01526 | LUNG |
| 596 | Lung | BSS01137 | LUNG |
| 597 | Lung | BSS01520 | LUNG |
| 598 | Lung | BSS01138 | LUNG |
| 599 | Lung | BSS01521 | LUNG |
| 600 | Lung | BSS01139 | LUNG |
| 601 | Lung | BSS01192 | LUNG |
| 602 | Lung | BSS01522 | LUNG |
| 603 | Lung | BSS01140 | LUNG |
| 604 | Lung | BSS01523 | LUNG |
| 605 | Lung | BSS01141 | LUNG |
| 606 | Lung | BSS01524 | LUNG |
| 607 | Lung | BSS01205 | LUNG |
| 608 | Lung | BSS01147 | LUNG |
| 609 | Lung | BSS01529 | LUNG |
| 610 | Lung | BSS01148 | LUNG |
| 611 | Lung | BSS01149 | LUNG |
| 612 | Lung | BSS01530 | LUNG |
| 613 | Lung | BSS01202 | LUNG |
| 614 | Lung | BSS01144 | LUNG |
| 615 | Lung | BSS01527 | LUNG |
| 616 | Lung | BSS01203 | LUNG |
| 617 | Lung | BSS01145 | LUNG |
| 618 | Lung | BSS01146 | LUNG |
| 619 | Lung | BSS01528 | LUNG |
| 620 | Lung | BSS01204 | LUNG |
| 621 | Lung | BSS01189 | LUNG |
| 622 | Lung | BSS01188 | LUNG |
| 623 | Lung | BSS01187 | LUNG |
| 624 | Lung | BSS01198 | LUNG |
| 625 | Lung | BSS01186 | LUNG |
| 626 | Lung | BSS01869 | LUNG |
| 627 | Lung | BSS01196 | LUNG |
| 628 | Lung | BSS01197 | LUNG |
| 629 | Lung | BSS01871 | LUNG |
| 630 | Lung | BSS01201 | LUNG |
| 631 | Lung | BSS01190 | LUNG |
| 632 | Lung | BSS01870 | LUNG |
| 633 | Lung | BSS01193 | LUNG |
| 634 | Lymphoblastoid | BSS00403 | LYMPHOBLASTOID CELL LINE |
| 635 | Lymphoblastoid | BSS00462 | LYMPHOBLASTOID CELL LINE |
| 636 | Lymphoblastoid | BSS00456 | LYMPHOBLASTOID CELL LINE |
| 637 | Lymphoblastoid | BSS00457 | LYMPHOBLASTOID CELL LINE |
| 638 | Lymphoblastoid | BSS00438 | LYMPHOBLASTOID CELL LINE |
| 639 | Lymphoblastoid | BSS00473 | LYMPHOBLASTOID CELL LINE |
| 640 | Lymphoblastoid | BSS00471 | LYMPHOBLASTOID CELL LINE |
| 641 | Lymphoblastoid | BSS00474 | LYMPHOBLASTOID CELL LINE |
| 642 | Lymphoblastoid | BSS00404 | LYMPHOBLASTOID CELL LINE |
| 643 | Lymphoblastoid | BSS00405 | LYMPHOBLASTOID CELL LINE |
| 644 | Lymphoblastoid | BSS00472 | LYMPHOBLASTOID CELL LINE |
| 645 | Lymphoblastoid | BSS00428 | LYMPHOBLASTOID CELL LINE |
| 646 | Lymphoblastoid | BSS00454 | LYMPHOBLASTOID CELL LINE |
| 647 | Lymphoblastoid | BSS00427 | LYMPHOBLASTOID CELL LINE |
| 648 | Lymphoblastoid | BSS00452 | LYMPHOBLASTOID CELL LINE |
| 649 | Lymphoblastoid | BSS00395 | LYMPHOBLASTOID CELL LINE |
| 650 | Lymphoblastoid | BSS00439 | LYMPHOBLASTOID CELL LINE |
| 651 | Mesench | BSS00039 | ADIPOCYTE FROM MSC |
| 652 | Mesench | BSS00250 | AMNIOTIC FLUID FROM MSC |
| 653 | Mesench | BSS00279 | EMBRYONIC FACIAL PROMINENCE |
| 654 | Mesench | BSS01260 | MESENCHYMAL STEM CELL |
| 655 | Muscle | BSS01293 | ARM MUSCLE |
| 656 | Muscle | BSS01294 | ARM MUSCLE |
| 657 | Muscle | BSS01290 | ARM MUSCLE |
| 658 | Muscle | BSS00352 | ARM MUSCLE |
| 659 | Muscle | BSS01291 | ARM MUSCLE |

**Figure S1:** Sample list with tissue group, unique identifier, and short name for 859 observed/imputed samples (full metadata in Table S1, page 1 of 4).

| | Group | BSSID | Extended Info |
|---|---|---|---|
| 86 | Brain | BSS00206 | CEREBELLUM |
| 87 | Brain | BSS00219 | CINGULATE GYRUS |
| 88 | Brain | BSS00220 | CINGULATE GYRUS |
| 89 | Brain | BSS00369 | FRONTAL CORTEX |
| 90 | Brain | BSS00371 | FRONTAL CORTEX |
| 91 | Brain | BSS00385 | GERMINAL MATRIX |
| 92 | Brain | BSS00386 | GLOBUS PALLIDUS |
| 93 | Brain | BSS01125 | HIPPOCAMPUS |
| 94 | Brain | BSS01126 | HIPPOCAMPUS |
| 95 | Brain | BSS01124 | HIPPOCAMPUS |
| 96 | Brain | BSS00729 | INFERIOR PARIETAL CORTEX |
| 97 | Brain | BSS01250 | MEDULLA OBLONGATA |
| 98 | Brain | BSS01270 | MIDBRAIN |
| 99 | Brain | BSS01271 | MIDDLE FRONTAL AREA |
| 100 | Brain | BSS01272 | MIDDLE FRONTAL AREA |
| 101 | Brain | BSS01273 | MIDDLE FRONTAL GYRUS |
| 102 | Brain | BSS01388 | OCCIPITAL LOBE |
| 103 | Brain | BSS01451 | PONS |
| 104 | Brain | BSS01452 | POSTERIOR CINGULATE CORTEX |
| 105 | Brain | BSS01469 | PUTAMEN |
| 106 | Brain | BSS01675 | SUBSTANTIA NIGRA |
| 107 | Brain | BSS01676 | SUBSTANTIA NIGRA |
| 108 | Brain | BSS01677 | SUPERIOR TEMPORAL GYRUS |
| 109 | Brain | BSS01714 | TEMPORAL LOBE |
| 110 | Brain | BSS01712 | TEMPORAL LOBE |
| 111 | Cancer | BSS01105 | ACUTE LYMPHOBLASTIC LEUKEMIA |
| 112 | Cancer | BSS00267 | ACUTE LYMPHOBLASTIC LEUKEMIA |
| 113 | Cancer | BSS01178 | ACUTE LYMPHOBLASTIC LEUKEMIA |
| 114 | Cancer | BSS01267 | OSTEOSARCOMA |
| 115 | Cancer | BSS01550 | OSTEOSARCOMA |
| 116 | Cancer | BSS00246 | DESMOPLASTIC MEDULLOBLASTOMA |
| 117 | Cancer | BSS01208 | GLIOBLASTOMA |
| 118 | Cancer | BSS00004 | GLIOBLASTOMA |
| 119 | Cancer | BSS00482 | GLIOBLASTOMA |
| 120 | Cancer | BSS01251 | MEDULLOBLASTOMA |
| 121 | Cancer | BSS01554 | NEUROBLASTOMA |
| 122 | Cancer | BSS01558 | NEUROBLASTOMA |
| 123 | Cancer | BSS00102 | NEUROBLASTOMA |
| 124 | Cancer | BSS01571 | NEUROBLASTOMA |
| 125 | Cancer | BSS01562 | NEUROBLASTOMA |
| 126 | Cancer | BSS01559 | NEUROEPITHELIOMA |
| 127 | Cancer | BSS00481 | NEUROGLIOMA |
| 128 | Cancer | BSS01535 | COLON CARCINOMA |
| 129 | Cancer | BSS01682 | COLORECTAL ADENOCARCINOMA |
| 130 | Cancer | BSS00708 | COLORECTAL ADENOCARCINOMA |
| 131 | Cancer | BSS01179 | COLORECTAL ADENOCARCINOMA |
| 132 | Cancer | BSS00159 | COLORECTAL ADENOCARCINOMA |
| 133 | Cancer | BSS00492 | COLORECTAL ADENOCARCINOMA |
| 134 | Cancer | BSS01412 | PARATHYROID ADENOMA |
| 135 | Cancer | BSS01411 | PARATHYROID ADENOMA |
| 136 | Cancer | BSS01386 | TESTICULAR EMBRYONAL CARCINOMA |
| 137 | Cancer | BSS01536 | MELANOMA |
| 138 | Cancer | BSS01551 | MELANOMA |
| 139 | Cancer | BSS00222 | MELANOMA |
| 140 | Cancer | BSS01365 | MYELOMA |
| 141 | Cancer | BSS01890 | EYE RETINOBLASTOMA |
| 142 | Cancer | BSS00702 | ACUTE PROMYELOCYTIC LEUKEMIA |
| 143 | Cancer | BSS01356 | ACUTE PROMYELOCYTIC LEUKEMIA |
| 144 | Cancer | BSS01391 | B CELL LYMPHOMA |
| 145 | Cancer | BSS01390 | B CELL LYMPHOMA |
| 146 | Cancer | BSS01065 | B CELL LYMPHOMA |
| 147 | Cancer | BSS00268 | B CELL LYMPHOMA |
| 148 | Cancer | BSS01664 | B CELL LYMPHOMA |
| 149 | Cancer | BSS01389 | B CELL LYMPHOMA |
| 150 | Cancer | BSS01350 | BURKITT LYMPHOMA |
| 151 | Cancer | BSS01351 | BURKITT LYMPHOMA |
| 152 | Cancer | BSS00491 | HAPLOID MYELOGENOUS LEUKEMIA |
| 153 | Cancer | BSS01038 | MYELOGENOUS LEUKEMIA |
| 154 | Cancer | BSS01039 | MYELOGENOUS LEUKEMIA |
| 155 | Cancer | BSS01056 | MYELOGENOUS LEUKEMIA |
| 156 | Cancer | BSS01057 | MYELOGENOUS LEUKEMIA |
| 157 | Cancer | BSS01059 | MYELOGENOUS LEUKEMIA |
| 158 | Cancer | BSS00221 | MYELOGENOUS LEUKEMIA |
| 159 | Cancer | BSS01066 | MYELOGENOUS LEUKEMIA |
| 160 | Cancer | BSS00762 | MYELOGENOUS LEUKEMIA |
| 161 | Cancer | BSS01104 | MYELOMA |
| 162 | Cancer | BSS01274 | MYELOMA |
| 163 | Cancer | BSS01537 | PLASMA CELL MYELOMA |
| 164 | Cancer | BSS00160 | KIDNEY CLEAR CELL CARCINOMA |
| 165 | Cancer | BSS00372 | KIDNEY RHABOID TUMOR |
| 166 | Cancer | BSS00037 | RENAL CELL ADENOCARCINOMA |
| 167 | Cancer | BSS01474 | RENAL CELL ADENOCARCINOMA |
| 168 | Cancer | BSS01481 | RENAL CELL CARCINOMA |
| 169 | Cancer | BSS00718 | HEPATOCELLULAR CARCINOMA |
| 373 | Epithelial | BSS00361 | FORESKIN KERATINOCYTE |
| 374 | Epithelial | BSS00367 | FORESKIN KERATINOCYTE |
| 375 | Epithelial | BSS00354 | FORESKIN KERATINOCYTE |
| 376 | Epithelial | BSS01071 | KERATINOCYTE |
| 377 | Epithelial | BSS01068 | KERATINOCYTE |
| 378 | Epithelial | BSS01209 | MAMMARY EPITHELIAL CELL |
| 379 | Epithelial | BSS01211 | MAMMARY EPITHELIAL CELL |
| 380 | Epithelial | BSS01213 | MAMMARY EPITHELIAL CELL |
| 381 | Epithelial | BSS01185 | MAMMARY LUMINAL EPITHELIAL CELL |
| 382 | Epithelial | BSS01340 | MAMMARY MYOEPITHELIAL CELL |
| 383 | Epithelial | BSS01341 | MAMMARY MYOEPITHELIAL CELL |
| 384 | Epithelial | BSS01181 | SKIN LEG |
| 385 | Epithelial | BSS01182 | SKIN LEG |
| 386 | Epithelial | BSS01587 | SKIN OF BODY |
| 387 | ES-deriv | BSS00112 | BIPOLAR NEURON DERIV |
| 388 | ES-deriv | BSS01366 | NEURAL DERIV |
| 389 | ES-deriv | BSS00272 | NEURAL PROGENITOR DERIV |
| 390 | ES-deriv | BSS01372 | NEURAL PROGENITOR DERIV |
| 391 | ES-deriv | BSS01370 | NEURAL PROGENITOR DERIV |
| 392 | ES-deriv | BSS01371 | NEURAL PROGENITOR DERIV |
| 393 | ES-deriv | BSS01375 | NEURON DERIV |
| 394 | ES-deriv | BSS00169 | CARDIAC MESODERM DERIV |
| 395 | ES-deriv | BSS00171 | CARDIAC MUSCLE DERIV |
| 396 | ES-deriv | BSS00556 | HEPATOCYTE DERIV |
| 397 | ES-deriv | BSS01261 | MESENCHYMAL STEM DERIV |
| 398 | ES-deriv | BSS01857 | TROPHOBLAST DERIV |
| 399 | ES-deriv | BSS01612 | SMOOTH MUSCLE DERIV |
| 400 | ES-deriv | BSS00273 | ECTODERMAL DERIV |
| 401 | ES-deriv | BSS00285 | ENDODERMAL CELL |
| 402 | ES-deriv | BSS00287 | ENDODERMAL DERIV |
| 403 | ES-deriv | BSS01263 | MESENDODERM DERIV |
| 404 | ES-deriv | BSS01264 | MESODERMAL DERIV |
| 405 | ESC | BSS00277 | ESC |
| 406 | ESC | BSS00315 | ESC |
| 407 | ESC | BSS01866 | ESC |
| 408 | ESC | BSS00483 | ESC |
| 409 | ESC | BSS00715 | ESC |
| 410 | ESC | BSS00716 | ESC |
| 411 | ESC | BSS00717 | ESC |
| 412 | ESC | BSS00484 | ESC |
| 413 | ESC | BSS00478 | ESC |
| 414 | Eye | BSS00329 | EYE |
| 415 | Eye | BSS00328 | EYE |
| 416 | Eye | BSS01504 | EYE RETINA |
| 417 | Eye | BSS01503 | EYE RETINA |
| 418 | Eye | BSS01502 | EYE RETINA |
| 419 | Heart | BSS00079 | AORTA |
| 420 | Heart | BSS00080 | AORTA |
| 421 | Heart | BSS00088 | ASCENDING AORTA |
| 422 | Heart | BSS00087 | ASCENDING AORTA |
| 423 | Heart | BSS00242 | CORONARY ARTERY |
| 424 | Heart | BSS00243 | CORONARY ARTERY |
| 425 | Heart | BSS00505 | HEART |
| 426 | Heart | BSS00498 | HEART |
| 427 | Heart | BSS00499 | HEART |
| 428 | Heart | BSS00500 | HEART |
| 429 | Heart | BSS00502 | HEART |
| 430 | Heart | BSS00503 | HEART |
| 431 | Heart | BSS00501 | HEART |
| 432 | Heart | BSS00516 | HEART |
| 433 | Heart | BSS00522 | HEART |
| 434 | Heart | BSS00518 | HEART |
| 435 | Heart | BSS00519 | HEART |
| 436 | Heart | BSS00514 | HEART |
| 437 | Heart | BSS00520 | HEART |
| 438 | Heart | BSS00495 | HEART |
| 439 | Heart | BSS00496 | HEART |
| 440 | Heart | BSS00494 | HEART |
| 441 | Heart | BSS00521 | HEART |
| 442 | Heart | BSS00517 | HEART |
| 443 | Heart | BSS00493 | HEART |
| 444 | Heart | BSS01127 | HEART LEFT ATRIUM |
| 445 | Heart | BSS00509 | HEART LEFT VENTRICLE |
| 446 | Heart | BSS00508 | HEART LEFT VENTRICLE |
| 447 | Heart | BSS00506 | HEART LEFT VENTRICLE |
| 448 | Heart | BSS00513 | HEART LEFT VENTRICLE |
| 449 | Heart | BSS00512 | HEART LEFT VENTRICLE |
| 450 | Heart | BSS00507 | HEART LEFT VENTRICLE |
| 451 | Heart | BSS01506 | HEART RIGHT ATRIUM |
| 452 | Heart | BSS01508 | HEART RIGHT ATRIUM |
| 453 | Heart | BSS01507 | HEART RIGHT ATRIUM |
| 454 | Heart | BSS00523 | HEART RIGHT VENTRICLE |
| 455 | Heart | BSS00524 | HEART RIGHT VENTRICLE |
| 456 | Heart | BSS00525 | HEART RIGHT VENTRICLE |
| 660 | Muscle | BSS01292 | ARM MUSCLE |
| 661 | Muscle | BSS01303 | ARM MUSCLE |
| 662 | Muscle | BSS01304 | ARM MUSCLE |
| 663 | Muscle | BSS01295 | ARM MUSCLE |
| 664 | Muscle | BSS01296 | ARM MUSCLE |
| 665 | Muscle | BSS01297 | ARM MUSCLE |
| 666 | Muscle | BSS01298 | ARM MUSCLE |
| 667 | Muscle | BSS01299 | ARM MUSCLE |
| 668 | Muscle | BSS01300 | ARM MUSCLE |
| 669 | Muscle | BSS01301 | ARM MUSCLE |
| 670 | Muscle | BSS01289 | ARM MUSCLE |
| 671 | Muscle | BSS01308 | BACK MUSCLE |
| 672 | Muscle | BSS01309 | BACK MUSCLE |
| 673 | Muscle | BSS01305 | BACK MUSCLE |
| 674 | Muscle | BSS01306 | BACK MUSCLE |
| 675 | Muscle | BSS01307 | BACK MUSCLE |
| 676 | Muscle | BSS01315 | BACK MUSCLE |
| 677 | Muscle | BSS01316 | BACK MUSCLE |
| 678 | Muscle | BSS01317 | BACK MUSCLE |
| 679 | Muscle | BSS01310 | BACK MUSCLE |
| 680 | Muscle | BSS01311 | BACK MUSCLE |
| 681 | Muscle | BSS01312 | BACK MUSCLE |
| 682 | Muscle | BSS01313 | BACK MUSCLE |
| 683 | Muscle | BSS01314 | BACK MUSCLE |
| 684 | Muscle | BSS00170 | CARDIAC MYOCYTE |
| 685 | Muscle | BSS00376 | GASTROCNEMIUS MEDIALIS |
| 686 | Muscle | BSS00378 | GASTROCNEMIUS MEDIALIS |
| 687 | Muscle | BSS00377 | GASTROCNEMIUS MEDIALIS |
| 688 | Muscle | BSS00379 | GASTROCNEMIUS MEDIALIS |
| 689 | Muscle | BSS01322 | LEG MUSCLE |
| 690 | Muscle | BSS01318 | LEG MUSCLE |
| 691 | Muscle | BSS01320 | LEG MUSCLE |
| 692 | Muscle | BSS01321 | LEG MUSCLE |
| 693 | Muscle | BSS01329 | LEG MUSCLE |
| 694 | Muscle | BSS01330 | LEG MUSCLE |
| 695 | Muscle | BSS01323 | LEG MUSCLE |
| 696 | Muscle | BSS01324 | LEG MUSCLE |
| 697 | Muscle | BSS01325 | LEG MUSCLE |
| 698 | Muscle | BSS01327 | LEG MUSCLE |
| 699 | Muscle | BSS00700 | LEG MUSCLE |
| 700 | Muscle | BSS01328 | LEG MUSCLE |
| 701 | Muscle | BSS01319 | LEG MUSCLE |
| 702 | Muscle | BSS01460 | PSOAS MUSCLE |
| 703 | Muscle | BSS01461 | PSOAS MUSCLE |
| 704 | Muscle | BSS01462 | PSOAS MUSCLE |
| 705 | Muscle | BSS01463 | PSOAS MUSCLE |
| 706 | Muscle | BSS01581 | SKELETAL MUSCLE |
| 707 | Muscle | BSS01577 | SKELETAL MUSCLE |
| 708 | Muscle | BSS01578 | SKELETAL MUSCLE |
| 709 | Muscle | BSS01572 | SKELETAL MUSCLE CELL |
| 710 | Muscle | BSS01845 | TONGUE |
| 711 | Muscle | BSS01846 | TONGUE |
| 712 | Muscle | BSS01331 | TRUNK MUSCLE |
| 713 | Muscle | BSS01333 | TRUNK MUSCLE |
| 714 | Muscle | BSS01334 | TRUNK MUSCLE |
| 715 | Muscle | BSS01332 | TRUNK MUSCLE |
| 716 | Myosat | BSS01338 | MYOCYTE |
| 717 | Myosat | BSS01344 | MYOTUBE |
| 718 | Myosat | BSS01155 | SKELETAL MUSCLE MYOBLAST |
| 719 | Myosat | BSS01573 | SKELETAL MUSCLE MYOBLAST |
| 720 | Myosat | BSS01574 | SKELETAL MUSCLE MYOBLAST |
| 721 | Myosat | BSS01576 | SKELETAL MUSCLE SATELLITE CELL |
| 722 | Neurosph | BSS01378 | NEUROSPHERE |
| 723 | Neurosph | BSS01379 | NEUROSPHERE |
| 724 | Neurosph | BSS01377 | NEUROSPHERE |
| 725 | Neurosph | BSS01392 | OLFACTORY NEUROSPHERE |
| 726 | Other | BSS00148 | BREAST EPITHELIUM |
| 727 | Other | BSS00145 | BREAST EPITHELIUM |
| 728 | Other | BSS00146 | BREAST EPITHELIUM |
| 729 | Other | BSS00304 | EPIDERMAL MELANOCYTE |
| 730 | Other | BSS00368 | FORESKIN MELANOCYTE |
| 731 | Other | BSS01156 | LIMB EMBRYO |
| 732 | Other | BSS01157 | LIMB EMBRYO |
| 733 | Other | BSS01216 | MAMMARY STEM CELL |
| 734 | Pancreas | BSS00121 | BODY OF PANCREAS |
| 735 | Pancreas | BSS00122 | BODY OF PANCREAS |
| 736 | Pancreas | BSS00123 | BODY OF PANCREAS |
| 737 | Pancreas | BSS00124 | BODY OF PANCREAS |
| 738 | Pancreas | BSS00758 | ISLET PRECURSOR CELL |
| 739 | Pancreas | BSS01406 | PANCREAS |
| 740 | Pancreas | BSS01407 | PANCREAS |
| 741 | Placenta & EEM | BSS00074 | AMNION |
| 742 | Placenta & EEM | BSS00076 | AMNION STEM CELL |
| 743 | Placenta & EEM | BSS00209 | CHORION |

**Figure S1:** (continued, 2 of 4)

| | Group | BSSID | Extended Info |
|---|---|---|---|
| 170 | Cancer | BSS00719 | HEPATOCELLULAR CARCINOMA |
| 171 | Cancer | BSS00558 | HEPATOCELLULAR CARCINOMA |
| 172 | Cancer | BSS01360 | LARGE CELL LUNG CANCER |
| 173 | Cancer | BSS01415 | LUNG ADENOCARCINOMA |
| 174 | Cancer | BSS00017 | LUNG EPITHELIAL CARCINOMA |
| 175 | Cancer | BSS00019 | LUNG EPITHELIAL CARCINOMA |
| 176 | Cancer | BSS00021 | LUNG EPITHELIAL CARCINOMA |
| 177 | Cancer | BSS00022 | LUNG EPITHELIAL CARCINOMA |
| 178 | Cancer | BSS00027 | LUNG EPITHELIAL CARCINOMA |
| 179 | Cancer | BSS00016 | LUNG EPITHELIAL CARCINOMA |
| 180 | Cancer | BSS00020 | LUNG EPITHELIAL CARCINOMA |
| 181 | Cancer | BSS00023 | LUNG EPITHELIAL CARCINOMA |
| 182 | Cancer | BSS00024 | LUNG EPITHELIAL CARCINOMA |
| 183 | Cancer | BSS00026 | LUNG EPITHELIAL CARCINOMA |
| 184 | Cancer | BSS00028 | LUNG EPITHELIAL CARCINOMA |
| 185 | Cancer | BSS00029 | LUNG EPITHELIAL CARCINOMA |
| 186 | Cancer | BSS00018 | LUNG EPITHELIAL CARCINOMA |
| 187 | Cancer | BSS00025 | LUNG EPITHELIAL CARCINOMA |
| 188 | Cancer | BSS00030 | LUNG EPITHELIAL CARCINOMA |
| 189 | Cancer | BSS00007 | LUNG EPITHELIAL CARCINOMA |
| 190 | Cancer | BSS00013 | LUNG EPITHELIAL CARCINOMA |
| 191 | Cancer | BSS00015 | LUNG EPITHELIAL CARCINOMA |
| 192 | Cancer | BSS01359 | SQUAMOUS CELL CARCINOMA |
| 193 | Cancer | BSS00035 | MUSCLE EWING SARCOMA |
| 194 | Cancer | BSS01549 | RHABDOMYOSARCOMA |
| 195 | Cancer | BSS00036 | ADENOID CYSTIC CARCINOMA |
| 196 | Cancer | BSS01240 | MAMMARY GLAND ADENOCARCINOMA |
| 197 | Cancer | BSS01243 | MAMMARY GLAND ADENOCARCINOMA |
| 198 | Cancer | BSS01244 | MAMMARY GLAND ADENOCARCINOMA |
| 199 | Cancer | BSS01235 | MAMMARY GLAND ADENOCARCINOMA |
| 200 | Cancer | BSS01226 | MAMMARY GLAND ADENOCARCINOMA |
| 201 | Cancer | BSS01699 | MAMMARY GLAND DUCTAL CARCINOMA |
| 202 | Cancer | BSS01705 | MAMMARY GLAND DUCTAL CARCINOMA |
| 203 | Cancer | BSS00003 | PANCREAS ADENOCARCINOMA |
| 204 | Cancer | BSS01405 | PANCREAS DUCT EPITHELIAL CARCINOMA |
| 205 | Cancer | BSS00541 | CERVIX ADENOCARCINOMA |
| 206 | Cancer | BSS00531 | CERVIX ADENOCARCINOMA |
| 207 | Cancer | BSS00529 | CERVIX ADENOCARCINOMA |
| 208 | Cancer | BSS00748 | ENDOMETRIAL ADENOCARCINOMA |
| 209 | Cancer | BSS00756 | ENDOMETRIAL ADENOCARCINOMA |
| 210 | Cancer | BSS00745 | ENDOMETRIAL ADENOCARCINOMA |
| 211 | Cancer | BSS01174 | PROSTATE ADENOCARCINOMA |
| 212 | Cancer | BSS01173 | PROSTATE ADENOCARCINOMA |
| 213 | Cancer | BSS01414 | PROSTATE ADENOCARCINOMA |
| 214 | Cancer | BSS00157 | PROSTATE CANCER |
| 215 | Cancer | BSS01888 | PROSTATE EPITHELIAL CARCINOMA |
| 216 | Cancer | BSS00001 | PROSTATE EPITHELIAL CARCINOMA |
| 217 | Cancer | BSS00002 | PROSTATE EPITHELIAL CARCINOMA |
| 218 | Cancer | BSS00709 | FIBROSARCOMA |
| 219 | Digestive | BSS00227 | COLON MUCOSA |
| 220 | Digestive | BSS00228 | COLON MUCOSA |
| 221 | Digestive | BSS00271 | DUODENUM MUCOSA |
| 222 | Digestive | BSS00270 | DUODENUM MUCOSA |
| 223 | Digestive | BSS00316 | ESOPHAGUS |
| 224 | Digestive | BSS00318 | ESOPHAGUS |
| 225 | Digestive | BSS00323 | ESOPHAGUS MUSCULARIS MUCOSA |
| 226 | Digestive | BSS00322 | ESOPHAGUS MUSCULARIS MUCOSA |
| 227 | Digestive | BSS00321 | ESOPHAGUS MUSCULARIS MUCOSA |
| 228 | Digestive | BSS00324 | ESOPHAGUS SQUAMOUS EPITHELIUM |
| 229 | Digestive | BSS00326 | ESOPHAGUS SQUAMOUS EPITHELIUM |
| 230 | Digestive | BSS00325 | ESOPHAGUS SQUAMOUS EPITHELIUM |
| 231 | Digestive | BSS00380 | GASTROESOPHAGEAL SPHINCTER |
| 232 | Digestive | BSS00381 | GASTROESOPHAGEAL SPHINCTER |
| 233 | Digestive | BSS01116 | LARGE INTESTINE |
| 234 | Digestive | BSS01117 | LARGE INTESTINE |
| 235 | Digestive | BSS01109 | LARGE INTESTINE |
| 236 | Digestive | BSS01110 | LARGE INTESTINE |
| 237 | Digestive | BSS01111 | LARGE INTESTINE |
| 238 | Digestive | BSS01112 | LARGE INTESTINE |
| 239 | Digestive | BSS01113 | LARGE INTESTINE |
| 240 | Digestive | BSS01114 | LARGE INTESTINE |
| 241 | Digestive | BSS01122 | LARGE INTESTINE |
| 242 | Digestive | BSS01118 | LARGE INTESTINE |
| 243 | Digestive | BSS01120 | LARGE INTESTINE |
| 244 | Digestive | BSS01121 | LARGE INTESTINE |
| 245 | Digestive | BSS01119 | LARGE INTESTINE |
| 246 | Digestive | BSS01427 | PEYERS PATCH |
| 247 | Digestive | BSS01428 | PEYERS PATCH |
| 248 | Digestive | BSS01426 | PEYERS PATCH |
| 249 | Digestive | BSS01282 | RECTUM MUCOSA |
| 250 | Digestive | BSS01283 | RECTUM MUCOSA |
| 251 | Digestive | BSS01542 | SIGMOID COLON |
| 252 | Digestive | BSS01546 | SIGMOID COLON |
| 253 | Digestive | BSS01547 | SIGMOID COLON |

| | Group | BSSID | Extended Info |
|---|---|---|---|
| 457 | Heart | BSS01815 | THORACIC AORTA |
| 458 | Heart | BSS01814 | THORACIC AORTA |
| 459 | Heart | BSS01839 | TIBIAL ARTERY |
| 460 | Heart | BSS01838 | TIBIAL ARTERY |
| 461 | Heart | BSS01837 | TIBIAL ARTERY |
| 462 | HSC & B-cell | BSS00097 | B CELL |
| 463 | HSC & B-cell | BSS01345 | B CELL |
| 464 | HSC & B-cell | BSS00098 | B CELL |
| 465 | HSC & B-cell | BSS00093 | B CELL |
| 466 | HSC & B-cell | BSS00096 | B CELL |
| 467 | HSC & B-cell | BSS00100 | B CELL |
| 468 | HSC & B-cell | BSS00101 | B CELL |
| 469 | HSC & B-cell | BSS00095 | B CELL |
| 470 | HSC & B-cell | BSS00179 | CD14 MONOCYTE |
| 471 | HSC & B-cell | BSS00181 | CD14 MONOCYTE |
| 472 | HSC & B-cell | BSS00180 | CD14 MONOCYTE |
| 473 | HSC & B-cell | BSS00178 | CD14 MONOCYTE |
| 474 | HSC & B-cell | BSS00182 | CD1C MYELOID DENDRITIC CELL |
| 475 | HSC & B-cell | BSS00233 | CD34 CMP |
| 476 | HSC & B-cell | BSS00230 | CD34 CMP |
| 477 | HSC & B-cell | BSS00236 | CD34 CMP |
| 478 | HSC & B-cell | BSS00238 | CD34 CMP |
| 479 | HSC & B-cell | BSS00240 | CD34 CMP |
| 480 | HSC & B-cell | BSS00241 | CD34 CMP |
| 481 | HSC & B-cell | BSS00234 | CD34 CMP |
| 482 | HSC & B-cell | BSS00235 | CD34 CMP |
| 483 | HSC & B-cell | BSS00229 | CD34 CMP |
| 484 | HSC & B-cell | BSS00237 | CD34 CMP |
| 485 | HSC & B-cell | BSS00239 | CD34 CMP |
| 486 | HSC & B-cell | BSS00231 | CD34 CMP |
| 487 | HSC & B-cell | BSS00232 | CD34 CMP |
| 488 | HSC & B-cell | BSS00384 | GERMINAL CENTER |
| 489 | HSC & B-cell | BSS00760 | LYMPHOCYTE |
| 490 | HSC & B-cell | BSS00544 | MPP |
| 491 | HSC & B-cell | BSS00545 | MPP |
| 492 | HSC & B-cell | BSS00546 | MPP |
| 493 | HSC & B-cell | BSS00547 | MPP |
| 494 | HSC & B-cell | BSS00548 | MPP |
| 495 | HSC & B-cell | BSS00549 | MPP |
| 496 | HSC & B-cell | BSS00550 | MPP |
| 497 | HSC & B-cell | BSS00551 | MPP |
| 498 | HSC & B-cell | BSS00552 | MPP |
| 499 | HSC & B-cell | BSS00543 | MPP |
| 500 | HSC & B-cell | BSS01381 | NEUTROPHIL |
| 501 | HSC & B-cell | BSS01380 | NEUTROPHIL |
| 502 | HSC & B-cell | BSS01353 | NK CELL |
| 503 | HSC & B-cell | BSS01355 | NK CELL |
| 504 | HSC & B-cell | BSS01354 | NK CELL |
| 505 | iPSC | BSS00742 | iPSC |
| 506 | iPSC | BSS00735 | iPSC |
| 507 | iPSC | BSS00741 | iPSC |
| 508 | iPSC | BSS00732 | iPSC |
| 509 | iPSC | BSS00733 | iPSC |
| 510 | iPSC | BSS00244 | iPSC |
| 511 | iPSC | BSS01107 | iPSC |
| 512 | iPSC | BSS01108 | iPSC |
| 513 | iPSC | BSS00738 | iPSC |
| 514 | iPSC | BSS00736 | iPSC |
| 515 | iPSC | BSS00737 | iPSC |
| 516 | iPSC | BSS00739 | iPSC |
| 517 | iPSC | BSS00731 | iPSC |
| 518 | iPSC | BSS00734 | iPSC |
| 519 | iPSC | BSS00477 | iPSC |
| 520 | Kidney | BSS01091 | KIDNEY |
| 521 | Kidney | BSS01132 | KIDNEY |
| 522 | Kidney | BSS01512 | KIDNEY |
| 523 | Kidney | BSS01133 | KIDNEY |
| 524 | Kidney | BSS01513 | KIDNEY |
| 525 | Kidney | BSS01084 | KIDNEY |
| 526 | Kidney | BSS01128 | KIDNEY |
| 527 | Kidney | BSS01509 | KIDNEY |
| 528 | Kidney | BSS01085 | KIDNEY |
| 529 | Kidney | BSS01129 | KIDNEY |
| 530 | Kidney | BSS01086 | KIDNEY |
| 531 | Kidney | BSS01510 | KIDNEY |
| 532 | Kidney | BSS01089 | KIDNEY |
| 533 | Kidney | BSS01130 | KIDNEY |
| 534 | Kidney | BSS01511 | KIDNEY |
| 535 | Kidney | BSS01090 | KIDNEY |
| 536 | Kidney | BSS01078 | KIDNEY |
| 537 | Kidney | BSS01100 | KIDNEY |
| 538 | Kidney | BSS01101 | KIDNEY |
| 539 | Kidney | BSS01135 | KIDNEY |
| 540 | Kidney | BSS01516 | KIDNEY |

| | Group | BSSID | Extended Info |
|---|---|---|---|
| 744 | Placenta & EEM | BSS00211 | CHORION |
| 745 | Placenta & EEM | BSS00212 | CHORION |
| 746 | Placenta & EEM | BSS00215 | CHORIONIC VILLUS |
| 747 | Placenta & EEM | BSS00216 | CHORIONIC VILLUS |
| 748 | Placenta & EEM | BSS00217 | CHORIONIC VILLUS |
| 749 | Placenta & EEM | BSS00214 | CHORIONIC VILLUS |
| 750 | Placenta & EEM | BSS01440 | PLACENTA |
| 751 | Placenta & EEM | BSS01436 | PLACENTA |
| 752 | Placenta & EEM | BSS01437 | PLACENTA |
| 753 | Placenta & EEM | BSS01435 | PLACENTA |
| 754 | Placenta & EEM | BSS01443 | PLACENTA |
| 755 | Placenta & EEM | BSS01444 | PLACENTA |
| 756 | Placenta & EEM | BSS01433 | PLACENTA |
| 757 | Placenta & EEM | BSS01432 | PLACENTA |
| 758 | Placenta & EEM | BSS01430 | PLACENTA |
| 759 | Placenta & EEM | BSS01448 | PLACENTA |
| 760 | Placenta & EEM | BSS01446 | PLACENTA |
| 761 | Placenta & EEM | BSS01441 | PLACENTA |
| 762 | Placenta & EEM | BSS01431 | PLACENTA |
| 763 | Placenta & EEM | BSS01438 | PLACENTA |
| 764 | Placenta & EEM | BSS00714 | TROPHOBLAST |
| 765 | Placenta & EEM | BSS01856 | TROPHOBLAST |
| 766 | Placenta & EEM | BSS01853 | TROPHOBLAST |
| 767 | Placenta & EEM | BSS01855 | TROPHOBLAST |
| 768 | Placenta & EEM | BSS01852 | TROPHOBLAST |
| 769 | Placenta & EEM | BSS01859 | TROPHOBLAST |
| 770 | Placenta & EEM | BSS01860 | TROPHOBLAST |
| 771 | Placenta & EEM | BSS01867 | UMBILICAL CORD |
| 772 | PNS | BSS01618 | SPINAL CORD |
| 773 | PNS | BSS01619 | SPINAL CORD |
| 774 | PNS | BSS01617 | SPINAL CORD |
| 775 | PNS | BSS01621 | SPINAL CORD |
| 776 | PNS | BSS01620 | SPINAL CORD |
| 777 | PNS | BSS01614 | SPINAL CORD |
| 778 | PNS | BSS01613 | SPINAL CORD |
| 779 | PNS | BSS01842 | TIBIAL NERVE |
| 780 | PNS | BSS01840 | TIBIAL NERVE |
| 781 | PNS | BSS01841 | TIBIAL NERVE |
| 782 | Reproductive | BSS01456 | PROSTATE GLAND |
| 783 | Reproductive | BSS01457 | PROSTATE GLAND |
| 784 | Reproductive | BSS01459 | PROSTATE GLAND |
| 785 | Reproductive | BSS01884 | UTERUS |
| 786 | Reproductive | BSS01886 | VAGINA |
| 787 | Reproductive | BSS01887 | VAGINA |
| 788 | Sm. Muscle | BSS01606 | BRAIN VASCULATURE SMOOTH MUSCLE CELL |
| 789 | Sm. Muscle | BSS01285 | COLON MUSCLE |
| 790 | Sm. Muscle | BSS01286 | COLON MUSCLE |
| 791 | Sm. Muscle | BSS01288 | DUODENUM MUSCLE |
| 792 | Sm. Muscle | BSS01287 | DUODENUM MUSCLE |
| 793 | Sm. Muscle | BSS01475 | RECTUM MUSCLE |
| 794 | Sm. Muscle | BSS01660 | STOMACH MUSCLE |
| 795 | Sm. Muscle | BSS01659 | STOMACH MUSCLE |
| 796 | Spleen | BSS01625 | SPLEEN |
| 797 | Spleen | BSS01628 | SPLEEN |
| 798 | Spleen | BSS01629 | SPLEEN |
| 799 | Spleen | BSS01633 | SPLEEN |
| 800 | Spleen | BSS01634 | SPLEEN |
| 801 | Spleen | BSS01631 | SPLEEN |
| 802 | Spleen | BSS01630 | SPLEEN |
| 803 | Stromal | BSS01661 | BONE MARROW STROMAL CELL |
| 804 | Stromal | BSS00144 | PERICYTE |
| 805 | Stromal | BSS00349 | CONJUNCTIVA FIBROBLAST |
| 806 | Stromal | BSS00347 | AORTA FIBROBLAST |
| 807 | Stromal | BSS00168 | CARDIAC FIBROBLAST |
| 808 | Stromal | BSS00166 | CARDIAC FIBROBLAST |
| 809 | Stromal | BSS00167 | CARDIAC FIBROBLAST |
| 810 | Stromal | BSS00064 | LUNG FIBROBLAST |
| 811 | Stromal | BSS01891 | LUNG FIBROBLAST |
| 812 | Stromal | BSS00342 | LUNG FIBROBLAST |
| 813 | Stromal | BSS00339 | LUNG FIBROBLAST |
| 814 | Stromal | BSS00062 | LUNG FIBROBLAST |
| 815 | Stromal | BSS00341 | LUNG FIBROBLAST |
| 816 | Stromal | BSS00720 | LUNG FIBROBLAST |
| 817 | Stromal | BSS00345 | PULMONARY ARTERY FIBROBLAST |
| 818 | Stromal | BSS00338 | GINGIVAL FIBROBLAST |
| 819 | Stromal | BSS00067 | GINGIVAL FIBROBLAST |
| 820 | Stromal | BSS00344 | PERIDONTAL LIGAMENT FIBROBLAST |
| 821 | Stromal | BSS00350 | VILLOUS MESENCHYME FIBROBLAST |
| 822 | Stromal | BSS00332 | BREAST FIBROBLAST |
| 823 | Stromal | BSS00333 | BREAST FIBROBLAST |
| 824 | Stromal | BSS00335 | DERMIS FIBROBLAST |
| 825 | Stromal | BSS00334 | DERMIS FIBROBLAST |
| 826 | Stromal | BSS00337 | DERMIS FIBROBLAST |
| 827 | Stromal | BSS00697 | FORESKIN FIBROBLAST |

**Figure S1:** (continued, 3 of 4)

| | Group | BSSID | Extended Info | | Group | BSSID | Extended Info | | Group | BSSID | Extended Info |
|---|---|---|---|---|---|---|---|---|---|---|---|
| 254 | Digestive | BSS01548 | SIGMOID COLON | 541 | Kidney | BSS01517 | KIDNEY | 828 | Stromal | BSS00353 | FORESKIN FIBROBLAST |
| 255 | Digestive | BSS01545 | SIGMOID COLON | 542 | Kidney | BSS01136 | KIDNEY | 829 | Stromal | BSS00343 | MAMMARY FIBROBLAST |
| 256 | Digestive | BSS01543 | SIGMOID COLON | 543 | Kidney | BSS01518 | KIDNEY | 830 | Stromal | BSS00275 | SKIN FIBROBLAST |
| 257 | Digestive | BSS01595 | SMALL INTESTINE | 544 | Kidney | BSS01099 | KIDNEY | 831 | Stromal | BSS00276 | SKIN FIBROBLAST |
| 258 | Digestive | BSS01596 | SMALL INTESTINE | 545 | Kidney | BSS01514 | KIDNEY | 832 | Stromal | BSS00393 | SKIN FIBROBLAST |
| 259 | Digestive | BSS01590 | SMALL INTESTINE | 546 | Kidney | BSS01134 | KIDNEY | 833 | Stromal | BSS00394 | SKIN FIBROBLAST |
| 260 | Digestive | BSS01591 | SMALL INTESTINE | 547 | Kidney | BSS01515 | KIDNEY | 834 | Stromal | BSS00063 | SKIN FIBROBLAST |
| 261 | Digestive | BSS01592 | SMALL INTESTINE | 548 | Kidney | BSS01131 | KIDNEY | 835 | Stromal | BSS00069 | SKIN FIBROBLAST |
| 262 | Digestive | BSS01593 | SMALL INTESTINE | 549 | Kidney | BSS01079 | KIDNEY | 836 | Stromal | BSS00346 | SKIN FIBROBLAST |
| 263 | Digestive | BSS01594 | SMALL INTESTINE | 550 | Kidney | BSS01096 | KIDNEY | 837 | Stromal | BSS01583 | SKIN FIBROBLAST |
| 264 | Digestive | BSS01603 | SMALL INTESTINE | 551 | Kidney | BSS01097 | KIDNEY | 838 | Stromal | BSS00278 | SKIN FIBROBLAST |
| 265 | Digestive | BSS01604 | SMALL INTESTINE | 552 | Kidney | BSS01088 | KIDNEY | 839 | Stromal | BSS00390 | SKIN FIBROBLAST |
| 266 | Digestive | BSS01600 | SMALL INTESTINE | 553 | Kidney | BSS00528 | KIDNEY CELL | 840 | Stromal | BSS00066 | SKIN FIBROBLAST |
| 267 | Digestive | BSS01602 | SMALL INTESTINE | 554 | Kidney | BSS00526 | KIDNEY CELL | 841 | Stromal | BSS00061 | SKIN FIBROBLAST |
| 268 | Digestive | BSS01597 | SMALL INTESTINE | 555 | Kidney | BSS01484 | RENAL CORTEX INTERSTITIUM | 842 | Stromal | BSS00068 | SKIN FIBROBLAST |
| 269 | Digestive | BSS01599 | SMALL INTESTINE | 556 | Kidney | BSS01485 | RENAL CORTEX INTERSTITIUM | 843 | Stromal | BSS00476 | SKIN FIBROBLAST |
| 270 | Digestive | BSS01588 | SMALL INTESTINE | 557 | Kidney | BSS01482 | RENAL CORTEX INTERSTITIUM | 844 | Stromal | BSS00113 | SKIN FIBROBLAST |
| 271 | Digestive | BSS01601 | SMALL INTESTINE | 558 | Kidney | BSS01483 | RENAL CORTEX INTERSTITIUM | 845 | Thymus | BSS01824 | THYMUS |
| 272 | Digestive | BSS01637 | STOMACH | 559 | Kidney | BSS01489 | RENAL CORTEX INTERSTITIUM | 846 | Thymus | BSS01819 | THYMUS |
| 273 | Digestive | BSS01642 | STOMACH | 560 | Kidney | BSS01490 | RENAL CORTEX INTERSTITIUM | 847 | Thymus | BSS01821 | THYMUS |
| 274 | Digestive | BSS01643 | STOMACH | 561 | Kidney | BSS01150 | RENAL CORTEX INTERSTITIUM | 848 | Thymus | BSS01823 | THYMUS |
| 275 | Digestive | BSS01644 | STOMACH | 562 | Kidney | BSS01531 | RENAL CORTEX INTERSTITIUM | 849 | Thymus | BSS01818 | THYMUS |
| 276 | Digestive | BSS01646 | STOMACH | 563 | Kidney | BSS01486 | RENAL CORTEX INTERSTITIUM | 850 | Thymus | BSS01826 | THYMUS |
| 277 | Digestive | BSS01647 | STOMACH | 564 | Kidney | BSS01487 | RENAL CORTEX INTERSTITIUM | 851 | Thymus | BSS01827 | THYMUS |
| 278 | Digestive | BSS01641 | STOMACH | 565 | Kidney | BSS01151 | RENAL CORTEX INTERSTITIUM | 852 | Thymus | BSS01828 | THYMUS |
| 279 | Digestive | BSS01658 | STOMACH | 566 | Kidney | BSS01532 | RENAL CORTEX INTERSTITIUM | 853 | Thymus | BSS01829 | THYMUS |
| 280 | Digestive | BSS01655 | STOMACH | 567 | Kidney | BSS01488 | RENAL CORTEX INTERSTITIUM | 854 | Thymus | BSS01820 | THYMUS |
| 281 | Digestive | BSS01656 | STOMACH | 568 | Kidney | BSS01495 | RENAL PELVIS | 855 | Thymus | BSS01825 | THYMUS |
| 282 | Digestive | BSS01657 | STOMACH | 569 | Kidney | BSS01496 | RENAL PELVIS | 856 | Urinary | BSS01878 | URINARY BLADDER |
| 283 | Digestive | BSS01636 | STOMACH | 570 | Kidney | BSS01493 | RENAL PELVIS | 857 | Urinary | BSS01876 | URINARY BLADDER |
| 284 | Digestive | BSS01638 | STOMACH | 571 | Kidney | BSS01494 | RENAL PELVIS | 858 | Urinary | BSS01879 | UROTHELIUM CELL |
| 285 | Digestive | BSS01650 | STOMACH | 572 | Kidney | BSS01500 | RENAL PELVIS | 859 | Urinary | BSS01880 | UROTHELIUM CELL |
| 286 | Digestive | BSS01653 | STOMACH | 573 | Kidney | BSS01501 | RENAL PELVIS | | | | |
| 287 | Digestive | BSS01649 | STOMACH | 574 | Kidney | BSS01152 | RENAL PELVIS | | | | |

**Figure S1:** (continued, 4 of 4)

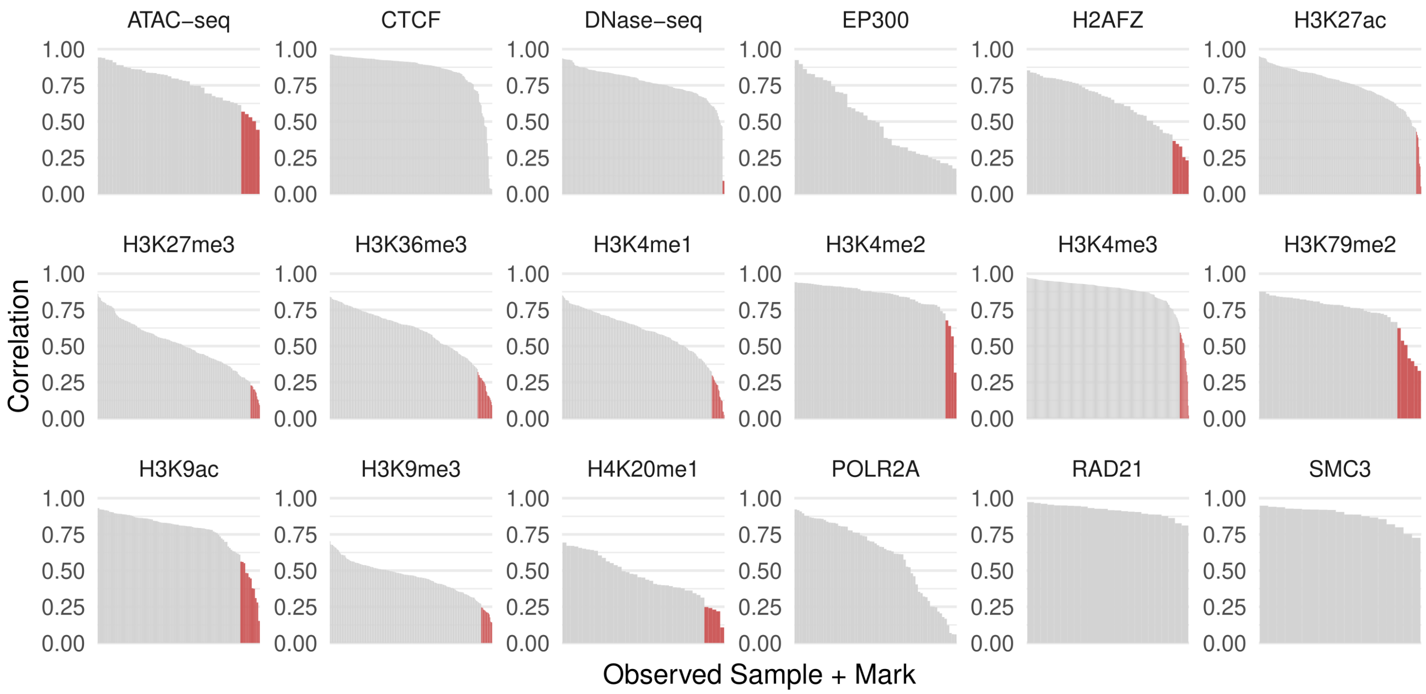


**Figure S2:** **(A)** Track agreement between observed and imputed in the Tier 1 marks, where both are available, by genome wide correlation. Top (green) and bottom (red) tracks are labeled according to automated elbow discovery in the ranked list.

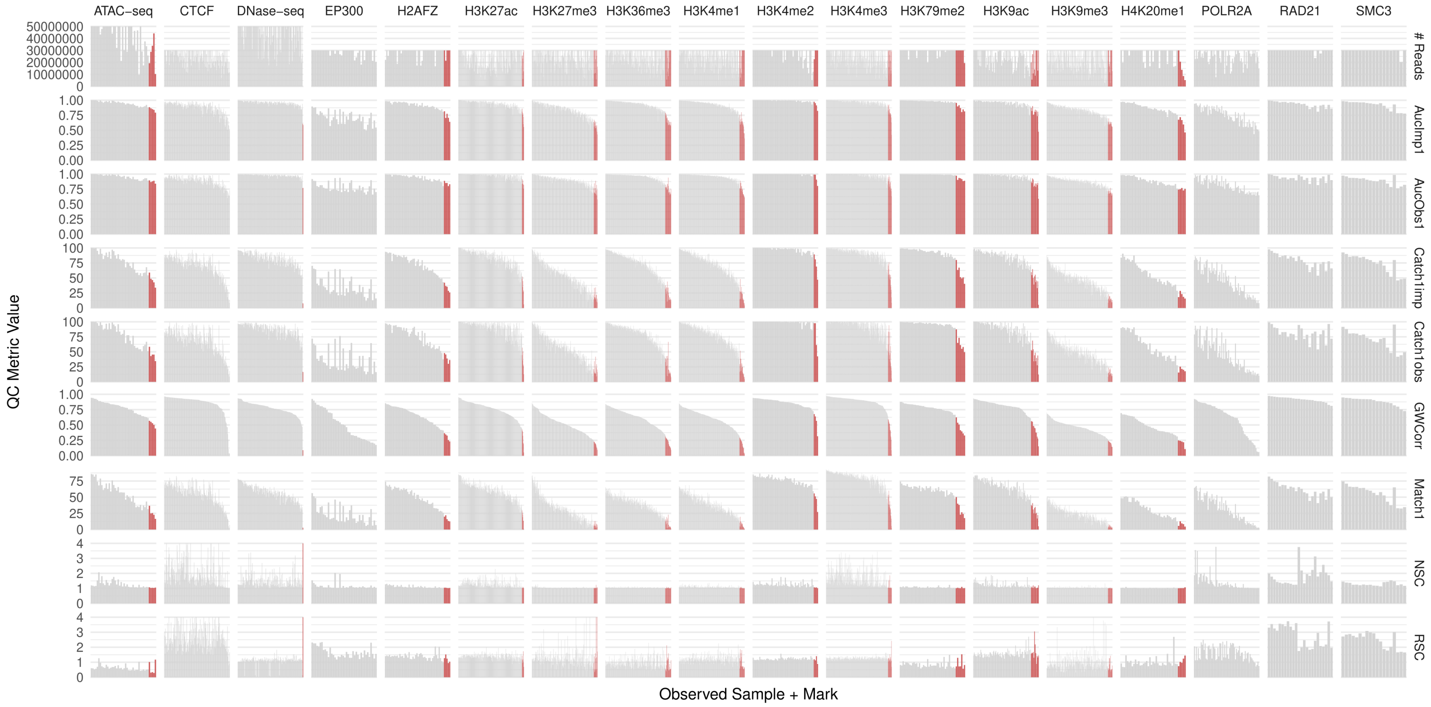


**Figure S2: (B)** All imputation metrics (from original ChromImpute paper: correlation, AUC, and peak recovery) and QC metrics (reads, NSC, RSC) across all 18 imputed Tier 1,2, and 3 assays, ordered according to correlation between imputed and observed within each assay. Bottom tracks for Tier 1 and 2 assays are labeled in red, as above.

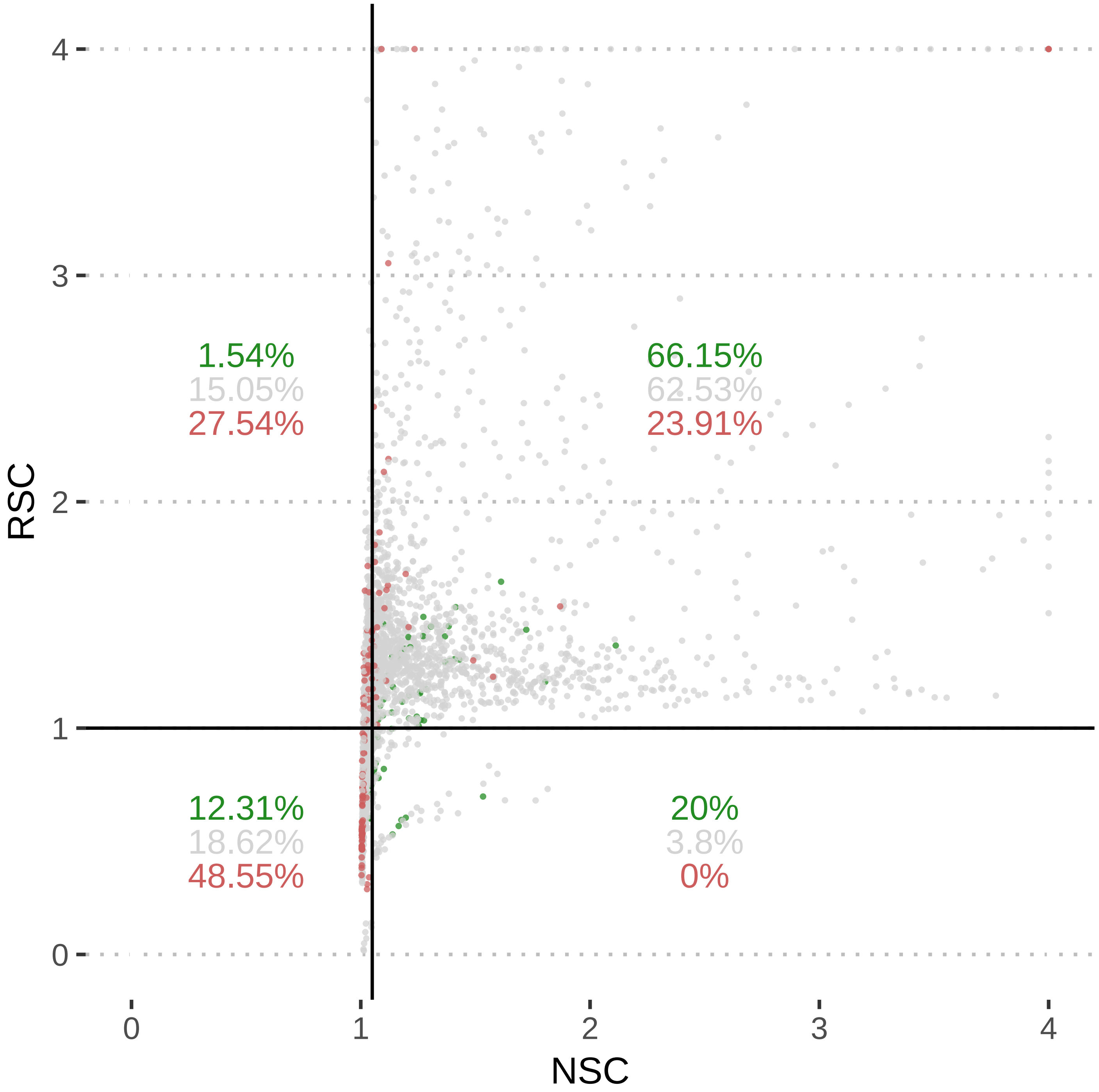


**Figure S3:** Imputation QC metrics reflect external ChIP-seq quality metrics (NSC and RSC). Normalized Strand Cross-correlation coefficient (NSC) against Relative Strand Cross-correlation coefficient (RSC) for observed tracks. Top (green) and bottom (red) tracks are labeled according to automated elbow discovery in imputed vs. observed correlation ranked list. Poor agreement tracks are strongly clustered in lower left quadrant of tracks failing QC. Critical values subdivide the plot for NSC (1.05) and RSC (1).

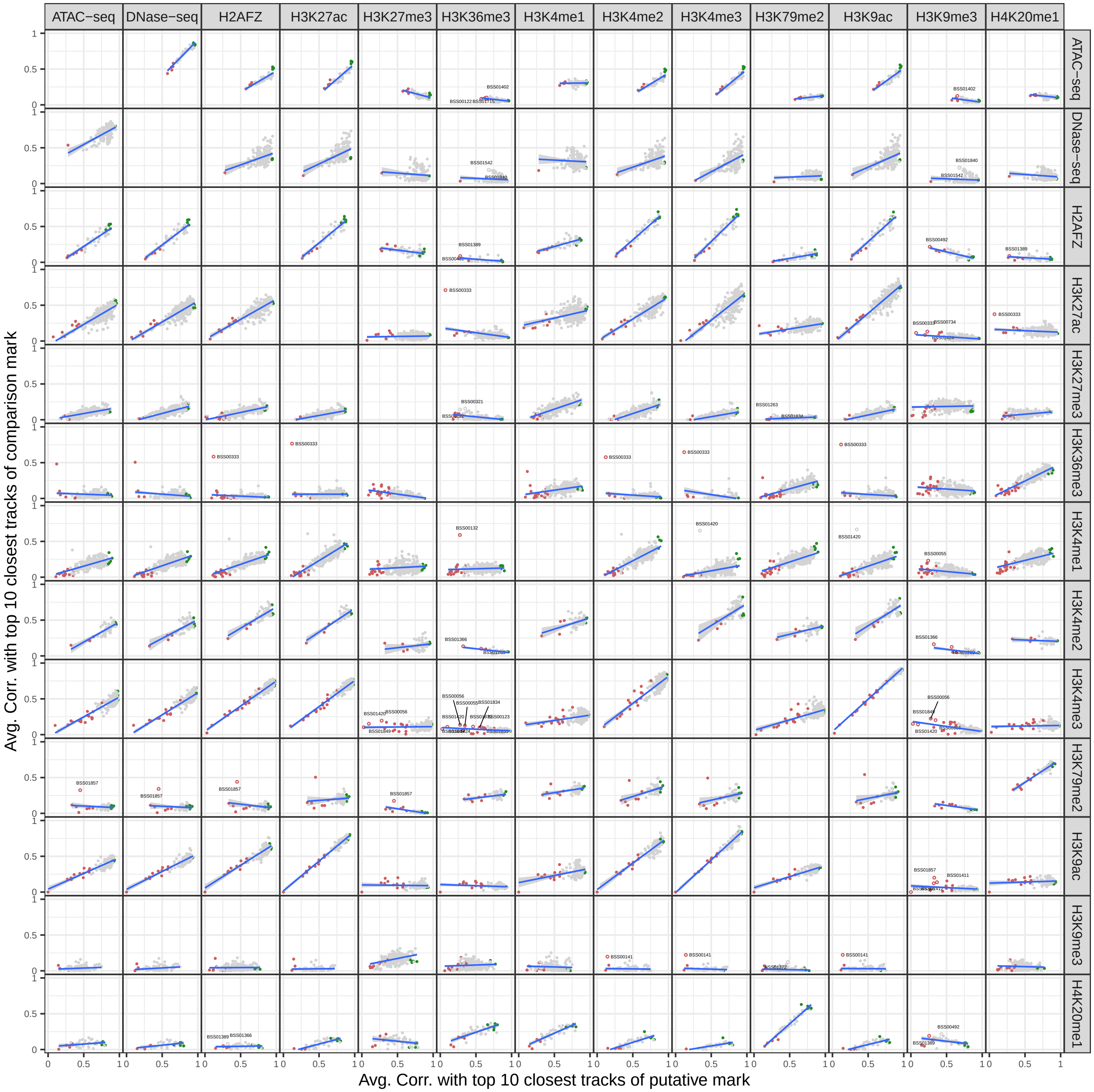


**Figure S4:** Imputed-observed agreement can be used to systematically flag antibody swaps. Each panel shows the average correlation with closest 10 tracks within the putative mark (each row) for each observed dataset against all other Tier 1 and 2 marks and assays. We used the overall mark-mark trend to flag outliers for visual inspection.

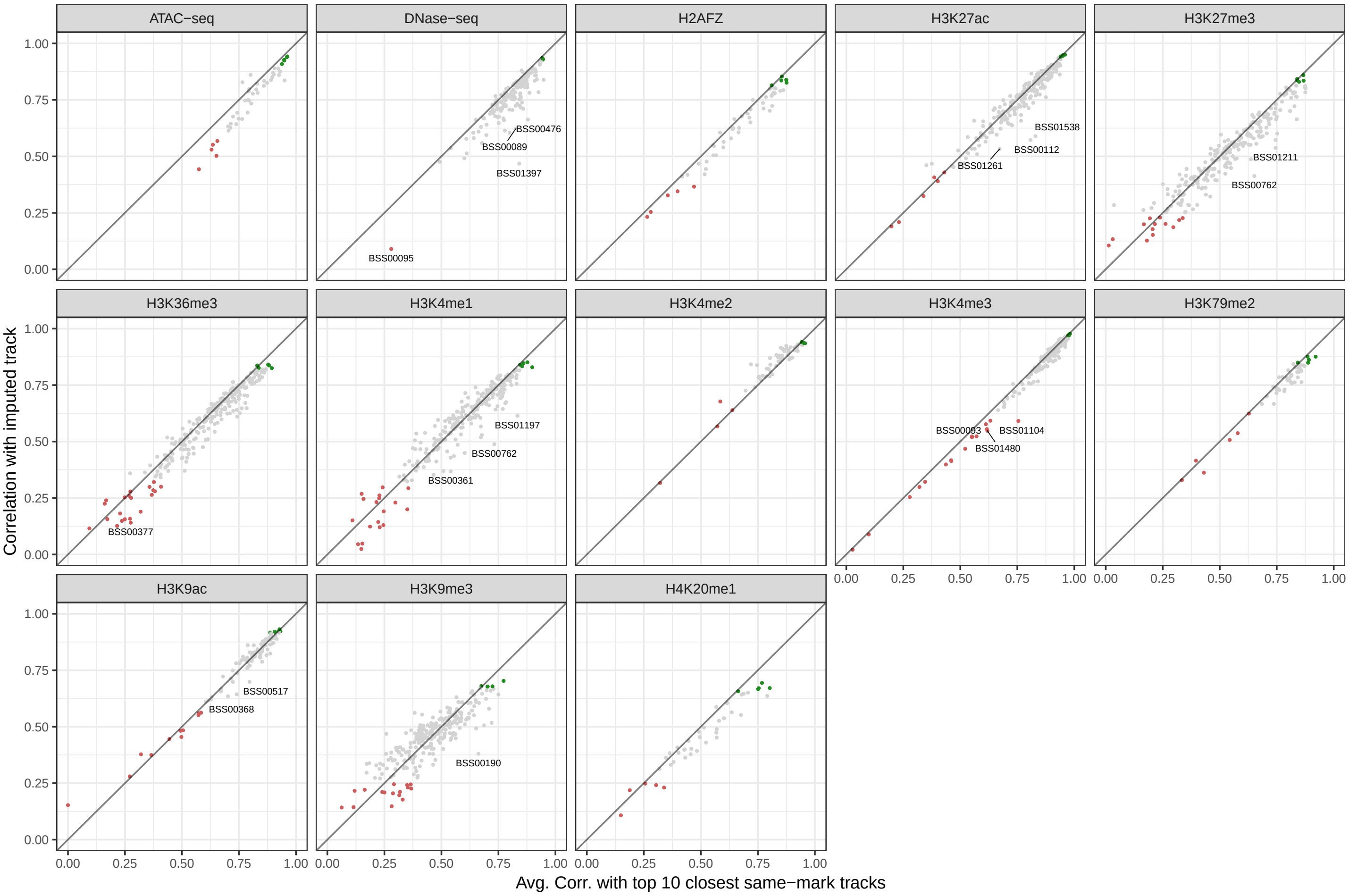


**Figure S5:** Imputed-observed agreement can be used to flag sample swaps. We compare the imputed-observed correlation to the average imputed-observed correlation within the top 10 closest samples for the putative mark and flagged outliers for visual inspection (after removing antibody swaps).

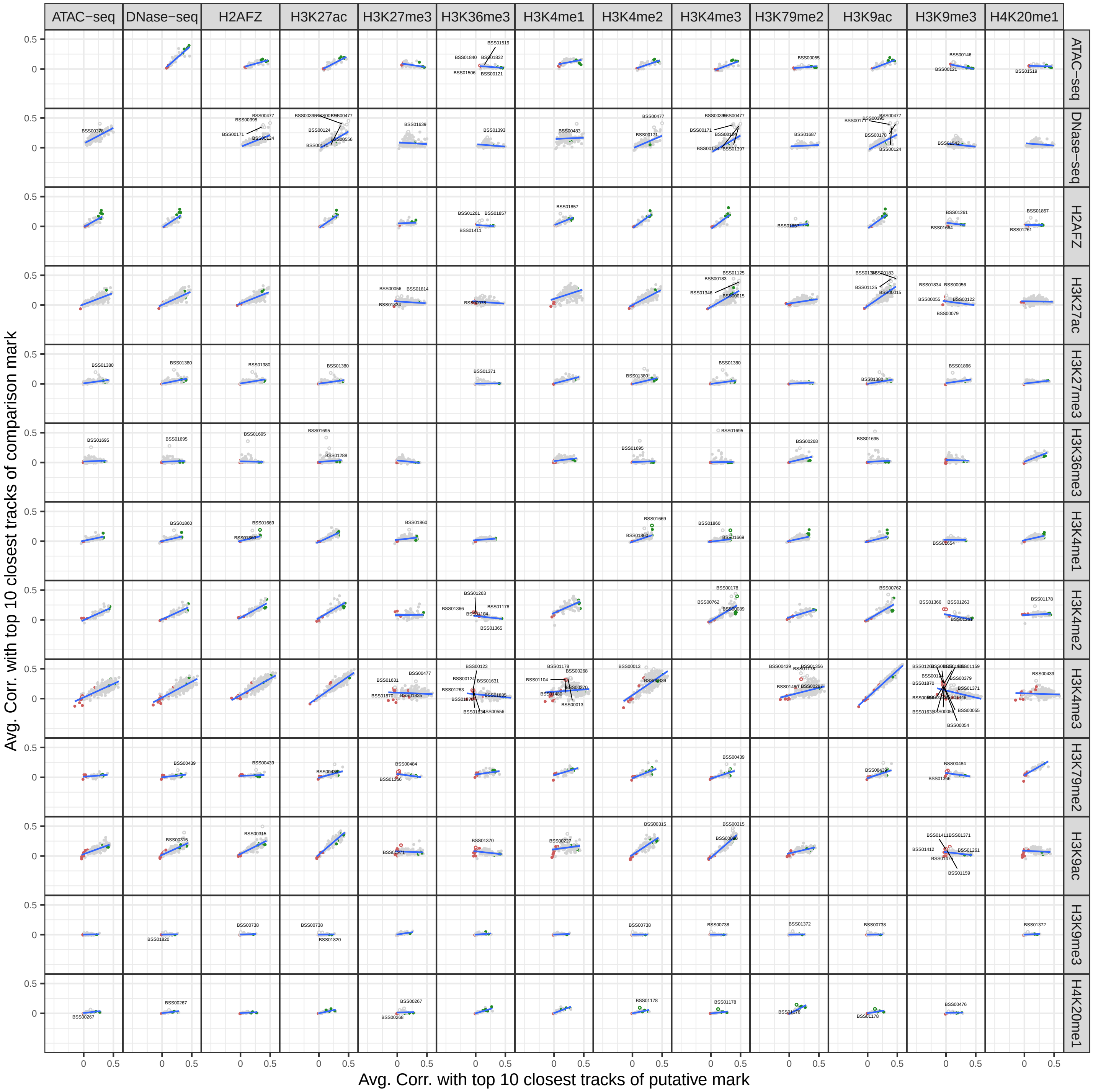


**Figure S6:** Imputed-observed agreement can be used to systematically flag secondary reactivities of antibodies. Each panel shows the average correlation of the difference between a pair of observed and imputed datasets with closest 10 tracks within the putative mark (each row) for each dataset against all other Tier 1 and 2 marks and assays. As with antibody swaps, we used the overall mark-mark trend to flag outliers for visual inspection.

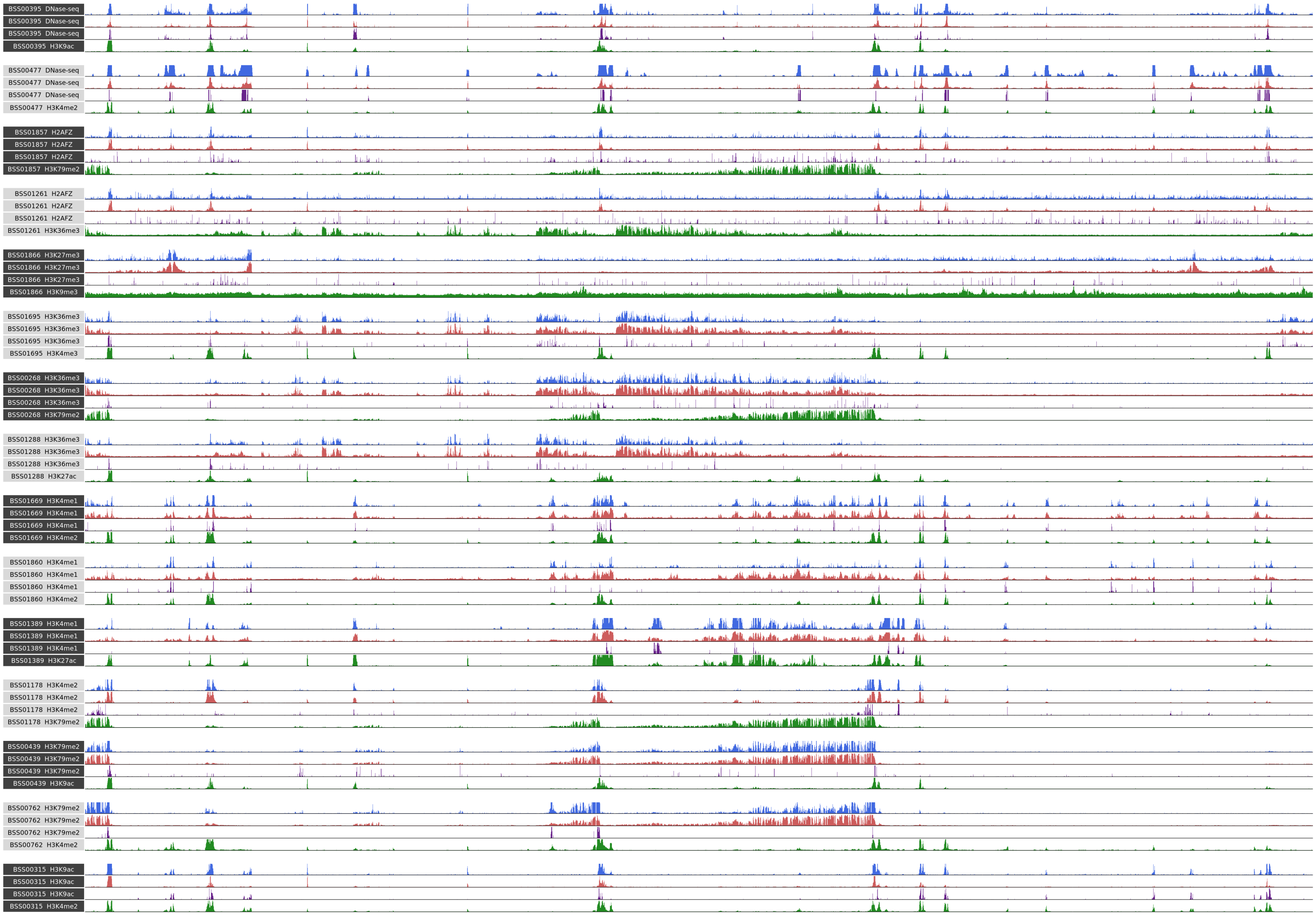


**Figure S7:** We can use imputation to identify antibodies with secondary reactivities. Track sets for 15 flagged samples highlight a disagreement from secondary reactivity (or sample swapping of one experiment of many), where the difference between observed and imputed best correlate with an external histone mark or assay. Each track set shows the Difference (purple) from Observed (blue) to Imputed (red) and the best Match (green) to the difference (a mark in the same sample) in chromosome 1 from 1.5Mb to 2Mb (chosen to show a range of diverse elements and marks).

| | id | mark | (low quality tracks) | | | | | |
|---|---|---|---|---|---|---|---|---|
| 1 | BSS01365 | H3K4me2 | 47 | BSS01857 | H3K9ac | 93 | BSS00734 | H3K27ac |
| 2 | BSS01104 | H3K4me2 | 48 | BSS01832 | H3K36me3 | 94 | BSS01715 | H3K9me3 |
| 3 | BSS01365 | H3K79me2 | 49 | BSS01841 | H3K36me3 | 95 | BSS00325 | H3K9me3 |
| 4 | BSS00556 | H3K4me3 | 50 | BSS01424 | H3K4me3 | 96 | BSS01850 | H3K27me3 |
| 5 | BSS01104 | H3K4me3 | 51 | BSS00055 | H3K4me1 | 97 | BSS01407 | H3K27me3 |
| 6 | BSS01263 | H3K4me3 | 52 | BSS00087 | H3K4me1 | 98 | BSS00381 | H3K4me1 |
| 7 | BSS01815 | ATAC-seq | 53 | BSS00080 | H3K4me1 | 99 | BSS00395 | H3K27me3 |
| 8 | BSS01263 | H3K4me2 | 54 | BSS00325 | H3K36me3 | 100 | BSS00381 | H3K9me3 |
| 9 | BSS01475 | H3K9ac | 55 | BSS00556 | H3K36me3 | 101 | BSS00321 | H3K4me1 |
| 10 | BSS01196 | H3K9ac | 56 | BSS01340 | H3K9ac | 102 | BSS01424 | H3K27ac |
| 11 | BSS00093 | H3K4me3 | 57 | BSS00284 | H3K36me3 | 103 | BSS00124 | H3K36me3 |
| 12 | BSS01506 | ATAC-seq | 58 | BSS00055 | H3K36me3 | 104 | BSS01507 | H3K27me3 |
| 13 | BSS01667 | H3K9ac | 59 | BSS00281 | H3K4me1 | 105 | BSS01715 | H3K36me3 |
| 14 | BSS01480 | H3K4me3 | 60 | BSS01543 | H3K36me3 | 106 | BSS01209 | H3K27me3 |
| 15 | BSS00484 | H3K79me2 | 61 | BSS01835 | H3K4me1 | 107 | BSS01832 | H3K9me3 |
| 16 | BSS01715 | ATAC-seq | 62 | BSS01080 | H3K36me3 | 108 | BSS01850 | H3K36me3 |
| 17 | BSS00123 | H3K4me3 | 63 | BSS00056 | H3K4me3 | 109 | BSS01411 | H3K36me3 |
| 18 | BSS01834 | H3K4me3 | 64 | BSS01389 | H2AFZ | 110 | BSS01370 | H3K36me3 |
| 19 | BSS01835 | H3K4me3 | 65 | BSS00493 | H3K36me3 | 111 | BSS01366 | H3K9ac |
| 20 | BSS01366 | H3K79me2 | 66 | BSS01835 | H3K36me3 | 112 | BSS01870 | H3K27me3 |
| 21 | BSS01402 | ATAC-seq | 67 | BSS00284 | H3K4me1 | 113 | BSS01834 | H3K4me1 |
| 22 | BSS01371 | H3K9ac | 68 | BSS01389 | H4K20me1 | 114 | BSS01887 | H3K36me3 |
| 23 | BSS01370 | H3K9ac | 69 | BSS00054 | H3K4me1 | 115 | BSS01831 | H3K9me3 |
| 24 | BSS00284 | H3K4me3 | 70 | BSS00122 | H3K9me3 | 116 | BSS01630 | H3K4me1 |
| 25 | BSS00197 | H3K9ac | 71 | BSS00123 | H3K9me3 | 117 | BSS01459 | H3K9me3 |
| 26 | BSS01341 | H3K9ac | 72 | BSS01850 | H3K9me3 | 118 | BSS01424 | H3K9me3 |
| 27 | BSS00122 | ATAC-seq | 73 | BSS01213 | H4K20me1 | 119 | BSS00377 | H3K36me3 |
| 28 | BSS00055 | H3K27ac | 74 | BSS00093 | H3K36me3 | 120 | BSS00159 | H3K27me3 |
| 29 | BSS01870 | H3K4me3 | 75 | BSS00132 | H3K4me1 | 121 | BSS01507 | H3K4me1 |
| 30 | BSS01412 | H3K79me2 | 76 | BSS00492 | H2AFZ | 122 | BSS01837 | H3K27me3 |
| 31 | BSS00141 | H3K4me3 | 77 | BSS01876 | H3K4me1 | 123 | BSS01426 | H3K36me3 |
| 32 | BSS00074 | H3K27ac | 78 | BSS01370 | H3K9me3 | 124 | BSS01837 | H3K4me1 |
| 33 | BSS00521 | H3K4me3 | 79 | BSS00439 | H4K20me1 | 125 | BSS01426 | H3K4me1 |
| 34 | BSS01857 | H3K79me2 | 80 | BSS00325 | H3K4me1 | 126 | BSS01459 | H3K36me3 |
| 35 | BSS01866 | H3K27ac | 81 | BSS01178 | H3K27me3 | 127 | BSS00484 | H4K20me1 |
| 36 | BSS01412 | H3K9ac | 82 | BSS00141 | H3K9me3 | 128 | BSS00531 | H3K27me3 |
| 37 | BSS01411 | H3K9ac | 83 | BSS01849 | H3K27me3 | 129 | BSS00333 | H3K36me3 |
| 38 | BSS00478 | H2AFZ | 84 | BSS01543 | H3K9me3 | 130 | BSS00093 | H3K27me3 |
| 39 | BSS00556 | H3K79me2 | 85 | BSS00284 | H3K27me3 | 131 | BSS01562 | DNase-seq |
| 40 | BSS00558 | H2AFZ | 86 | BSS00054 | H3K36me3 | 132 | BSS00095 | DNase-seq |
| 41 | BSS01411 | H3K79me2 | 87 | BSS00493 | H3K9me3 | 133 | BSS01420 | H3K4me3 |
| 42 | BSS00477 | H2AFZ | 88 | BSS00341 | H4K20me1 | 134 | BSS00333 | H3K27ac |
| 43 | BSS00281 | H3K27ac | 89 | BSS00325 | H3K27me3 | 135 | BSS01849 | H3K4me1 |
| 44 | BSS00055 | H3K4me3 | 90 | BSS00074 | H3K9me3 | 136 | BSS01519 | H3K4me1 |
| 45 | BSS00378 | H3K36me3 | 91 | BSS01426 | H3K9me3 | 137 | BSS01543 | H3K4me1 |
| 46 | BSS01366 | H3K4me2 | 92 | BSS00483 | H3K9me3 | 138 | BSS01849 | H3K4me3 |

| | id | mark | potential.abswap |
|---|---|---|---|
| 1 | BSS00333 | H3K36me3 | H3K27ac |
| 2 | BSS00333 | H3K27ac | H3K36me3 |
| 3 | BSS01857 | H3K9ac | H3K79me2 |
| 4 | BSS01857 | H3K79me2 | H3K9ac |
| 5 | BSS00141 | H3K9me3 | H3K9ac |
| 6 | BSS00132 | H3K4me1 | H3K36me3 |
| 7 | BSS01420 | H3K4me1 | H3K4me3 |

| | id | mark | potential.sampswap |
|---|---|---|---|
| 1 | BSS00089 | DNase-seq | BSS00339 |
| 2 | BSS00095 | DNase-seq | BSS01452 |
| 3 | BSS00476 | DNase-seq | BSS01397 |
| 4 | BSS01397 | DNase-seq | BSS00334 |
| 5 | BSS00112 | H3K27ac | BSS01366 |
| 6 | BSS01261 | H3K27ac | BSS00387 |
| 7 | BSS01538 | H3K27ac | BSS00703 |
| 8 | BSS00762 | H3K27me3 | BSS00547 |
| 9 | BSS01211 | H3K27me3 | BSS01224 |
| 10 | BSS00377 | H3K36me3 | BSS01715 |
| 11 | BSS00361 | H3K4me1 | BSS00365 |
| 12 | BSS00762 | H3K4me1 | BSS01038 |
| 13 | BSS01197 | H3K4me1 | BSS01144 |
| 14 | BSS00093 | H3K4me3 | BSS00702 |
| 15 | BSS01104 | H3K4me3 | BSS01365 |
| 16 | BSS01480 | H3K4me3 | BSS00702 |
| 17 | BSS00368 | H3K9ac | BSS00207 |
| 18 | BSS00517 | H3K9ac | BSS00502 |
| 19 | BSS00190 | H3K9me3 | BSS00196 |

| | id | mark | potential.secondary |
|---|---|---|---|
| 1 | BSS00395 | DNase-seq | H3K9ac |
| 2 | BSS00477 | DNase-seq | H3K4me2 |
| 3 | BSS01857 | H2AFZ | H3K79me2 |
| 4 | BSS01261 | H2AFZ | H3K36me3 |
| 5 | BSS01866 | H3K27me3 | H3K9me3 |
| 6 | BSS01695 | H3K36me3 | H3K4me3 |
| 7 | BSS00268 | H3K36me3 | H3K79me2 |
| 8 | BSS01288 | H3K36me3 | H3K27ac |
| 9 | BSS01669 | H3K4me1 | H3K4me2 |
| 10 | BSS01860 | H3K4me1 | H3K4me2 |
| 11 | BSS01389 | H3K4me1 | H3K27ac |
| 12 | BSS01178 | H3K4me2 | H3K79me2 |
| 13 | BSS00439 | H3K79me2 | H3K9ac |
| 14 | BSS00762 | H3K79me2 | H3K4me2 |
| 15 | BSS00315 | H3K9ac | H3K4me2 |

**Figure S8:** Table of flagged samples. **(left)** low agreement tracks, **(right, top)** potential antibody swaps, **(right, middle)** potential sample swaps, **(right, bottom)** potential secondary antibody reactivities or single replicate or experiment swaps.

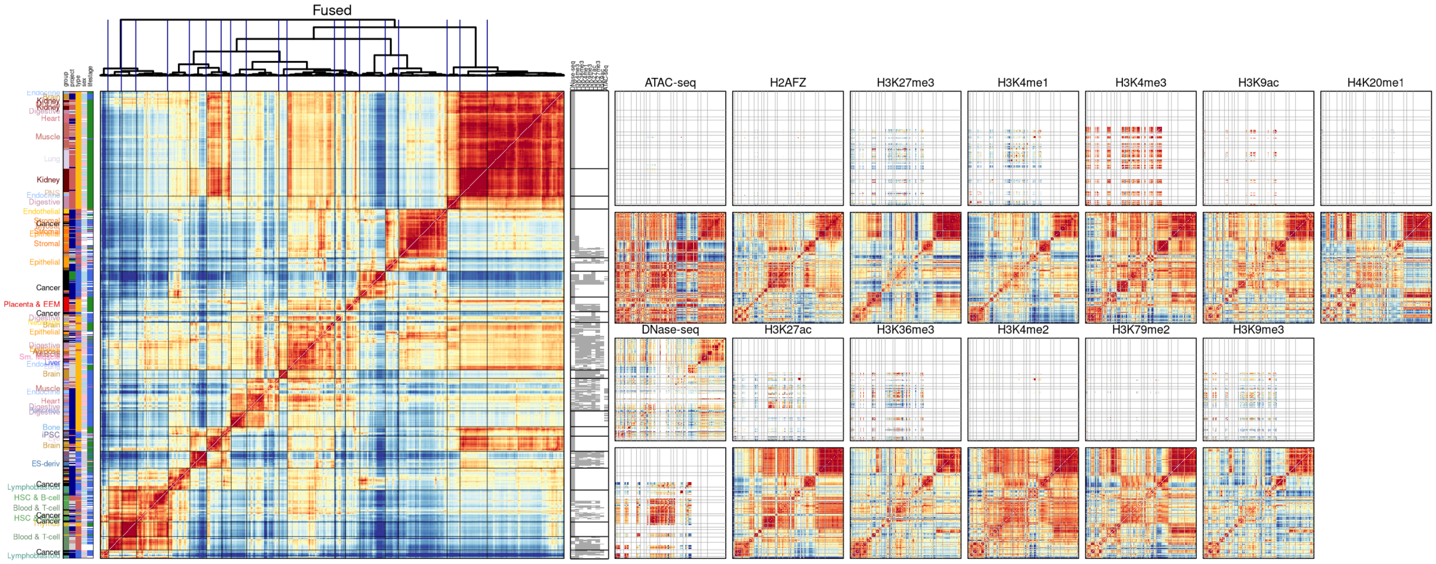


**Figure S9:** Hierarchically clustered genome-wide correlation across samples in all 13 imputed Tier 1 and 2 assays. Observed (top) vs. imputed (bottom) matrices shown. Clustering conducted on fused matrix (left panel, constructed as in main figure). Observed data availability matrix (grey is available, white is unavailable) is shown for the top nine marks and accessibility assays by number of observed datasets

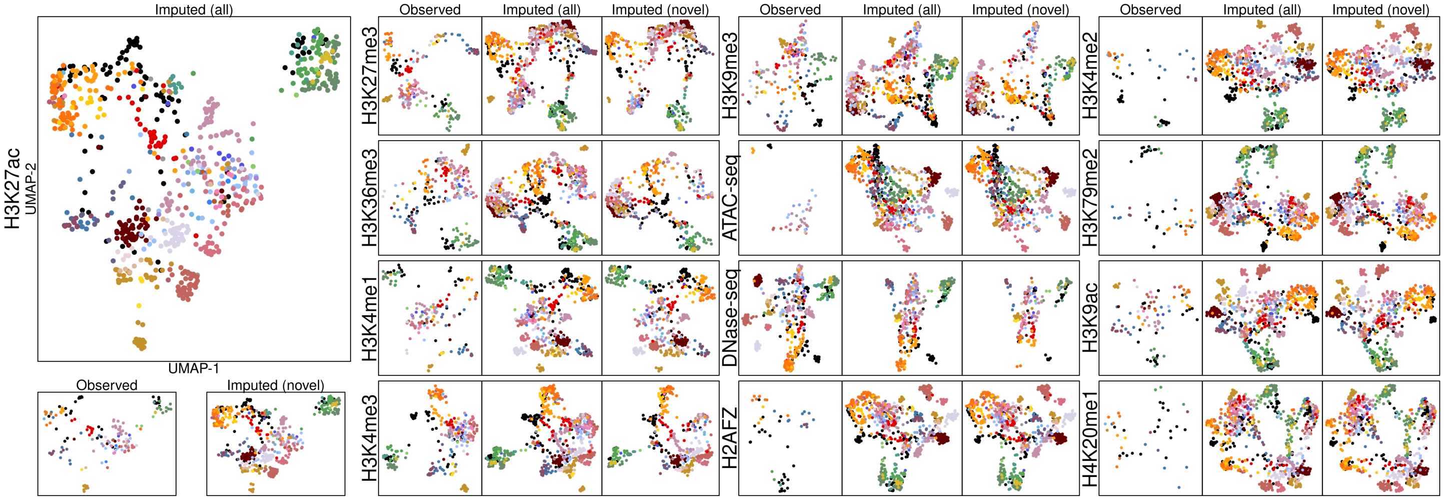


**Figure S10:** Joint UMAP embeddings of observed and imputed data within each Tier 1 and 2 mark/assay. Separately, observed, imputed samples, and all novel imputed tracks are plotted for each mark/assay and colored according to tissue group. Imputed UMAP highlights differences in cell types: H3K27ac clusters hematopoietic cells and tissues (in green) closely, reflecting lineage, whereas H3K27me3 clusters iPSC, ESCs, and derived cells (in purple/blue), reflecting differentiation stage. UMAP embedding was calculated from spearman correlation of tracks within regions marked by mark or assay relevant states within the Roadmap compendium.

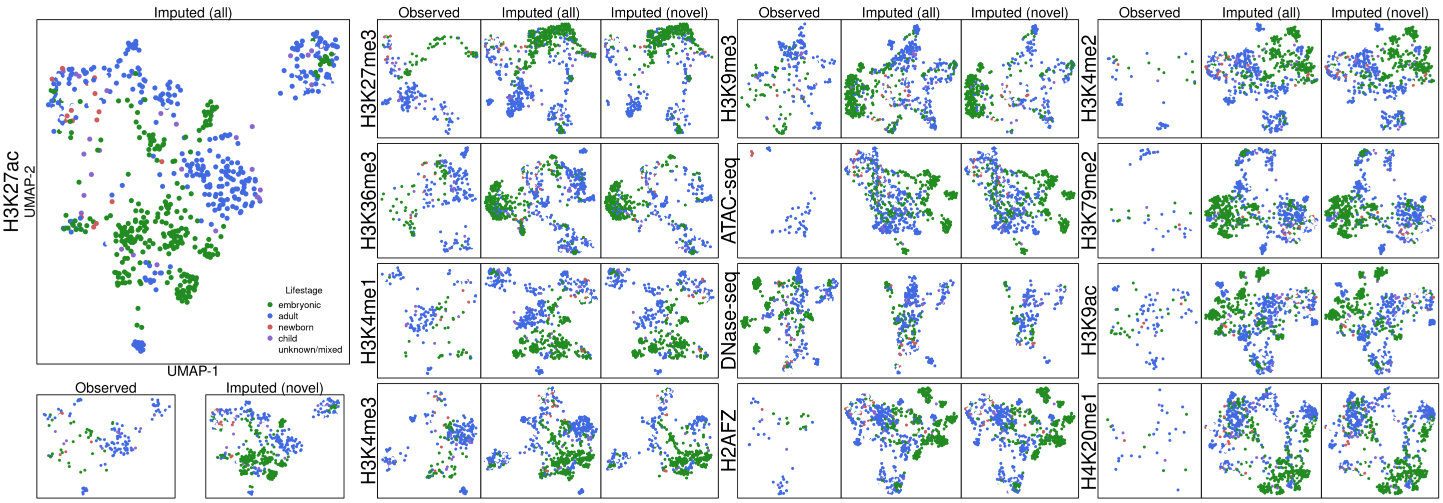


**Figure S11: (A)** Joint UMAP embeddings of observed and imputed data, with observed and imputed data (points) labeled by the sample's biological lifestage.

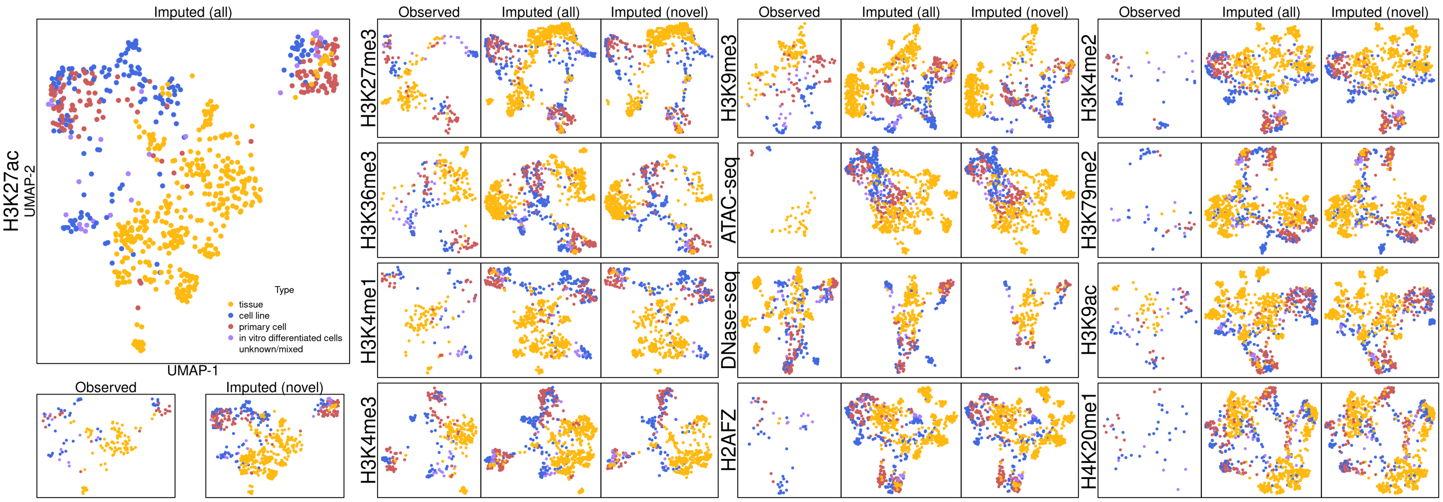


**Figure S11: (B)** Joint UMAP embeddings of observed and imputed data, with observed and imputed data (points) labeled by the sample type.

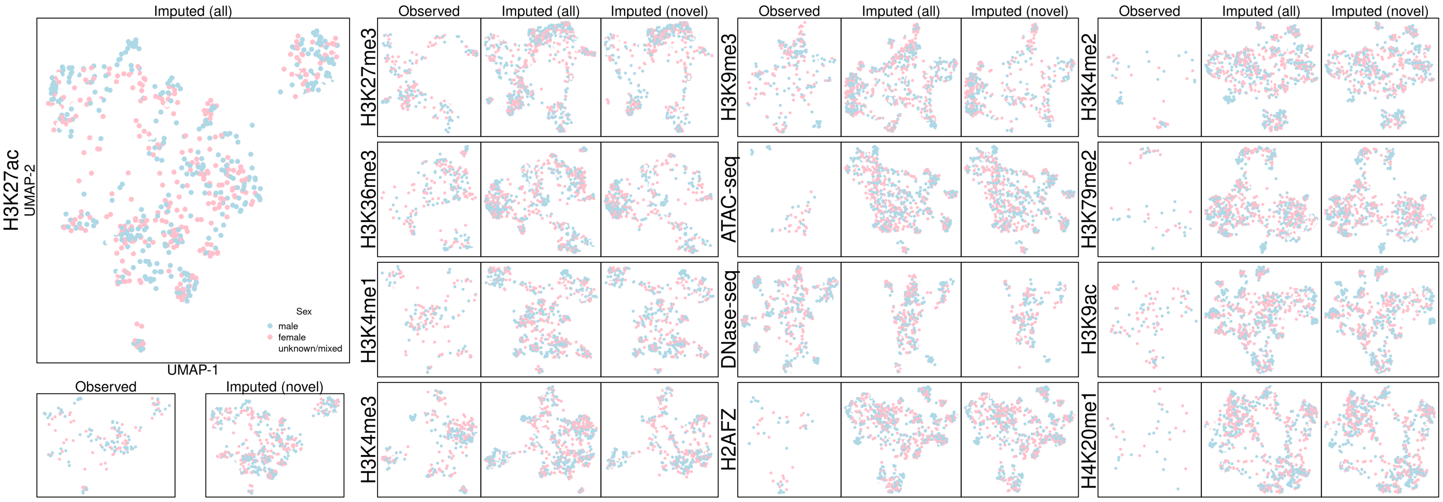


**Figure S11: (C)** Joint UMAP embeddings of observed and imputed data, with observed and imputed data (points) labeled by the sample biological sex.

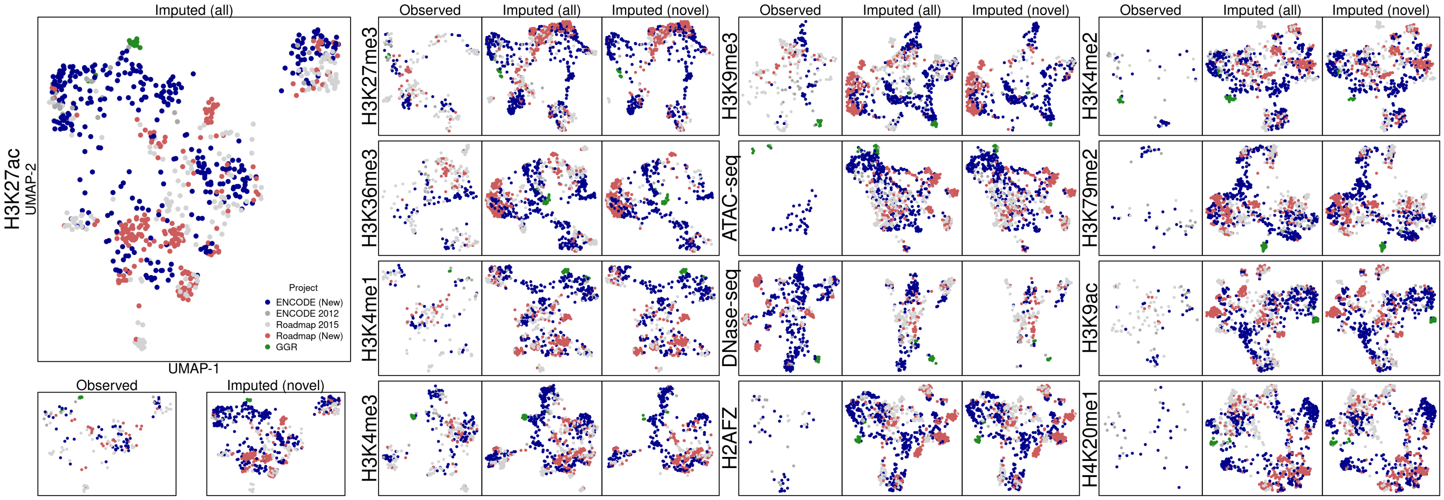


**Figure S11: (D)** Joint UMAP embeddings of observed and imputed data, with observed and imputed data (points) labeled by the epigenomic reference project of origin.

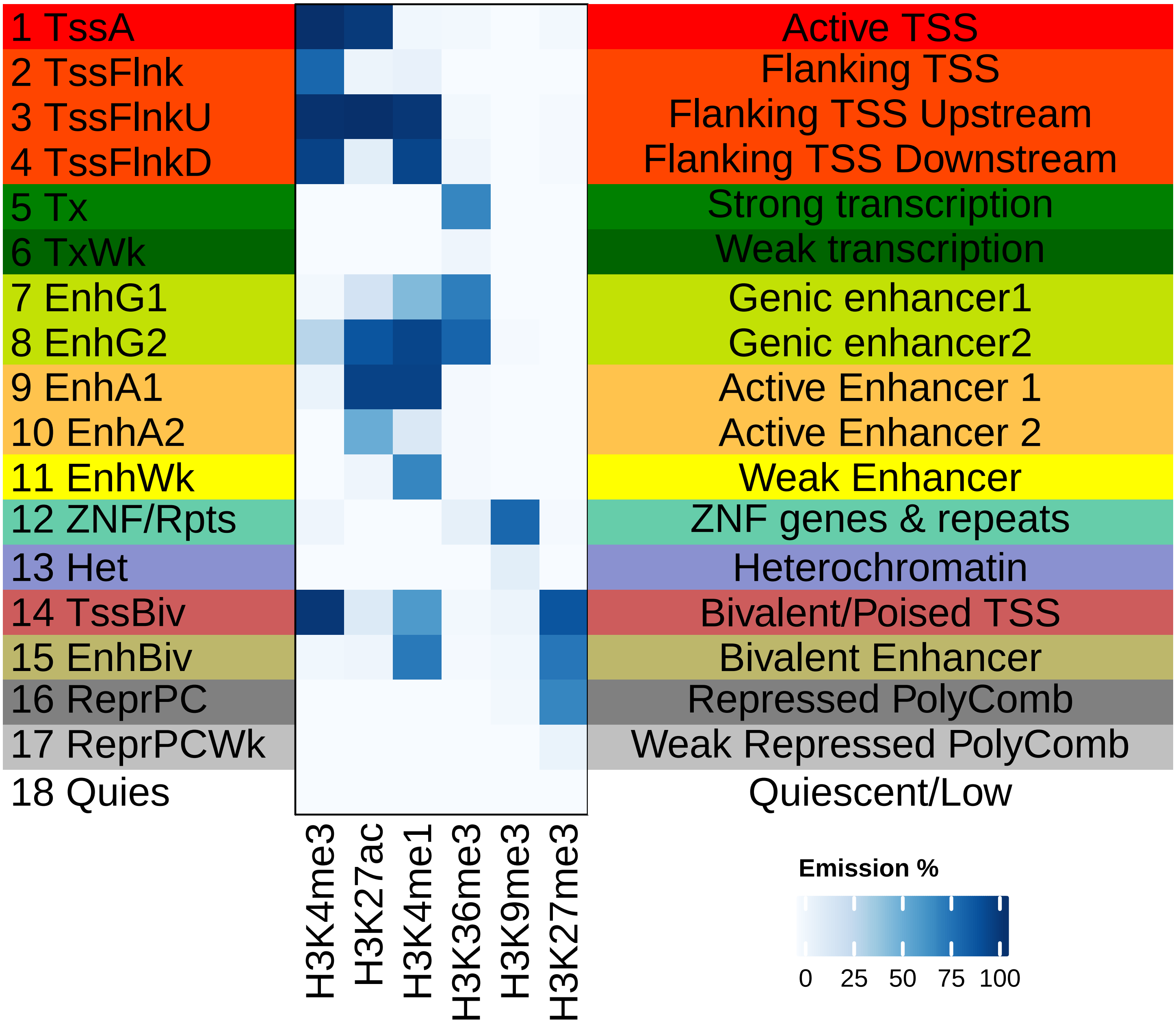


**Figure S12:** Epigenomic state mnemonics for ChromHMM 18-state model (left) with emissions matrix (center) and state definitions (right). The 18-state model was trained on Roadmap data for the Roadmap 2015 paper.

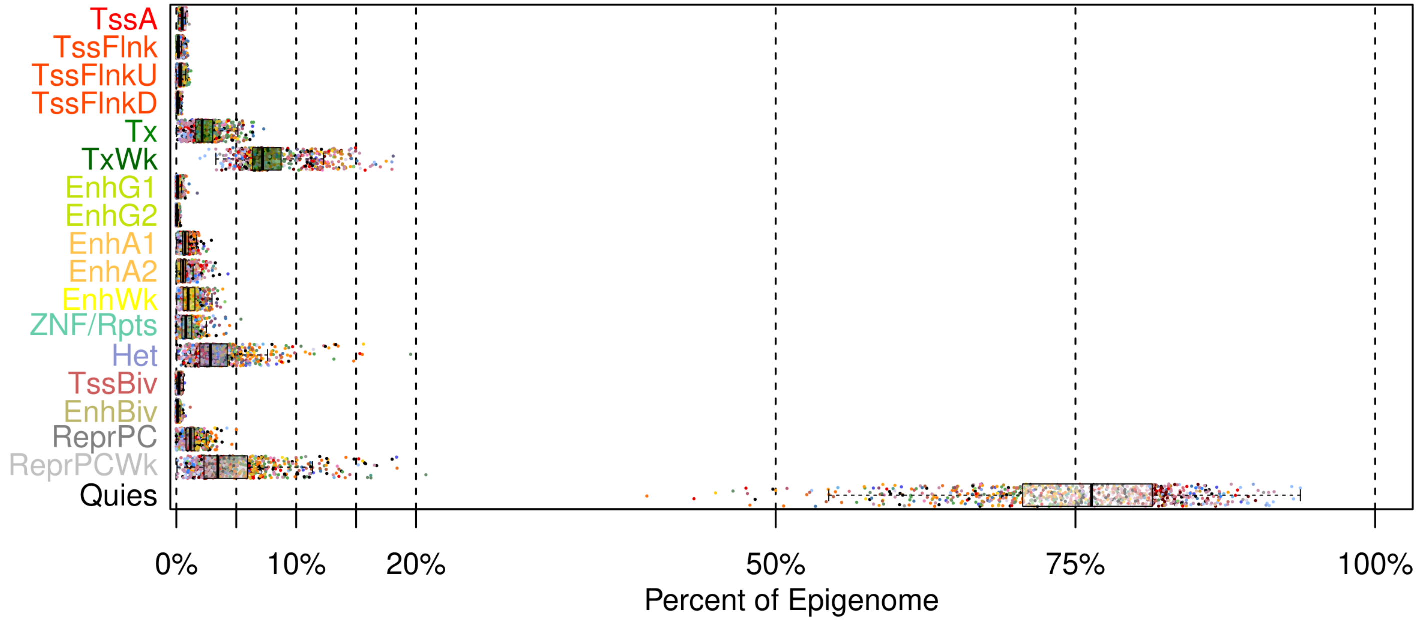


**Figure S13:** Distributions of per-state epigenome fractions (boxplots) across 833 epigenomes (points, colored by tissue group) according to the ChromHMM 18-state model annotations.

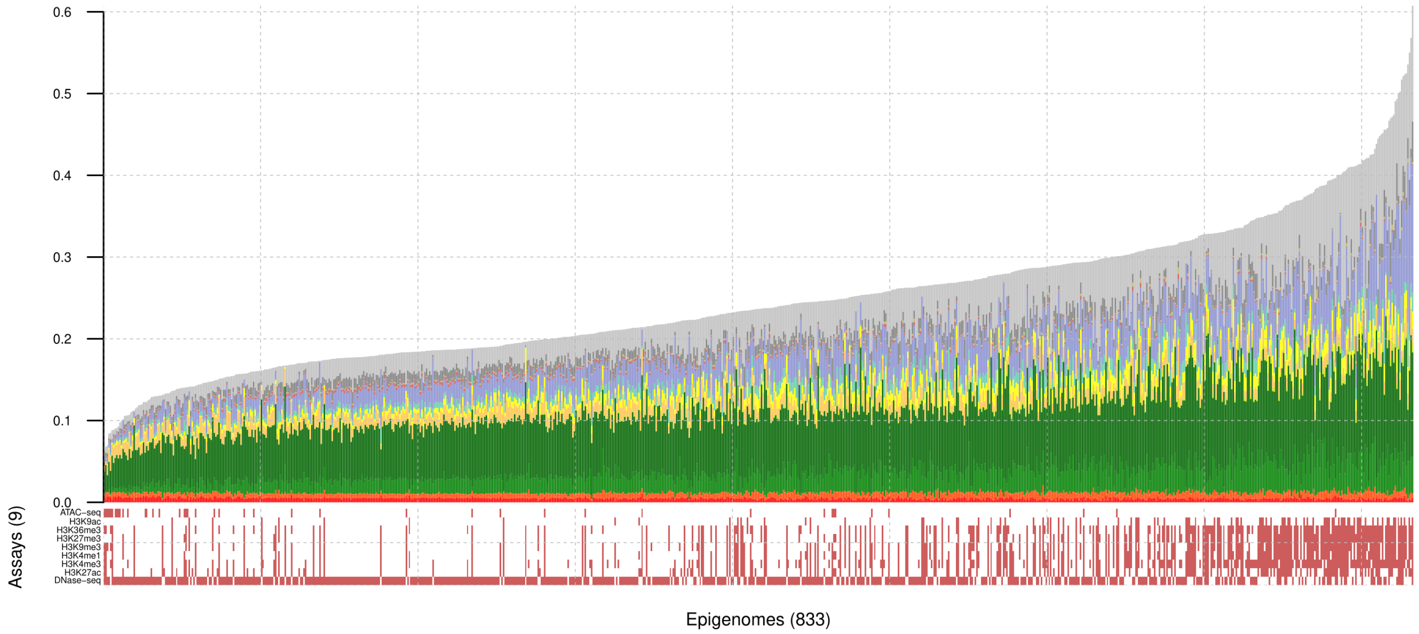


**Figure S14:** Epigenome state fractions for the ChromHMM 18-state model across 833 epigenomes after QC. Lower panel shows availability of 9 top marks, ordered by number of observed datasets. Epigenomes are ordered by percent of the genome not annotated as quiescent.

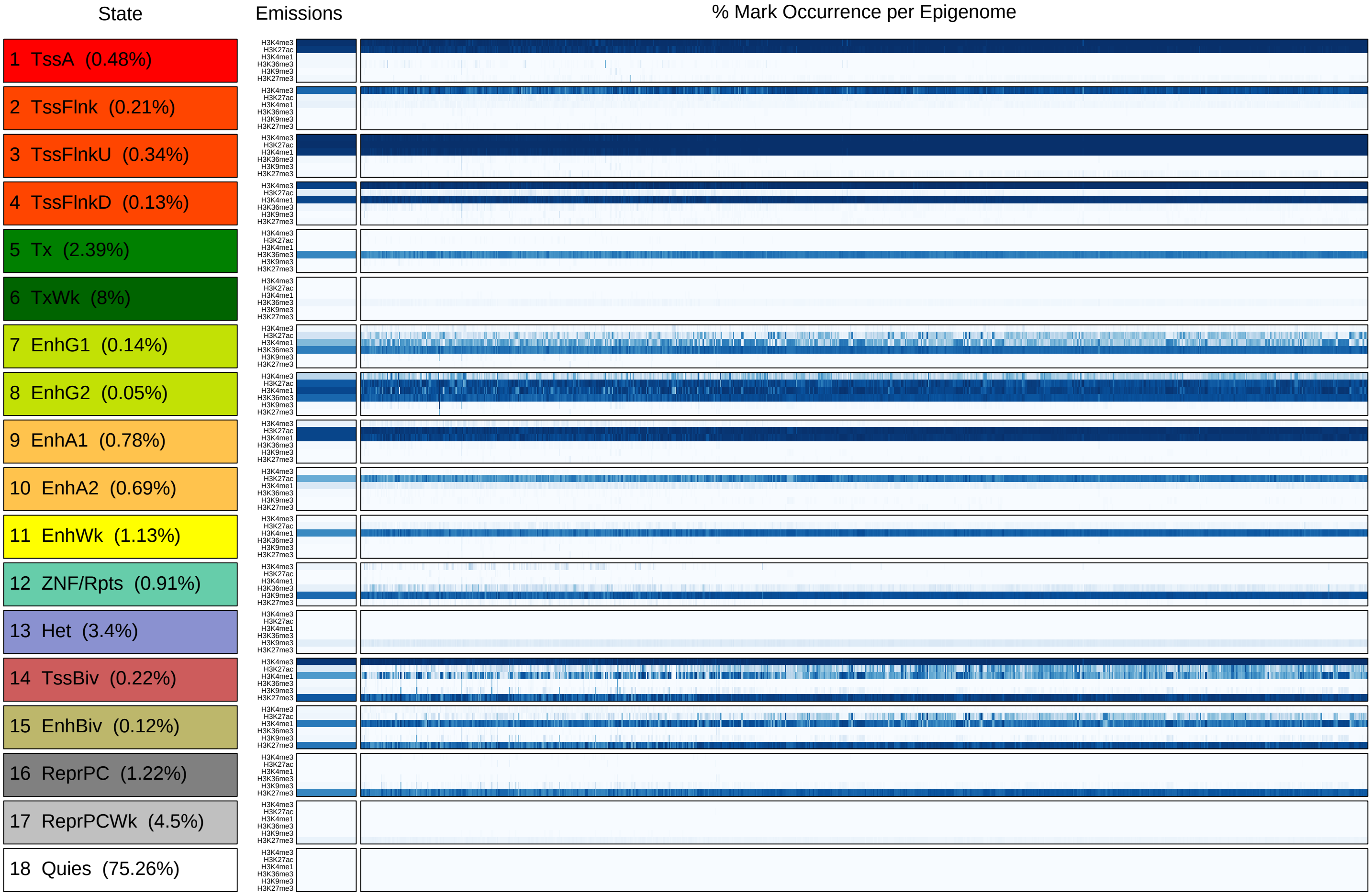


**Figure S15:** Comparison of per state (left panel) model emissions (middle panel) against mark occurrence in state calls (right panel) across 833 epigenomes (columns in right panel). Observed occurrence matched the emissions closely, with three exceptions. These exceptions corresponded to bivalent chromatin states and transcribed enhancers (cumulatively covering 0.48% of the genome on average), which showed discrepancies for 12.1% of the imputed epigenomes on average, likely stemming from their low frequency in the genome, and the frequent co-occurrence of H3K27ac and H3K4me1

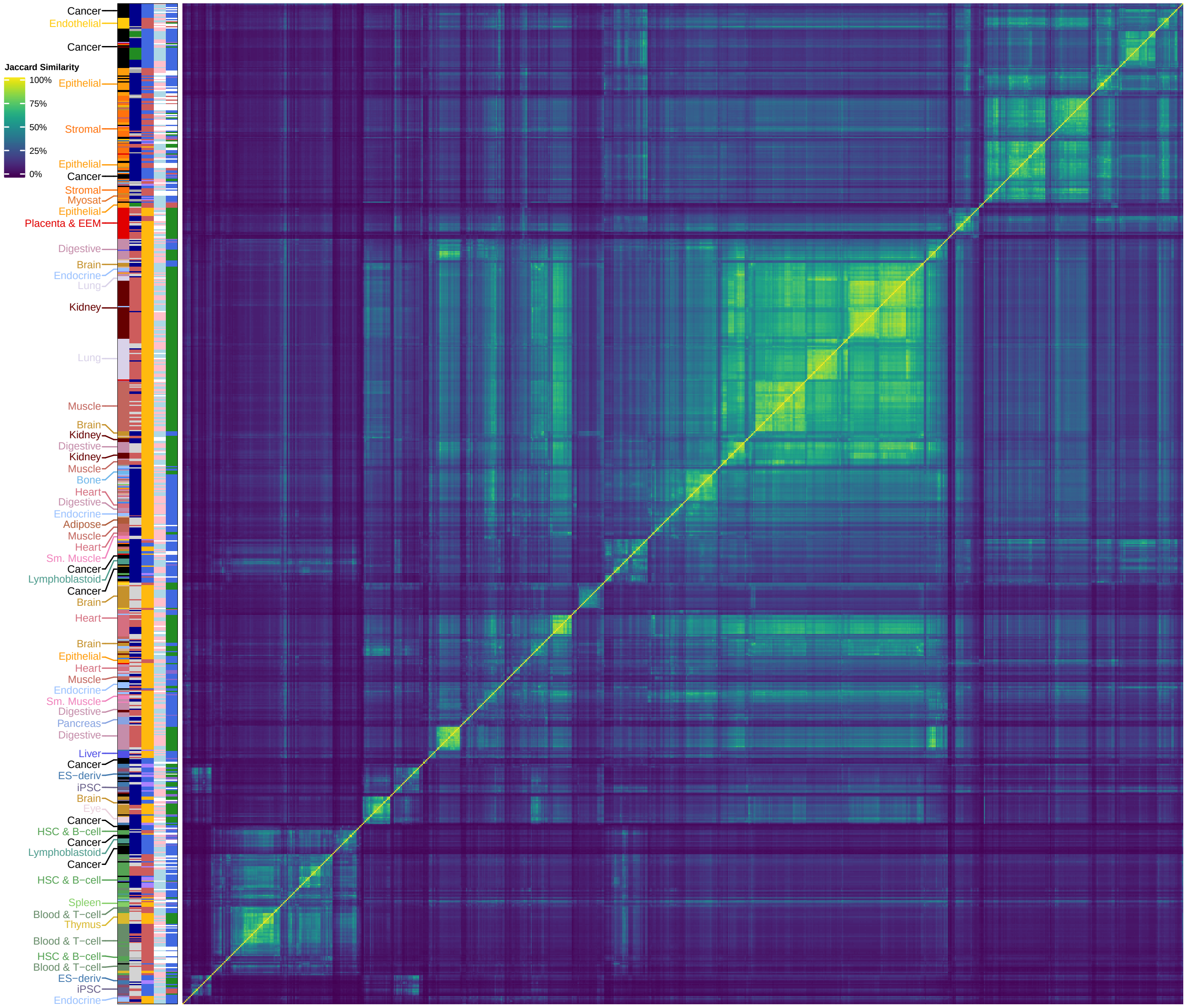


**Figure S16:** Jaccard similarity matrix (heatmap) across 833 epigenomes (metadata on left) from binarized enhancer activity matrix (2.1M enhancers by 833 epigenomes). Similarity matrix clustered by complete-linkage clustering. Consecutive blocks of at least six samples from the same group are labeled on the left.

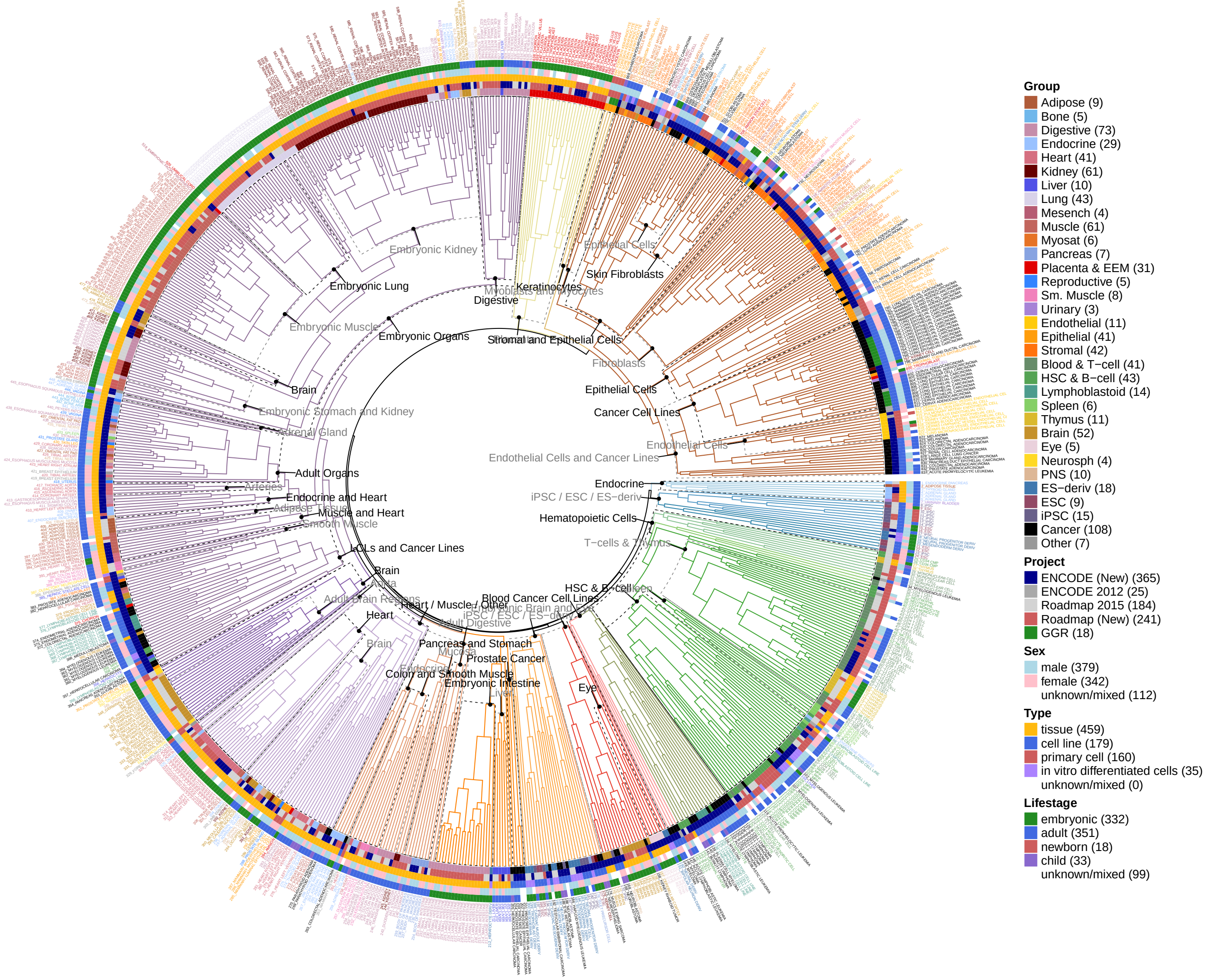


**Figure S17:** Tree constructed from complete-linkage hierarchical clustering of jaccard similarity matrix of enhancer activity across epigenomes (Supplementary Figure S5A). 50 subtrees covering the full tree are annotated with their major subgroups. Tree is cut and colored to create 20 clusters for purposes of visualization. Leaves are labeled with metadata and reduced sample names and colored according to their tissue group. Metadata (heatmap track) from inside-out: tissue group, project, sample type, and donor sex and lifestage.

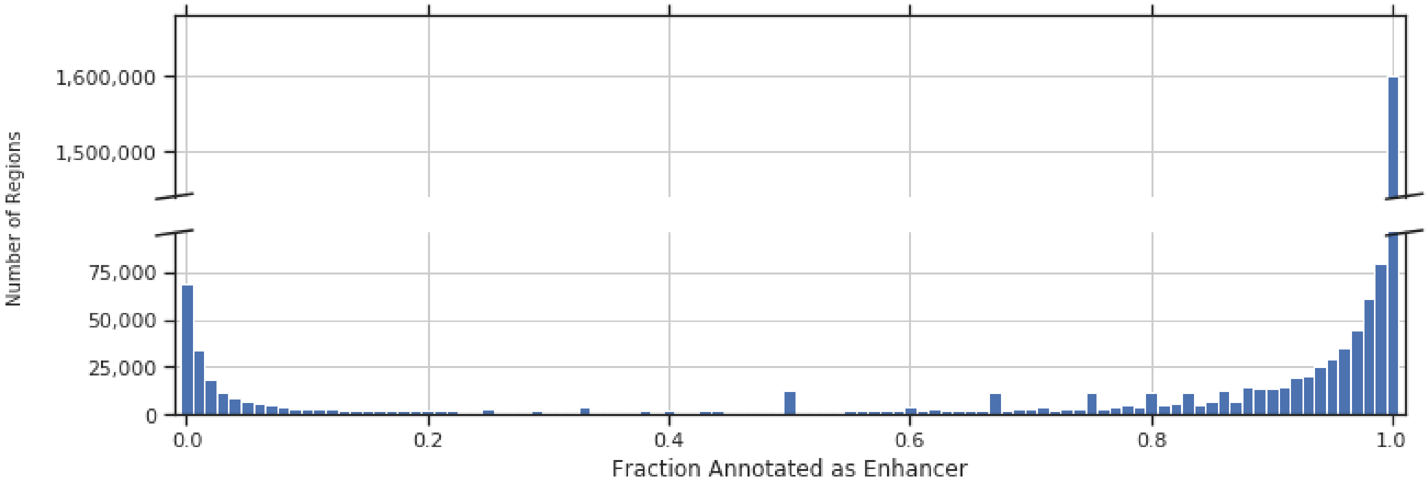


**Figure S18: (A)** Number of DHS sites annotated as enhancers instead of promoters across 833 epigenomes. Most regions are either labeled enhancer (at least 75% of occurrences are enhancers) or promoter (at least 75% of occurrences are promoters) across all of their active occurrences. Using these cutoffs, we labeled 2,069,090 enhancers, 204,104 promoters, and 122,358 dyadic elements (neither specifically promoter or enhancer).

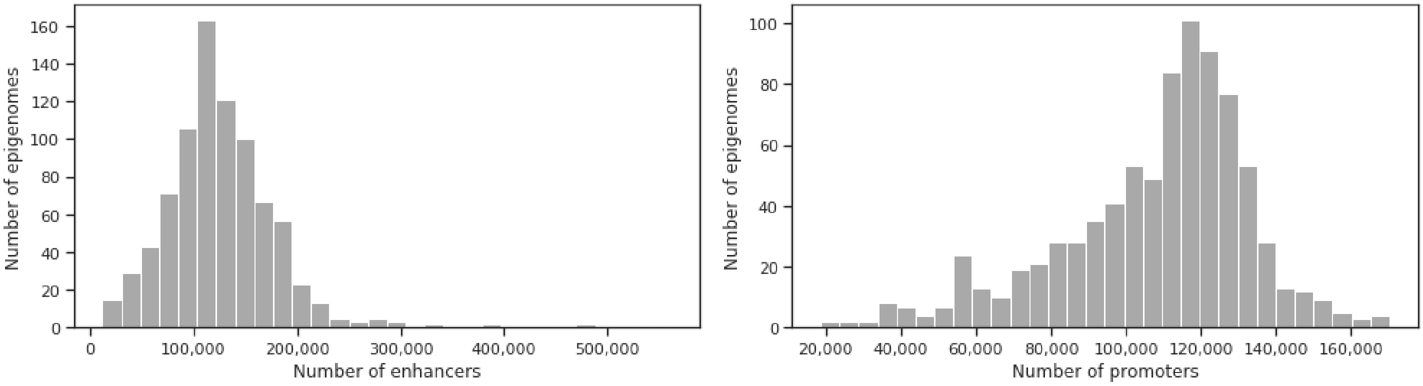


**Figure S18: (B)** Histogram of number of active enhancers (left) and promoters (right) per sample (row-margins of binary activity matrix).

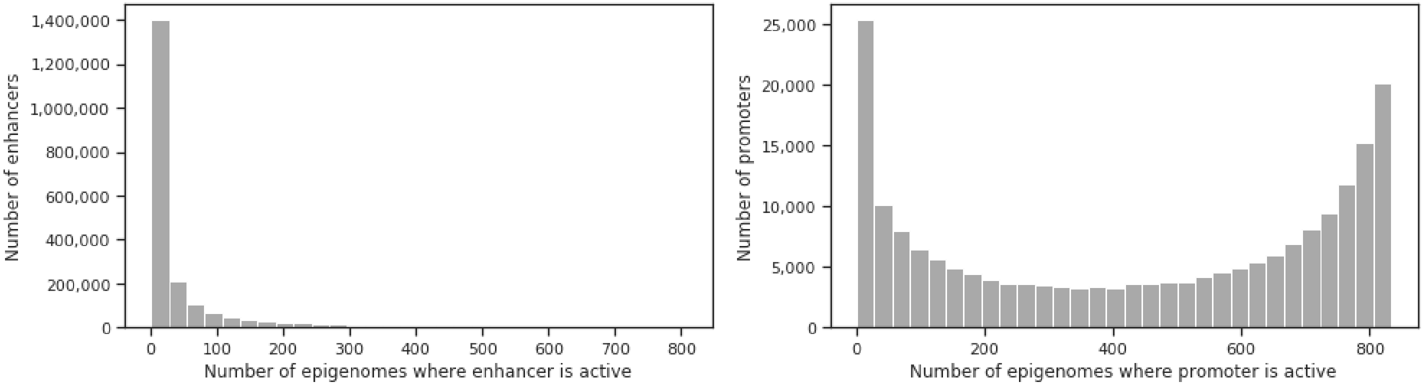


**Figure S18: (C)** Histogram of the number of samples for which each enhancer (left) or promoter (right) is an active enhancer (column-margins of binary activity matrix)

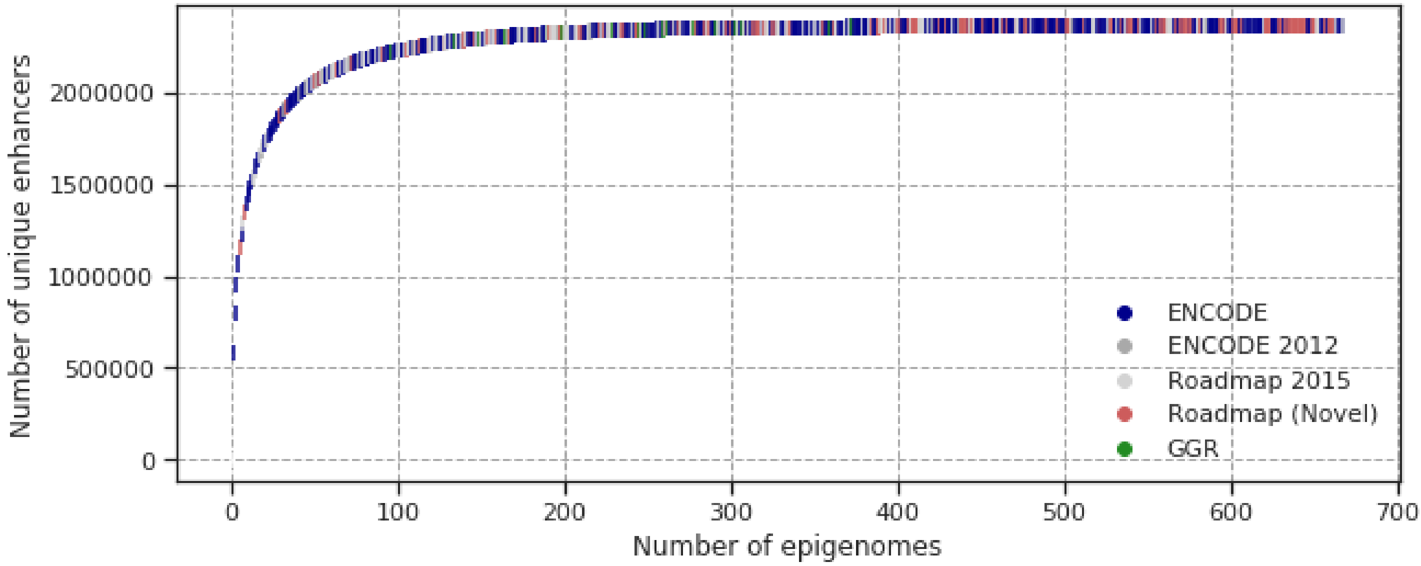


**Figure S19: (A)** Rarefaction curve for enhancer recovery for all enhancer and dyadic elements (2.3M total) across 833 samples (points, colored by project of origin). Curve was created by iteratively adding the sample contributing the most novel active elements until all 2.3M elements were accounted for.

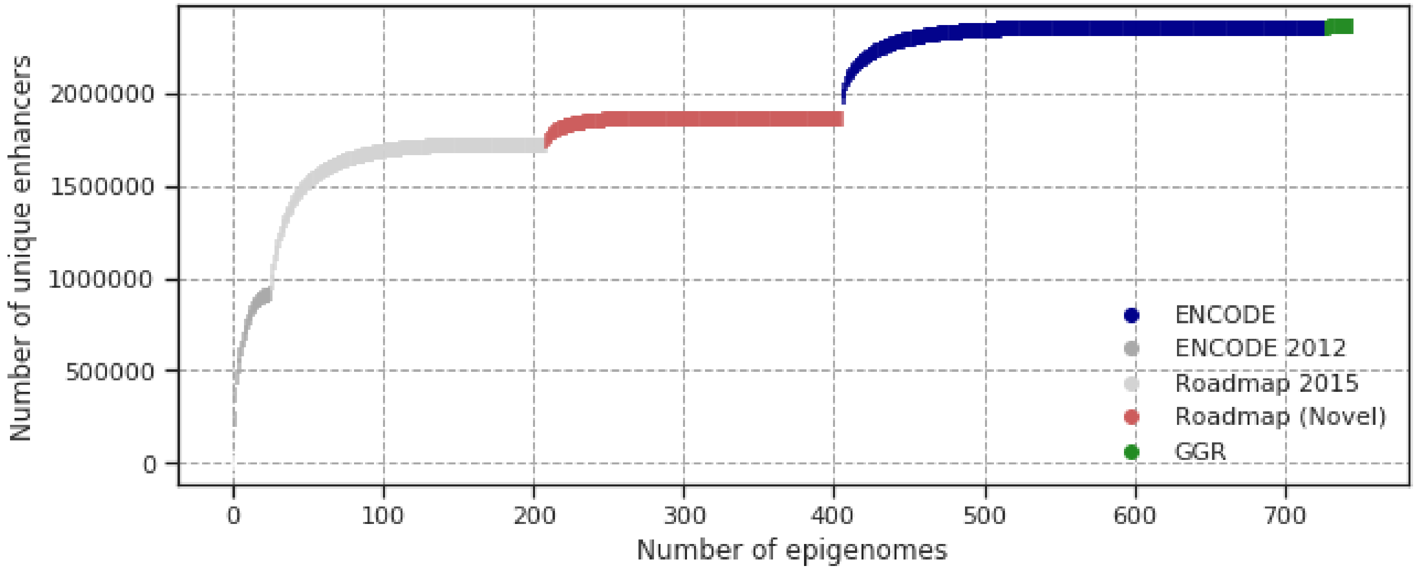


**Figure S19: (B)** Rarefaction curve for enhancer recovery, shown in order of project completion, for all enhancer and dyadic elements (2.3M total) across 833 samples (points, colored by project of origin). Curve was created by iteratively adding the sample contributing the most novel active elements until all 2.3M elements were accounted for. Samples were considered in order of project publication/completion, only taking samples from the next project when the current project did not contribute any more enhancers.

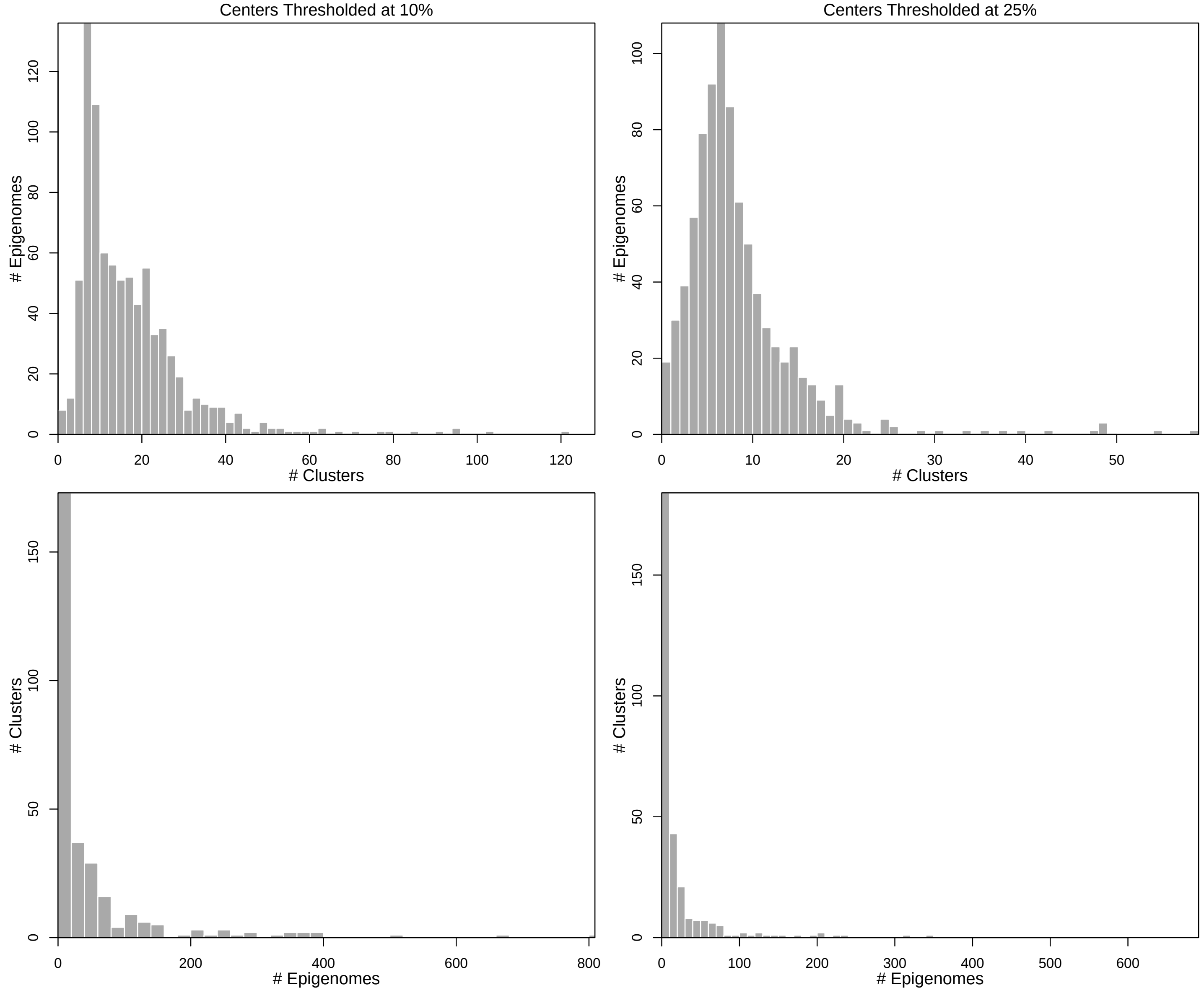


**Figure S20:** Module inclusion distributions according to two different module inclusion cutoffs of 10% and 25% (columns). Number of modules (N=300) for each of 833 epigenomes with module inclusion of at least 10% or 25% (top). Number of epigenomes (N=833) for each of 300 modules at the same inclusion cutoffs.

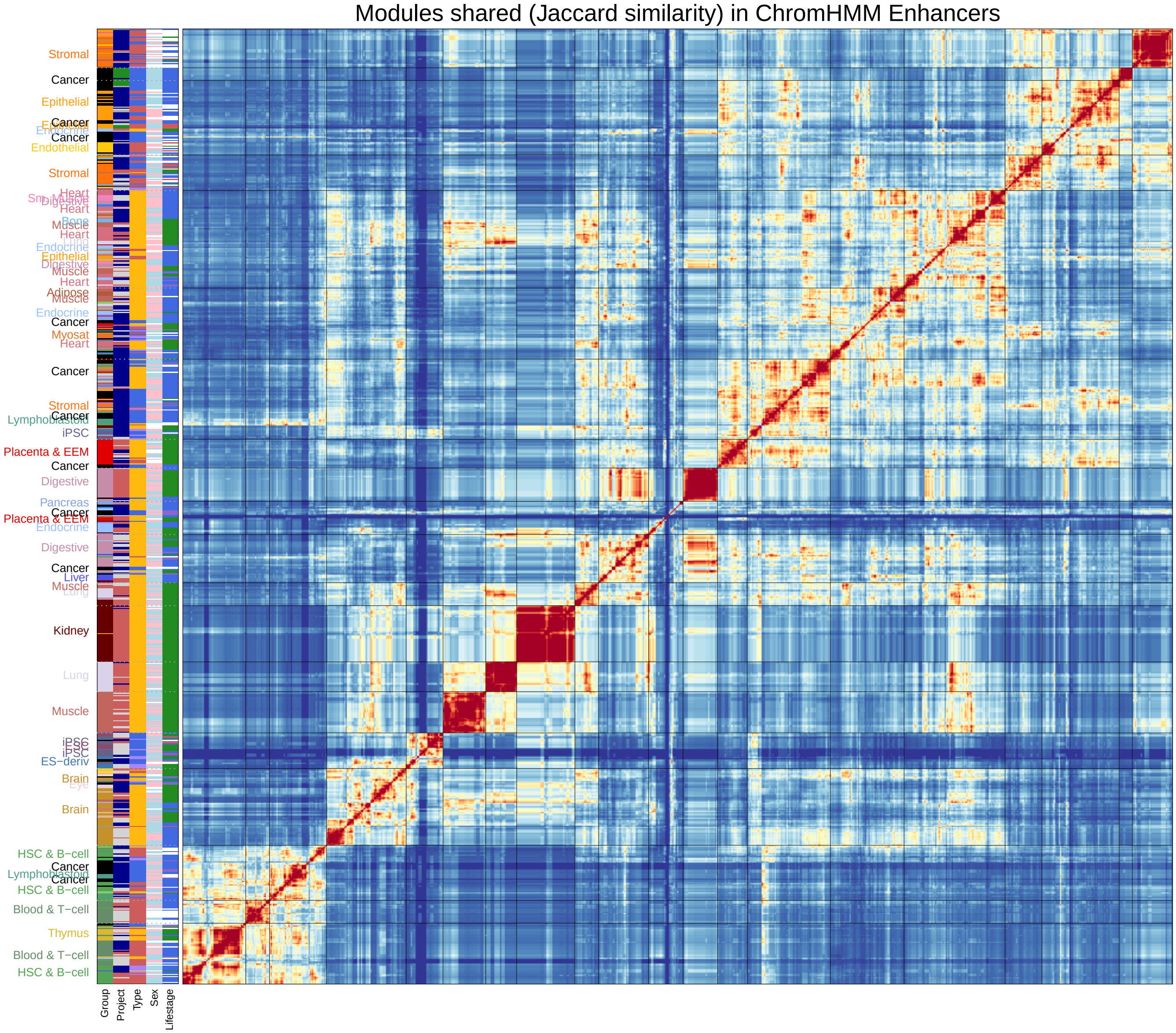


**Figure S21:** Sample similarity (heatmap) across 833 epigenomes by their number of shared modules.(jaccard similarity, intersection over union of modules of each pair of epigenomes). Similarity matrix clustered by Ward's method. Consecutive blocks of at least six samples from the same group are labeled on the left.

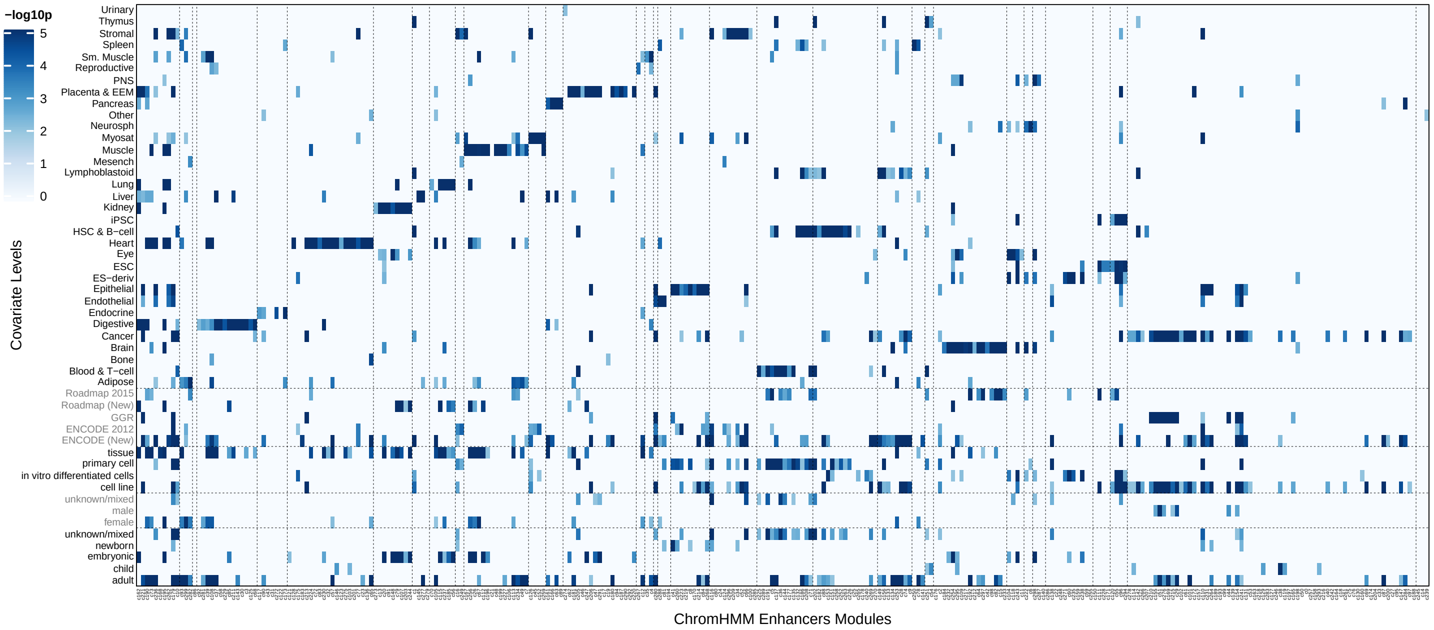


**Figure S22:** Full module enrichments across main metadata facets of group, project, type, sex, and lifestage, for which the reduced version is shown as a panel in Figure 3A. Significance assessed according to hypergeometric test on the module centers matrix with module inclusion of at least 25%.

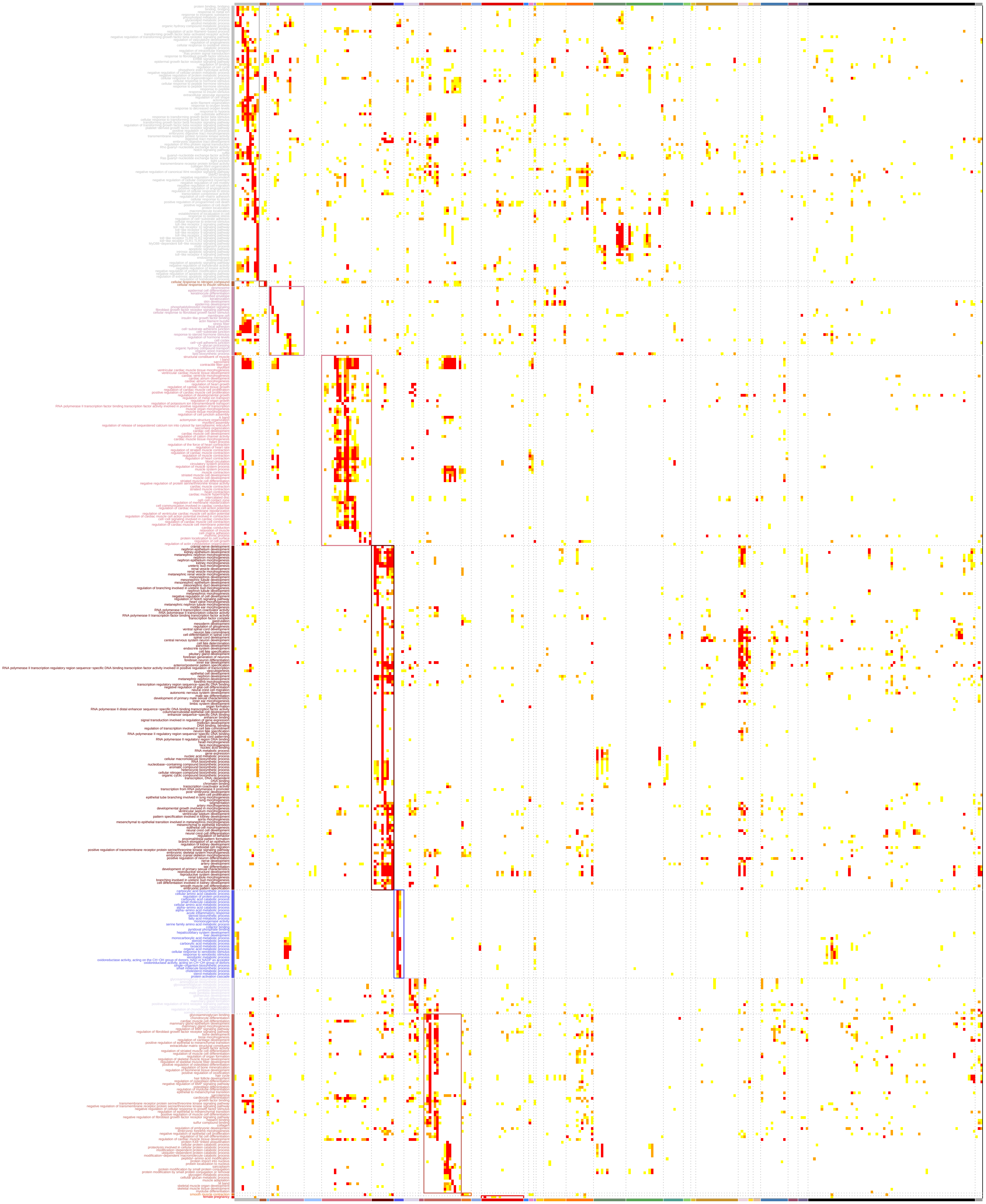

**Figure S23:** All 865 module-specific GO terms (BP, MF, CC), including all terms enriched in less than 10% of modules and with a maximum enrichment of at least $-\log_{10} p > 4$. This is the full version of figure 3B).

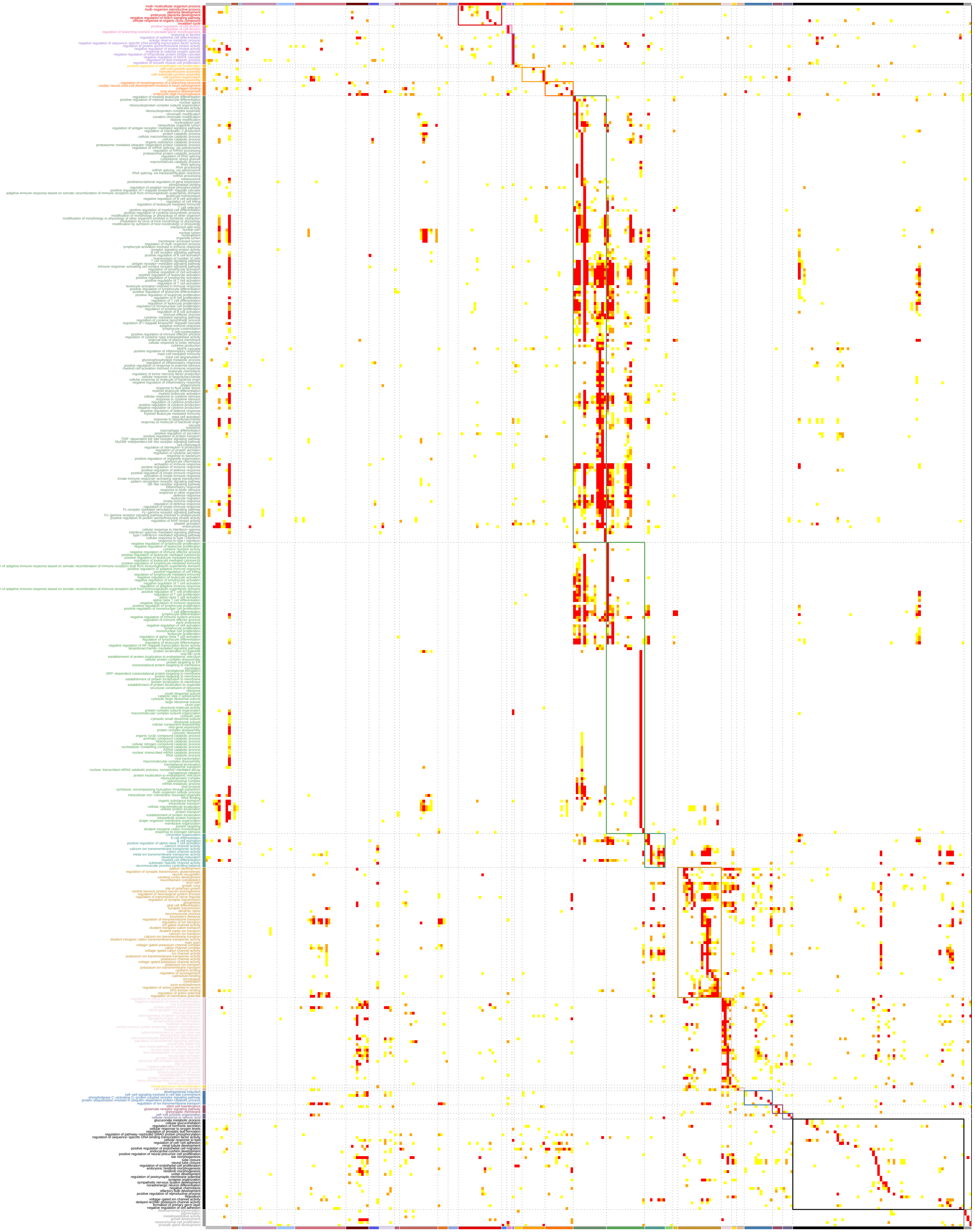

**Figure S23:** (continued)

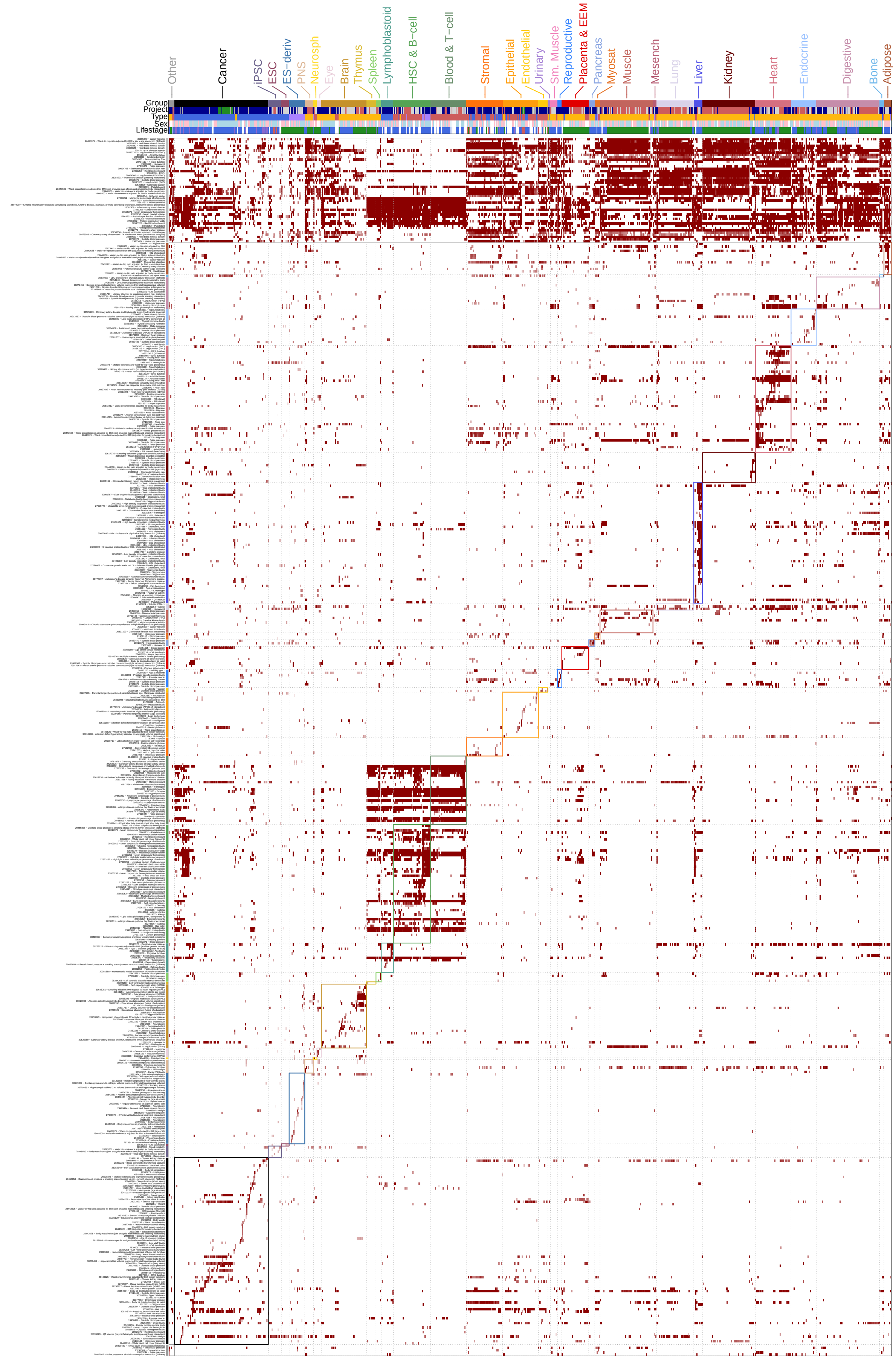


**Figure S24:** All epigenome trait enrichments at FDR < 1%, for 534 traits (rows) and 833 epigenomes (columns)

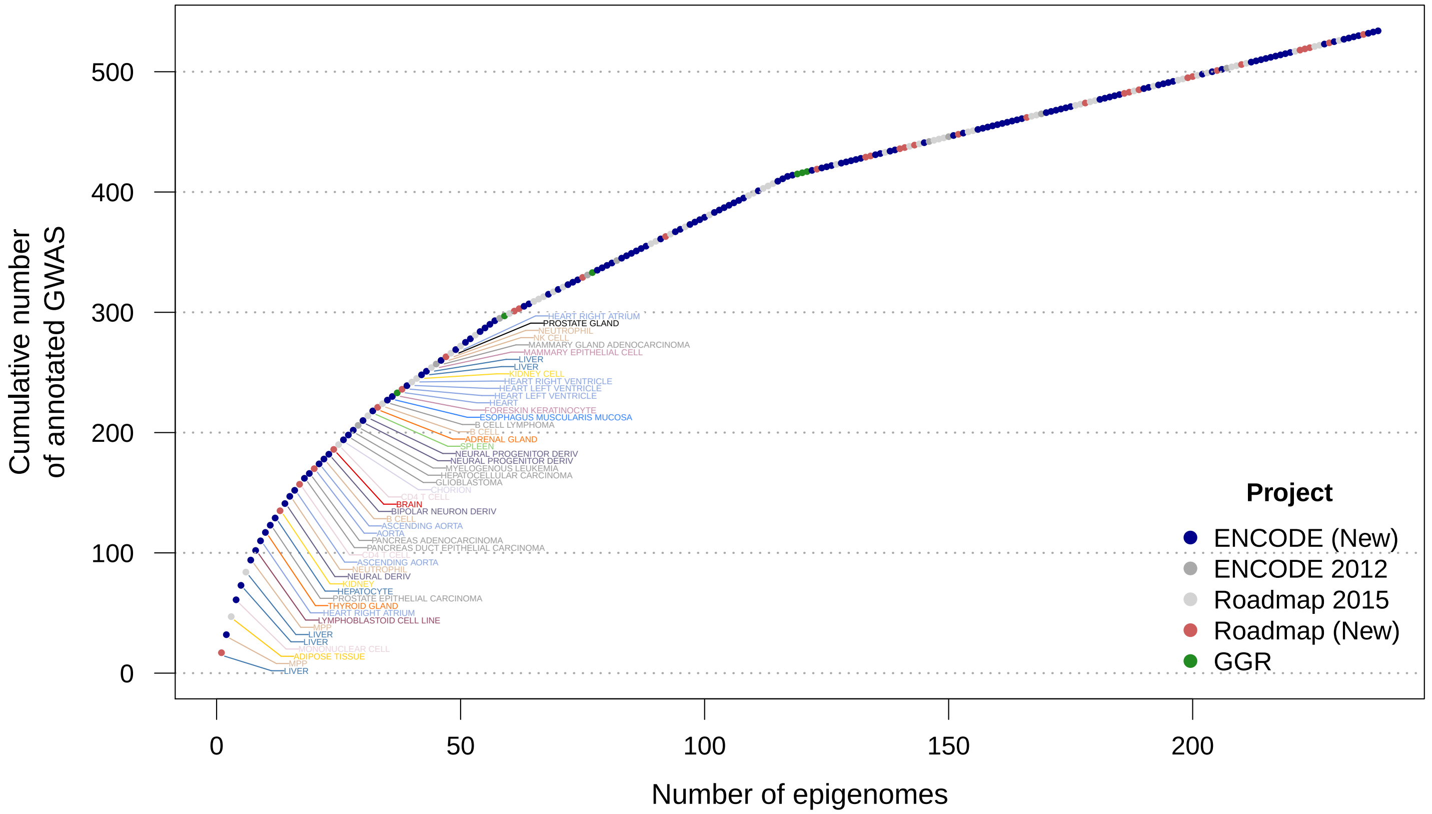


**Figure S25:** Rarefaction curve for all maximal enrichments. The rarefaction curve was calculated by iteratively adding the sample that contained the maximal enrichment for most remaining GWAS until all GWAS were accounted for. Points (238 samples) are colored according to the sample's project and the top 50 contributing samples are labeled and colored according to their tissue group.

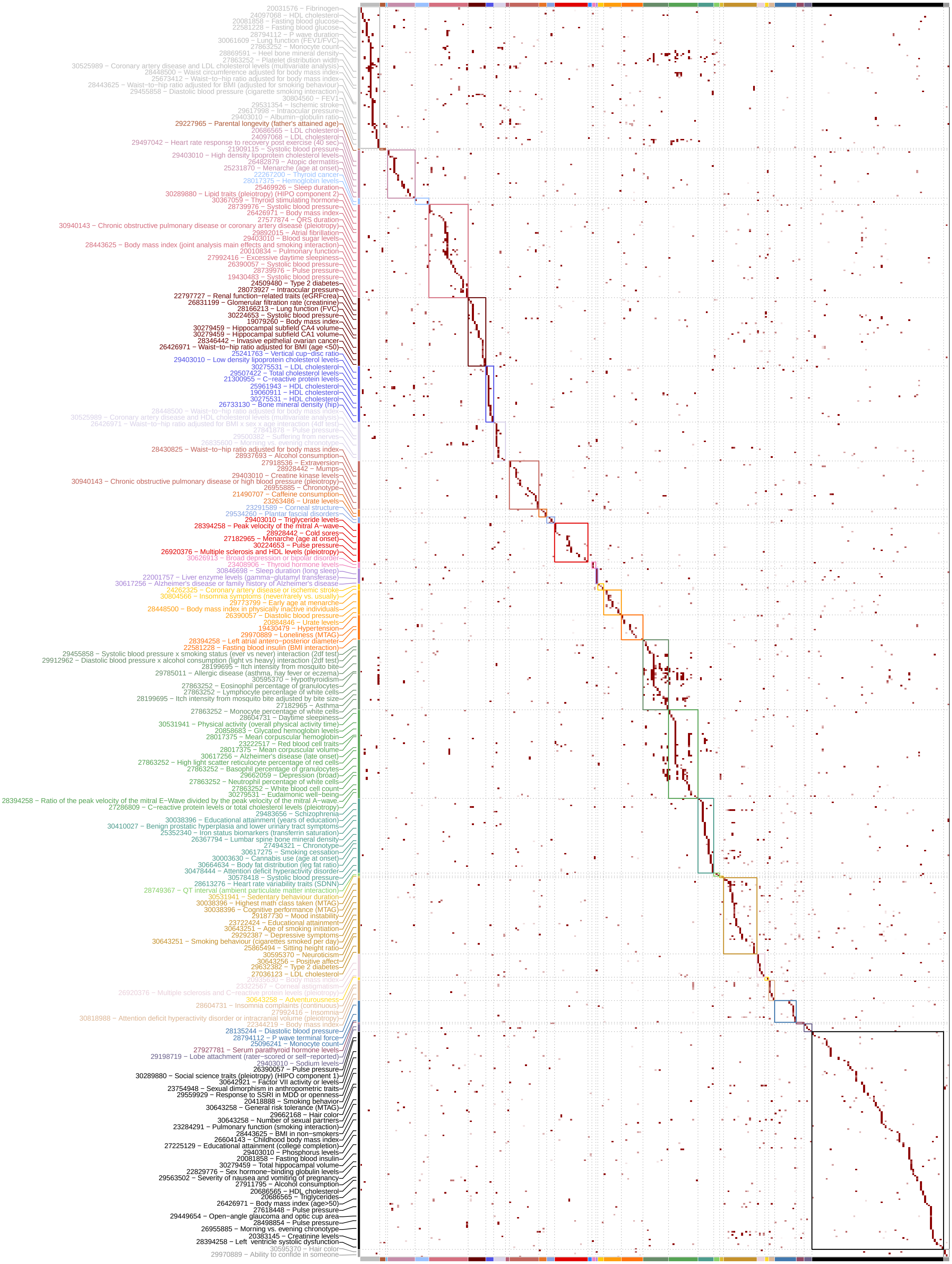


**Figure S26:** Modules trait enrichments at FDR $< 1\%$, for 804 traits (rows) and 300 modules (columns). Modules are ordered as in Figure 3A. Representative traits are chosen by a bag-of-words approach as in Figures 3B and 4A.

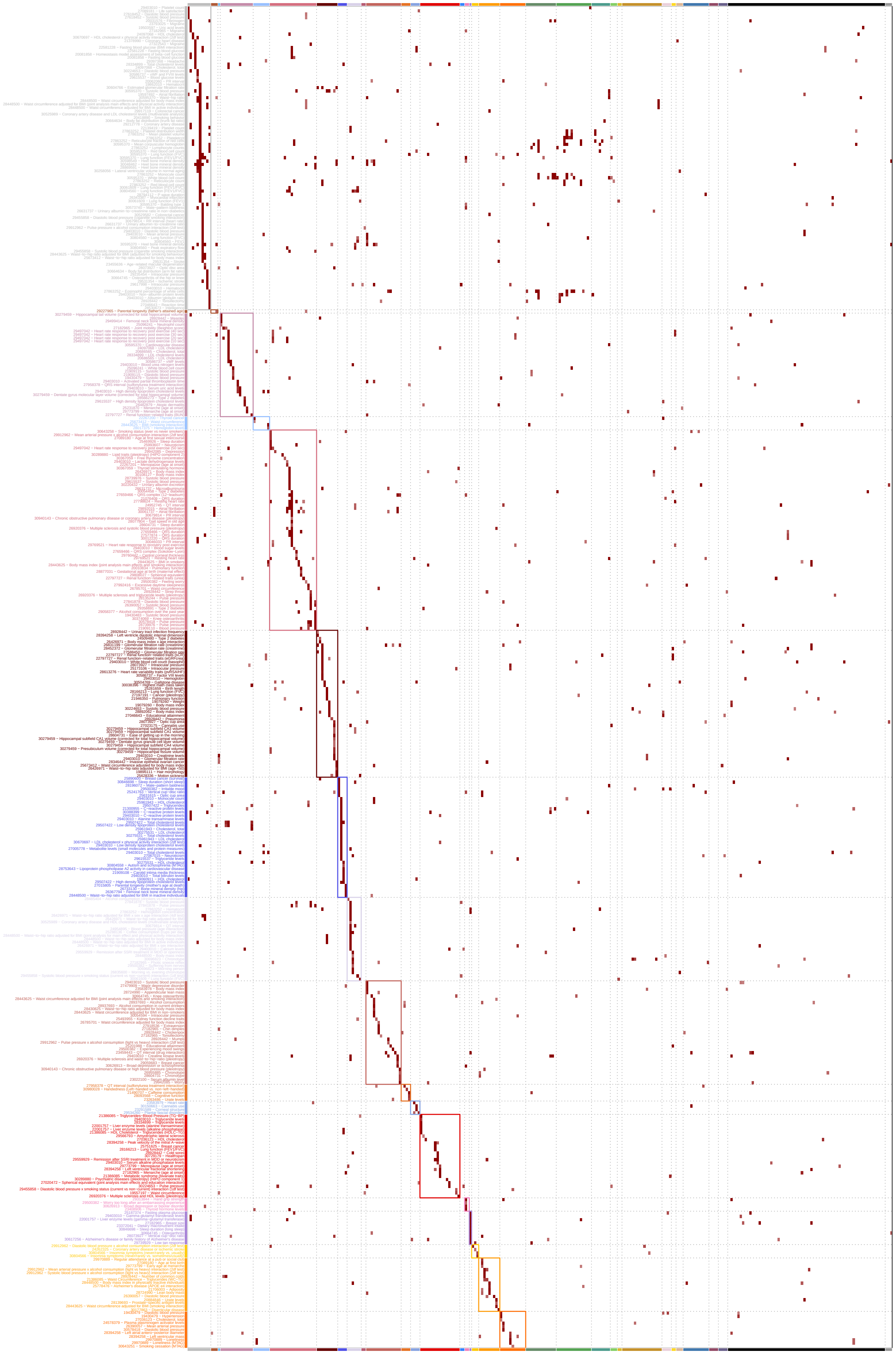

**Figure S27:** All module trait enrichments at FDR < 1%, for 804 traits (rows) and 300 modules (columns). Modules are ordered as in Figure 3A.

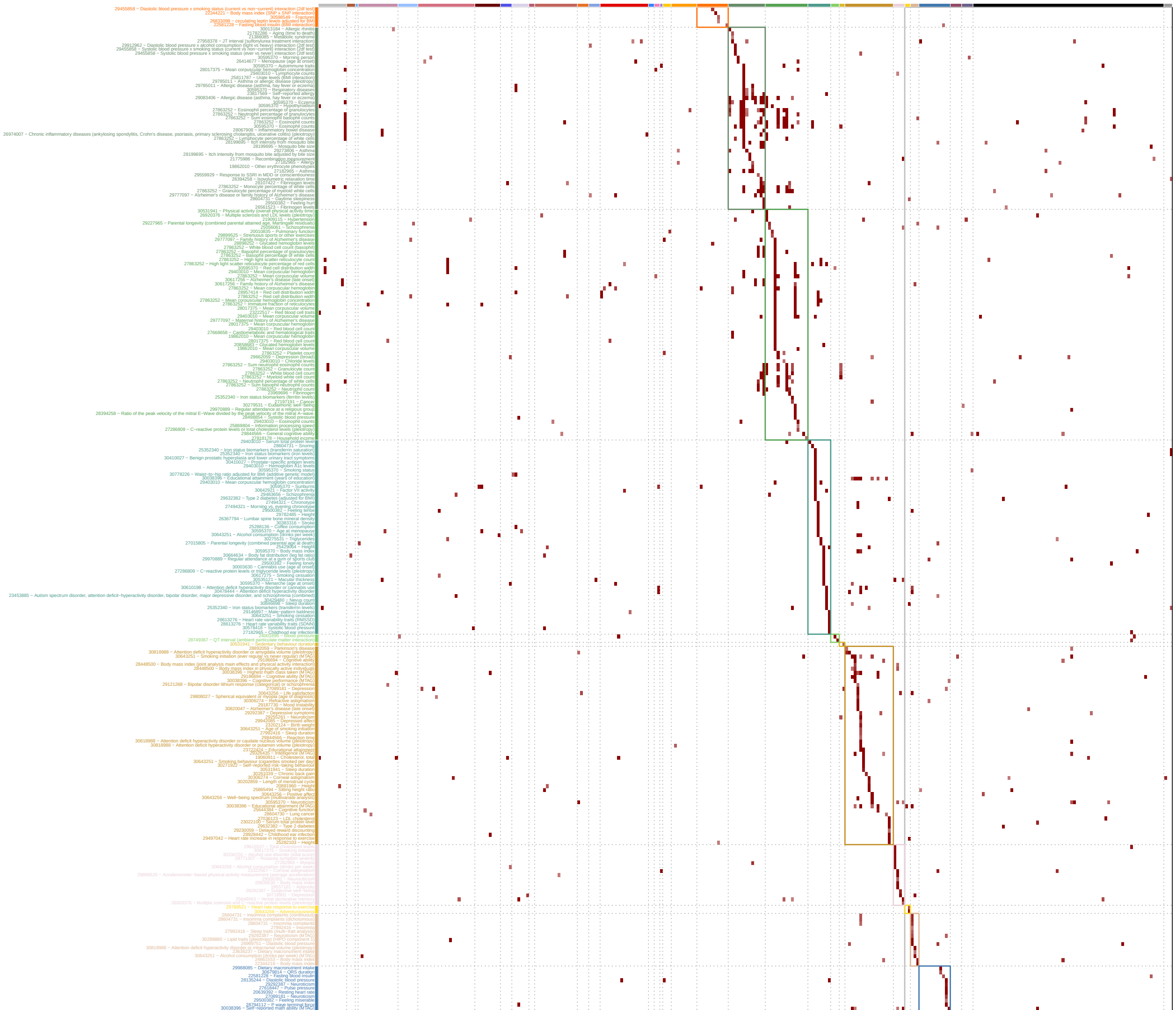

**Figure S27:** (continued)

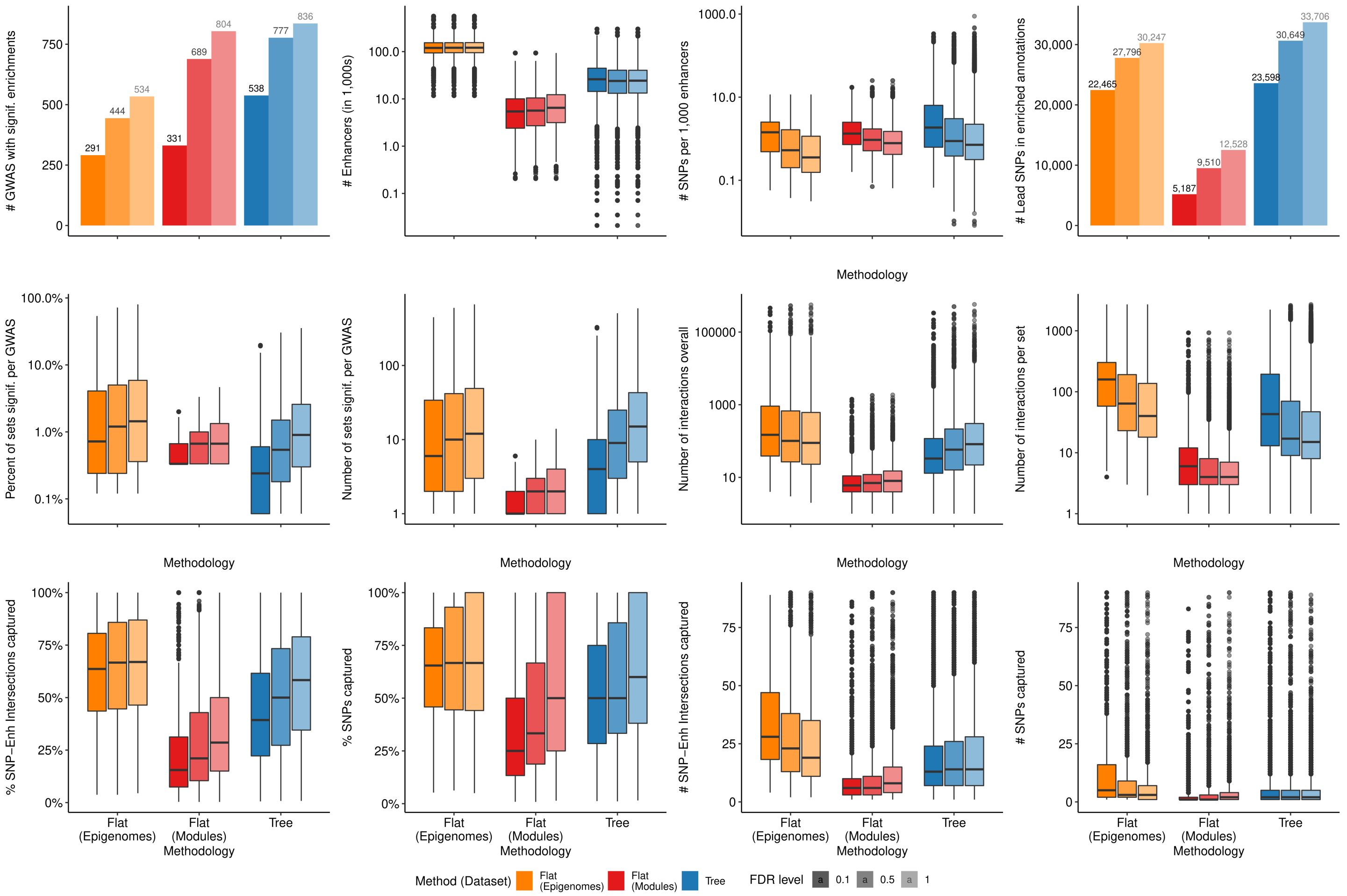


**Figure S28:** Extended method validation figures comparing epigenomic GWAS enrichments using different methodologies (x-axis) for three FDR cutoffs (shades). First and last figures from top row are shown as Figure 4E.

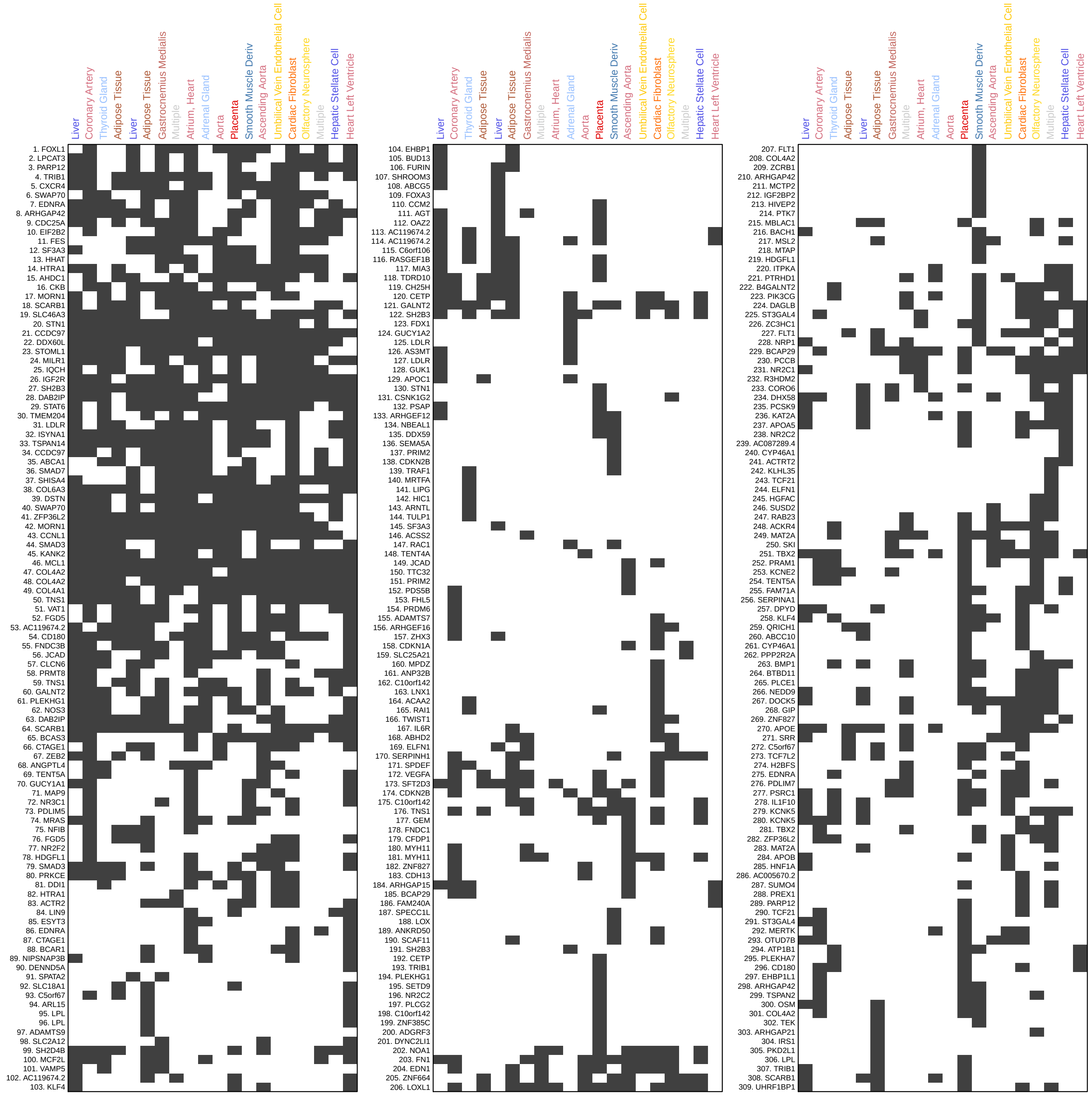


**Figure S29: (A)** Loci of lead SNPs for CAD in top 20 significantly enriched nodes. Matrix is split in three and shows 309 SNPs against 20 nodes, by presence (black) or absence (white) in the node's enhancers. Loci are ordered by clustering their jaccard similarity across nodes using the Ward method. Each SNP is annotated with its nearest protein-coding gene.

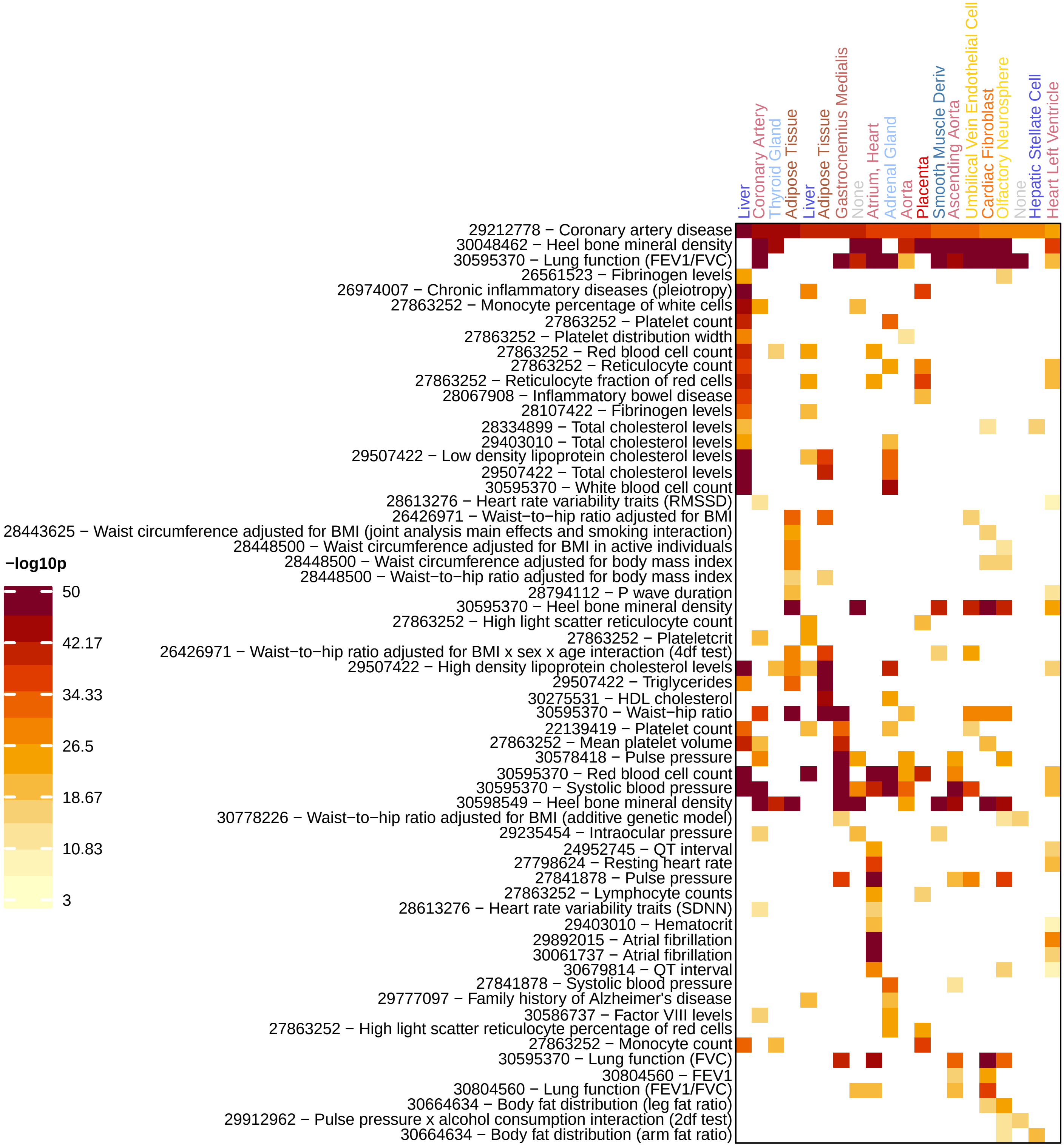


**Figure S29: (B)** Shared enrichments with CAD for 61 of 926 traits sharing at least two enrichments with the top 20 enriched nodes of CAD. Matrix is diagonalized according to maximal enrichment (nominal p-value, only enrichments passing FDR $< 0.1\%$ are shown).

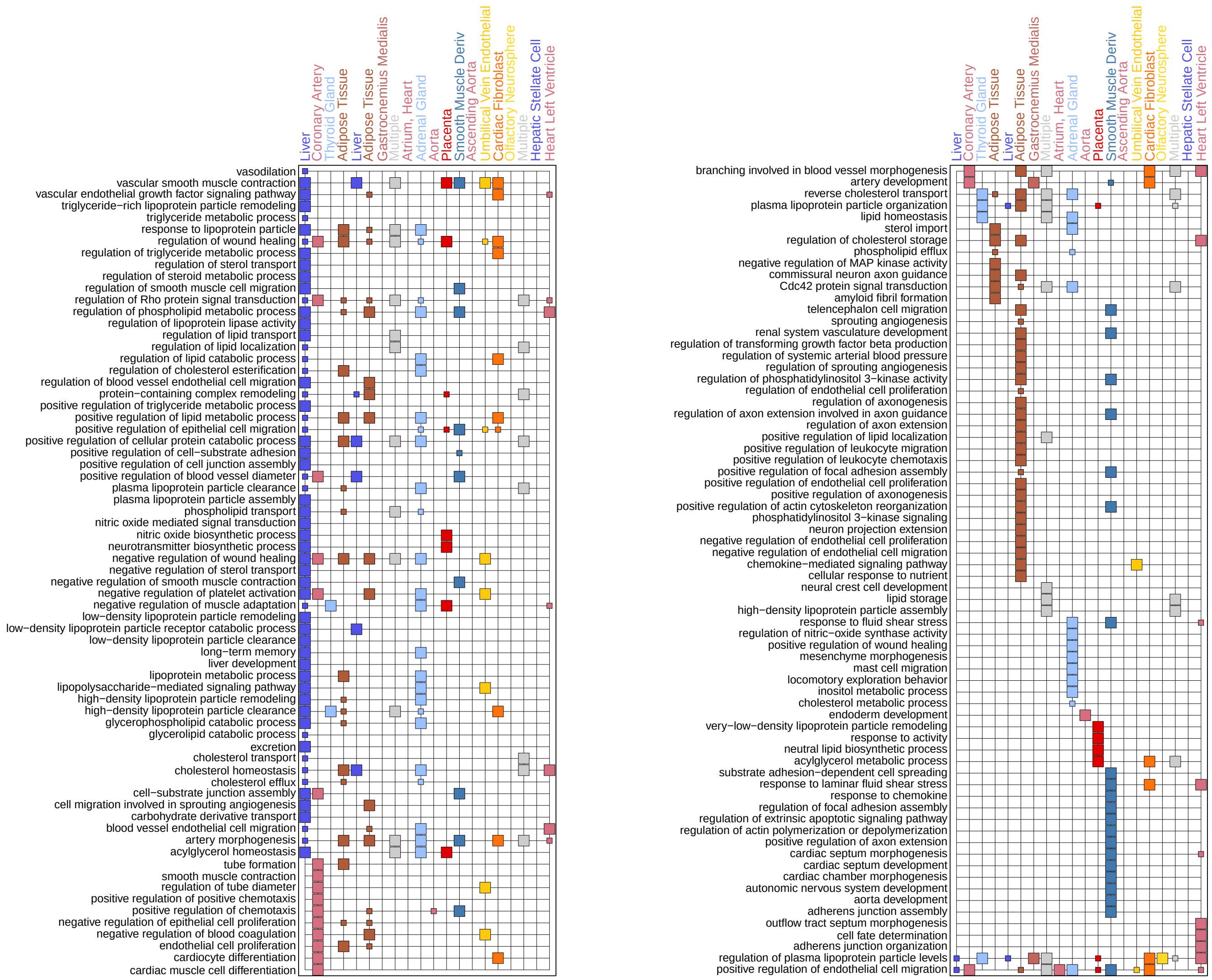

**Figure S29:** **(C)** Extended GO terms for CAD lead SNPs in enriched nodes. All go terms with at least $-\log_{10} q > 2$ enriched in less than 25% of nodes.

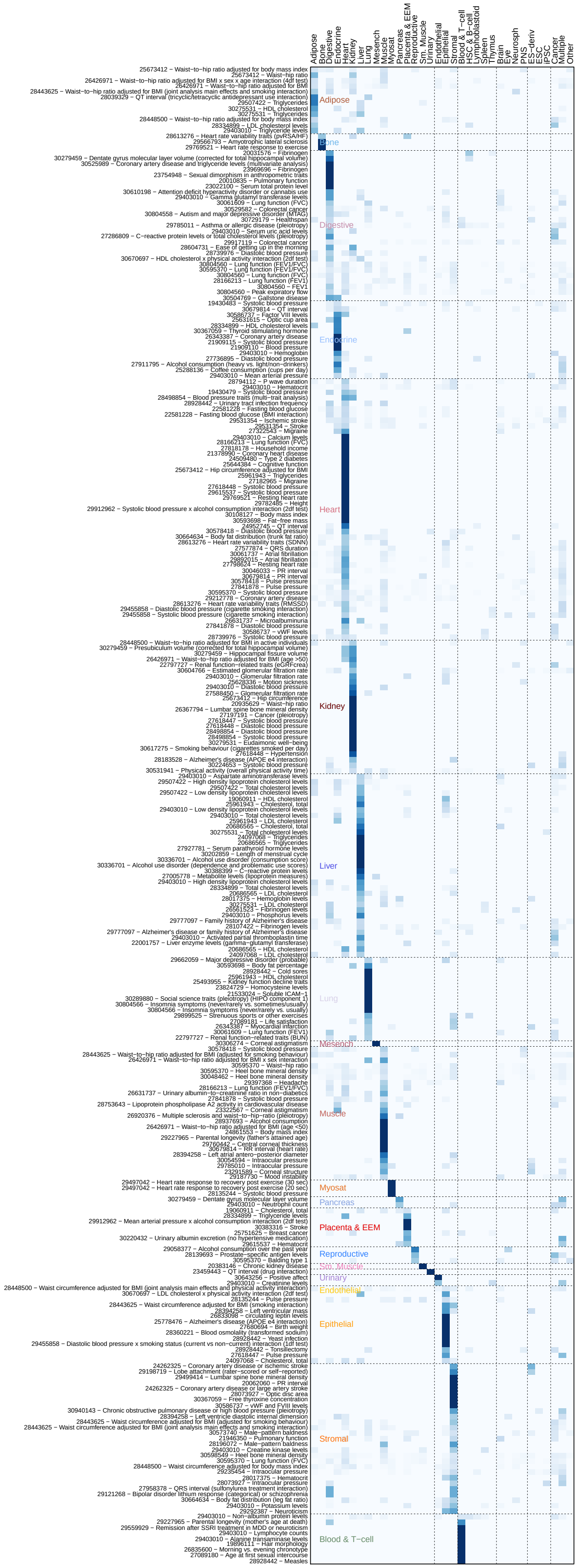

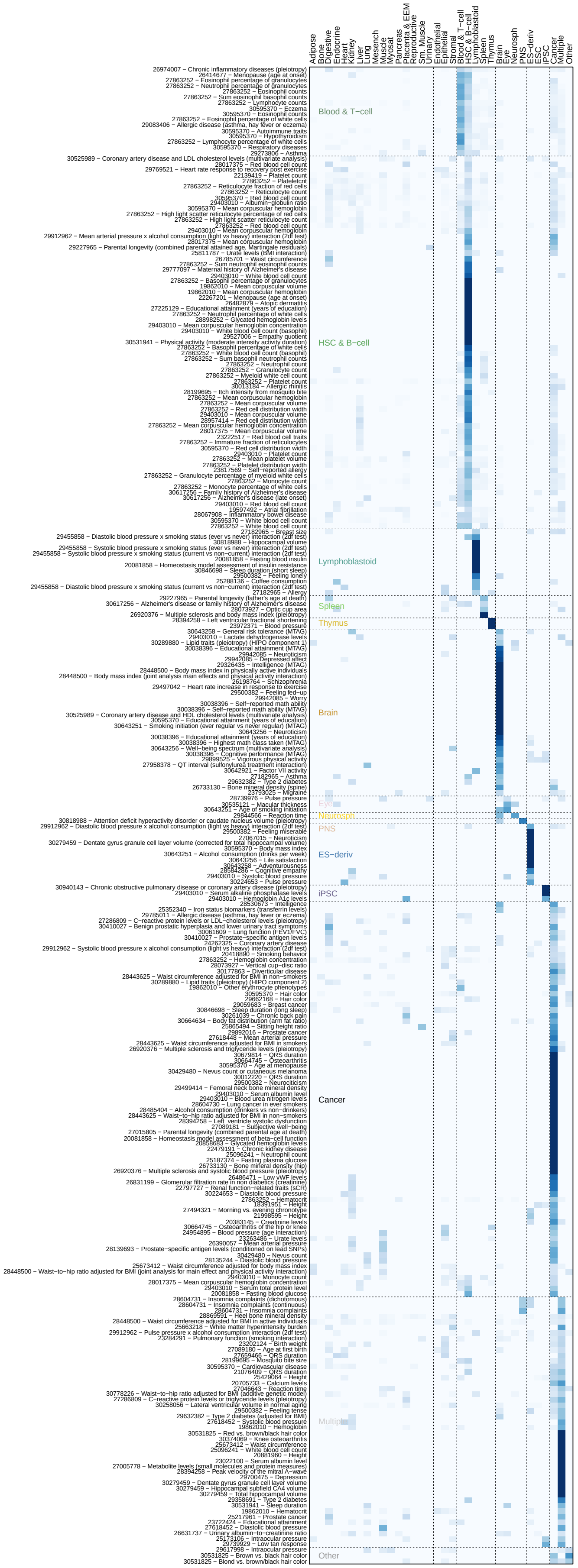

**Figure S30:** Overall tissue-level prioritization of 538 traits at FDR < 0.1%. Heatmap represents trait (rows) vs. tissue matrix and is split for visibility. Traits are diagonalized according to their top tissue enrichment, and values are the trait-normalized and tissue aggregated $-\log_{10}$p-values.

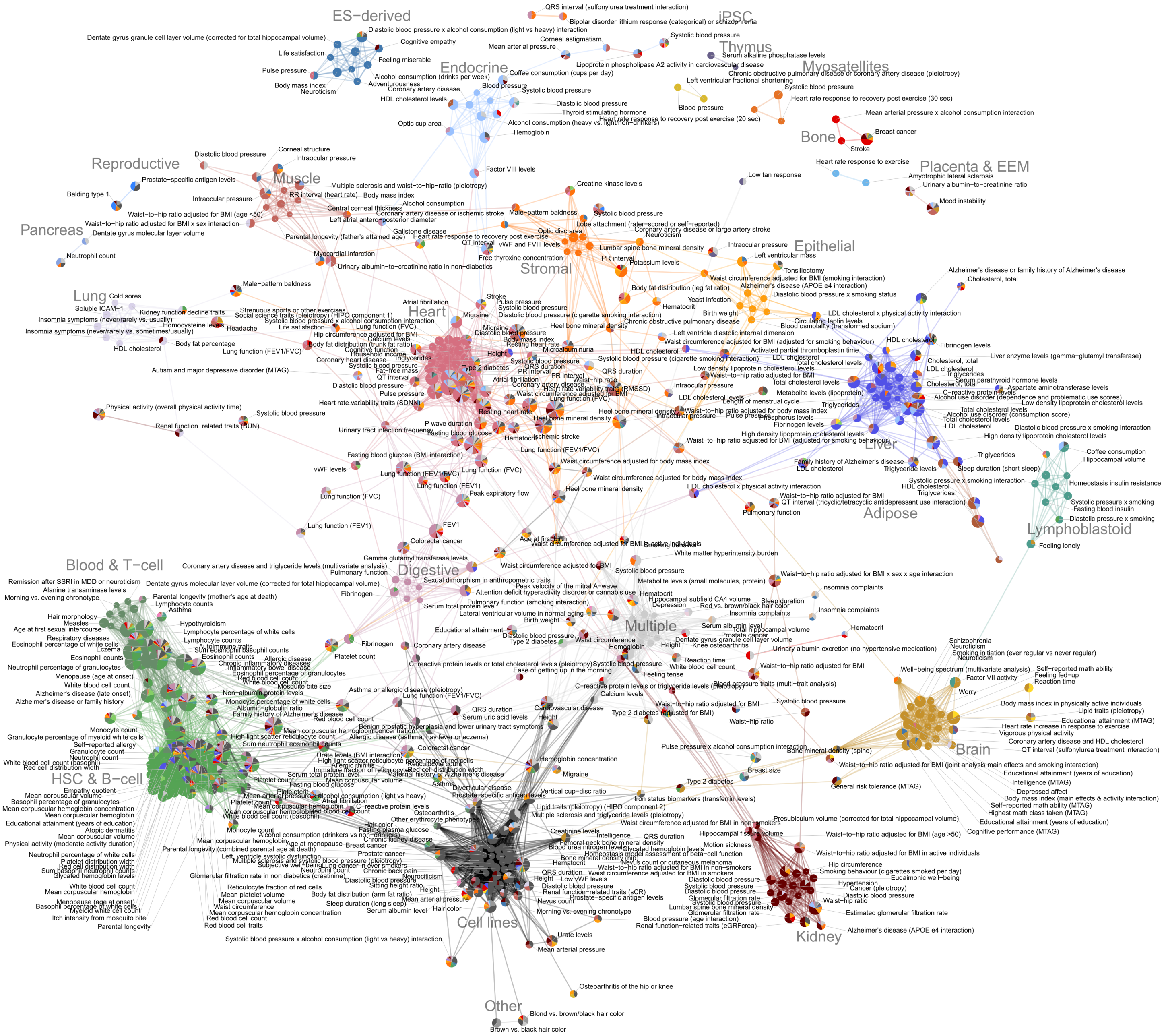


**Figure S31:** Trait-trait network (as Figure 7) across 511 traits (of 538 total, with edges) by similarity of epigenetic enrichments (cosine sim. ¿= 0.75), laid out using the Fruchterman-Reingold algorithm. Traits (nodes) are colored by contributing groups (pie chart by fraction of -log10p, size by maximal -log10p) and interactions (edges) by the group with maximal dot product of enrichments between two traits. All 511 traits labeled.

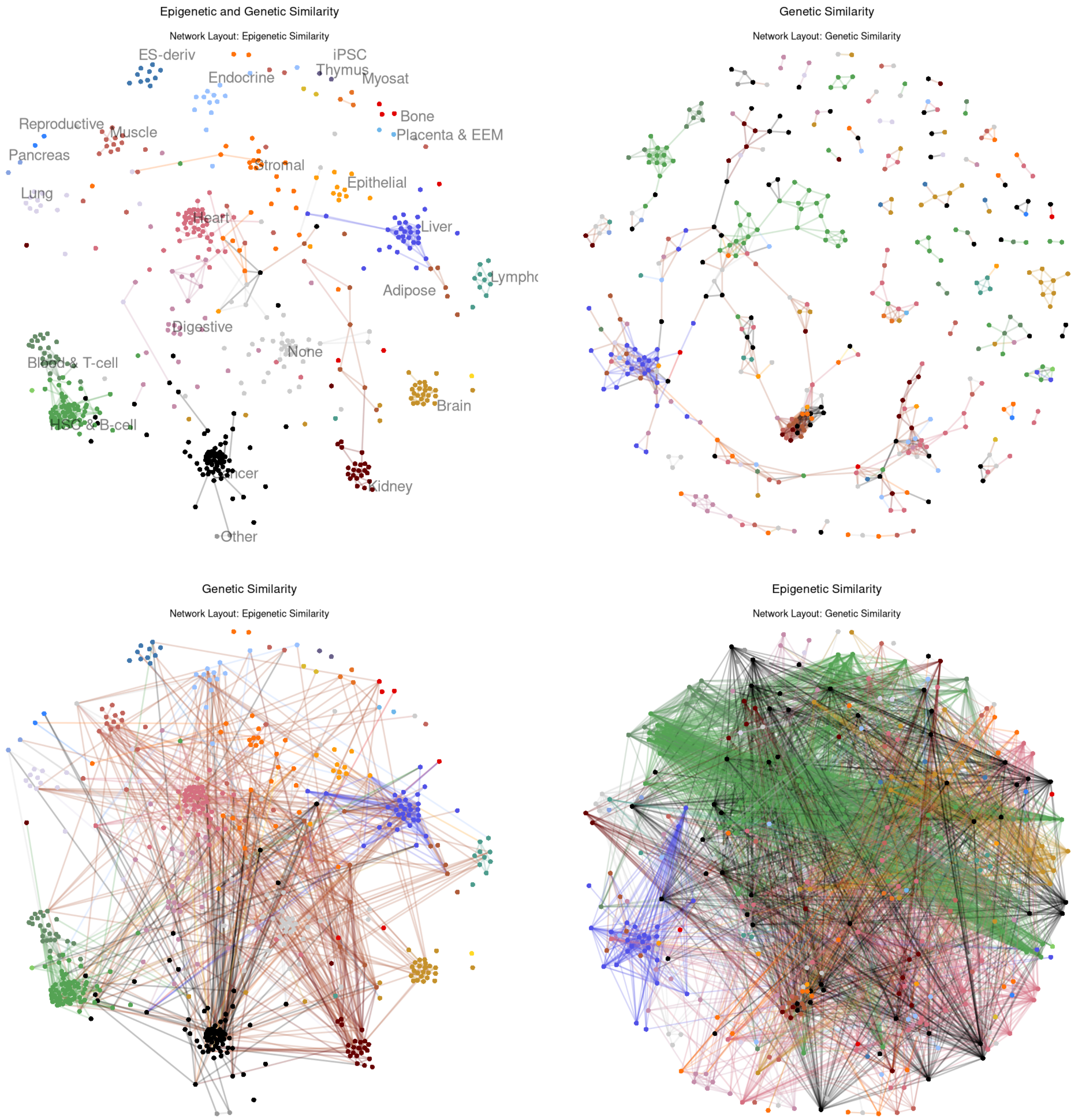


**Figure S32:** Comparison networks with genetics and epigenetics. (top left) Edges with both high epigenetic similarity and any genetic overlap on epigenetic similarity layout. (bottom left) All trait pairs with genetic overlap (> 5% jaccard similarity of lead SNPs overlapping when binned into 10k bp bins starting from the start of each chromosome, network using epigenetic layout. (top right) Network of trait pairs with any genetic overlap laid out by genetics. (bottom right) All trait pairs with high epigenetic similarity on the genetic layout network

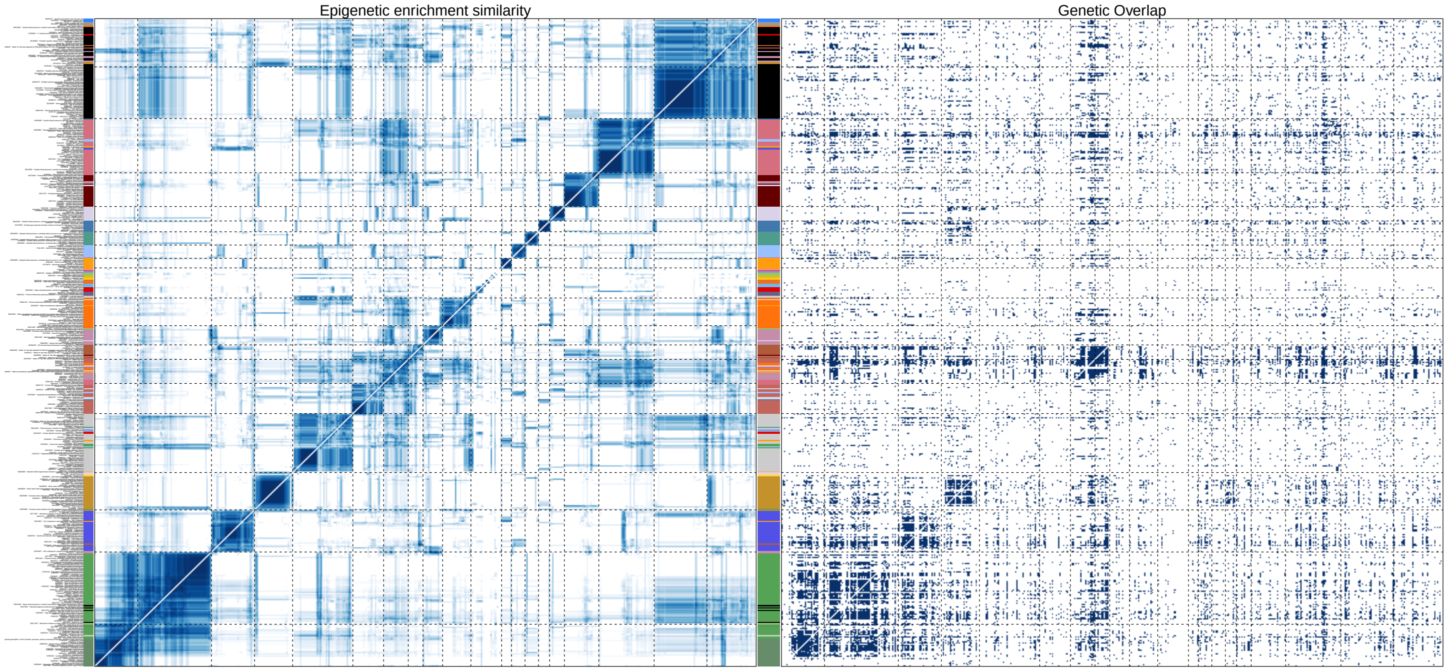


**Figure S33: (A)** Trait-trait similarity matrix by cosine similarity of epigenetic enrichments (left) and by 0/1 genetic similarity (defined as $> 5\%$ jaccard similarity of lead SNPs overlapping when binned into 10k bp bins starting from the start of each chromosome). Matrices are ordered by hierarchical clustering according to Ward's method on the epigenetic matrix.

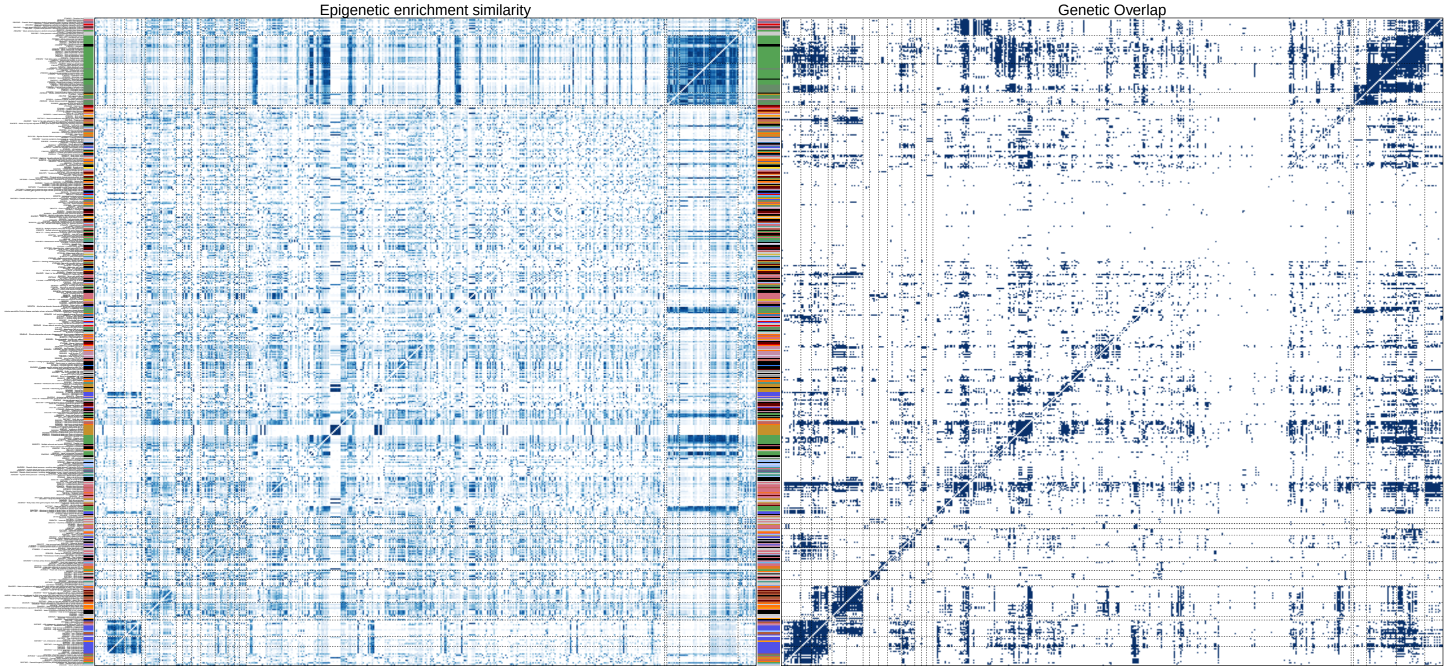


**Figure S33: (B)** Epigenetic similarity (left) and genetic similarity (right) matrices, as above. Matrices ordered by hierarchical clustering according to Ward's method on the genetic similarity matrix.

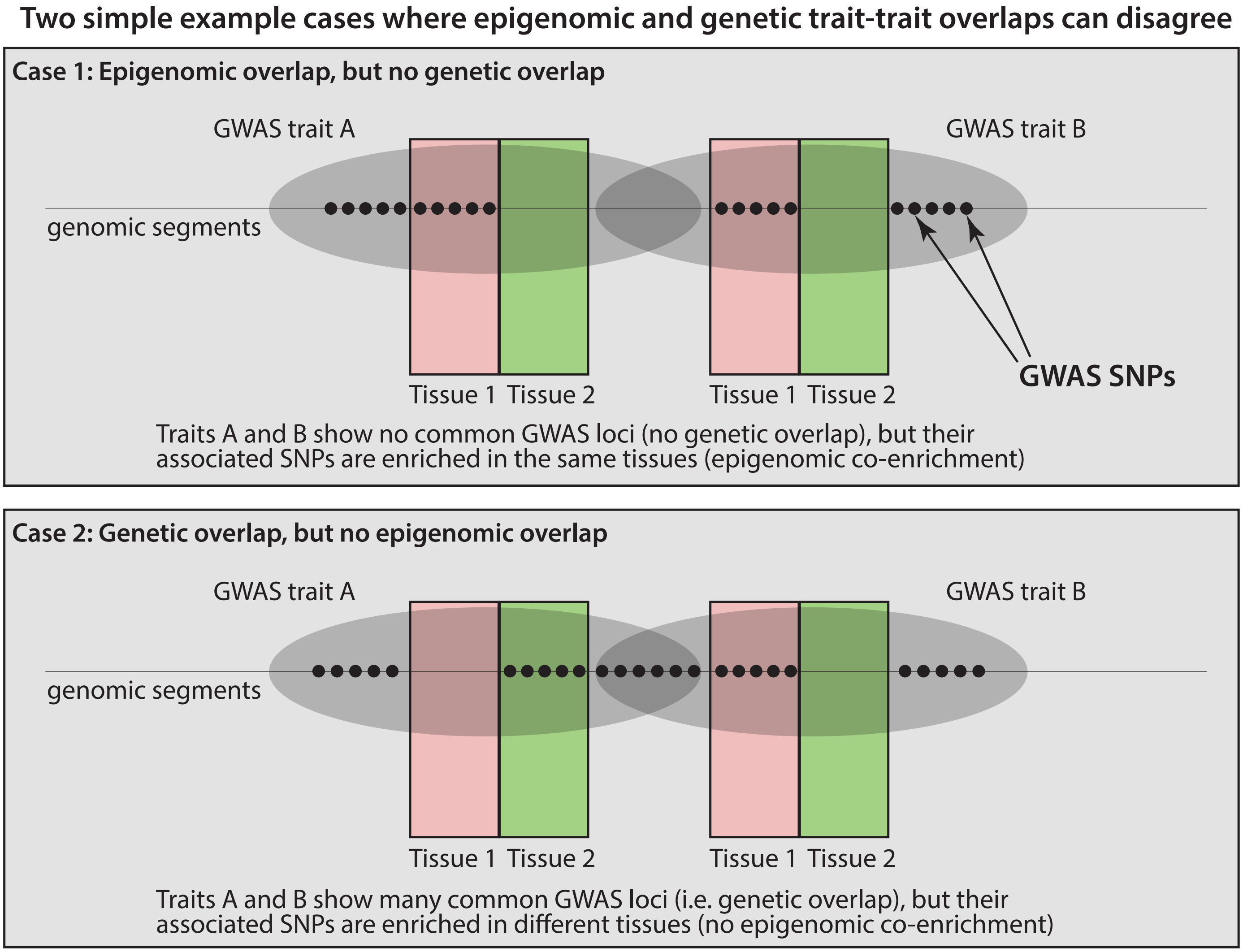


**Figure S34:** Two example cases where the epigenetic and the genetic trait-trait links may not agree.
