## Supplemental Data S1 for "Integrative analysis of 10,000 epigenomic maps across 800 samples for regulatory genomics and disease dissection"

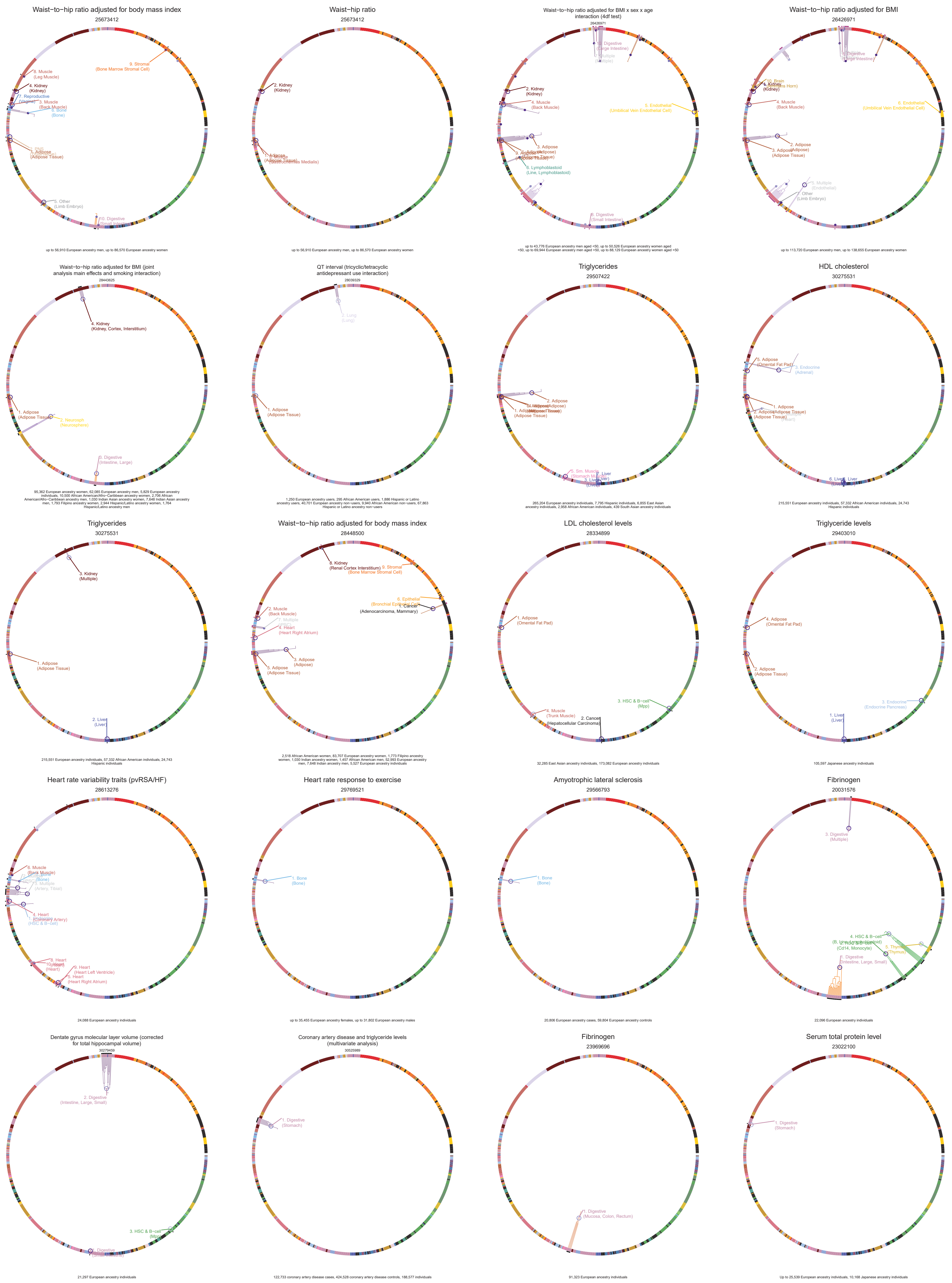

Waist−to−hip ratio adjusted for body mass index
25673412
up to 56,910 European ancestry men, up to 86,570 European ancestry women
Waist−hip ratio
25673412
up to 56,910 European ancestry men, up to 86,570 European ancestry women
Waist−to−hip ratio adjusted for BMI x sex x age interaction (4df test)
26426971
up to 43,776 European ancestry men aged <50, up to 50,526 European ancestry women aged <50, up to 69,944 European ancestry men aged >50, up to 88,129 European ancestry women aged >50
Waist−to−hip ratio adjusted for BMI
26426971
up to 113,720 European ancestry men, up to 138,655 European ancestry women
Waist−to−hip ratio adjusted for BMI (joint analysis main effects and smoking interaction)
28443625
95,362 European ancestry women, 62,085 European ancestry men, 5,829 European ancestry individuals, 10,500 African American/Afro−Caribbean ancestry women, 2,706 African American/Afro−Caribbean ancestry men, 1,030 Indian Asian ancestry women, 7,648 Indian Asian ancestry men, 1,793 Filipino ancestry women, 2,944 Hispanic/Latino ancestry women, 1,764 Hispanic/Latino ancestry men
QT interval (tricyclic/tetracyclic antidepressant use interaction)
28039329
1,250 European ancestry users, 295 African American users, 1,886 Hispanic or Latino ancestry users, 40,701 European ancestry non−users, 9,940 African American non−users, 67,863 Hispanic or Latino ancestry non−users
Triglycerides
29507422
265,204 European ancestry individuals, 7,795 Hispanic individuals, 6,855 East Asian ancestry individuals, 2,958 African American individuals, 439 South Asian ancestry individuals
HDL cholesterol
30275531
215,551 European ancestry individuals, 57,332 African American individuals, 24,743 Hispanic individuals
Triglycerides
30275531
215,551 European ancestry individuals, 57,332 African American individuals, 24,743 Hispanic individuals
Waist−to−hip ratio adjusted for body mass index
28448500
2,518 African American women, 83,707 European ancestry women, 1,773 Filipino ancestry women, 1,030 Indian ancestry women, 1,457 African American men, 52,993 European ancestry men, 7,648 Indian ancestry men, 5,527 European ancestry individuals
LDL cholesterol levels
28334899
32,285 East Asian ancestry individuals, 173,082 European ancestry individuals
Triglyceride levels
29403010
105,597 Japanese ancestry individuals
Heart rate variability traits (pvRSA/HF)
28613276
24,088 European ancestry individuals
Heart rate response to exercise
29769521
up to 35,455 European ancestry females, up to 31,802 European ancestry males
Amyotrophic lateral sclerosis
29566793
20,806 European ancestry cases, 59,804 European ancestry controls
Fibrinogen
20031576
22,096 European ancestry individuals
Dentate gyrus molecular layer volume (corrected for total hippocampal volume)
30279459
21,297 European ancestry individuals
Coronary artery disease and triglyceride levels (multivariate analysis)
30525989
122,733 coronary artery disease cases, 424,528 coronary artery disease controls, 188,577 individuals
Fibrinogen
23969696
91,323 European ancestry individuals
Serum total protein level
23022100
Up to 25,539 European ancestry individuals, 10,168 Japanese ancestry individuals

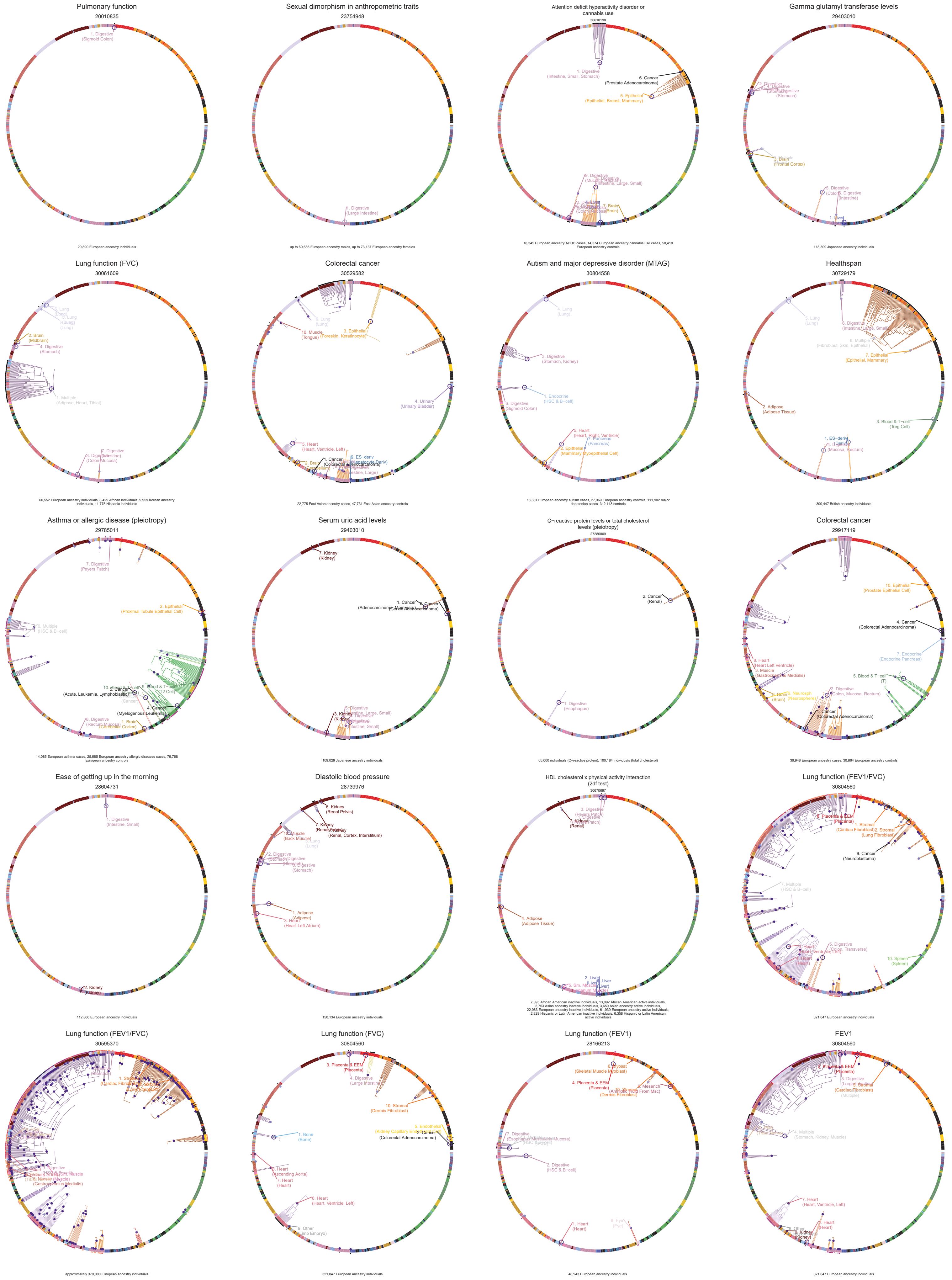
Pulmonary function
20010835
20,890 European ancestry individuals
Sexual dimorphism in anthropometric traits
23754948
up to 60,586 European ancestry males, up to 73,137 European ancestry females
Attention deficit hyperactivity disorder or cannabis use
30610198
18,345 European ancestry ADHD cases, 14,374 European ancestry cannabis use cases, 50,410 European ancestry controls
Gamma glutamyl transferase levels
29403010
118,309 Japanese ancestry individuals
Lung function (FVC)
30061609
60,552 European ancestry individuals, 8,429 African individuals, 9,959 Korean ancestry individuals, 11,775 Hispanic individuals
Colorectal cancer
30529582
22,775 East Asian ancestry cases, 47,731 East Asian ancestry controls
Autism and major depressive disorder (MTAG)
30804558
18,381 European ancestry autism cases, 27,969 European ancestry controls, 111,902 major depression cases, 312,113 controls
Healthspan
30729179
300,447 British ancestry individuals
Asthma or allergic disease (pleiotropy)
29785011
14,085 European asthma cases, 25,685 European ancestry allergic diseases cases, 76,768 European ancestry controls
Serum uric acid levels
29403010
109,029 Japanese ancestry individuals
C−reactive protein levels or total cholesterol levels (pleiotropy)
27286809
65,000 individuals (C−reactive protein), 100,184 individuals (total cholesterol)
Colorectal cancer
29917119
36,948 European ancestry cases, 30,864 European ancestry controls
Ease of getting up in the morning
28604731
112,866 European ancestry individuals
Diastolic blood pressure
28739976
150,134 European ancestry individuals
HDL cholesterol x physical activity interaction (2df test)
30670697
7,395 African American inactive individuals, 13,092 African American active individuals, 2,753 Asian ancestry inactive individuals, 3,650 Asian ancestry active individuals, 22,963 European ancestry inactive individuals, 61,939 European ancestry active individuals, 2,829 Hispanic or Latin American inactive individuals, 6,358 Hispanic or Latin American active individuals
Lung function (FEV1/FVC)
30804560
321,047 European ancestry individuals
Lung function (FEV1/FVC)
30595370
approximately 370,000 European ancestry individuals
Lung function (FVC)
30804560
321,047 European ancestry individuals
Lung function (FEV1)
28166213
48,943 European ancestry individuals
FEV1
30804560
321,047 European ancestry individuals

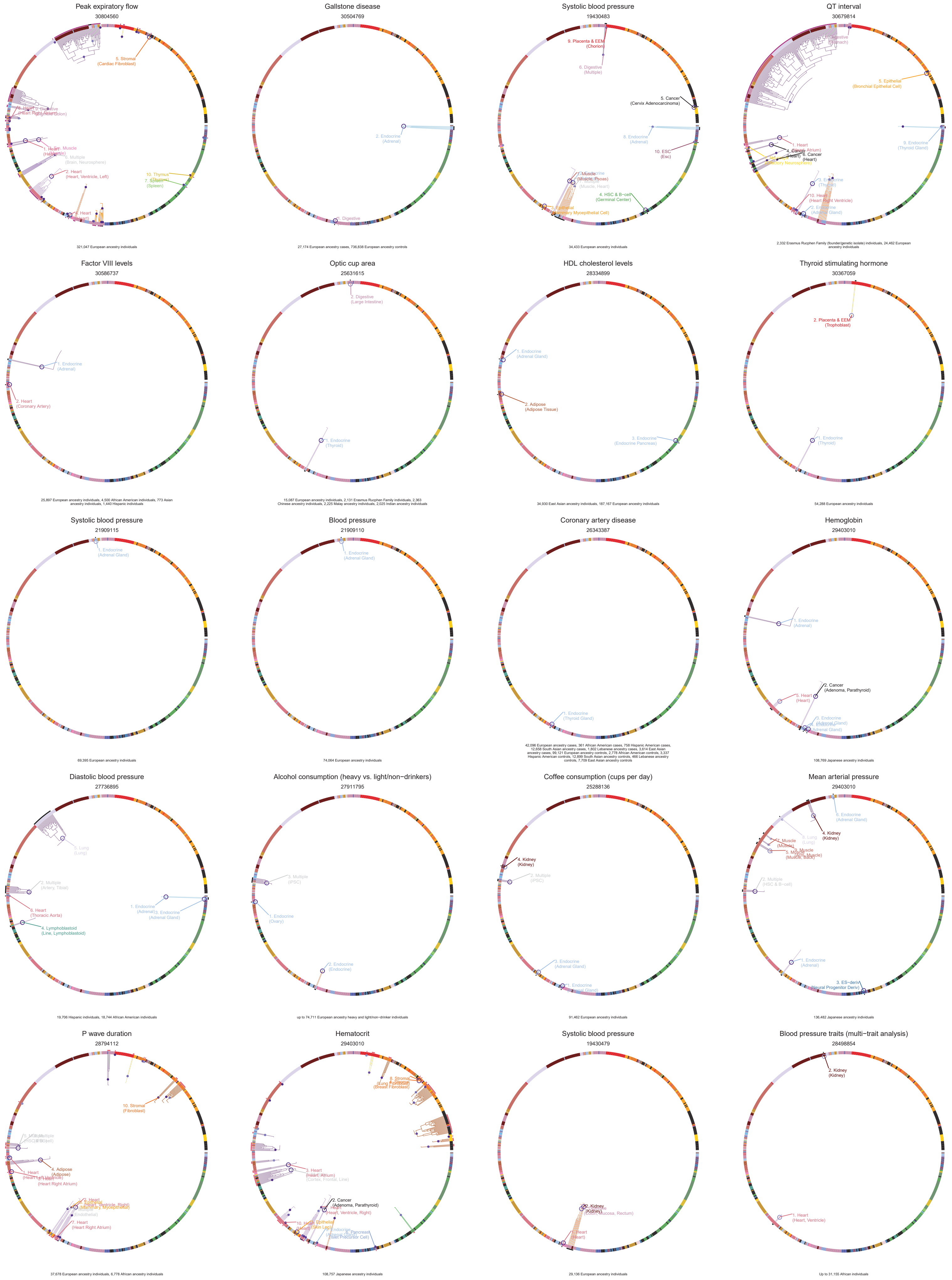
Peak expiratory flow
30804560
5. Stromal
(Cardiac Fibroblast)
2. Heart
(Heart, Ventricle, Left)
10. Thymus
7. Spleen
(Spleen)
321,047 European ancestry individuals
Gallstone disease
30504769
2. Endocrine
(Adrenal)
1. Digestive
27,174 European ancestry cases, 736,838 European ancestry controls
Systolic blood pressure
19430483
9. Placenta & EEM
(Chorion)
6. Digestive
(Multiple)
5. Cancer
(Cervix Adenocarcinoma)
8. Endocrine
(Adrenal)
10. ESC
(Esc)
4. HSC & B−cell
(Germinal Center)
34,433 European ancestry individuals
QT interval
30679814
5. Epithelial
(Bronchial Epithelial Cell)
9. Endocrine
(Thyroid Gland)
6. Cancer
(Heart)
10. Heart
(Heart Right Ventricle)
2,332 Erasmus Rucphen Family (founder/genetic isolate) individuals, 24,462 European ancestry individuals
Factor VIII levels
30586737
1. Endocrine
(Adrenal)
2. Heart
(Coronary Artery)
25,897 European ancestry individuals, 4,500 African American individuals, 773 Asian ancestry individuals, 1,440 Hispanic individuals
Optic cup area
25631615
2. Digestive
(Large Intestine)
1. Endocrine
(Thyroid)
15,087 European ancestry individuals, 2,131 Erasmus Rucphen Family individuals, 2,363 Chinese ancestry individuals, 2,225 Malay ancestry individuals, 2,025 Indian ancestry individuals
HDL cholesterol levels
28334899
1. Endocrine
(Adrenal Gland)
2. Adipose
(Adipose Tissue)
3. Endocrine
(Endocrine Pancreas)
34,930 East Asian ancestry individuals, 187,167 European ancestry individuals
Thyroid stimulating hormone
30367059
2. Placenta & EEM
(Trophoblast)
1. Endocrine
(Thyroid)
54,288 European ancestry individuals
Systolic blood pressure
21909115
1. Endocrine
(Adrenal Gland)
69,395 European ancestry individuals
Blood pressure
21909110
1. Endocrine
(Adrenal Gland)
74,064 European ancestry individuals
Coronary artery disease
26343387
1. Endocrine
(Thyroid Gland)
42,096 European ancestry cases, 361 African American cases, 758 Hispanic American cases, 12,658 South Asian ancestry cases, 1,802 Lebanese ancestry cases, 3,614 East Asian ancestry cases, 99,121 European ancestry controls, 2,778 African American controls, 3,337 Hispanic American controls, 12,899 South Asian ancestry controls, 466 Lebanese ancestry controls, 7,709 East Asian ancestry controls
Hemoglobin
29403010
1. Endocrine
(Adrenal)
2. Cancer
(Adenoma, Parathyroid)
5. Heart
(Heart)
108,769 Japanese ancestry individuals
Diastolic blood pressure
27736895
5. Lung
(Lung)
2. Multiple
(Artery, Tibial)
6. Heart
(Thoracic Aorta)
4. Lymphoblastoid
(Line, Lymphoblastoid)
3. Endocrine
(Adrenal Gland)
19,706 Hispanic individuals, 18,744 African American individuals
Alcohol consumption (heavy vs. light/non−drinkers)
27911795
3. Multiple
(iPSC)
1. Endocrine
(Ovary)
2. Endocrine
(Endocrine)
up to 74,711 European ancestry heavy and light/non−drinker individuals
Coffee consumption (cups per day)
25288136
4. Kidney
(Kidney)
2. Multiple
(iPSC)
3. Endocrine
(Adrenal Gland)
91,462 European ancestry individuals
Mean arterial pressure
29403010
6. Endocrine
(Adrenal Gland)
4. Kidney
(Kidney)
2. Multiple
(HSC & B−cell)
1. Endocrine
(Adrenal)
3. ES−deriv
(Neural Progenitor Deriv)
136,482 Japanese ancestry individuals
P wave duration
28794112
10. Stromal
(Fibroblast)
4. Adipose
(Adipose)
7. Heart
(Heart Right Atrium)
37,678 European ancestry individuals, 6,778 African ancestry individuals
Hematocrit
29403010
3. Heart
(Heart, Atrium)
2. Cancer
(Adenoma, Parathyroid)
108,757 Japanese ancestry individuals
Systolic blood pressure
19430479
1. Heart
(Heart)
29,136 European ancestry individuals
Blood pressure traits (multi−trait analysis)
28498854
2. Kidney
(Kidney)
1. Heart
(Heart, Ventricle)
Up to 31,155 African individuals

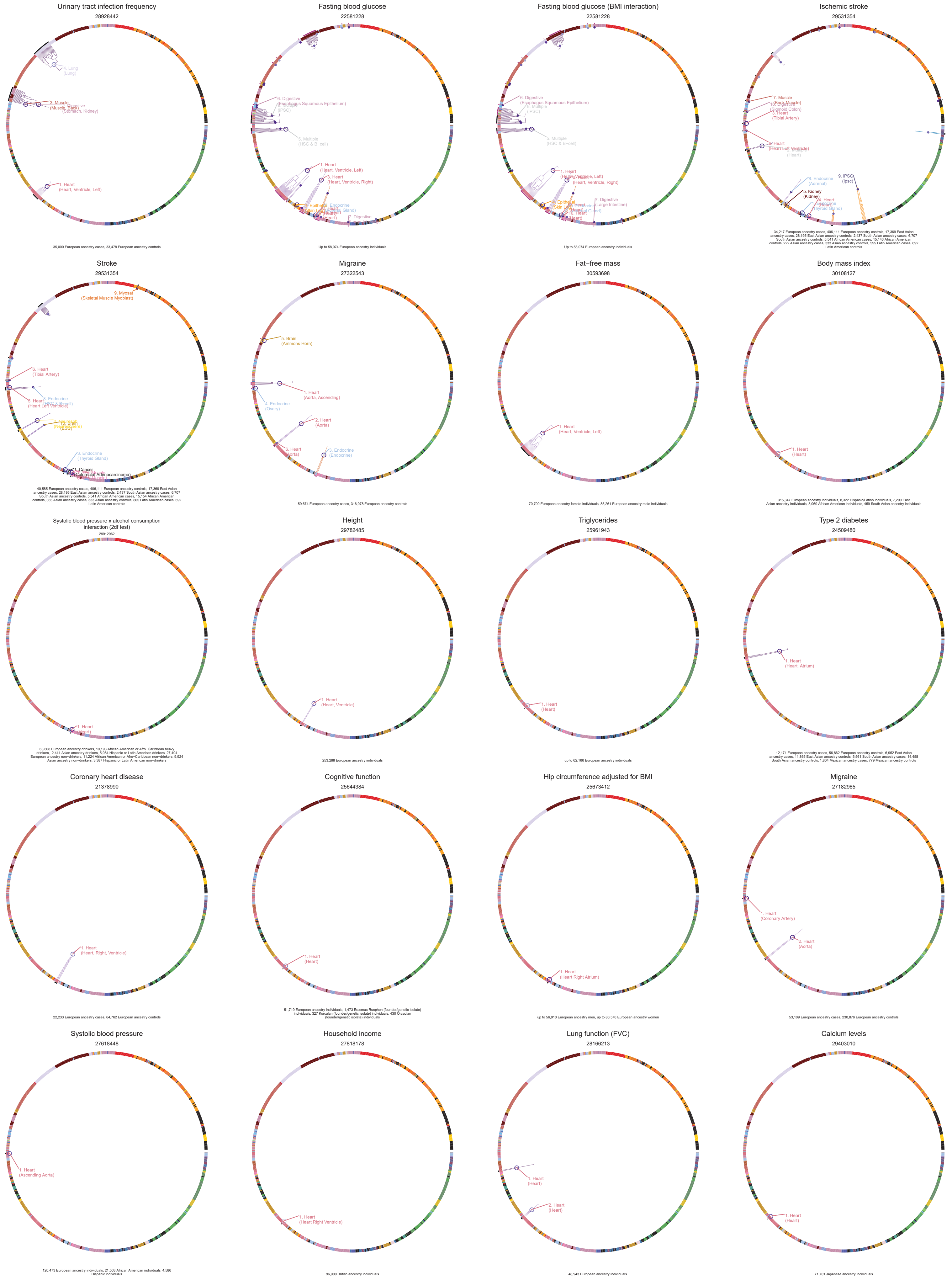

Urinary tract infection frequency
28928442
35,000 European ancestry cases, 33,478 European ancestry controls
Fasting blood glucose
22581228
Up to 58,074 European ancestry individuals
Fasting blood glucose (BMI interaction)
22581228
Up to 58,074 European ancestry individuals
Ischemic stroke
29531354
34,217 European ancestry cases, 406,111 European ancestry controls, 17,369 East Asian ancestry cases, 28,195 East Asian ancestry controls, 2,437 South Asian ancestry cases, 6,707 South Asian ancestry controls, 5,541 African American cases, 15,146 African American controls, 222 Asian ancestry cases, 333 Asian ancestry controls, 555 Latin American cases, 692 Latin American controls
Stroke
29531354
40,585 European ancestry cases, 406,111 European ancestry controls, 17,369 East Asian ancestry cases, 28,195 East Asian ancestry controls, 2,437 South Asian ancestry cases, 6,707 South Asian ancestry controls, 5,541 African American cases, 15,154 African American controls, 365 Asian ancestry cases, 333 Asian ancestry controls, 865 Latin American cases, 692 Latin American controls
Migraine
27322543
59,674 European ancestry cases, 316,078 European ancestry controls
Fat−free mass
30593698
70,700 European ancestry female individuals, 85,261 European ancestry male individuals
Body mass index
30108127
315,347 European ancestry individuals, 8,322 Hispanic/Latino individuals, 7,290 East Asian ancestry individuals, 3,069 African American individuals, 459 South Asian ancestry individuals
Systolic blood pressure x alcohol consumption interaction (2df test)
29912962
63,608 European ancestry drinkers, 10,193 African American or Afro−Caribbean heavy drinkers, 2,441 Asian ancestry drinkers, 5,084 Hispanic or Latin American drinkers, 27,494 European ancestry non−drinkers, 11,224 African American or Afro−Caribbean non−drinkers, 9,924 Asian ancestry non−drinkers, 3,387 Hispanic or Latin American non−drinkers
Height
29782485
253,288 European ancestry individuals
Triglycerides
25961943
up to 62,166 European ancestry individuals
Type 2 diabetes
24509480
12,171 European ancestry cases, 56,862 European ancestry controls, 6,952 East Asian ancestry cases, 11,865 East Asian ancestry controls, 5,561 South Asian ancestry cases, 14,458 South Asian ancestry controls, 1,804 Mexican ancestry cases, 779 Mexican ancestry controls
Coronary heart disease
21378990
22,233 European ancestry cases, 64,762 European ancestry controls
Cognitive function
25644384
51,719 European ancestry individuals, 1,473 Erasmus Rucphen (founder/genetic isolate) individuals, 327 Korculan (founder/genetic isolate) individuals, 430 Orcadian (founder/genetic isolate) individuals
Hip circumference adjusted for BMI
25673412
up to 56,910 European ancestry men, up to 86,570 European ancestry women
Migraine
27182965
53,109 European ancestry cases, 230,876 European ancestry controls
Systolic blood pressure
27618448
120,473 European ancestry individuals, 21,503 African American individuals, 4,586 Hispanic individuals
Household income
27818178
96,900 British ancestry individuals
Lung function (FVC)
28166213
48,943 European ancestry individuals
Calcium levels
29403010
71,701 Japanese ancestry individuals

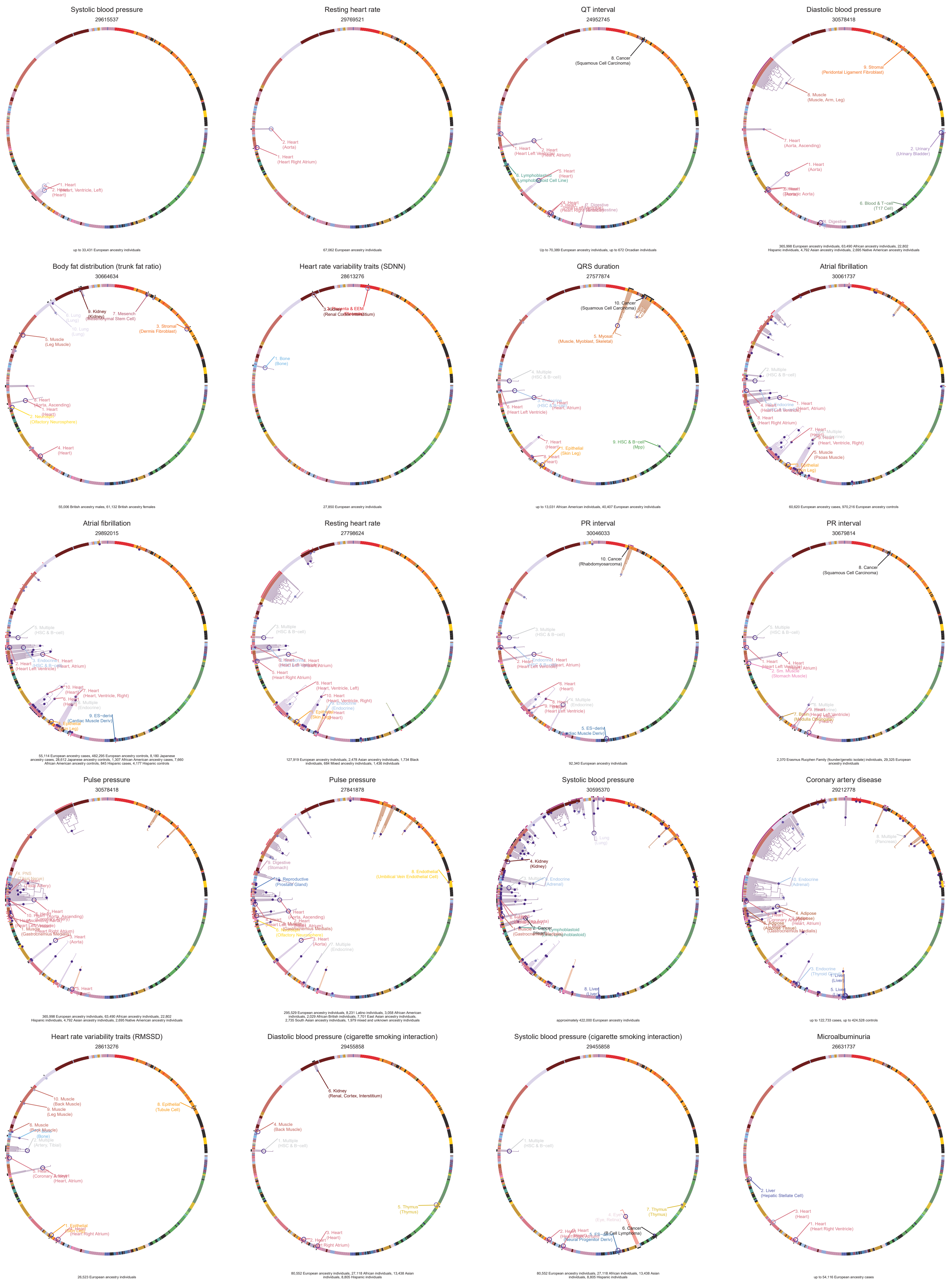
Systolic blood pressure
29615537
up to 33,431 European ancestry individuals
Resting heart rate
29769521
67,062 European ancestry individuals
QT interval
24952745
Up to 70,389 European ancestry individuals, up to 672 Orcadian individuals
Diastolic blood pressure
30578418
365,998 European ancestry individuals, 63,490 African ancestry individuals, 22,802 Hispanic individuals, 4,792 Asian ancestry individuals, 2,695 Native American ancestry individuals
Body fat distribution (trunk fat ratio)
30664634
55,006 British ancestry males, 61,132 British ancestry females
Heart rate variability traits (SDNN)
28613276
27,850 European ancestry individuals
QRS duration
27577874
up to 13,031 African American individuals, 40,407 European ancestry individuals
Atrial fibrillation
30061737
60,620 European ancestry cases, 970,216 European ancestry controls
Atrial fibrillation
29892015
55,114 European ancestry cases, 482,295 European ancestry controls, 8,180 Japanese ancestry cases, 28,612 Japanese ancestry controls, 1,307 African American ancestry cases, 7,660 African American ancestry controls, 845 Hispanic cases, 4,177 Hispanic controls
Resting heart rate
27798624
127,919 European ancestry individuals, 2,478 Asian ancestry individuals, 1,734 Black individuals, 684 Mixed ancestry individuals, 1,436 individuals
PR interval
30046033
92,340 European ancestry individuals
PR interval
30679814
2,370 Erasmus Rucphen Family (founder/genetic isolate) individuals, 29,325 European ancestry individuals
Pulse pressure
30578418
365,998 European ancestry individuals, 63,490 African ancestry individuals, 22,802 Hispanic individuals, 4,792 Asian ancestry individuals, 2,695 Native American ancestry individuals
Pulse pressure
27841878
295,529 European ancestry individuals, 8,231 Latino individuals, 3,058 African American individuals, 2,029 African British individuals, 7,701 East Asian ancestry individuals, 2,735 South Asian ancestry individuals, 1,979 mixed and unknown ancestry individuals
Systolic blood pressure
30595370
approximately 422,000 European ancestry individuals
Coronary artery disease
29212778
up to 122,733 cases, up to 424,528 controls
Heart rate variability traits (RMSSD)
28613276
26,523 European ancestry individuals
Diastolic blood pressure (cigarette smoking interaction)
29455858
80,552 European ancestry individuals, 27,118 African individuals, 13,438 Asian individuals, 8,805 Hispanic individuals
Systolic blood pressure (cigarette smoking interaction)
29455858
80,552 European ancestry individuals, 27,118 African individuals, 13,438 Asian individuals, 8,805 Hispanic individuals
Microalbuminuria
26631737
up to 54,116 European ancestry cases

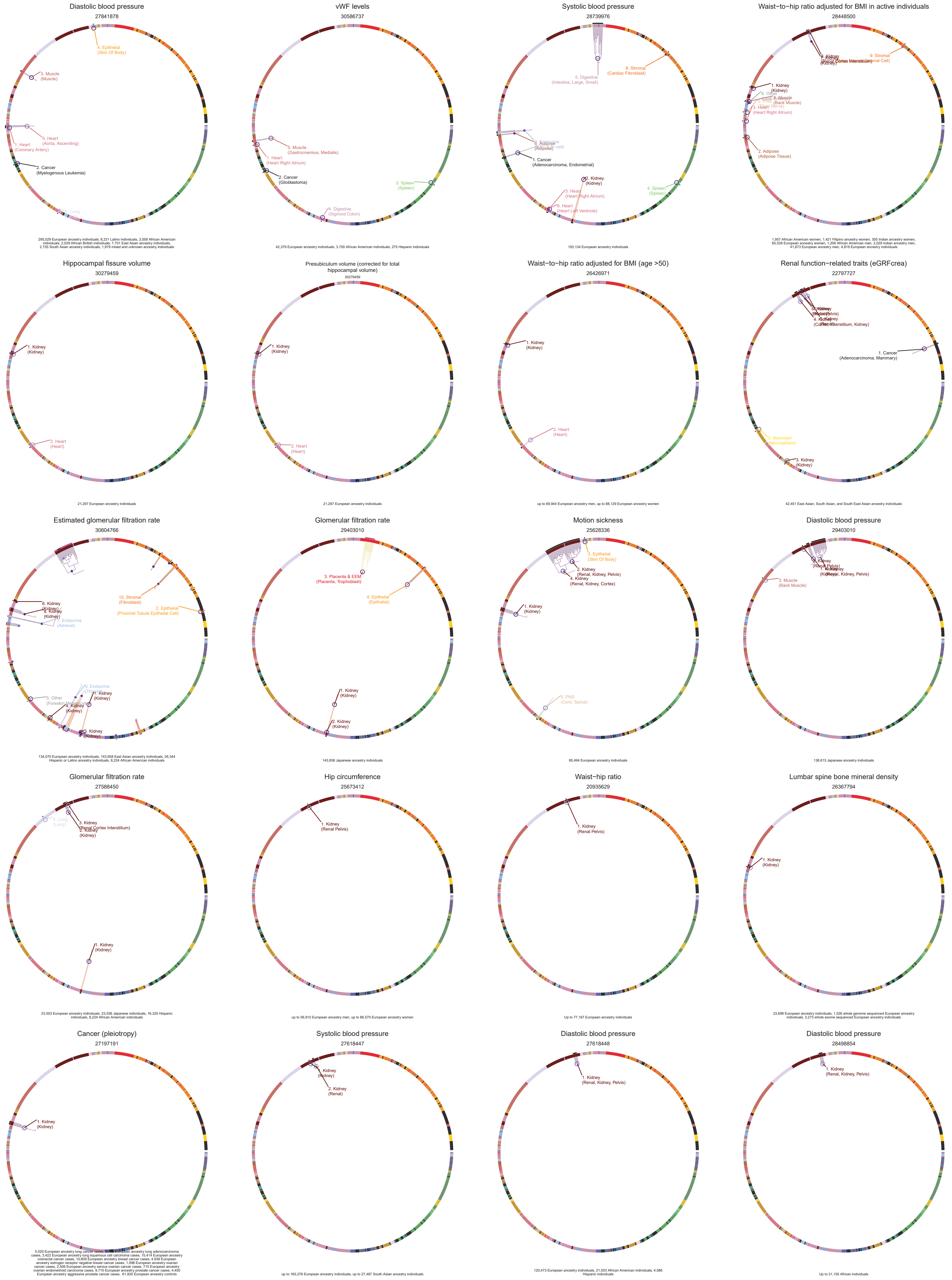

Diastolic blood pressure
27841878
4. Epithelial (Skin Of Body)
3. Muscle (Muscle)
5. Heart (Aorta, Ascending)
1. Heart (Coronary Artery)
2. Cancer (Myelogenous Leukemia)
295,529 European ancestry individuals, 8,231 Latino individuals, 3,058 African American individuals, 2,029 African British individuals, 7,701 East Asian ancestry individuals, 2,735 South Asian ancestry individuals, 1,979 mixed and unknown ancestry individuals
vWF levels
30586737
5. Muscle (Gastrocnemius, Medialis)
1. Heart (Heart Right Atrium)
2. Cancer (Glioblastoma)
3. Spleen (Spleen)
4. Digestive (Sigmoid Colon)
42,379 European ancestry individuals, 3,700 African American individuals, 275 Hispanic individuals
Systolic blood pressure
28739976
5. Digestive (Intestine, Large, Small)
8. Stromal (Cardiac Fibroblast)
6. Adipose (Adipose)
1. Cancer (Adenocarcinoma, Endometrial)
2. Kidney (Kidney)
3. Heart (Heart Right Atrium)
9. Heart (Heart Left Ventricle)
4. Spleen (Spleen)
150,134 European ancestry individuals
Waist−to−hip ratio adjusted for BMI in active individuals
28448500
9. Stromal
1. Kidney (Kidney)
6. Other
4. Muscle (Back Muscle)
3. Heart (Heart Right Atrium)
2. Adipose (Adipose Tissue)
1,957 African American women, 1,421 Filipino ancestry women, 305 Indian ancestry women, 65,539 European ancestry women, 1,206 African American men, 2,029 Indian ancestry men, 41,673 European ancestry men, 4,819 European ancestry individuals
Hippocampal fissure volume
30279459
1. Kidney (Kidney)
2. Heart (Heart)
21,297 European ancestry individuals
Presubiculum volume (corrected for total hippocampal volume)
30279459
1. Kidney (Kidney)
2. Heart (Heart)
21,297 European ancestry individuals
Waist−to−hip ratio adjusted for BMI (age >50)
26426971
1. Kidney (Kidney)
2. Heart (Heart)
up to 69,944 European ancestry men, up to 88,129 European ancestry women
Renal function−related traits (eGRFcrea)
22797727
4. Kidney (Cortex Interstitium, Kidney)
1. Cancer (Adenocarcinoma, Mammary)
2. Neurosph (Neurosphere)
3. Kidney (Kidney)
42,451 East Asian, South Asian, and South East Asian ancestry individuals
Estimated glomerular filtration rate
30604766
6. Kidney (Kidney)
8. Kidney (Kidney)
7. Endocrine (Adrenal)
10. Stromal (Fibroblast)
2. Epithelial (Proximal Tubule Epithelial Cell)
9. Endocrine
5. Other
1. Kidney (Kidney)
4. Kidney (Kidney)
3. Kidney (Kidney)
134,070 European ancestry individuals, 143,658 East Asian ancestry individuals, 26,344 Hispanic or Latino ancestry individuals, 8,224 African American individuals
Glomerular filtration rate
29403010
3. Placenta & EEM (Placenta, Trophoblast)
4. Epithelial (Epithelial)
1. Kidney (Kidney)
2. Kidney (Kidney)
143,658 Japanese ancestry individuals
Motion sickness
25628336
3. Epithelial (Skin Of Body)
2. Kidney (Renal, Kidney, Pelvis)
4. Kidney (Renal, Kidney, Cortex)
1. Kidney (Kidney)
5. PNS (Cord, Spinal)
80,494 European ancestry individuals
Diastolic blood pressure
29403010
3. Muscle (Back Muscle)
136,615 Japanese ancestry individuals
Glomerular filtration rate
27588450
4. Lung (Lung)
3. Kidney (Renal Cortex Interstitium)
2. Kidney (Kidney)
1. Kidney (Kidney)
23,553 European ancestry individuals, 23,536 Japanese individuals, 16,325 Hispanic individuals, 8,224 African American individuals
Hip circumference
25673412
1. Kidney (Renal Pelvis)
up to 56,910 European ancestry men, up to 86,570 European ancestry women
Waist−hip ratio
20935629
1. Kidney (Renal Pelvis)
Up to 77,167 European ancestry individuals
Lumbar spine bone mineral density
26367794
1. Kidney (Kidney)
23,699 European ancestry individuals, 1,526 whole genome sequenced European ancestry individuals, 3,273 whole exome sequenced European ancestry individuals
Cancer (pleiotropy)
27197191
1. Kidney (Kidney)
Systolic blood pressure
27618447
2. Kidney (Renal)
up to 165,276 European ancestry individuals, up to 27,487 South Asian ancestry individuals
Diastolic blood pressure
27618448
1. Kidney (Renal, Kidney, Pelvis)
120,473 European ancestry individuals, 21,503 African American individuals, 4,586 Hispanic individuals
Diastolic blood pressure
28498854
1. Kidney (Renal, Kidney, Pelvis)
Up to 31,155 African individuals

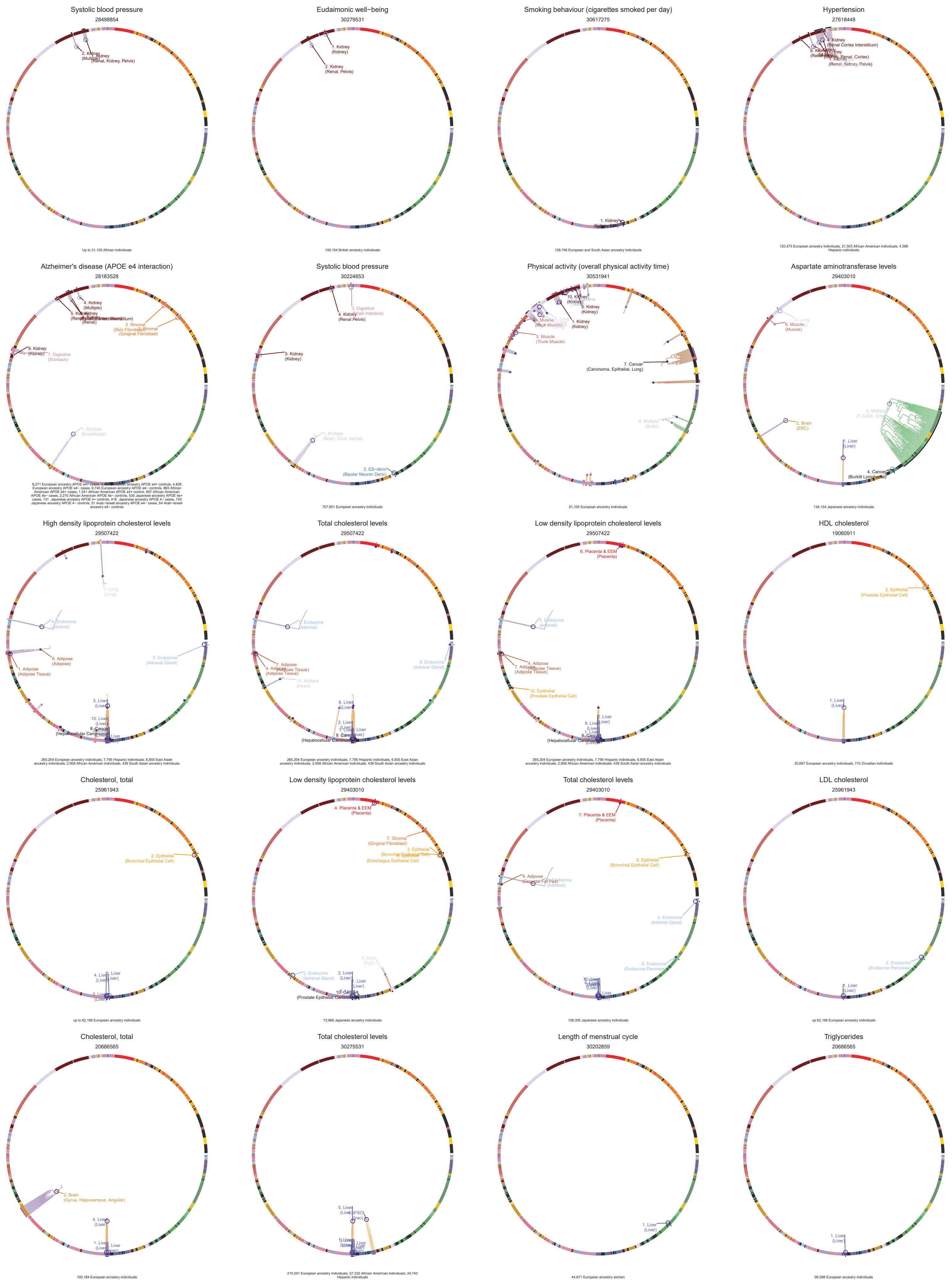
Systolic blood pressure
28498854
Up to 31,155 African individuals
Eudaimonic well−being
30279531
108,154 British ancestry individuals
Smoking behaviour (cigarettes smoked per day)
30617275
128,746 European and South Asian ancestry individuals
Hypertension
27618448
120,473 European ancestry individuals, 21,503 African American individuals, 4,586 Hispanic individuals
Alzheimer's disease (APOE e4 interaction)
28183528
Systolic blood pressure
30224653
757,601 European ancestry individuals
Physical activity (overall physical activity time)
30531941
91,105 European ancestry individuals
Aspartate aminotransferase levels
29403010
134,154 Japanese ancestry individuals
High density lipoprotein cholesterol levels
29507422
265,204 European ancestry individuals, 7,795 Hispanic individuals, 6,855 East Asian ancestry individuals, 2,958 African American individuals, 439 South Asian ancestry individuals
Total cholesterol levels
29507422
265,204 European ancestry individuals, 7,795 Hispanic individuals, 6,855 East Asian ancestry individuals, 2,958 African American individuals, 439 South Asian ancestry individuals
Low density lipoprotein cholesterol levels
29507422
265,204 European ancestry individuals, 7,795 Hispanic individuals, 6,855 East Asian ancestry individuals, 2,958 African American individuals, 439 South Asian ancestry individuals
HDL cholesterol
19060911
20,697 European ancestry individuals, 715 Orcadian individuals
Cholesterol, total
25961943
up to 62,166 European ancestry individuals
Low density lipoprotein cholesterol levels
29403010
72,866 Japanese ancestry individuals
Total cholesterol levels
29403010
128,305 Japanese ancestry individuals
LDL cholesterol
25961943
up 62,166 European ancestry individuals
Cholesterol, total
20686565
100,184 European ancestry individuals
Total cholesterol levels
30275531
215,551 European ancestry individuals, 57,332 African American individuals, 24,743 Hispanic individuals
Length of menstrual cycle
30202859
44,871 European ancestry women
Triglycerides
20686565
96,598 European ancestry individuals

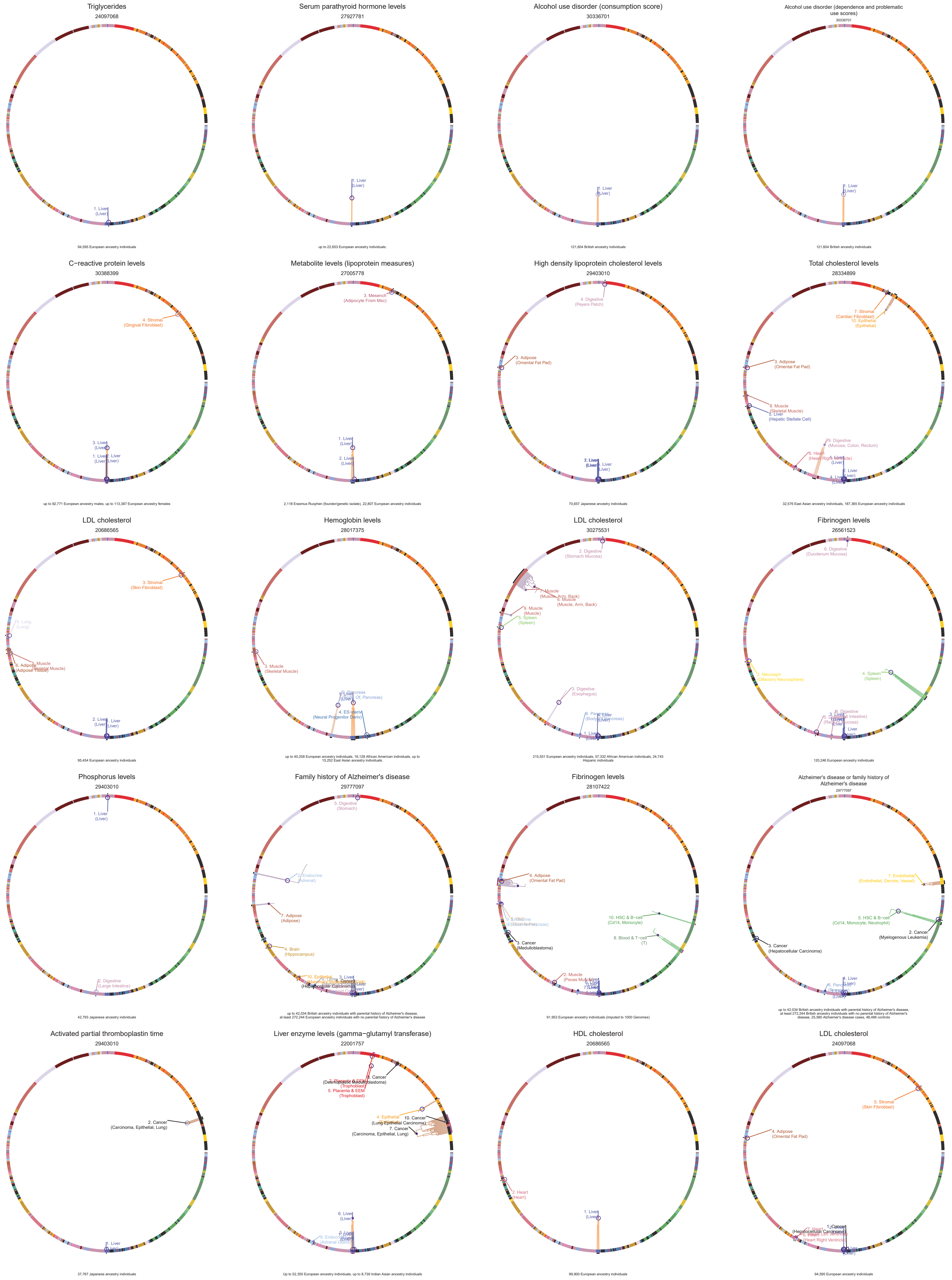
Triglycerides
24097068
94,595 European ancestry individuals
Serum parathyroid hormone levels
27927781
up to 22,653 European ancestry individuals.
Alcohol use disorder (consumption score)
30336701
121,604 British ancestry individuals
Alcohol use disorder (dependence and problematic use scores)
30336701
121,604 British ancestry individuals
C−reactive protein levels
30388399
up to 92,771 European ancestry males, up to 113,387 European ancestry females
Metabolite levels (lipoprotein measures)
27005778
2,118 Erasmus Rucphen (founder/genetic isolate), 22,807 European ancestry individuals
High density lipoprotein cholesterol levels
29403010
70,657 Japanese ancestry individuals
Total cholesterol levels
28334899
32,576 East Asian ancestry individuals, 187,365 European ancestry individuals
LDL cholesterol
20686565
95,454 European ancestry individuals
Hemoglobin levels
28017375
up to 40,258 European ancestry individuals, 16,128 African American individuals, up to 15,252 East Asian ancestry individuals
LDL cholesterol
30275531
215,551 European ancestry individuals, 57,332 African American individuals, 24,743 Hispanic individuals
Fibrinogen levels
26561523
120,246 European ancestry individuals
Phosphorus levels
29403010
42,793 Japanese ancestry individuals
Family history of Alzheimer's disease
29777097
up to 42,034 British ancestry individuals with parental history of Alzheimer's disease, at least 272,244 European ancestry individuals with no parental history of Alzheimer's disease
Fibrinogen levels
28107422
91,953 European ancestry individuals (imputed to 1000 Genomes)
Alzheimer's disease or family history of Alzheimer's disease
29777097
up to 42,034 British ancestry individuals with parental history of Alzheimer's disease, at least 272,244 British ancestry individuals with no parental history of Alzheimer's disease, 25,580 Alzheimer's disease cases, 48,466 controls
Activated partial thromboplastin time
29403010
37,767 Japanese ancestry individuals
Liver enzyme levels (gamma−glutamyl transferase)
22001757
Up to 52,350 European ancestry individuals, up to 8,739 Indian Asian ancestry individuals
HDL cholesterol
20686565
99,900 European ancestry individuals
LDL cholesterol
24097068
94,595 European ancestry individuals

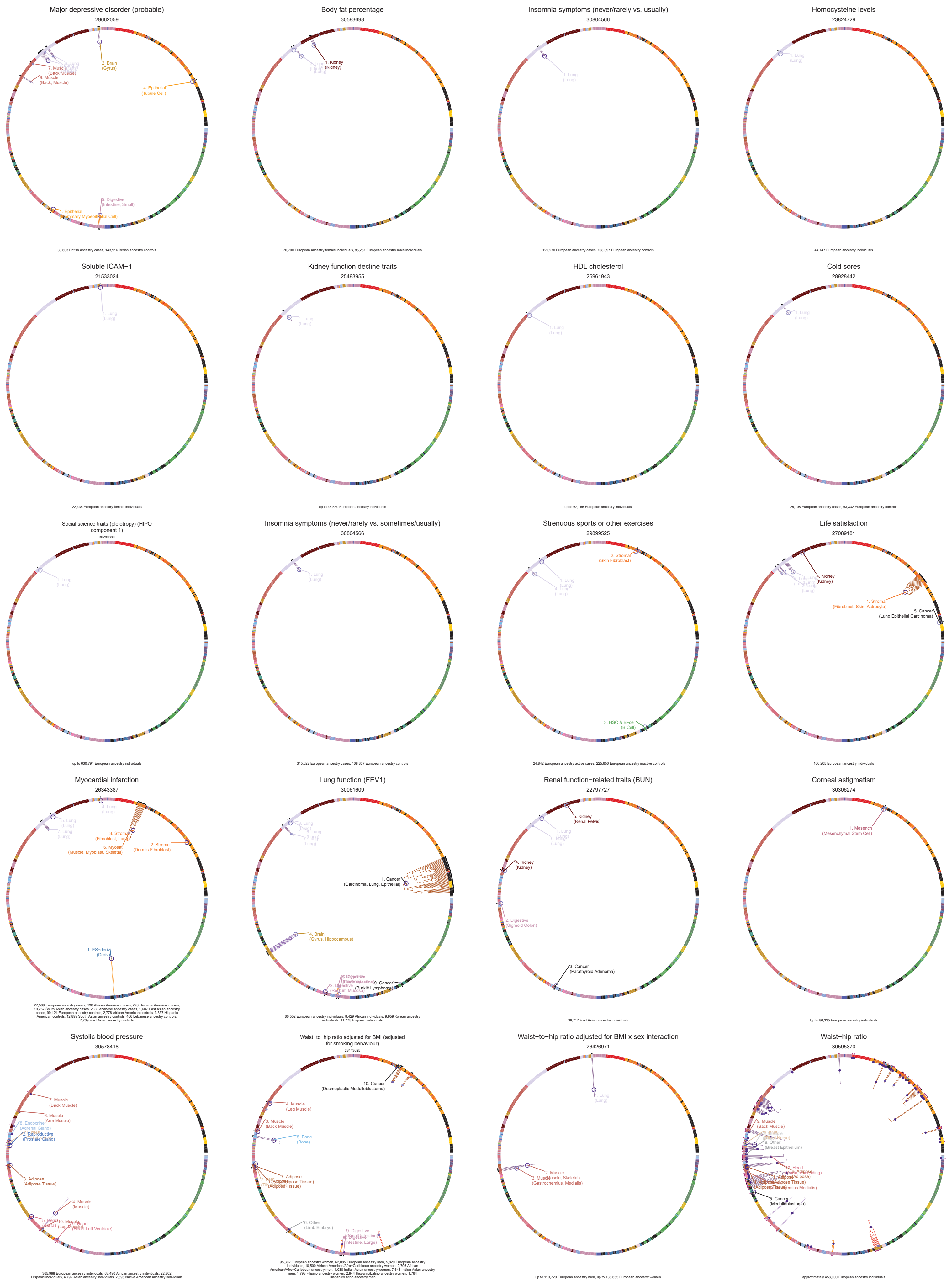
Major depressive disorder (probable)
29662059
30,603 British ancestry cases, 143,916 British ancestry controls
Body fat percentage
30593698
70,700 European ancestry female individuals, 85,261 European ancestry male individuals
Insomnia symptoms (never/rarely vs. usually)
30804566
129,270 European ancestry cases, 108,357 European ancestry controls
Homocysteine levels
23824729
44,147 European ancestry individuals
Soluble ICAM−1
21533024
22,435 European ancestry female individuals
Kidney function decline traits
25493955
up to 45,530 European ancestry individuals
HDL cholesterol
25961943
up to 62,166 European ancestry individuals
Cold sores
28928442
25,108 European ancestry cases, 63,332 European ancestry controls
Social science traits (pleiotropy) (HIPO component 1)
30289880
up to 630,791 European ancestry individuals
Insomnia symptoms (never/rarely vs. sometimes/usually)
30804566
345,022 European ancestry cases, 108,357 European ancestry controls
Strenuous sports or other exercises
29899525
124,842 European ancestry active cases, 225,650 European ancestry inactive controls
Life satisfaction
27089181
166,205 European ancestry individuals
Myocardial infarction
26343387
27,509 European ancestry cases, 130 African American cases, 278 Hispanic American cases, 10,257 South Asian ancestry cases, 288 Lebanese ancestry cases, 1,687 East Asian ancestry cases, 99,121 European ancestry controls, 2,778 African American controls, 3,337 Hispanic American controls, 12,899 South Asian ancestry controls, 466 Lebanese ancestry controls, 7,709 East Asian ancestry controls
Lung function (FEV1)
30061609
60,552 European ancestry individuals, 8,429 African individuals, 9,959 Korean ancestry individuals, 11,775 Hispanic individuals
Renal function−related traits (BUN)
22797727
39,717 East Asian ancestry individuals
Corneal astigmatism
30306274
Up to 86,335 European ancestry individuals
Systolic blood pressure
30578418
365,998 European ancestry individuals, 63,490 African ancestry individuals, 22,802 Hispanic individuals, 4,792 Asian ancestry individuals, 2,695 Native American ancestry individuals
Waist−to−hip ratio adjusted for BMI (adjusted for smoking behaviour)
28443625
95,362 European ancestry women, 62,085 European ancestry men, 5,829 European ancestry individuals, 10,500 African American/Afro−Caribbean ancestry women, 2,706 African American/Afro−Caribbean ancestry men, 1,030 Indian Asian ancestry women, 7,648 Indian Asian ancestry men, 1,793 Filipino ancestry women, 2,944 Hispanic/Latino ancestry women, 1,764 Hispanic/Latino ancestry men
Waist−to−hip ratio adjusted for BMI x sex interaction
26426971
up to 113,720 European ancestry men, up to 138,655 European ancestry women
Waist−hip ratio
30595370
approximately 458,000 European ancestry individuals

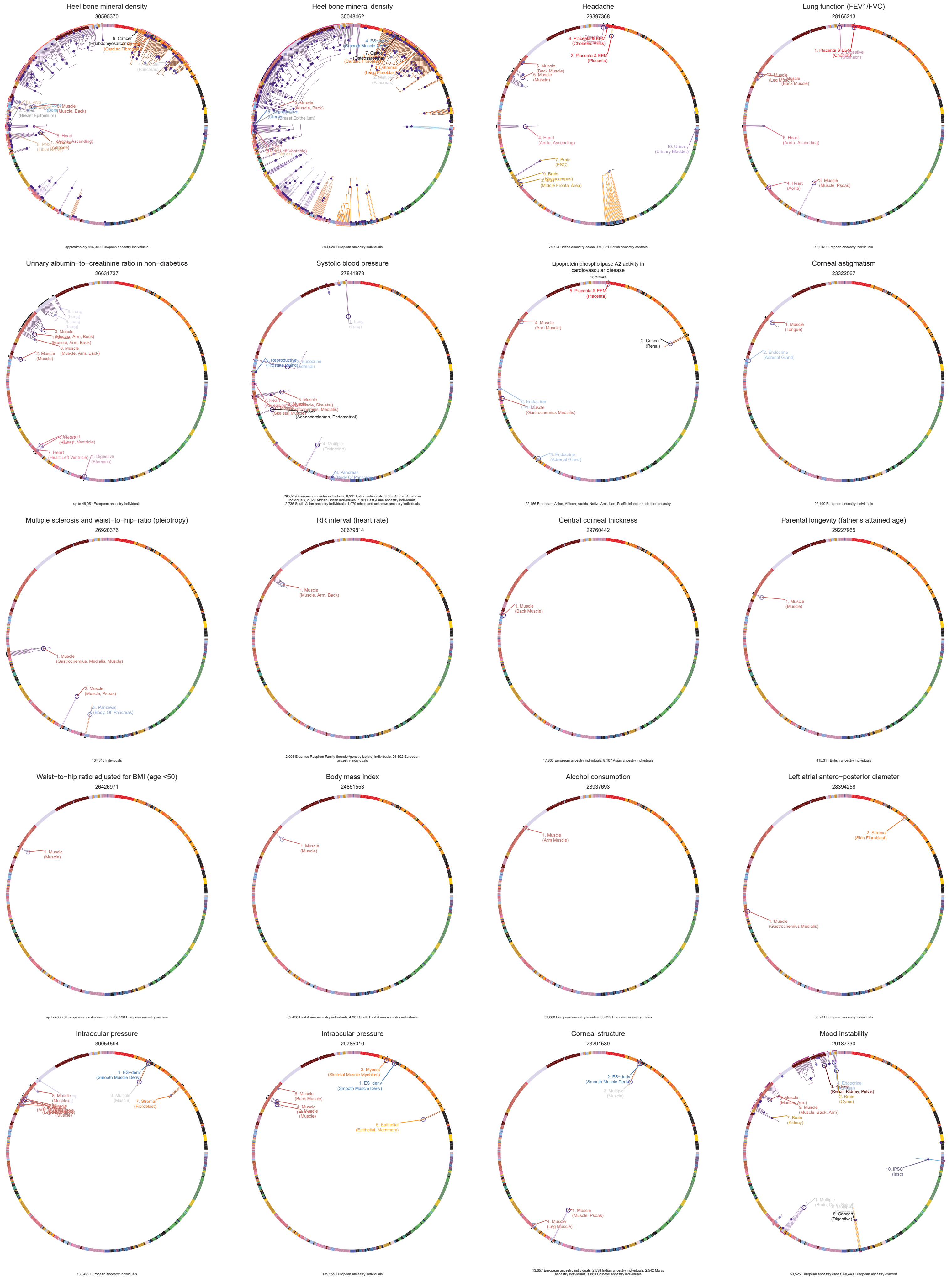
Heel bone mineral density
30595370
approximately 446,000 European ancestry individuals
Heel bone mineral density
30048462
394,929 European ancestry individuals
Headache
29397368
2. Placenta & EEM
(Placenta)
4. Heart
(Aorta, Ascending)
10. Urinary
(Urinary Bladder)
74,461 British ancestry cases, 149,321 British ancestry controls
Lung function (FEV1/FVC)
28166213
6. Heart
(Aorta, Ascending)
4. Heart
(Aorta)
3. Muscle
(Muscle, Psoas)
48,943 European ancestry individuals
Urinary albumin−to−creatinine ratio in non−diabetics
26631737
2. Muscle
(Muscle)
7. Heart
(Heart Left Ventricle)
4. Digestive
(Stomach)
up to 46,051 European ancestry individuals
Systolic blood pressure
27841878
4. Multiple
(Endocrine)
295,529 European ancestry individuals, 8,231 Latino individuals, 3,058 African American individuals, 2,029 African British individuals, 7,701 East Asian ancestry individuals, 2,735 South Asian ancestry individuals, 1,979 mixed and unknown ancestry individuals
Lipoprotein phospholipase A2 activity in cardiovascular disease
28753643
5. Placenta & EEM
(Placenta)
4. Muscle
(Arm Muscle)
2. Cancer
(Renal)
3. Endocrine
(Adrenal Gland)
22,156 European, Asian, African, Arabic, Native American, Pacific Islander and other ancestry
Corneal astigmatism
23322567
1. Muscle
(Tongue)
2. Endocrine
(Adrenal Gland)
22,100 European ancestry individuals
Multiple sclerosis and waist−to−hip−ratio (pleiotropy)
26920376
1. Muscle
(Gastrocnemius, Medialis, Muscle)
2. Muscle
(Muscle, Psoas)
3. Pancreas
(Body, Of, Pancreas)
104,315 individuals
RR interval (heart rate)
30679814
1. Muscle
(Muscle, Arm, Back)
2,006 Erasmus Rucphen Family (founder/genetic isolate) individuals, 26,692 European ancestry individuals
Central corneal thickness
29760442
1. Muscle
(Back Muscle)
17,803 European ancestry individuals, 8,107 Asian ancestry individuals
Parental longevity (father's attained age)
29227965
1. Muscle
(Muscle)
415,311 British ancestry individuals
Waist−to−hip ratio adjusted for BMI (age <50)
26426971
1. Muscle
(Muscle)
up to 43,776 European ancestry men, up to 50,526 European ancestry women
Body mass index
24861553
1. Muscle
(Muscle)
82,438 East Asian ancestry individuals, 4,301 South East Asian ancestry individuals
Alcohol consumption
28937693
1. Muscle
(Arm Muscle)
59,088 European ancestry females, 53,029 European ancestry males
Left atrial antero−posterior diameter
28394258
2. Stromal
(Skin Fibroblast)
1. Muscle
(Gastrocnemius Medialis)
30,201 European ancestry individuals
Intraocular pressure
30054594
1. ES−deriv
(Smooth Muscle Deriv)
7. Stromal
(Fibroblast)
133,492 European ancestry individuals
Intraocular pressure
29785010
3. Myosat
(Skeletal Muscle Myoblast)
1. ES−deriv
(Smooth Muscle Deriv)
5. Epithelial
(Epithelial, Mammary)
139,555 European ancestry individuals
Corneal structure
23291589
2. ES−deriv
(Smooth Muscle Deriv)
3. Multiple
(Muscle)
1. Muscle
(Muscle, Psoas)
4. Muscle
(Leg Muscle)
13,057 European ancestry individuals, 2,538 Indian ancestry individuals, 2,542 Malay ancestry individuals, 1,883 Chinese ancestry individuals
Mood instability
29187730
3. Kidney
(Renal, Kidney, Pelvis)
2. Brain
(Gyrus)
10. iPSC
(Ipsc)
8. Cancer
(Digestive)
53,525 European ancestry cases, 60,443 European ancestry controls

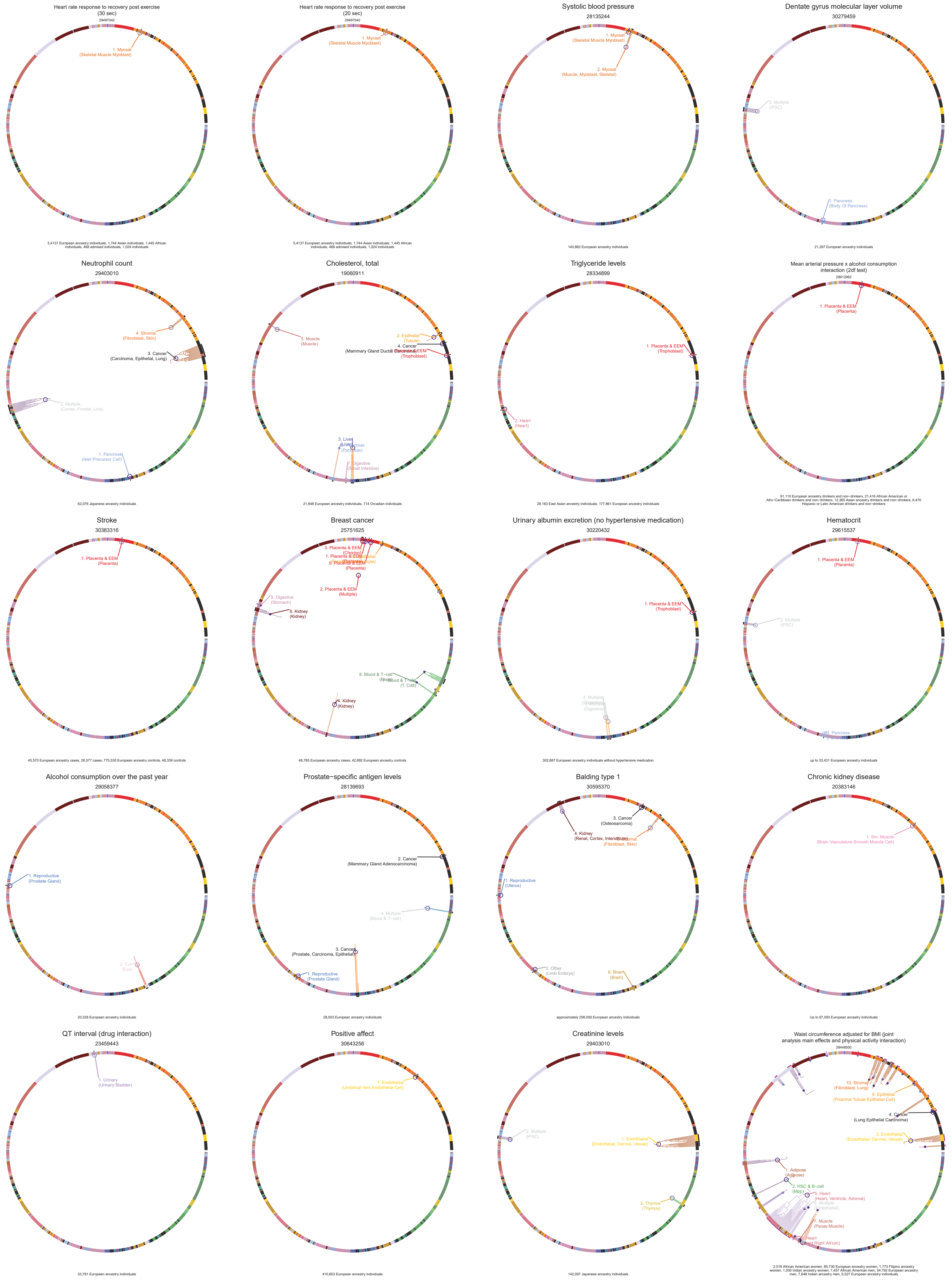

Heart rate response to recovery post exercise (30 sec)
29497042
1. Myosat (Skeletal Muscle Myoblast)
5,4137 European ancestry individuals, 1,744 Asian individuals, 1,445 African individuals, 468 admixed individuals, 1,024 individuals
Heart rate response to recovery post exercise (20 sec)
29497042
1. Myosat (Skeletal Muscle Myoblast)
5,4137 European ancestry individuals, 1,744 Asian individuals, 1,445 African individuals, 468 admixed individuals, 1,024 individuals
Systolic blood pressure
28135244
1. Myosat (Skeletal Muscle Myoblast)
2. Myosat (Muscle, Myoblast, Skeletal)
140,882 European ancestry individuals
Dentate gyrus molecular layer volume
30279459
2. Multiple (iPSC)
1. Pancreas (Body Of Pancreas)
21,297 European ancestry individuals
Neutrophil count
29403010
4. Stromal (Fibroblast, Skin)
3. Cancer (Carcinoma, Epithelial, Lung)
2. Multiple (Cortex, Frontal, Line)
1. Pancreas (Islet Precursor Cell)
62,076 Japanese ancestry individuals
Cholesterol, total
19060911
5. Muscle (Muscle)
2. Epithelial (Tubule)
4. Cancer (Mammary Gland Ductal Carcinoma)
3. Liver
6. Pancreas (Pancreas)
7. Digestive (Small Intestine)
21,848 European ancestry individuals, 714 Orcadian individuals
Triglyceride levels
28334899
1. Placenta & EEM (Trophoblast)
2. Heart (Heart)
28,183 East Asian ancestry individuals, 177,861 European ancestry individuals
Mean arterial pressure x alcohol consumption interaction (2df test)
29912962
1. Placenta & EEM (Placenta)
91,110 European ancestry drinkers and non−drinkers, 21,416 African American or Afro−Caribbean drinkers and non−drinkers, 12,365 Asian ancestry drinkers and non−drinkers, 8,470 Hispanic or Latin American drinkers and non−drinkers
Stroke
30383316
1. Placenta & EEM (Placenta)
45,570 European ancestry cases, 26,577 cases, 775,530 European ancestry controls, 48,339 controls
Breast cancer
25751625
3. Placenta & EEM (Chorion)
1. Placenta & EEM
5. Placenta & EEM (Placenta)
2. Placenta & EEM (Multiple)
9. Digestive (Stomach)
6. Kidney (Kidney)
8. Blood & T−cell
7. Blood & T−cell (T, Cd8)
4. Kidney (Kidney)
46,785 European ancestry cases, 42,892 European ancestry controls
Urinary albumin excretion (no hypertensive medication)
30220432
1. Placenta & EEM (Trophoblast)
3. Multiple
2. Digestive (Digestive)
302,687 European ancestry individuals without hypertensive medication
Hematocrit
29615537
1. Placenta & EEM (Placenta)
2. Multiple (iPSC)
3. Pancreas
up to 33,431 European ancestry individuals
Alcohol consumption over the past year
29058377
1. Reproductive (Prostate Gland)
2. Eye (Eye)
20,328 European ancestry individuals
Prostate−specific antigen levels
28139693
2. Cancer (Mammary Gland Adenocarcinoma)
4. Multiple (Blood & T−cell)
3. Cancer (Prostate, Carcinoma, Epithelial)
1. Reproductive (Prostate Gland)
28,503 European ancestry individuals
Balding type 1
30595370
3. Cancer (Osteosarcoma)
4. Kidney (Renal, Cortex, Interstitium)
5. Stromal (Fibroblast, Skin)
1. Reproductive (Uterus)
2. Other (Limb Embryo)
6. Brain (Brain)
approximately 208,000 European ancestry individuals
Chronic kidney disease
20383146
1. Sm. Muscle (Brain Vasculature Smooth Muscle Cell)
Up to 67,093 European ancestry individuals
QT interval (drug interaction)
23459443
1. Urinary (Urinary Bladder)
33,781 European ancestry individuals
Positive affect
30643256
1. Endothelial (Umbilical Vein Endothelial Cell)
410,603 European ancestry individuals
Creatinine levels
29403010
2. Multiple (iPSC)
1. Endothelial (Endothelial, Dermis, Vessel)
3. Thymus (Thymus)
142,097 Japanese ancestry individuals
Waist circumference adjusted for BMI (joint analysis main effects and physical activity interaction)
28448500
10. Stromal (Fibroblast, Lung)
8. Epithelial (Proximal Tubule Epithelial Cell)
4. Cancer (Lung Epithelial Carcinoma)
3. Endothelial (Endothelial, Dermis, Vessel)
1. Adipose (Adipose)
2. HSC & B−cell (Mpp)
5. Heart (Heart, Ventricle, Adrenal)
6. Multiple (Endothelial)
7. Muscle (Psoas Muscle)
9. Heart (Heart Right Atrium)
2,518 African American women, 85,730 European ancestry women, 1,773 Filipino ancestry women, 1,030 Indian ancestry women, 1,457 African American men, 54,792 European ancestry men, 7,648 Indian ancestry men, 5,527 European ancestry individuals

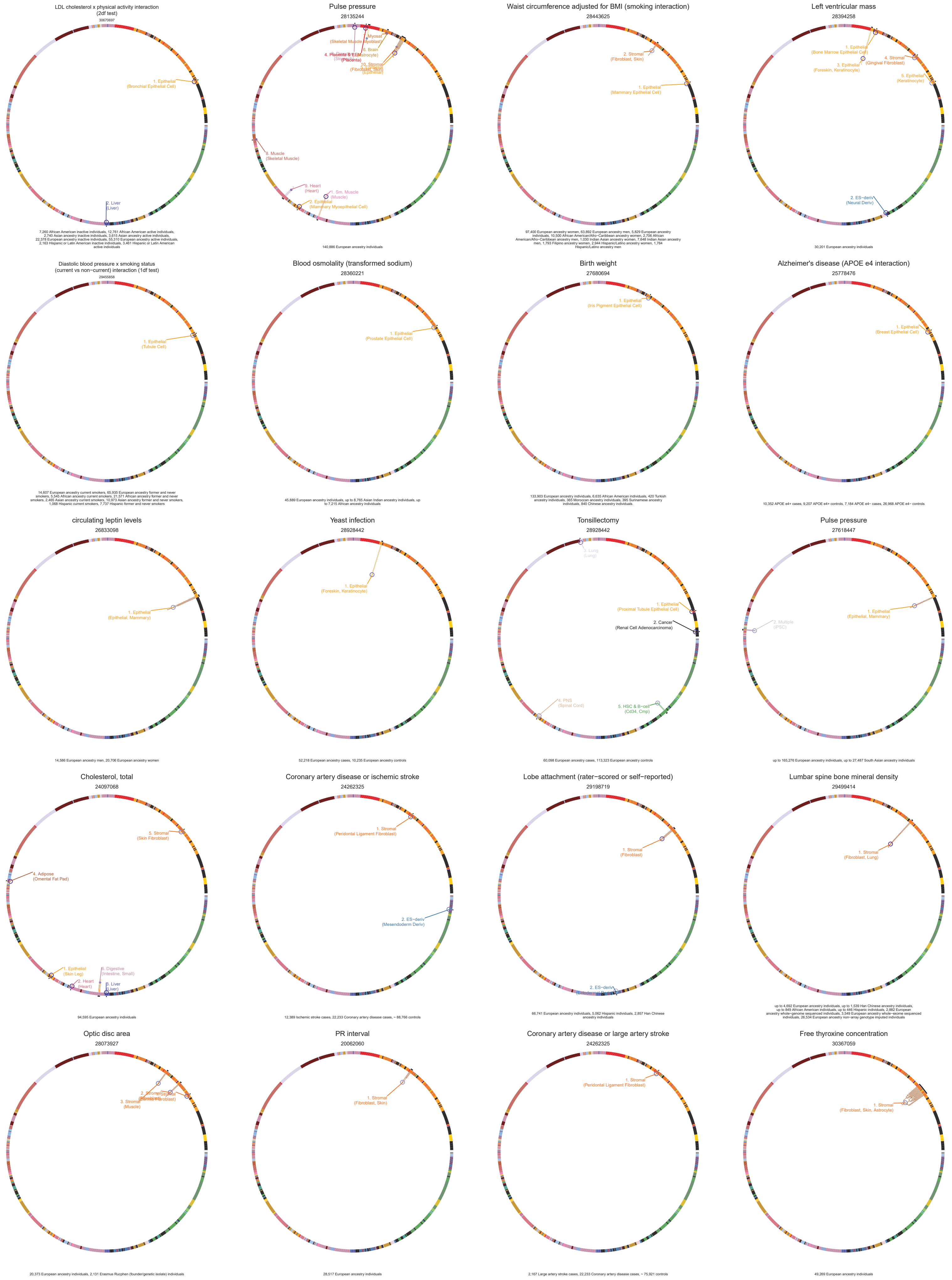

LDL cholesterol x physical activity interaction
(2df test)
30670697
1. Epithelial
(Bronchial Epithelial Cell)
2. Liver
(Liver)
7,260 African American inactive individuals, 12,761 African American active individuals, 2,740 Asian ancestry inactive individuals, 3,615 Asian ancestry active individuals, 22,378 European ancestry inactive individuals, 55,310 European ancestry active individuals, 2,163 Hispanic or Latin American inactive individuals, 3,461 Hispanic or Latin American active individuals
Pulse pressure
28135244
8. Muscle
(Skeletal Muscle)
9. Heart
(Heart)
1. Sm. Muscle
(Muscle)
2. Epithelial
(Mammary Myoepithelial Cell)
140,886 European ancestry individuals
Waist circumference adjusted for BMI (smoking interaction)
28443625
2. Stromal
(Fibroblast, Skin)
1. Epithelial
(Mammary Epithelial Cell)
97,400 European ancestry women, 63,892 European ancestry men, 5,829 European ancestry individuals, 10,500 African American/Afro−Caribbean ancestry women, 2,706 African American/Afro−Caribbean ancestry men, 1,030 Indian Asian ancestry women, 7,648 Indian Asian ancestry men, 1,793 Filipino ancestry women, 2,944 Hispanic/Latino ancestry women, 1,764 Hispanic/Latino ancestry men
Left ventricular mass
28394258
1. Epithelial
(Bone Marrow Epithelial Cell)
3. Epithelial
(Foreskin, Keratinocyte)
4. Stromal
(Gingival Fibroblast)
5. Epithelial
(Keratinocyte)
2. ES−deriv
(Neural Deriv)
30,201 European ancestry individuals
Diastolic blood pressure x smoking status
(current vs non−current) interaction (1df test)
29455858
1. Epithelial
(Tubule Cell)
14,607 European ancestry current smokers, 65,935 European ancestry former and never smokers, 5,545 African ancestry current smokers, 21,571 African ancestry former and never smokers, 2,465 Asian ancestry current smokers, 10,973 Asian ancestry former and never smokers, 1,068 Hispanic current smokers, 7,737 Hispanic former and never smokers
Blood osmolality (transformed sodium)
28360221
1. Epithelial
(Prostate Epithelial Cell)
45,889 European ancestry individuals, up to 8,765 Asian Indian ancestry individuals, up to 7,215 African ancestry individuals
Birth weight
27680694
1. Epithelial
(Iris Pigment Epithelial Cell)
133,903 European ancestry individuals, 6,635 African American individuals, 420 Turkish ancestry individuals, 365 Moroccan ancestry individuals, 395 Surinamese ancestry individuals, 840 Chinese ancestry individuals.
Alzheimer's disease (APOE e4 interaction)
25778476
1. Epithelial
(Breast Epithelial Cell)
10,352 APOE e4+ cases, 9,207 APOE e4+ controls, 7,184 APOE e4− cases, 26,968 APOE e4− controls
circulating leptin levels
26833098
1. Epithelial
(Epithelial, Mammary)
14,586 European ancestry men, 20,706 European ancestry women
Yeast infection
28928442
1. Epithelial
(Foreskin, Keratinocyte)
52,218 European ancestry cases, 10,235 European ancestry controls
Tonsillectomy
28928442
3. Lung
(Lung)
1. Epithelial
(Proximal Tubule Epithelial Cell)
2. Cancer
(Renal Cell Adenocarcinoma)
4. PNS
(Spinal Cord)
5. HSC & B−cell
(Cd34, Cmp)
60,098 European ancestry cases, 113,323 European ancestry controls
Pulse pressure
27618447
2. Multiple
(PSC)
1. Epithelial
(Epithelial, Mammary)
up to 165,276 European ancestry individuals, up to 27,487 South Asian ancestry individuals
Cholesterol, total
24097068
5. Stromal
(Skin Fibroblast)
4. Adipose
(Omental Fat Pad)
1. Epithelial
(Skin Leg)
2. Heart
(Heart)
6. Digestive
(Intestine, Small)
3. Liver
(Liver)
94,595 European ancestry individuals
Coronary artery disease or ischemic stroke
24262325
1. Stromal
(Peridontal Ligament Fibroblast)
2. ES−deriv
(Mesendoderm Deriv)
12,389 Ischemic stroke cases, 22,233 Coronary artery disease cases, ~ 88,766 controls
Lobe attachment (rater−scored or self−reported)
29198719
1. Stromal
(Fibroblast)
2. ES−deriv
66,741 European ancestry individuals, 5,062 Hispanic individuals, 2,857 Han Chinese ancestry individuals
Lumbar spine bone mineral density
29499414
1. Stromal
(Fibroblast, Lung)
up to 4,692 European ancestry individuals, up to 1,539 Han Chinese ancestry individuals, up to 849 African American individuals, up to 446 Hispanic individuals, 2,882 European ancestry whole−genome sequenced individuals, 3,549 European ancestry whole−exome sequenced individuals, 26,534 European ancestry non−array genotype imputed individuals
Optic disc area
28073927
3. Stromal
(Muscle)
20,373 European ancestry individuals, 2,131 Erasmus Rucphen (founder/genetic isolate) individuals
PR interval
20062060
1. Stromal
(Fibroblast, Skin)
28,517 European ancestry individuals
Coronary artery disease or large artery stroke
24262325
1. Stromal
(Peridontal Ligament Fibroblast)
2,167 Large artery stroke cases, 22,233 Coronary artery disease cases, ~ 75,921 controls
Free thyroxine concentration
30367059
1. Stromal
(Fibroblast, Skin, Astrocyte)
49,269 European ancestry individuals

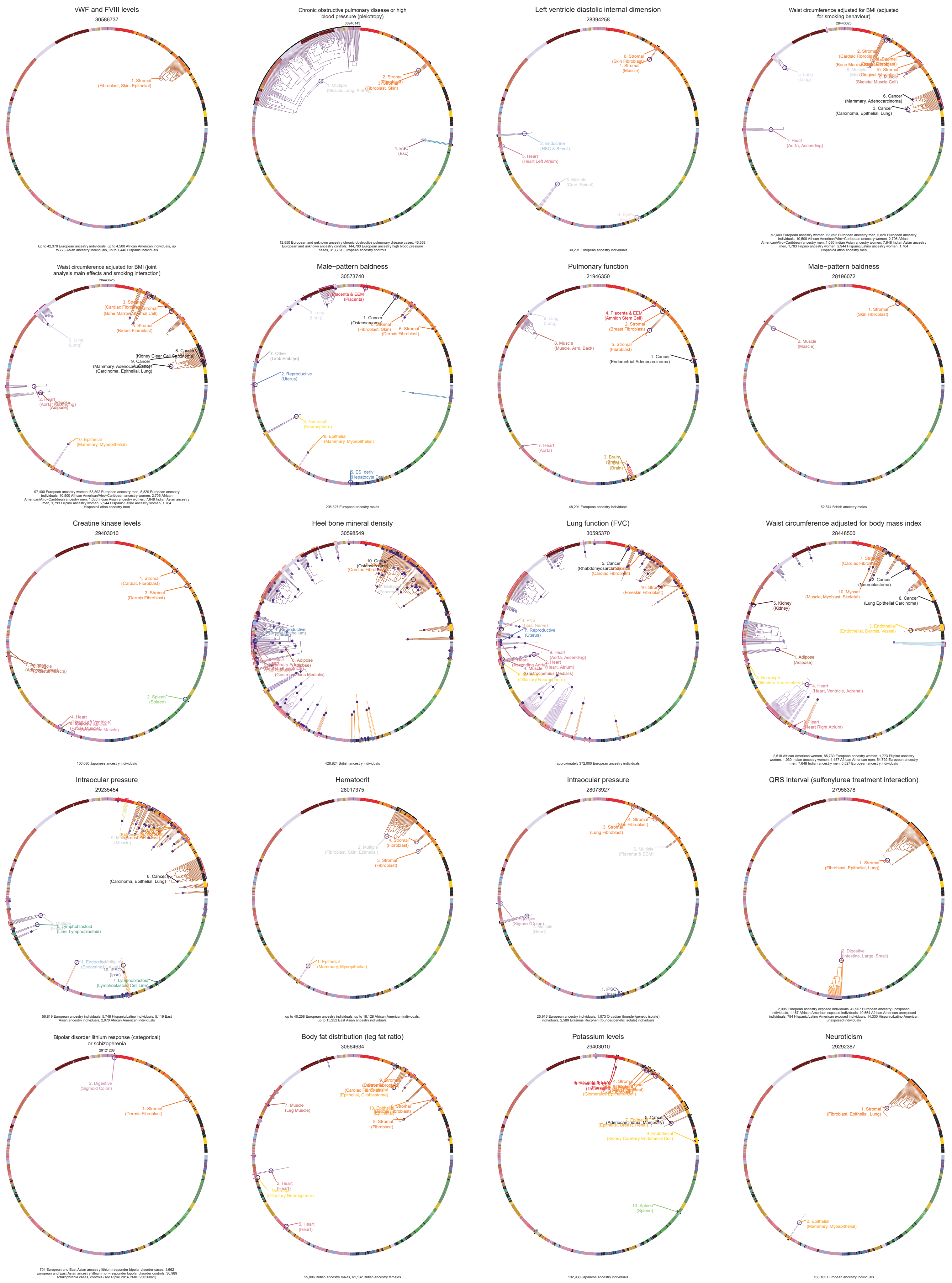
vWF and FVIII levels
30586737
Up to 42,379 European ancestry individuals, up to 4,500 African American individuals, up to 773 Asian ancestry individuals, up to 1,440 Hispanic individuals
Chronic obstructive pulmonary disease or high blood pressure (pleiotropy)
30940143
12,550 European and unknown ancestry chronic obstructive pulmonary disease cases, 46,368 European and unknown ancestry controls, 144,793 European ancestry high blood pressure cases, 313,761 European ancestry controls
Left ventricle diastolic internal dimension
28394258
30,201 European ancestry individuals
Waist circumference adjusted for BMI (adjusted for smoking behaviour)
28443625
97,400 European ancestry women, 63,892 European ancestry men, 5,829 European ancestry individuals, 10,500 African American/Afro−Caribbean ancestry women, 2,706 African American/Afro−Caribbean ancestry men, 1,030 Indian Asian ancestry women, 7,648 Indian Asian ancestry men, 1,793 Filipino ancestry women, 2,944 Hispanic/Latino ancestry women, 1,764 Hispanic/Latino ancestry men
Waist circumference adjusted for BMI (joint analysis main effects and smoking interaction)
28443625
97,400 European ancestry women, 63,892 European ancestry men, 5,829 European ancestry individuals, 10,500 African American/Afro−Caribbean ancestry women, 2,706 African American/Afro−Caribbean ancestry men, 1,030 Indian Asian ancestry women, 7,648 Indian Asian ancestry men, 1,793 Filipino ancestry women, 2,944 Hispanic/Latino ancestry women, 1,764 Hispanic/Latino ancestry men
Male−pattern baldness
30573740
205,327 European ancestry males
Pulmonary function
21946350
48,201 European ancestry individuals
Male−pattern baldness
28196072
52,874 British ancestry males
Creatine kinase levels
29403010
106,080 Japanese ancestry individuals
Heel bone mineral density
30598549
426,824 British ancestry individuals
Lung function (FVC)
30595370
approximately 372,000 European ancestry individuals
Waist circumference adjusted for body mass index
28448500
2,518 African American women, 85,730 European ancestry women, 1,773 Filipino ancestry women, 1,030 Indian ancestry women, 1,457 African American men, 54,792 European ancestry men, 7,648 Indian ancestry men, 5,527 European ancestry individuals
Intraocular pressure
29235454
56,819 European ancestry individuals, 5,748 Hispanic/Latino individuals, 5,119 East Asian ancestry individuals, 2,070 African American individuals
Hematocrit
28017375
up to 40,258 European ancestry individuals, up to 16,128 African American individuals, up to 15,252 East Asian ancestry individuals.
Intraocular pressure
28073927
25,916 European ancestry individuals, 1,073 Orcadian (founder/genetic isolate) individuals, 2,589 Erasmus Rucphen (founder/genetic isolate) individuals
QRS interval (sulfonylurea treatment interaction)
27958378
2,095 European ancestry exposed individuals, 42,907 European ancestry unexposed individuals, 1,167 African American exposed individuals, 10,564 African American unexposed individuals, 794 Hispanic/Latino American exposed individuals, 14,330 Hispanic/Latino American unexposed individuals
Bipolar disorder lithium response (categorical) or schizophrenia
29121268
704 European and East Asian ancestry lithium responder bipolar disorder cases, 1,662 European and East Asian ancestry lithium non−responder bipolar disorder cases, 36,989 schizophrenia cases, controls (see Ripke 2014 PMID:25056061).
Body fat distribution (leg fat ratio)
30664634
55,006 British ancestry males, 61,132 British ancestry females
Potassium levels
29403010
132,938 Japanese ancestry individuals
Neuroticism
29292387
168,105 European ancestry individuals

Non−albumin protein levels
29403010

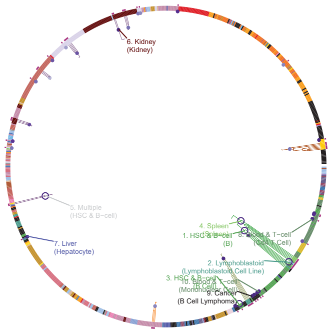


98,538 Japanese ancestry individuals

Parental longevity (mother's age at death)
29227965

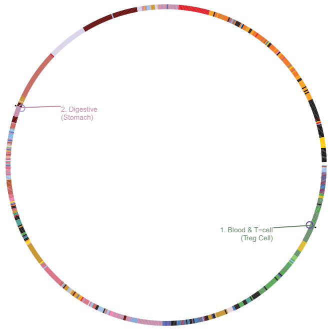


246,941 British ancestry individuals

Remission after SSRI treatment in MDD or neuroticism
29559929

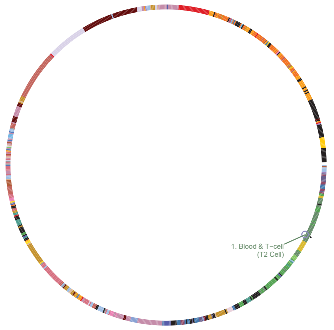


865 individuals with SSRI response data, 170,911 individuals with personality trait data

Lymphocyte counts
29403010

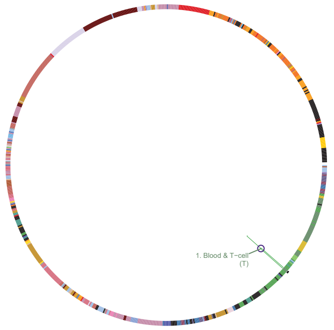


62,076 Japanese ancestry individuals

Measles
28928442

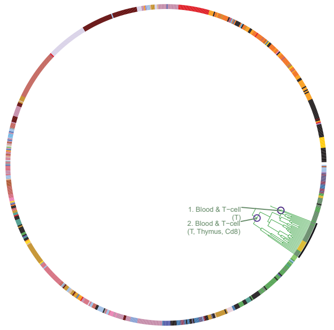


38,219 European ancestry cases, 47,279 European ancestry controls

Morning vs. evening chronotype
26835600

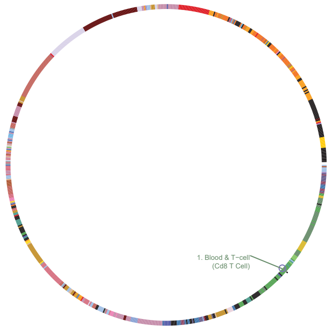


38,937 European ancestry morning chronotype individuals, 50,346 European ancestry evening chronotype individuals

Hair morphology
19896111

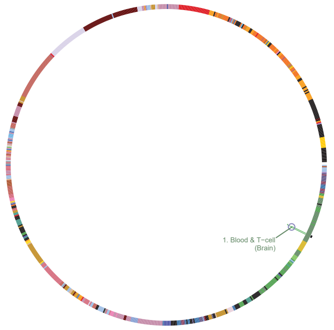


4,845 European ancestry individuals from 2,892 families, 16,140 European ancestry individuals

Age at first sexual intercourse
27089180

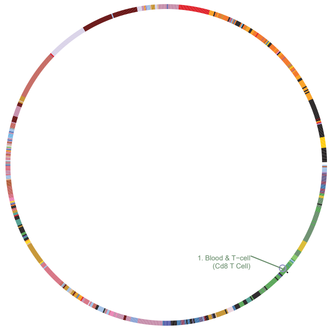


59,357 British ancestry males, 66,310 British ancestry females

Alanine transaminase levels
29403010

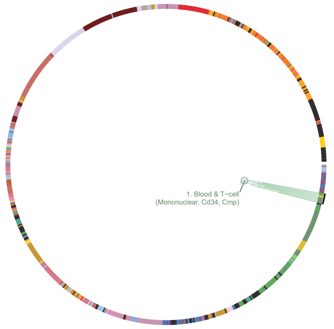


134,182 Japanese ancestry individuals

Chronic inflammatory diseases (ankylosing spondylitis, Crohn's disease, psoriasis, primary sclerosing cholangitis, ulcerative colitis) (pleiotropy)
26974007

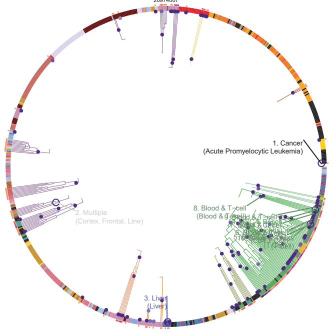


8,726 European ancestry ankylosing spondylitis cases, 19,085 European ancestry Crohn...s disease cases, 6,530 European ancestry psoriasis cases, 3,408 European ancestry primary sclerosing cholangitis cases, 14,513 European ancestry ulcerative colitis cases, 34,213 European ancestry controls

Menopause (age at onset)
26414677

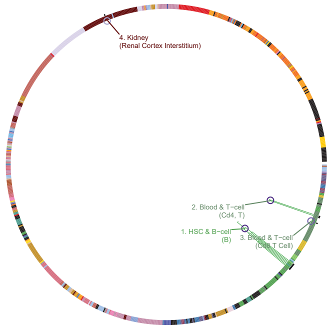


up to 69,626 European ancestry women

Eosinophil percentage of granulocytes
27863252

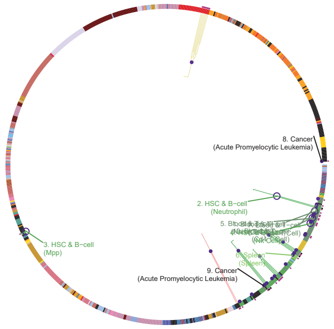


170,536 European ancestry individuals

Neutrophil percentage of granulocytes
27863252

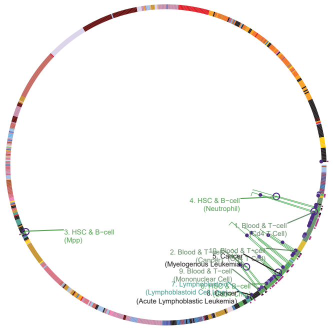


170,672 European ancestry individuals

Eosinophil counts
27863252

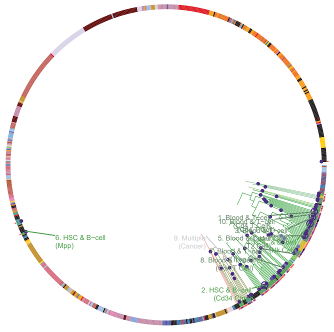


172,275 European ancestry individuals

Sum eosinophil basophil counts
27863252

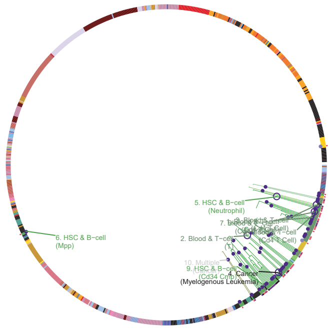


171,771 European ancestry individuals

Lymphocyte counts
27863252

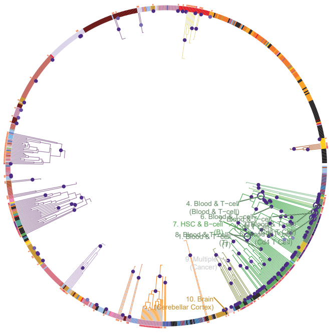


171,643 European ancestry individuals

Eczema
30595370

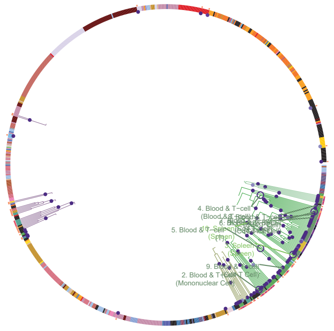


approximately 459,000 European ancestry individuals

Eosinophil counts
30595370

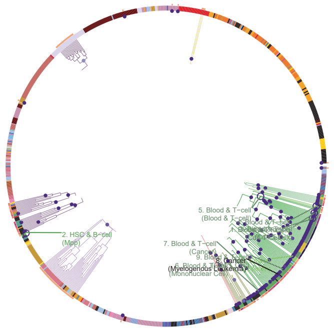


approximately 440,000 European ancestry individuals

Eosinophil percentage of white cells
27863252

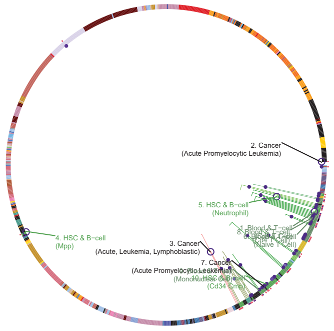


172,378 European ancestry individuals

Allergic disease (asthma, hay fever or eczema)
29083406

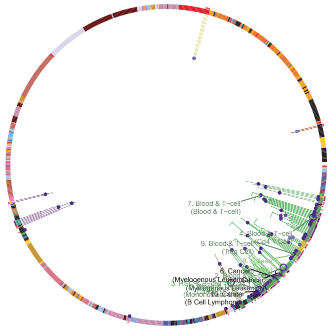


180,129 European ancestry cases, 180,709 European ancestry controls

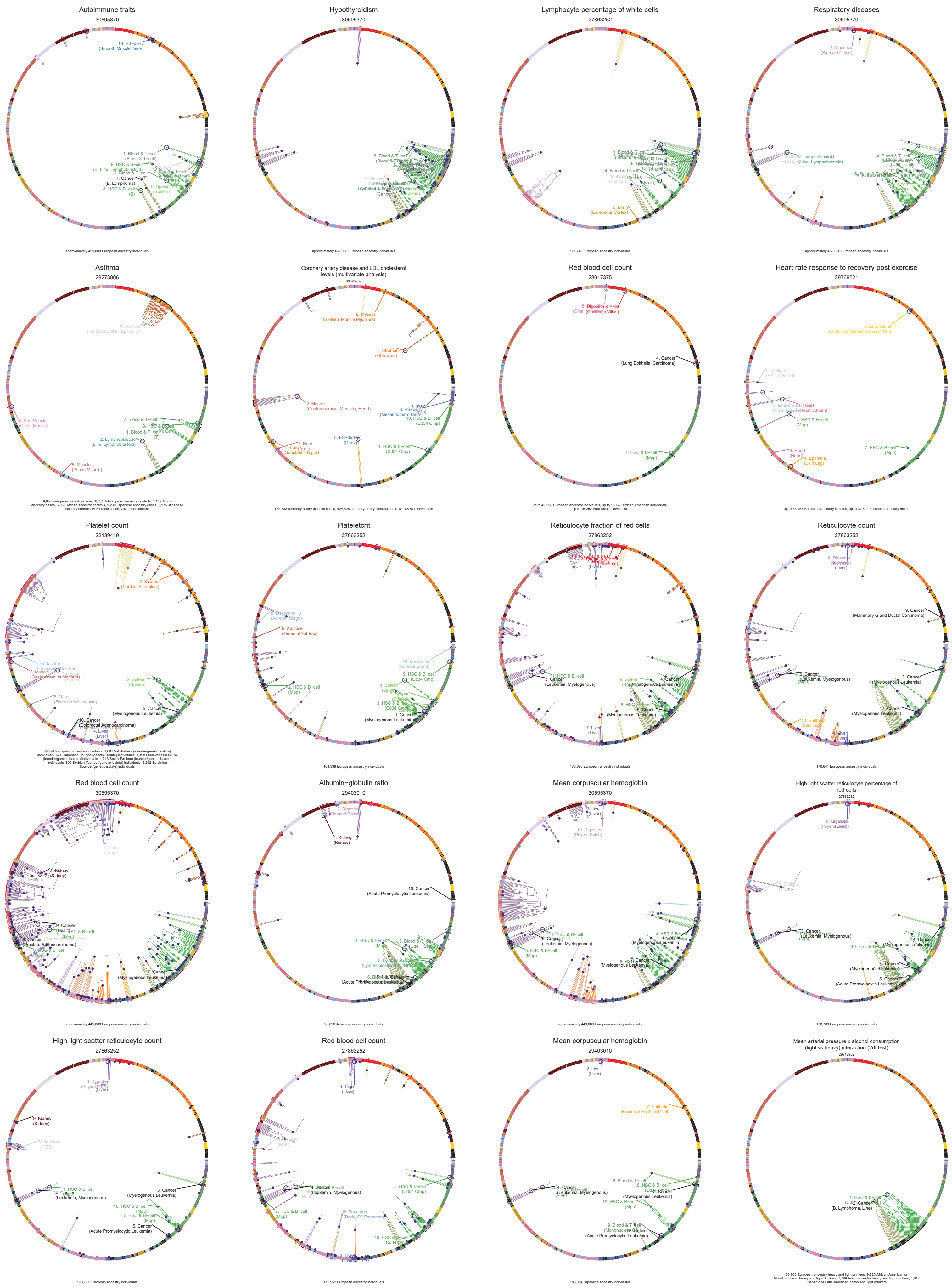

Autoimmune traits
30595370
approximately 459,000 European ancestry individuals
Hypothyroidism
30595370
approximately 459,000 European ancestry individuals
Lymphocyte percentage of white cells
27863252
171,748 European ancestry individuals
Respiratory diseases
30595370
approximately 459,000 European ancestry individuals
Asthma
29273806
19,954 European ancestry cases, 107,715 European ancestry controls, 2,149 African ancestry cases, 6,055 African ancestry controls, 1,239 Japanese ancestry cases, 3,976 Japanese ancestry controls, 606 Latino cases, 792 Latino controls
Coronary artery disease and LDL cholesterol levels (multivariate analysis)
30525989
122,733 coronary artery disease cases, 424,528 coronary artery disease controls, 188,577 individuals
Red blood cell count
28017375
up to 40,258 European ancestry individuals, up to 16,128 African American individuals, up to 15,252 East Asian individuals
Heart rate response to recovery post exercise
29769521
up to 35,455 European ancestry females, up to 31,802 European ancestry males
Platelet count
22139419
38,891 European ancestry individuals, 1,661 Val Borbera (founder/genetic isolate) individuals, 521 Carlantino (founder/genetic isolate) individuals, 1,198 Friuli Venezia Giulia (founder/genetic isolate) individuals, 1,213 South Tyrolean (founder/genetic isolate) individuals, 890 Sorbian (founder/genetic isolate) individuals, 4,292 Sardinian (founder/genetic isolate) individuals
Plateletcrit
27863252
164,339 European ancestry individuals
Reticulocyte fraction of red cells
27863252
170,690 European ancestry individuals
Reticulocyte count
27863252
170,641 European ancestry individuals
Red blood cell count
30595370
approximately 445,000 European ancestry individuals
Albumin−globulin ratio
29403010
98,626 Japanese ancestry individuals
Mean corpuscular hemoglobin
30595370
approximately 443,000 European ancestry individuals
High light scatter reticulocyte percentage of red cells
27863252
170,763 European ancestry individuals
High light scatter reticulocyte count
27863252
170,761 European ancestry individuals
Red blood cell count
27863252
172,952 European ancestry individuals
Mean corpuscular hemoglobin
29403010
108,054 Japanese ancestry individuals
Mean arterial pressure x alcohol consumption (light vs heavy) interaction (2df test)
29912962
56,705 European ancestry heavy and light drinkers, 9,725 African American or Afro−Caribbean heavy and light drinkers, 1,789 Asian ancestry heavy and light drinkers, 4,812 Hispanic or Latin American heavy and light drinkers

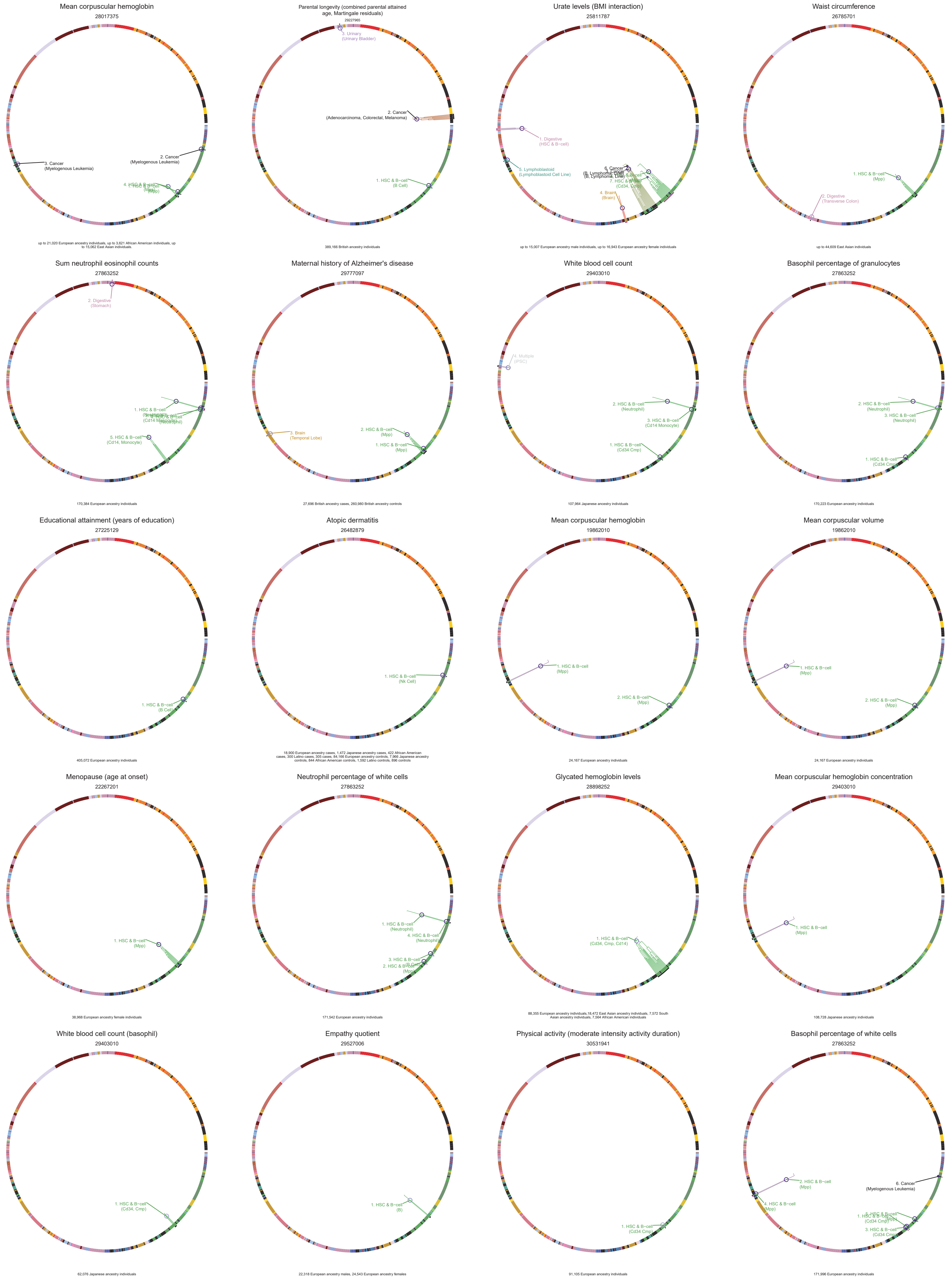

Mean corpuscular hemoglobin
28017375
up to 21,020 European ancestry individuals, up to 3,621 African American individuals, up to 15,062 East Asian individuals.
Parental longevity (combined parental attained age, Martingale residuals)
29227965
389,166 British ancestry individuals
Urate levels (BMI interaction)
25811787
up to 15,007 European ancestry male individuals, up to 16,943 European ancestry female individuals
Waist circumference
26785701
up to 44,609 East Asian individuals
Sum neutrophil eosinophil counts
27863252
170,384 European ancestry individuals
Maternal history of Alzheimer's disease
29777097
27,696 British ancestry cases, 260,980 British ancestry controls
White blood cell count
29403010
107,964 Japanese ancestry individuals
Basophil percentage of granulocytes
27863252
170,223 European ancestry individuals
Educational attainment (years of education)
27225129
405,072 European ancestry individuals
Atopic dermatitis
26482879
18,900 European ancestry cases, 1,472 Japanese ancestry cases, 422 African American cases, 300 Latino cases, 305 cases, 84,166 European ancestry controls, 7,966 Japanese ancestry controls, 844 African American controls, 1,592 Latino controls, 896 controls
Mean corpuscular hemoglobin
19862010
24,167 European ancestry individuals
Mean corpuscular volume
19862010
24,167 European ancestry individuals
Menopause (age at onset)
22267201
38,968 European ancestry female individuals
Neutrophil percentage of white cells
27863252
171,542 European ancestry individuals
Glycated hemoglobin levels
28898252
88,355 European ancestry individuals,18,472 East Asian ancestry individuals, 7,572 South Asian ancestry individuals, 7,564 African American individuals
Mean corpuscular hemoglobin concentration
29403010
108,728 Japanese ancestry individuals
White blood cell count (basophil)
29403010
62,076 Japanese ancestry individuals
Empathy quotient
29527006
22,318 European ancestry males, 24,543 European ancestry females
Physical activity (moderate intensity activity duration)
30531941
91,105 European ancestry individuals
Basophil percentage of white cells
27863252
171,996 European ancestry individuals

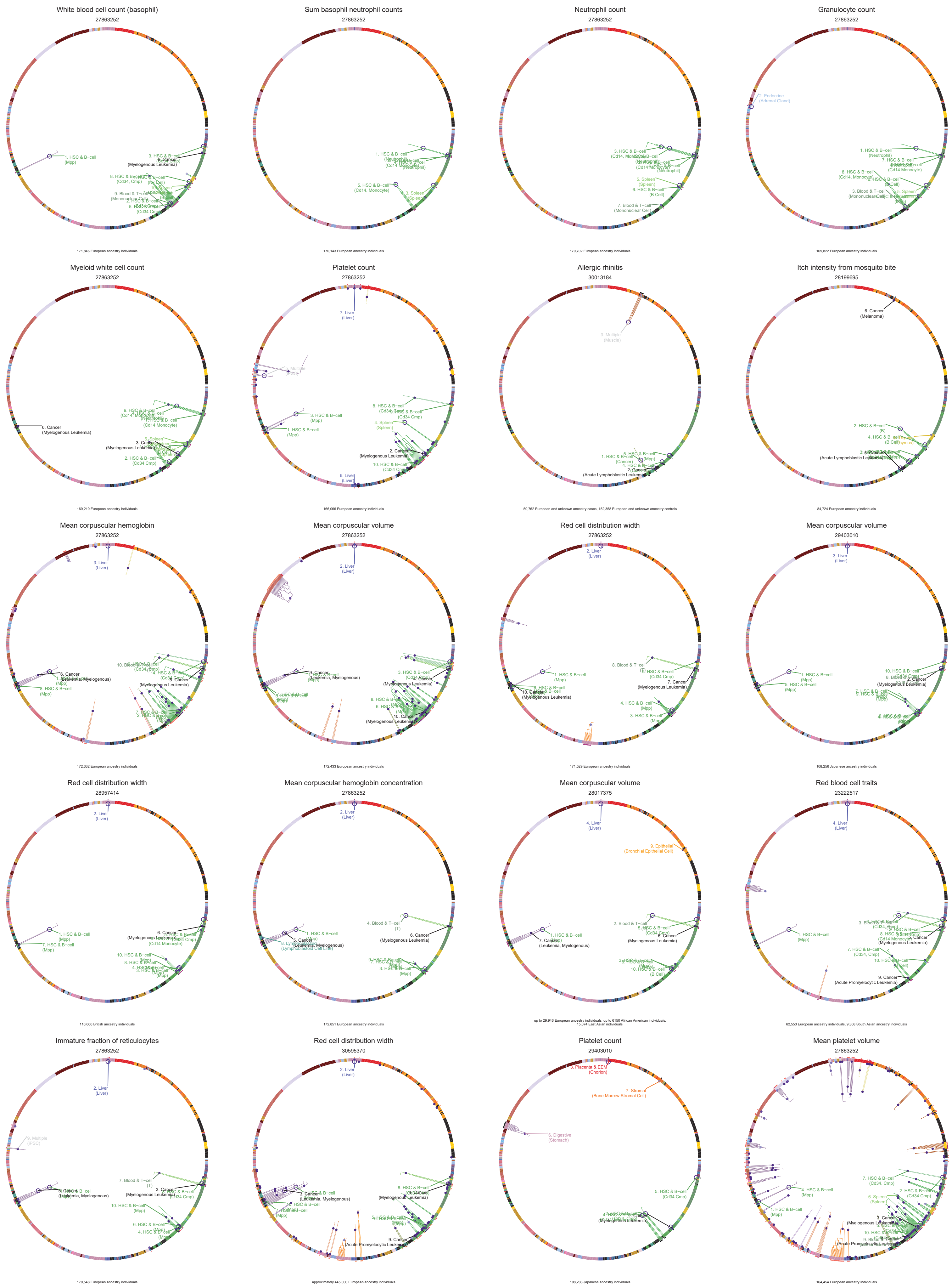
White blood cell count (basophil)
27863252
171,846 European ancestry individuals
Sum basophil neutrophil counts
27863252
170,143 European ancestry individuals
Neutrophil count
27863252
170,702 European ancestry individuals
Granulocyte count
27863252
169,822 European ancestry individuals
Myeloid white cell count
27863252
169,219 European ancestry individuals
Platelet count
27863252
166,066 European ancestry individuals
Allergic rhinitis
30013184
59,762 European and unknown ancestry cases, 152,358 European and unknown ancestry controls
Itch intensity from mosquito bite
28199695
84,724 European ancestry individuals
Mean corpuscular hemoglobin
27863252
172,332 European ancestry individuals
Mean corpuscular volume
27863252
172,433 European ancestry individuals
Red cell distribution width
27863252
171,529 European ancestry individuals
Mean corpuscular volume
29403010
108,256 Japanese ancestry individuals
Red cell distribution width
28957414
116,666 British ancestry individuals
Mean corpuscular hemoglobin concentration
27863252
172,851 European ancestry individuals
Mean corpuscular volume
28017375
up to 29,946 European ancestry individuals, up to 6150 African American individuals, 15,074 East Asian individuals
Red blood cell traits
23222517
62,553 European ancestry individuals, 9,308 South Asian ancestry individuals
Immature fraction of reticulocytes
27863252
170,548 European ancestry individuals
Red cell distribution width
30595370
approximately 445,000 European ancestry individuals
Platelet count
29403010
108,208 Japanese ancestry individuals
Mean platelet volume
27863252
164,454 European ancestry individuals

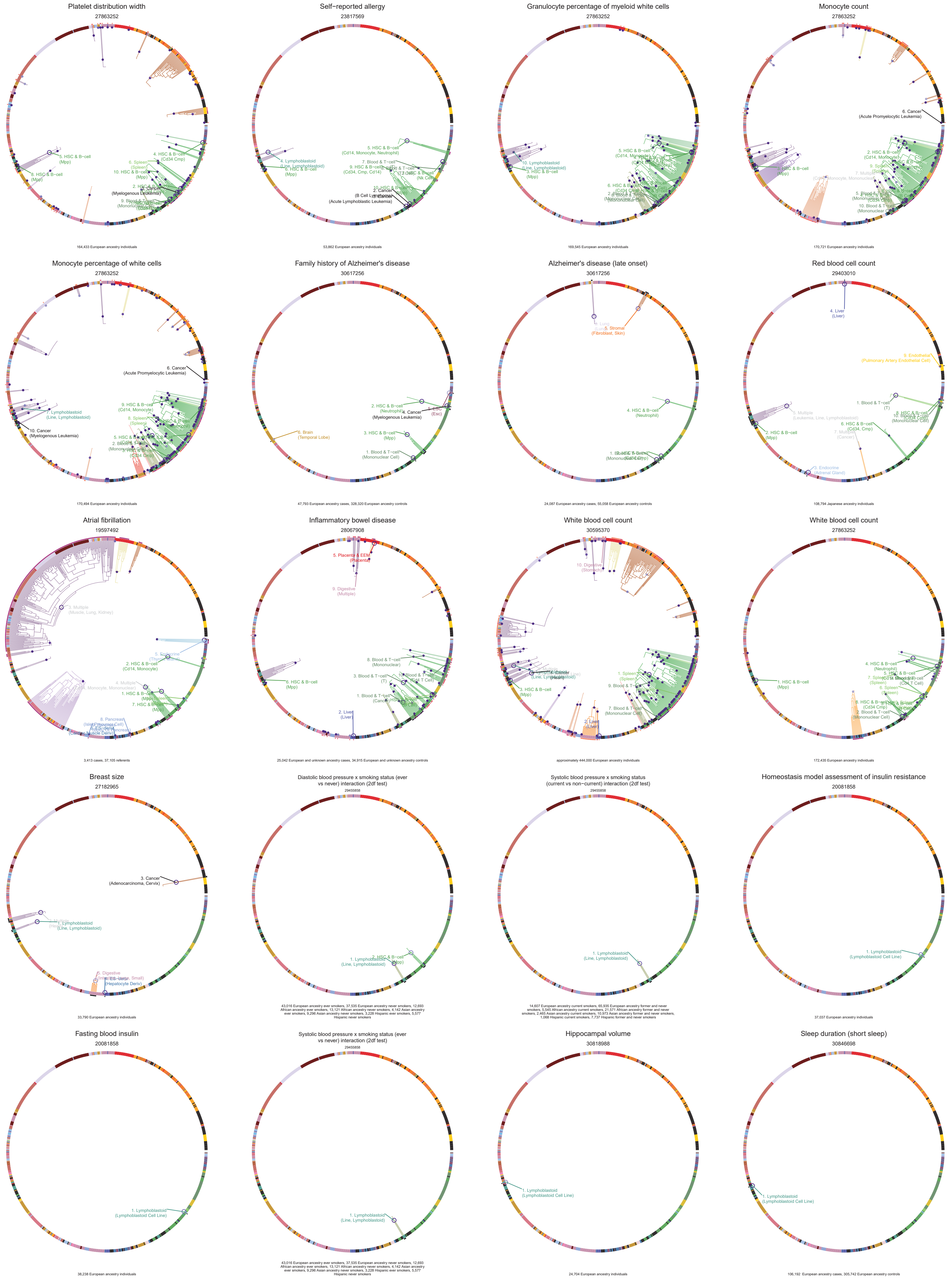

Platelet distribution width
27863252
5. HSC & B−cell
(Mpp)
8. HSC & B−cell
(Mpp)
6. Spleen
(Spleen)
4. HSC & B−cell
(Cd34 Cmp)
10. HSC & B−cell
(Mpp)
(Myelogenous Leukemia)
164,433 European ancestry individuals
Self−reported allergy
23817569
4. Lymphoblastoid
(Line, Lymphoblastoid)
8. HSC & B−cell
(Mpp)
5. HSC & B−cell
(Cd14, Monocyte, Neutrophil)
7. Blood & T−cell
9. HSC & B−cell
(Cd34, Cmp, Cd14)
2. Cancer
(B Cell Lymphoma)
(Acute Lymphoblastic Leukemia)
53,862 European ancestry individuals
Granulocyte percentage of myeloid white cells
27863252
10. Lymphoblastoid
(Line, Lymphoblastoid)
3. HSC & B−cell
(Mpp)
5. HSC & B−cell
169,545 European ancestry individuals
Monocyte count
27863252
6. Cancer
(Acute Promyelocytic Leukemia)
2. HSC & B−cell
(Cd14, Monocyte)
(Mpp)
10. Blood & T−cell
(Mononuclear Cell)
170,721 European ancestry individuals
Monocyte percentage of white cells
27863252
6. Cancer
(Acute Promyelocytic Leukemia)
7. Lymphoblastoid
(Line, Lymphoblastoid)
10. Cancer
(Myelogenous Leukemia)
9. HSC & B−cell
(Cd14, Monocyte)
8. Spleen
(Spleen)
170,494 European ancestry individuals
Family history of Alzheimer's disease
30617256
6. Brain
(Temporal Lobe)
2. HSC & B−cell
(Neutrophil)
4. Cancer
(Myelogenous Leukemia)
(Esc)
3. HSC & B−cell
(Mpp)
1. Blood & T−cell
(Mononuclear Cell)
47,793 European ancestry cases, 328,320 European ancestry controls
Alzheimer's disease (late onset)
30617256
5. Stromal
(Fibroblast, Skin)
4. HSC & B−cell
(Neutrophil)
24,087 European ancestry cases, 55,058 European ancestry controls
Red blood cell count
29403010
4. Liver
(Liver)
9. Endothelial
(Pulmonary Artery Endothelial Cell)
1. Blood & T−cell
(T)
8. Multiple
(Leukemia, Line, Lymphoblastoid)
2. HSC & B−cell
(Mpp)
6. HSC & B−cell
(Cd34, Cmp)
7. Multiple
(Cancer)
3. Endocrine
(Adrenal Gland)
108,794 Japanese ancestry individuals
Atrial fibrillation
19597492
3. Multiple
(Muscle, Lung, Kidney)
2. HSC & B−cell
(Cd14, Monocyte)
4. Multiple
1. HSC & B−cell
7. HSC & B−cell
8. Pancreas
3,413 cases, 37,105 referents
Inflammatory bowel disease
28067908
5. Placenta & EEM
(Placenta)
9. Digestive
(Multiple)
6. HSC & B−cell
(Mpp)
8. Blood & T−cell
(Mononuclear)
3. Blood & T−cell
(T)
1. Blood & T−cell
2. Liver
(Liver)
25,042 European and unknown ancestry cases, 34,915 European and unknown ancestry controls
White blood cell count
30595370
10. Digestive
(Stomach)
3. HSC & B−cell
(Mpp)
1. Spleen
(Spleen)
9. Blood & T−cell
7. Blood & T−cell
(Mononuclear Cell)
2. Liver
approximately 444,000 European ancestry individuals
White blood cell count
27863252
1. HSC & B−cell
(Mpp)
4. HSC & B−cell
(Neutrophil)
7. Spleen
(Spleen)
6. Spleen
(Spleen)
2. Blood & T−cell
(Mononuclear Cell)
172,435 European ancestry individuals
Breast size
27182965
3. Cancer
(Adenocarcinoma, Cervix)
1. Lymphoblastoid
(Line, Lymphoblastoid)
5. Digestive
(Hepatocyte Deriv)
33,790 European ancestry individuals
Diastolic blood pressure x smoking status (ever vs never) interaction (2df test)
29455858
1. Lymphoblastoid
(Line, Lymphoblastoid)
2. HSC & B−cell
43,016 European ancestry ever smokers, 37,535 European ancestry never smokers, 12,693 African ancestry ever smokers, 13,121 African ancestry never smokers, 4,142 Asian ancestry ever smokers, 9,296 Asian ancestry never smokers, 3,228 Hispanic ever smokers, 5,577 Hispanic never smokers
Systolic blood pressure x smoking status (current vs non−current) interaction (2df test)
29455858
1. Lymphoblastoid
(Line, Lymphoblastoid)
14,607 European ancestry current smokers, 65,935 European ancestry former and never smokers, 5,545 African ancestry current smokers, 21,571 African ancestry former and never smokers, 2,465 Asian ancestry current smokers, 10,973 Asian ancestry former and never smokers, 1,068 Hispanic current smokers, 7,737 Hispanic former and never smokers
Homeostasis model assessment of insulin resistance
20081858
1. Lymphoblastoid
(Lymphoblastoid Cell Line)
37,037 European ancestry individuals
Fasting blood insulin
20081858
1. Lymphoblastoid
(Lymphoblastoid Cell Line)
38,238 European ancestry individuals
Systolic blood pressure x smoking status (ever vs never) interaction (2df test)
29455858
1. Lymphoblastoid
(Line, Lymphoblastoid)
43,016 European ancestry ever smokers, 37,535 European ancestry never smokers, 12,693 African ancestry ever smokers, 13,121 African ancestry never smokers, 4,142 Asian ancestry ever smokers, 9,296 Asian ancestry never smokers, 3,228 Hispanic ever smokers, 5,577 Hispanic never smokers
Hippocampal volume
30818988
1. Lymphoblastoid
(Lymphoblastoid Cell Line)
24,704 European ancestry individuals
Sleep duration (short sleep)
30846698
1. Lymphoblastoid
(Lymphoblastoid Cell Line)
106,192 European ancestry cases, 305,742 European ancestry controls

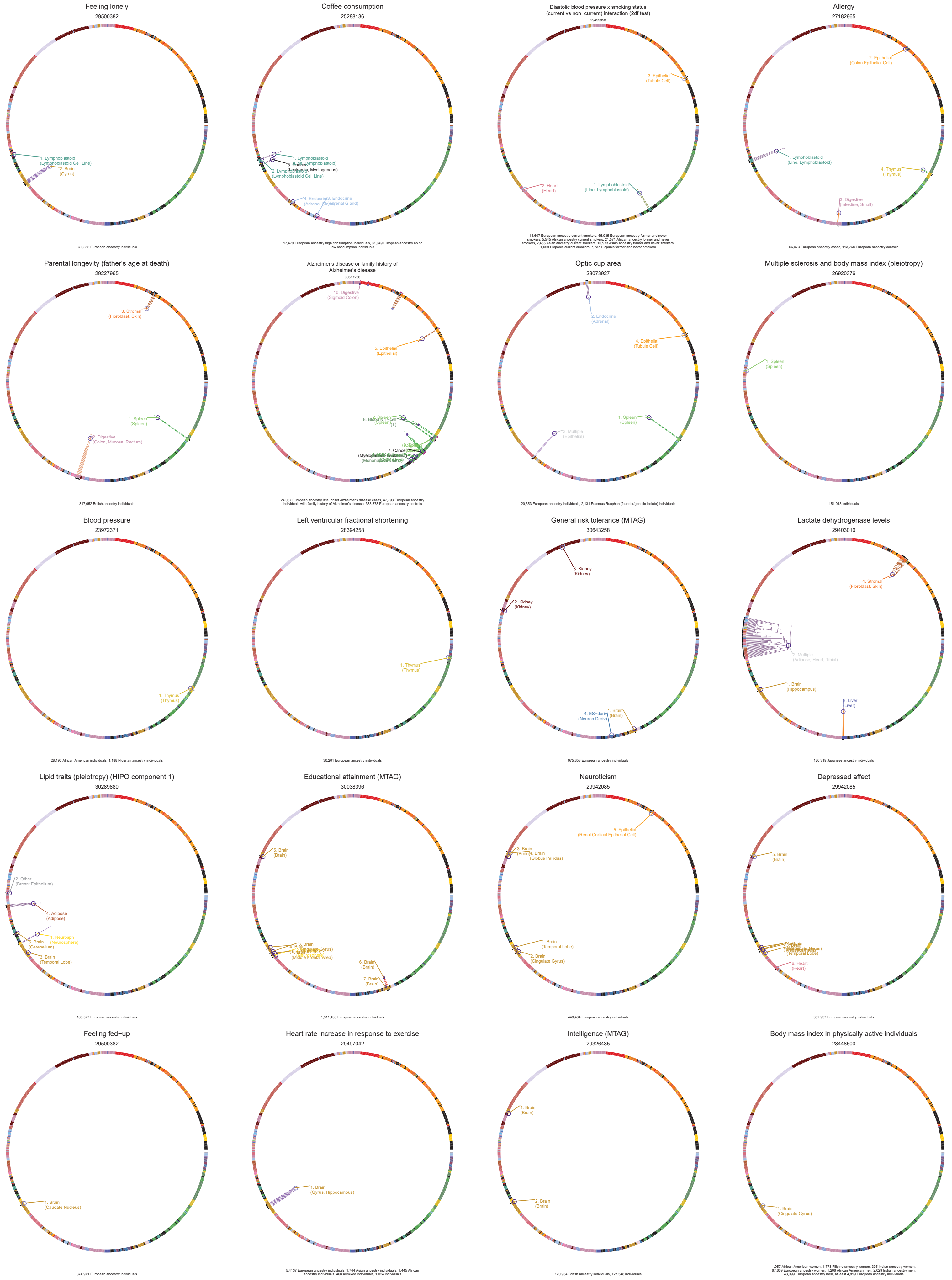

Feeling lonely
29500382
1. Lymphoblastoid (Lymphoblastoid Cell Line)
2. Brain (Gyrus)
376,352 European ancestry individuals
Coffee consumption
25288136
1. Lymphoblastoid (Line, Lymphoblastoid)
5. Cancer (Leukemia, Myelogenous)
2. Lymphoblastoid (Lymphoblastoid Cell Line)
17,479 European ancestry high consumption individuals, 31,049 European ancestry no or low consumption individuals
Diastolic blood pressure x smoking status (current vs non−current) interaction (2df test)
29455858
3. Epithelial (Tubule Cell)
2. Heart (Heart)
1. Lymphoblastoid (Line, Lymphoblastoid)
14,607 European ancestry current smokers, 65,935 European ancestry former and never smokers, 5,545 African ancestry current smokers, 21,571 African ancestry former and never smokers, 2,465 Asian ancestry current smokers, 10,973 Asian ancestry former and never smokers, 1,068 Hispanic current smokers, 7,737 Hispanic former and never smokers
Allergy
27182965
2. Epithelial (Colon Epithelial Cell)
1. Lymphoblastoid (Line, Lymphoblastoid)
4. Thymus (Thymus)
3. Digestive (Intestine, Small)
66,973 European ancestry cases, 113,768 European ancestry controls
Parental longevity (father's age at death)
29227965
3. Stromal (Fibroblast, Skin)
1. Spleen (Spleen)
2. Digestive (Colon, Mucosa, Rectum)
317,652 British ancestry individuals
Alzheimer's disease or family history of Alzheimer's disease
30617256
10. Digestive (Sigmoid Colon)
5. Epithelial (Epithelial)
24,087 European ancestry late−onset Alzheimer's disease cases, 47,793 European ancestry individuals with family history of Alzheimer's disease, 383,378 European ancestry controls
Optic cup area
28073927
2. Endocrine (Adrenal)
4. Epithelial (Tubule Cell)
1. Spleen (Spleen)
3. Multiple (Epithelial)
20,353 European ancestry individuals, 2,131 Erasmus Rucphen (founder/genetic isolate) individuals
Multiple sclerosis and body mass index (pleiotropy)
26920376
1. Spleen (Spleen)
151,013 individuals
Blood pressure
23972371
1. Thymus (Thymus)
28,190 African American individuals, 1,188 Nigerian ancestry individuals
Left ventricular fractional shortening
28394258
1. Thymus (Thymus)
30,201 European ancestry individuals
General risk tolerance (MTAG)
30643258
3. Kidney (Kidney)
2. Kidney (Kidney)
1. Brain (Brain)
4. ES−deriv (Neuron Deriv)
975,353 European ancestry individuals
Lactate dehydrogenase levels
29403010
4. Stromal (Fibroblast, Skin)
2. Multiple (Adipose, Heart, Tibial)
1. Brain (Hippocampus)
3. Liver (Liver)
126,319 Japanese ancestry individuals
Lipid traits (pleiotropy) (HIPO component 1)
30289880
2. Other (Breast Epithelium)
4. Adipose (Adipose)
1. Neurosph (Neurosphere)
5. Brain (Cerebellum)
3. Brain (Temporal Lobe)
188,577 European ancestry individuals
Educational attainment (MTAG)
30038396
5. Brain (Brain)
6. Brain (Brain)
7. Brain (Brain)
1,311,438 European ancestry individuals
Neuroticism
29942085
5. Epithelial (Renal Cortical Epithelial Cell)
3. Brain (Brain)
4. Brain (Globus Pallidus)
1. Brain (Temporal Lobe)
2. Brain (Cingulate Gyrus)
449,484 European ancestry individuals
Depressed affect
29942085
5. Brain (Brain)
6. Heart (Heart)
357,957 European ancestry individuals
Feeling fed−up
29500382
1. Brain (Caudate Nucleus)
374,971 European ancestry individuals
Heart rate increase in response to exercise
29497042
1. Brain (Gyrus, Hippocampus)
5,4137 European ancestry individuals, 1,744 Asian ancestry individuals, 1,445 African ancestry individuals, 468 admixed individuals, 1,024 individuals
Intelligence (MTAG)
29326435
1. Brain (Brain)
2. Brain (Brain)
120,934 British ancestry individuals, 127,548 individuals
Body mass index in physically active individuals
28448500
1. Brain (Cingulate Gyrus)
1,957 African American women, 1,773 Filipino ancestry women, 305 Indian ancestry women, 67,609 European ancestry women, 1,206 African American men, 2,029 Indian ancestry men, 43,399 European ancestry men, at least 4,819 European ancestry individuals

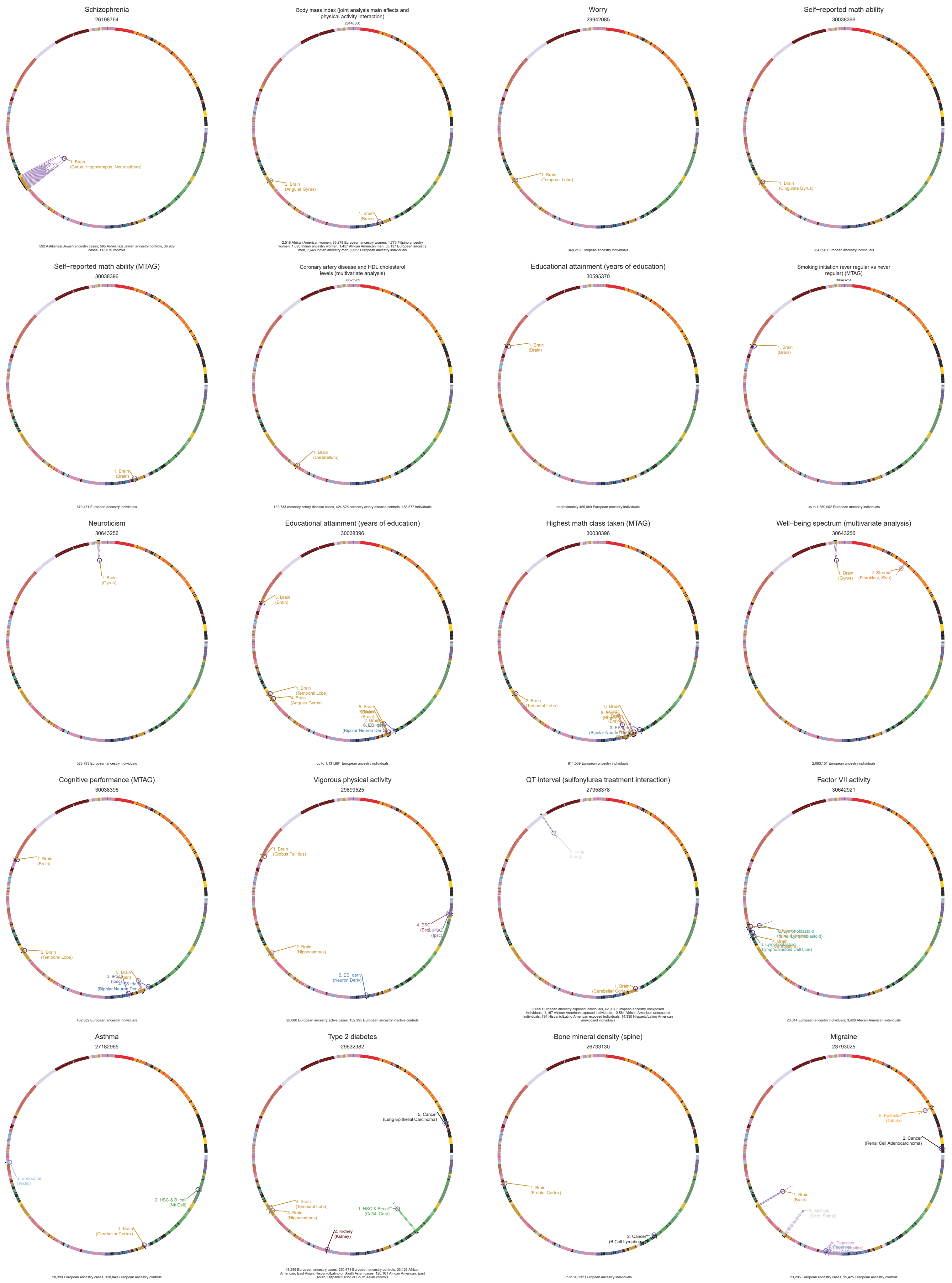

Schizophrenia
26198764
1. Brain
(Gyrus, Hippocampus, Neurosphere)
592 Ashkenazi Jewish ancestry cases, 505 Ashkenazi Jewish ancestry controls, 36,989 cases, 113,075 controls
Body mass index (joint analysis main effects and physical activity interaction)
28448500
2. Brain
(Angular Gyrus)
1. Brain
(Brain)
2,518 African American women, 86,278 European ancestry women, 1,773 Filipino ancestry women, 1,030 Indian ancestry women, 1,457 African American men, 55,137 European ancestry men, 7,648 Indian ancestry men, 5,527 European ancestry individuals
Worry
29942085
1. Brain
(Temporal Lobe)
348,219 European ancestry individuals
Self−reported math ability
30038396
1. Brain
(Cingulate Gyrus)
564,698 European ancestry individuals
Self−reported math ability (MTAG)
30038396
1. Brain
(Brain)
670,471 European ancestry individuals
Coronary artery disease and HDL cholesterol levels (multivariate analysis)
30525989
1. Brain
(Cerebellum)
122,733 coronary artery disease cases, 424,528 coronary artery disease controls, 188,577 individuals
Educational attainment (years of education)
30595370
1. Brain
(Brain)
approximately 455,000 European ancestry individuals
Smoking initiation (ever regular vs never regular) (MTAG)
30643251
1. Brain
(Brain)
up to 1,359,002 European ancestry individuals
Neuroticism
30643256
1. Brain
(Gyrus)
523,783 European ancestry individuals
Educational attainment (years of education)
30038396
3. Brain
(Brain)
1. Brain
(Temporal Lobe)
4. Brain
(Angular Gyrus)
5. Brain
(Bipolar Neuron Deriv)
up to 1,131,881 European ancestry individuals
Highest math class taken (MTAG)
30038396
2. Brain
(Temporal Lobe)
6. Brain
(Bipolar Neuron Deriv)
811,539 European ancestry individuals
Well−being spectrum (multivariate analysis)
30643256
1. Brain
(Gyrus)
2. Stromal
(Fibroblast, Skin)
2,083,151 European ancestry individuals
Cognitive performance (MTAG)
30038396
1. Brain
(Brain)
2. Brain
(Temporal Lobe)
5. Brain
(Bipolar Neuron Deriv)
402,382 European ancestry individuals
Vigorous physical activity
29899525
1. Brain
(Globus Pallidus)
4. ESC
3. iPSC
(Ipsc)
2. Brain
(Hippocampus)
5. ES−deriv
(Neuron Deriv)
98,060 European ancestry active cases, 162,995 European ancestry inactive controls
QT interval (sulfonylurea treatment interaction)
27958378
2. Lung
(Lung)
1. Brain
(Cerebellar Cortex)
2,095 European ancestry exposed individuals, 42,907 European ancestry unexposed individuals, 1,167 African American exposed individuals, 10,564 African American unexposed individuals, 794 Hispanic/Latino American exposed individuals, 14,330 Hispanic/Latino American unexposed individuals
Factor VII activity
30642921
4. Brain
3. Lymphoblastoid
(Lymphoblastoid Cell Line)
20,014 European ancestry individuals, 3,420 African American individuals
Asthma
27182965
3. Endocrine
(Testis)
2. HSC & B−cell
(Nk Cell)
1. Brain
(Cerebellar Cortex)
28,399 European ancestry cases, 128,843 European ancestry controls
Type 2 diabetes
29632382
5. Cancer
(Lung Epithelial Carcinoma)
4. Brain
(Temporal Lobe)
3. Brain
(Hippocampus)
1. HSC & B−cell
(Cd34, Cmp)
2. Kidney
(Kidney)
48,286 European ancestry cases, 250,671 European ancestry controls, 33,126 African American, East Asian, Hispanic/Latino or South Asian cases, 120,161 African American, East Asian, Hispanic/Latino or South Asian controls
Bone mineral density (spine)
26733130
1. Brain
(Frontal Cortex)
2. Cancer
up to 20,132 European ancestry individuals
Migraine
23793025
5. Epithelial
(Tubule)
2. Cancer
(Renal Cell Adenocarcinoma)
1. Brain
(Brain)
4. Digestive
23,285 European ancestry cases, 95,425 European ancestry controls

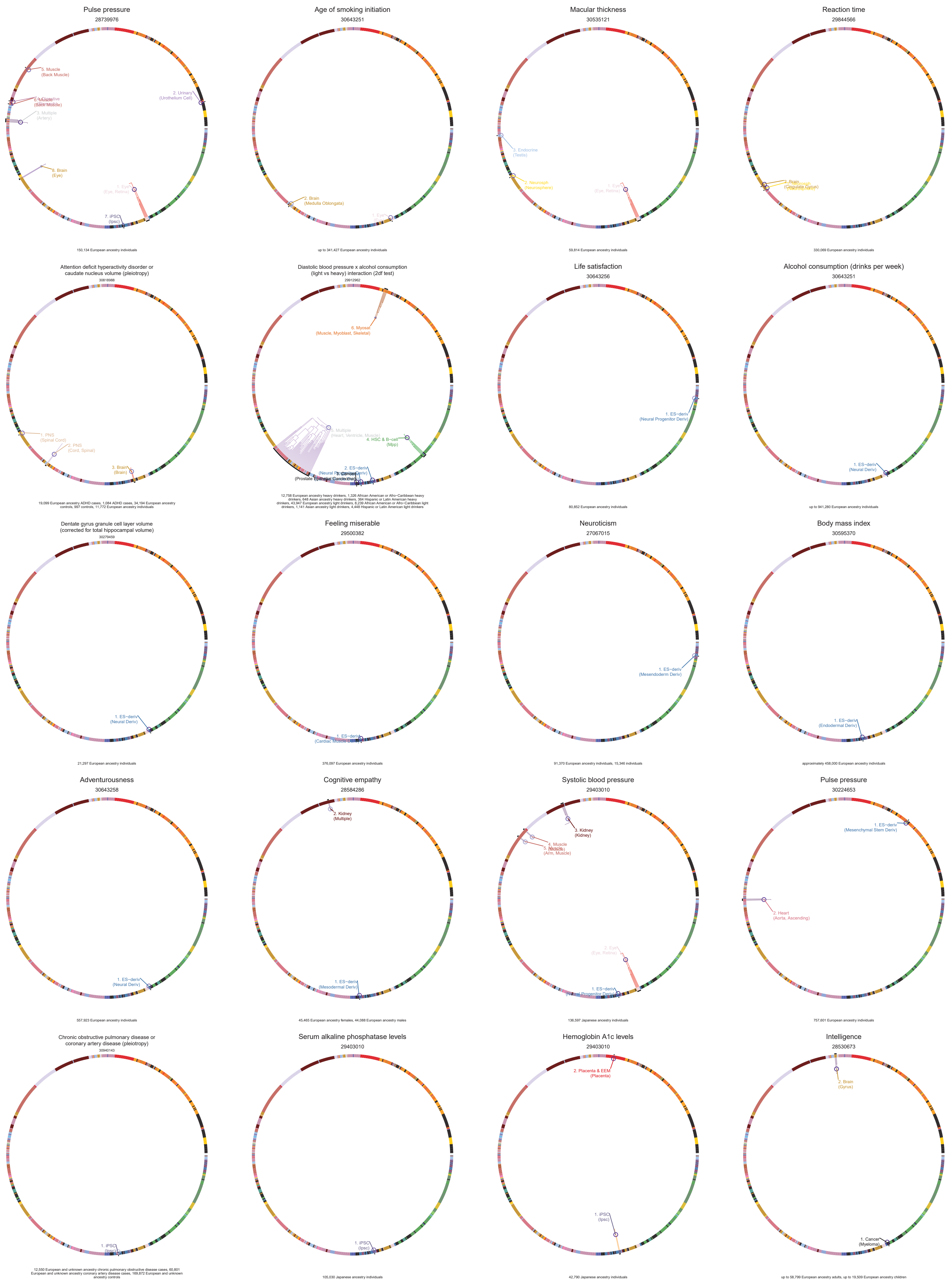

Pulse pressure
28739976
5. Muscle
(Back Muscle)
2. Urinary
(Urothelium Cell)
3. Multiple
(Artery)
8. Brain
(Eye)
1. Eye
(Eye, Retina)
7. iPSC
(Ipsc)
150,134 European ancestry individuals
Age of smoking initiation
30643251
2. Brain
(Medulla Oblongata)
up to 341,427 European ancestry individuals
Macular thickness
30535121
3. Endocrine
(Testis)
2. Neurosph
(Neurosphere)
1. Eye
(Eye, Retina)
59,814 European ancestry individuals
Reaction time
29844566
(Cingulate Gyrus)
330,069 European ancestry individuals
Attention deficit hyperactivity disorder or
caudate nucleus volume (pleiotropy)
30818988
1. PNS
(Spinal Cord)
2. PNS
(Cord, Spinal)
3. Brain
(Brain)
19,099 European ancestry ADHD cases, 1,084 ADHD cases, 34,194 European ancestry controls, 997 controls, 11,772 European ancestry individuals
Diastolic blood pressure x alcohol consumption
(light vs heavy) interaction (2df test)
29912962
6. Myosat
(Muscle, Myoblast, Skeletal)
5. Multiple
(Heart, Ventricle, Muscle)
4. HSC & B-cell
(Mpp)
2. ES-deriv
12,758 European ancestry heavy drinkers, 1,326 African American or Afro-Caribbean heavy drinkers, 648 Asian ancestry heavy drinkers, 364 Hispanic or Latin American heavy drinkers, 43,947 European ancestry light drinkers, 8,239 African American or Afro-Caribbean light drinkers, 1,141 Asian ancestry light drinkers, 4,448 Hispanic or Latin American light drinkers
Life satisfaction
30643256
1. ES-deriv
(Neural Progenitor Deriv)
80,852 European ancestry individuals
Alcohol consumption (drinks per week)
30643251
1. ES-deriv
(Neural Deriv)
up to 941,280 European ancestry individuals
Dentate gyrus granule cell layer volume
(corrected for total hippocampal volume)
30279459
1. ES-deriv
(Neural Deriv)
21,297 European ancestry individuals
Feeling miserable
29500382
1. ES-deriv
376,097 European ancestry individuals
Neuroticism
27067015
1. ES-deriv
(Mesendoderm Deriv)
91,370 European ancestry individuals, 15,346 individuals
Body mass index
30595370
1. ES-deriv
(Endodermal Deriv)
approximately 458,000 European ancestry individuals
Adventurousness
30643258
1. ES-deriv
(Neural Deriv)
557,923 European ancestry individuals
Cognitive empathy
28584286
2. Kidney
(Multiple)
1. ES-deriv
(Mesodermal Deriv)
45,465 European ancestry females, 44,068 European ancestry males
Systolic blood pressure
29403010
3. Kidney
(Kidney)
4. Muscle
(Arm, Muscle)
2. Eye
(Eye, Retina)
1. ES-deriv
136,597 Japanese ancestry individuals
Pulse pressure
30224653
1. ES-deriv
(Mesenchymal Stem Deriv)
2. Heart
(Aorta, Ascending)
757,601 European ancestry individuals
Chronic obstructive pulmonary disease or
coronary artery disease (pleiotropy)
30940143
1. iPSC
(Ipsc)
12,550 European and unknown ancestry chronic pulmonary obstructive disease cases, 60,801 European and unknown ancestry coronary artery disease cases, 169,872 European and unknown ancestry controls
Serum alkaline phosphatase levels
29403010
1. iPSC
(Ipsc)
105,030 Japanese ancestry individuals
Hemoglobin A1c levels
29403010
2. Placenta & EEM
(Placenta)
1. iPSC
(Ipsc)
42,790 Japanese ancestry individuals
Intelligence
28530673
2. Brain
(Gyrus)
1. Cancer
(Myeloma)
up to 58,799 European ancestry adults, up to 19,509 European ancestry children

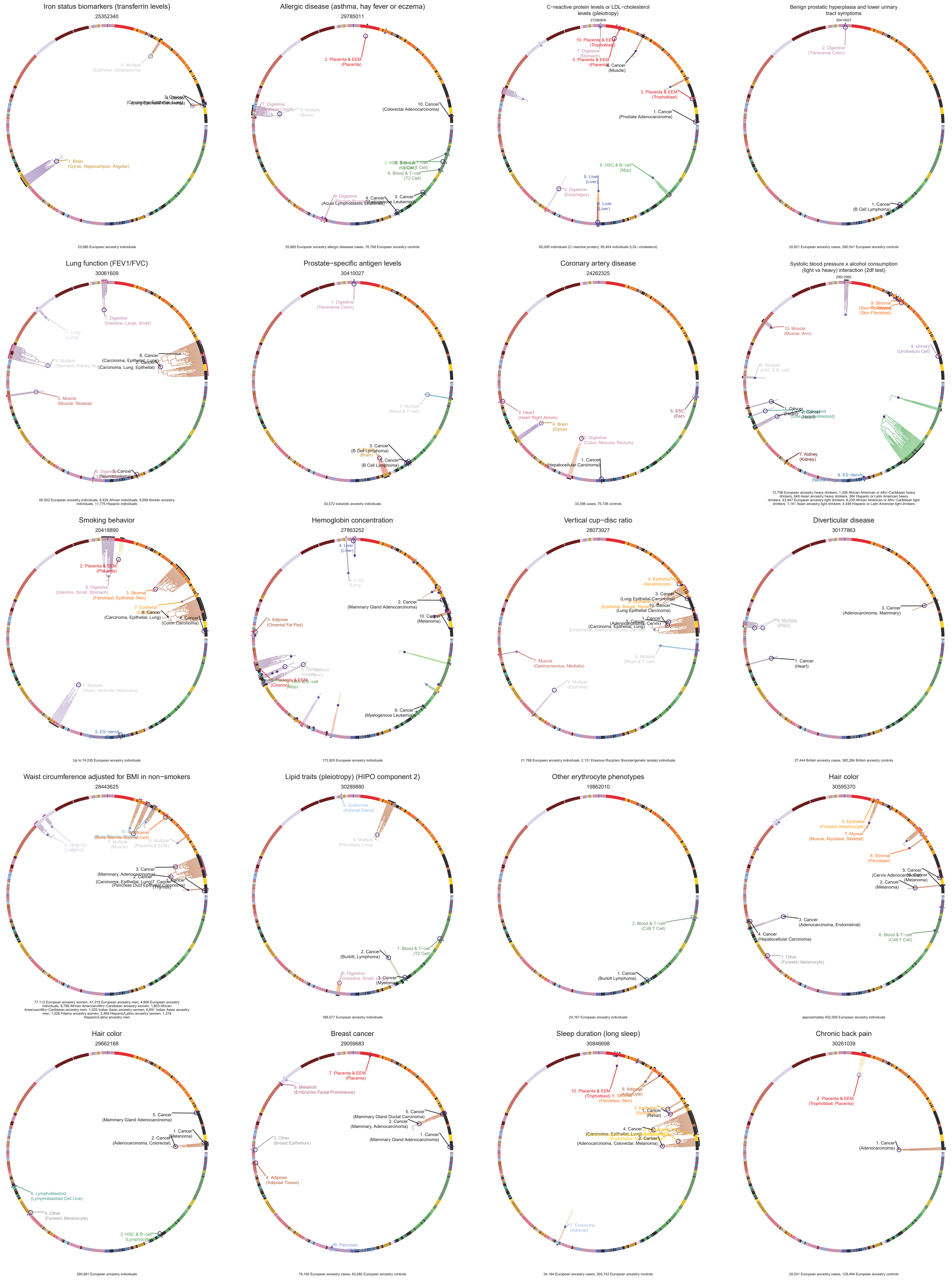

Iron status biomarkers (transferrin levels)
25352340
23,986 European ancestry individuals
Allergic disease (asthma, hay fever or eczema)
29785011
25,685 European ancestry allergic diseases cases, 76,768 European ancestry controls
C−reactive protein levels or LDL−cholesterol levels (pleiotropy)
27286809
65,000 individuals (C−reactive protein), 95,454 individuals (LDL−cholesterol)
Benign prostatic hyperplasia and lower urinary tract symptoms
30410027
20,621 European ancestry cases, 280,541 European ancestry controls
Lung function (FEV1/FVC)
30061609
60,552 European ancestry individuals, 8,429 African individuals, 9,959 Korean ancestry individuals, 11,775 Hispanic individuals
Prostate−specific antigen levels
30410027
33,572 Icelandic ancestry individuals
Coronary artery disease
24262325
33,398 cases, 75,726 controls
Systolic blood pressure x alcohol consumption (light vs heavy) interaction (2df test)
29912962
12,758 European ancestry heavy drinkers, 1,326 African American or Afro−Caribbean heavy drinkers, 648 Asian ancestry heavy drinkers, 364 Hispanic or Latin American heavy drinkers, 43,947 European ancestry light drinkers, 8,239 African American or Afro−Caribbean light drinkers, 1,141 Asian ancestry light drinkers, 4,448 Hispanic or Latin American light drinkers
Smoking behavior
20418890
Up to 74,035 European ancestry individuals
Hemoglobin concentration
27863252
172,925 European ancestry individuals
Vertical cup−disc ratio
28073927
21,768 European ancestry individuals, 2,131 Erasmus Rucphen (founder/genetic isolate) individuals
Diverticular disease
30177863
27,444 British ancestry cases, 382,284 British ancestry controls
Waist circumference adjusted for BMI in non−smokers
28443625
77,113 European ancestry women, 47,319 European ancestry men, 4,856 European ancestry individuals, 8,799 African American/Afro−Caribbean ancestry women, 1,803 African American/Afro−Caribbean ancestry men, 1,020 Indian Asian ancestry women, 6,691 Indian Asian ancestry men, 1,526 Filipino ancestry women, 2,469 Hispanic/Latino ancestry women, 1,318 Hispanic/Latino ancestry men
Lipid traits (pleiotropy) (HIPO component 2)
30289880
188,577 European ancestry individuals
Other erythrocyte phenotypes
19862010
24,167 European ancestry individuals
Hair color
30595370
approximately 452,000 European ancestry individuals
Hair color
29662168
290,891 European ancestry individuals
Breast cancer
29059683
76,192 European ancestry cases, 63,082 European ancestry controls
Sleep duration (long sleep)
30846698
34,184 European ancestry cases, 305,742 European ancestry controls
Chronic back pain
30261039
29,531 European ancestry cases, 128,494 European ancestry controls

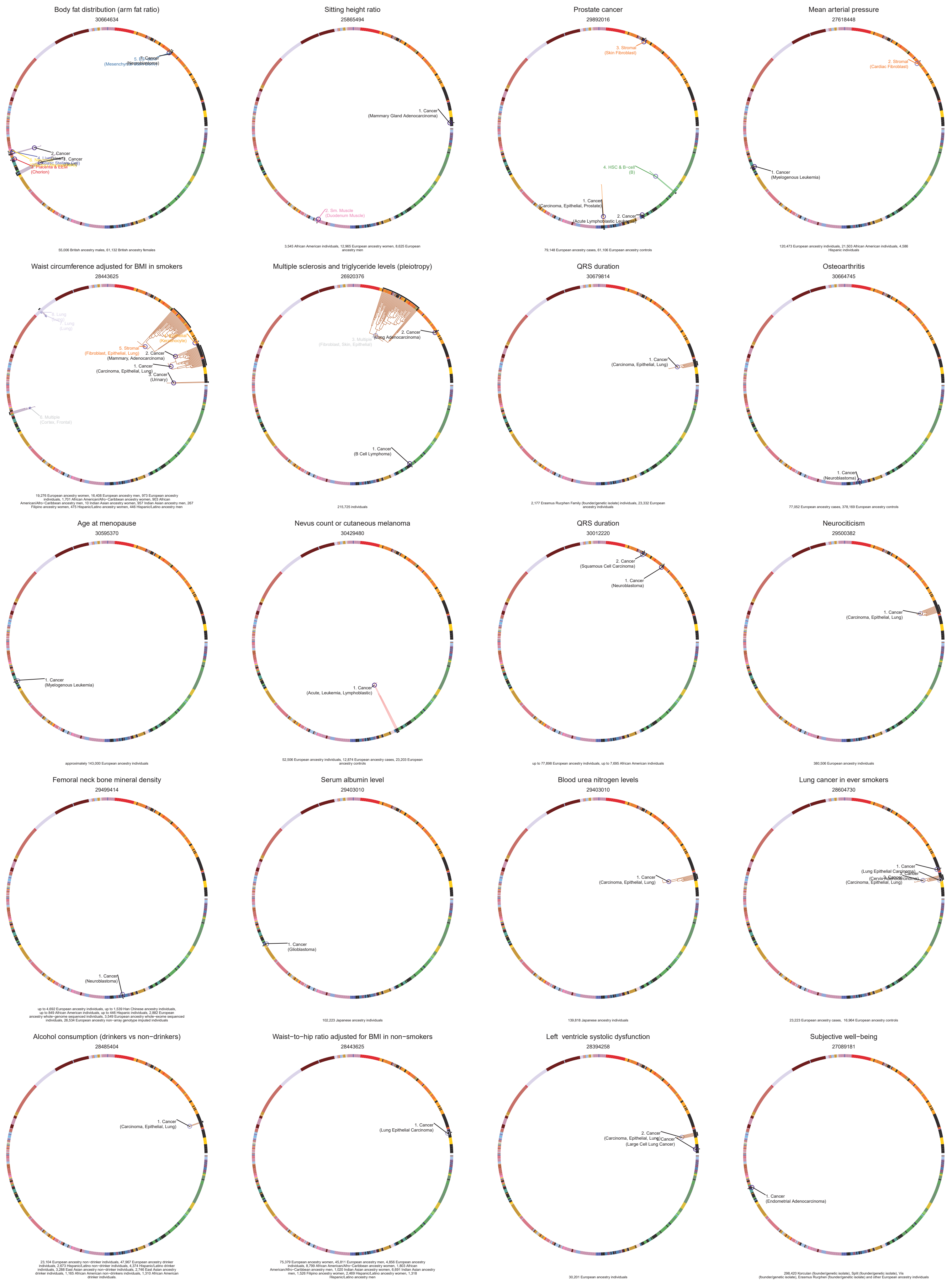

Body fat distribution (arm fat ratio)
30664634
2. Cancer
1. Cancer
(Hepatic Stellate Cell)
4. Liver
3. Placenta & EEM
(Chorion)
55,006 British ancestry males, 61,132 British ancestry females
Sitting height ratio
25865494
1. Cancer
(Mammary Gland Adenocarcinoma)
2. Sm. Muscle
(Duodenum Muscle)
3,545 African American individuals, 12,965 European ancestry women, 8,625 European ancestry men
Prostate cancer
29892016
3. Stromal
(Skin Fibroblast)
4. HSC & B-cell
(B)
1. Cancer
(Carcinoma, Epithelial, Prostate)
2. Cancer
(Acute Lymphoblastic Leukemia)
79,148 European ancestry cases, 61,106 European ancestry controls
Mean arterial pressure
27618448
2. Stromal
(Cardiac Fibroblast)
1. Cancer
(Myelogenous Leukemia)
120,473 European ancestry individuals, 21,503 African American individuals, 4,586 Hispanic individuals
Waist circumference adjusted for BMI in smokers
28443625
5. Stromal
(Fibroblast, Epithelial, Lung)
2. Cancer
(Mammary, Adenocarcinoma)
1. Cancer
(Carcinoma, Epithelial, Lung)
3. Cancer
(Urinary)
(Keratinocyte)
8. Multiple
(Cortex, Frontal)
19,276 European ancestry women, 16,408 European ancestry men, 973 European ancestry individuals, 1,701 African American/Afro-Caribbean ancestry women, 903 African American/Afro-Caribbean ancestry men, 10 Indian Asian ancestry women, 957 Indian Asian ancestry men, 267 Filipino ancestry women, 475 Hispanic/Latino ancestry women, 446 Hispanic/Latino ancestry men
Multiple sclerosis and triglyceride levels (pleiotropy)
26920376
2. Cancer
(Lung Adenocarcinoma)
3. Multiple
(Fibroblast, Skin, Epithelial)
1. Cancer
(B Cell Lymphoma)
215,725 individuals
QRS duration
30679814
1. Cancer
(Carcinoma, Epithelial, Lung)
2,177 Erasmus Rucphen Family (founder/genetic isolate) individuals, 23,332 European ancestry individuals
Osteoarthritis
30664745
1. Cancer
(Neuroblastoma)
77,052 European ancestry cases, 378,169 European ancestry controls
Age at menopause
30595370
1. Cancer
(Myelogenous Leukemia)
approximately 143,000 European ancestry individuals
Nevus count or cutaneous melanoma
30429480
1. Cancer
(Acute, Leukemia, Lymphoblastic)
52,506 European ancestry individuals, 12,874 European ancestry cases, 23,203 European ancestry controls
QRS duration
30012220
2. Cancer
(Squamous Cell Carcinoma)
1. Cancer
(Neuroblastoma)
up to 77,898 European ancestry individuals, up to 7,695 African American individuals
Neurociticism
29500382
1. Cancer
(Carcinoma, Epithelial, Lung)
380,506 European ancestry individuals
Femoral neck bone mineral density
29499414
1. Cancer
(Neuroblastoma)
up to 4,692 European ancestry individuals, up to 1,539 Han Chinese ancestry individuals, up to 849 African American individuals, up to 446 Hispanic individuals, 2,882 European ancestry whole−genome sequenced individuals, 3,549 European ancestry whole−exome sequenced individuals, 26,534 European ancestry non−array genotype imputed individuals
Serum albumin level
29403010
1. Cancer
(Glioblastoma)
102,223 Japanese ancestry individuals
Blood urea nitrogen levels
29403010
1. Cancer
(Carcinoma, Epithelial, Lung)
139,818 Japanese ancestry individuals
Lung cancer in ever smokers
28604730
1. Cancer
(Lung Epithelial Carcinoma)
(Carcinoma, Epithelial, Lung)
23,223 European ancestry cases, 16,964 European ancestry controls
Alcohol consumption (drinkers vs non−drinkers)
28485404
1. Cancer
(Carcinoma, Epithelial, Lung)
23,104 European ancestry non−drinker individuals, 47,967 European ancestry drinker individuals, 2,673 Hispanic/Latino non−drinker individuals, 4,374 Hispanic/Latino drinker individuals, 3,288 East Asian ancestry non−drinker individuals, 2,746 East Asian ancestry drinker individuals, 1,165 African American non−drinkers individuals, 1,310 African American drinker individuals.
Waist−to−hip ratio adjusted for BMI in non−smokers
28443625
1. Cancer
(Lung Epithelial Carcinoma)
75,379 European ancestry women, 45,811 European ancestry men, 4,856 European ancestry individuals, 8,799 African American/Afro−Caribbean ancestry women, 1,803 African American/Afro−Caribbean ancestry men, 1,020 Indian Asian ancestry women, 6,691 Indian Asian ancestry men, 1,526 Filipino ancestry women, 2,469 Hispanic/Latino ancestry women, 1,318 Hispanic/Latino ancestry men
Left ventricle systolic dysfunction
28394258
2. Cancer
(Carcinoma, Epithelial, Lung)
1. Cancer
(Large Cell Lung Cancer)
30,201 European ancestry individuals
Subjective well−being
27089181
1. Cancer
(Endometrial Adenocarcinoma)
298,420 Korculan (founder/genetic isolate), Split (founder/genetic isolate), Vis (founder/genetic isolate), Erasmus Rucphen (founder/genetic isolate) and other European ancestry individuals

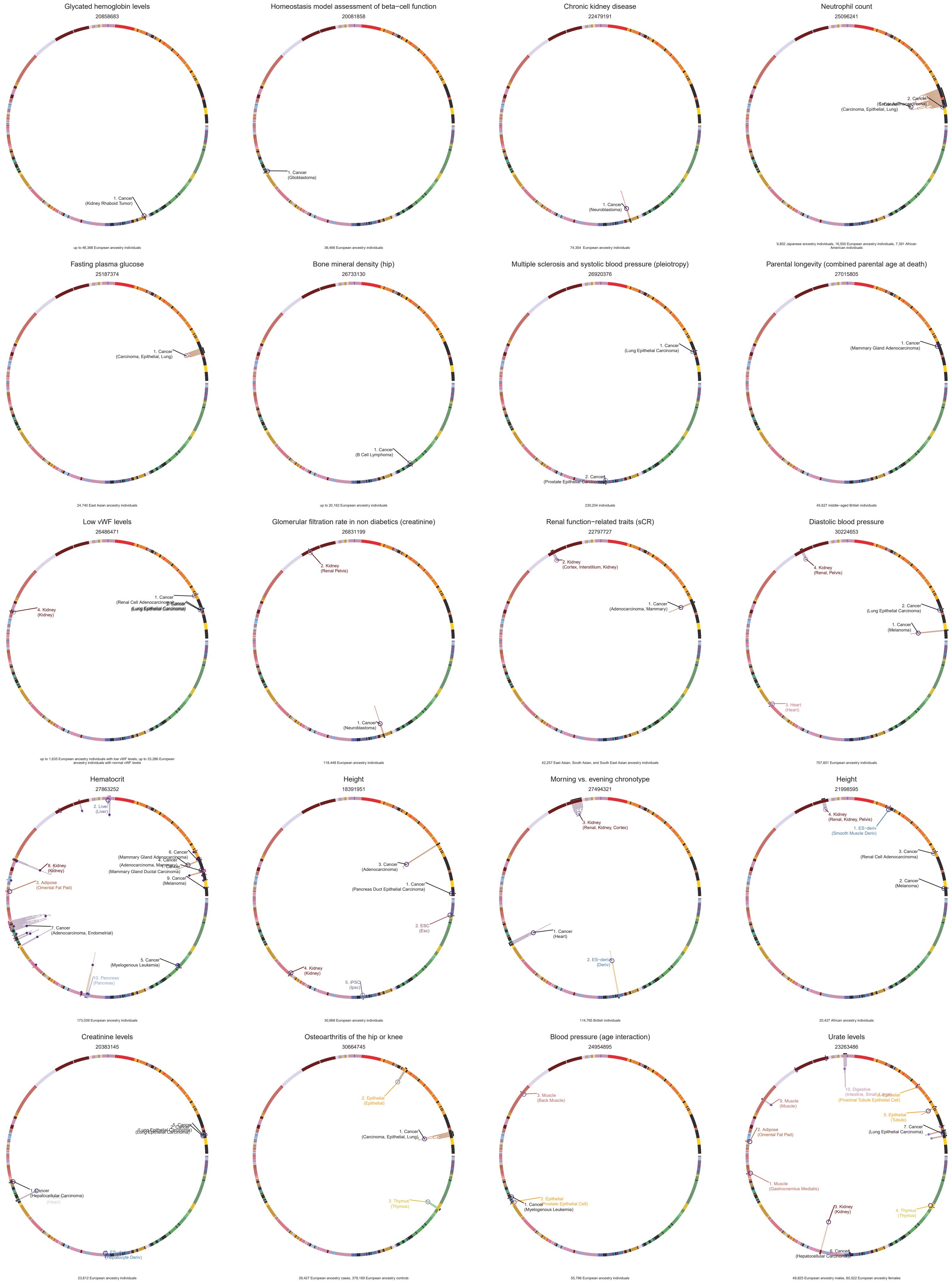

Glycated hemoglobin levels
20858683
1. Cancer
(Kidney Rhaboid Tumor)
up to 46,368 European ancestry individuals
Homeostasis model assessment of beta−cell function
20081858
1. Cancer
(Glioblastoma)
36,466 European ancestry individuals
Chronic kidney disease
22479191
1. Cancer
(Neuroblastoma)
74,354 European ancestry individuals
Neutrophil count
25096241
2. Cancer
(Carcinoma, Epithelial, Lung)
9,802 Japanese ancestry individuals, 16,550 European ancestry individuals, 7,391 African American individuals
Fasting plasma glucose
25187374
1. Cancer
(Carcinoma, Epithelial, Lung)
24,740 East Asian ancestry individuals
Bone mineral density (hip)
26733130
1. Cancer
(B Cell Lymphoma)
up to 20,162 European ancestry individuals
Multiple sclerosis and systolic blood pressure (pleiotropy)
26920376
1. Cancer
(Lung Epithelial Carcinoma)
2. Cancer
(Prostate Epithelial Carcinoma)
230,204 individuals
Parental longevity (combined parental age at death)
27015805
1. Cancer
(Mammary Gland Adenocarcinoma)
45,627 middle−aged British individuals
Low vWF levels
26486471
1. Cancer
(Renal Cell Adenocarcinoma)
4. Kidney
(Kidney)
up to 1,635 European ancestry individuals with low vWF levels, up to 33,286 European ancestry individuals with normal vWF levels
Glomerular filtration rate in non diabetics (creatinine)
26831199
2. Kidney
(Renal Pelvis)
1. Cancer
(Neuroblastoma)
118,448 European ancestry individuals
Renal function−related traits (sCR)
22797727
2. Kidney
(Cortex, Interstitium, Kidney)
1. Cancer
(Adenocarcinoma, Mammary)
42,257 East Asian, South Asian, and South East Asian ancestry individuals
Diastolic blood pressure
30224653
4. Kidney
(Renal, Pelvis)
2. Cancer
(Lung Epithelial Carcinoma)
1. Cancer
(Melanoma)
3. Heart
(Heart)
757,601 European ancestry individuals
Hematocrit
27863252
2. Liver
(Liver)
6. Cancer
(Mammary Gland Adenocarcinoma)
4. Cancer
(Adenocarcinoma, Mammary)
(Mammary Gland Ductal Carcinoma)
9. Cancer
(Melanoma)
8. Kidney
(Kidney)
3. Adipose
(Omental Fat Pad)
7. Cancer
(Adenocarcinoma, Endometrial)
5. Cancer
(Myelogenous Leukemia)
10. Pancreas
(Pancreas)
173,039 European ancestry individuals
Height
18391951
3. Cancer
(Adenocarcinoma)
1. Cancer
(Pancreas Duct Epithelial Carcinoma)
2. ESC
(Esc)
4. Kidney
(Kidney)
5. iPSC
(Ipsc)
30,968 European ancestry individuals
Morning vs. evening chronotype
27494321
3. Kidney
(Renal, Kidney, Cortex)
1. Cancer
(Heart)
2. ES−deriv
(Deriv)
114,765 British individuals
Height
21998595
4. Kidney
(Renal, Kidney, Pelvis)
1. ES−deriv
(Smooth Muscle Deriv)
3. Cancer
(Renal Cell Adenocarcinoma)
2. Cancer
(Melanoma)
20,427 African ancestry individuals
Creatinine levels
20383145
(Lung Epithelial Carcinoma)
1. Cancer
(Hepatocellular Carcinoma)
(Heart)
(Hepatocyte Deriv)
23,812 European ancestry individuals
Osteoarthritis of the hip or knee
30664745
2. Epithelial
(Epithelial)
1. Cancer
(Carcinoma, Epithelial, Lung)
3. Thymus
(Thymus)
39,427 European ancestry cases, 378,169 European ancestry controls
Blood pressure (age interaction)
24954895
3. Muscle
(Back Muscle)
2. Epithelial
(Prostate Epithelial Cell)
1. Cancer
(Myelogenous Leukemia)
55,796 European ancestry individuals
Urate levels
23263486
10. Digestive
(Proximal Tubule Epithelial Cell)
9. Muscle
(Muscle)
5. Epithelial
(Tubule)
7. Cancer
(Lung Epithelial Carcinoma)
2. Adipose
(Omental Fat Pad)
1. Muscle
(Gastrocnemius Medialis)
3. Kidney
(Kidney)
4. Thymus
(Thymus)
6. Cancer
(Hepatocellular Carcinoma)
49,825 European ancestry males, 60,522 European ancestry females

Mean arterial pressure
26390057

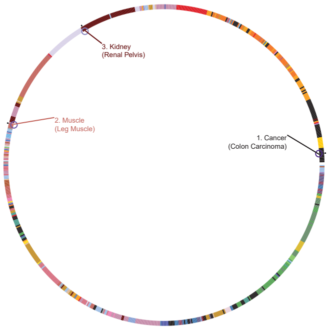


31,516 East Asian ancestry individuals, 35,352 European ancestry individuals, 33,126 South Asian ancestry individuals

Prostate−specific antigen levels (conditioned on lead SNPs)
28139693

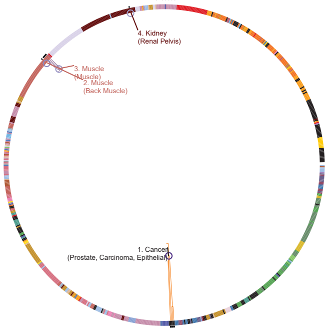


28,503 European ancestry individuals

Nevus count
30429480

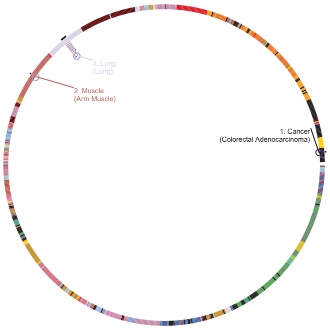


52,506 European ancestry individuals

Diastolic blood pressure
28135244

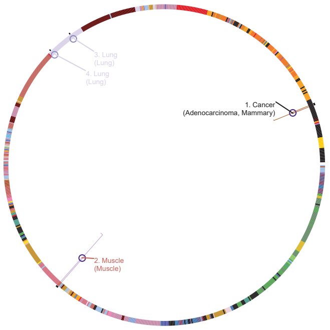


140,886 European ancestry individuals

Waist circumference adjusted for body mass index
25673412

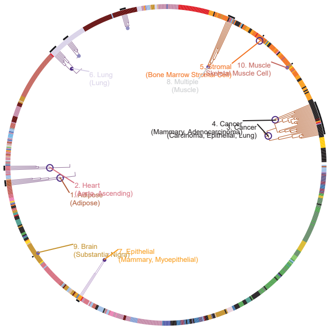


up to 56,910 European ancestry men, up to 86,570 European ancestry women

Waist−to−hip ratio adjusted for BMI (joint analysis for main effect and physical activity interaction)
28448500

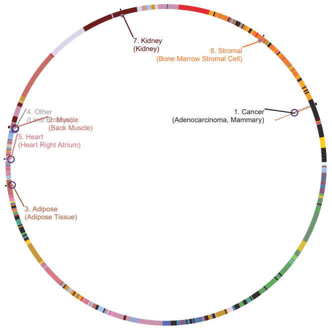


2,518 African American women, 83,707 European ancestry women, 1,773 Filipino ancestry women, 1,030 Indian ancestry women, 1,457 African American men, 52,993 European ancestry men, 7,648 Indian ancestry men, 5,527 European ancestry individuals

Monocyte count
29403010

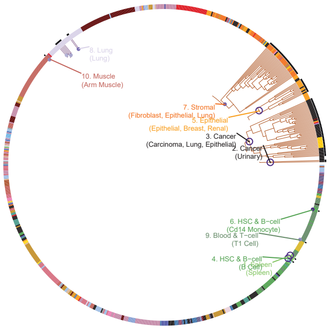


62,076 Japanese ancestry individuals

Mean corpuscular hemoglobin concentration
28017375

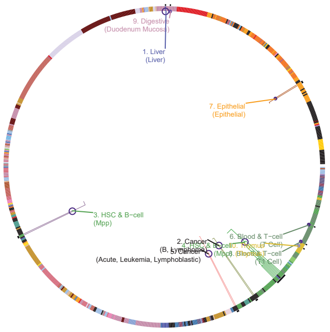


up to 21,020 European ancestry individuals, up to 3,621 African American individuals, up to 15,062 East Asian individuals.

Serum total protein level
29403010

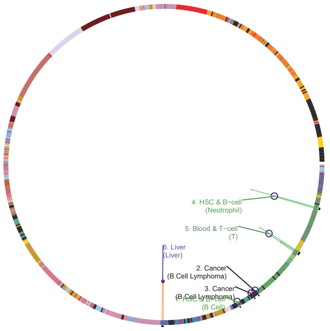


113,509 Japanese ancestry individuals

Fasting blood glucose
20081858

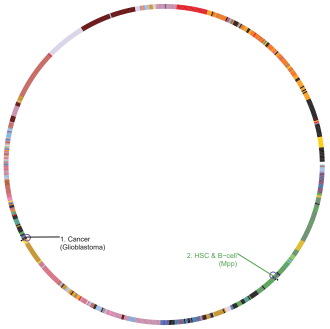


46,186 European ancestry individuals

Insomnia complaints (continuous)
28604731

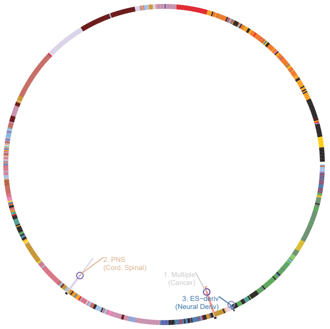


12,863 European ancestry male cases, 19,521 European ancestry female cases, 40,776 European ancestry male controls, 39,846 European ancestry female controls

Insomnia complaints (dichotomous)
28604731

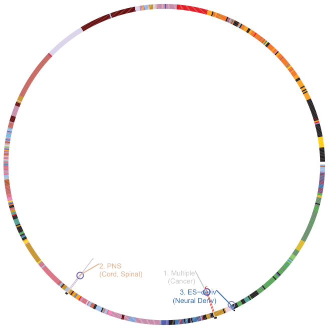


32,384 European ancestry cases, 27,128 European ancestry controls

Insomnia complaints
28604731

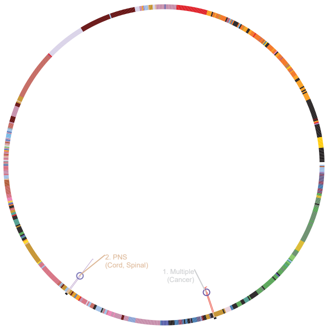


12,863 European ancestry male cases, 19,521 European ancestry female cases, 40,776 European ancestry male controls, 39,846 European ancestry female controls

Heel bone mineral density
28869591

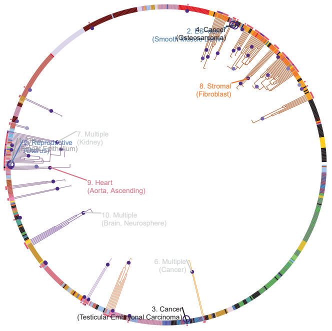


76,067 European ancestry women, 66,420 European ancestry men

Waist circumference adjusted for BMI in active individuals
28448500

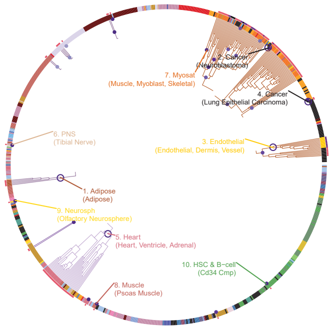


1,957 African American women, 1,773 Filipino ancestry women, 305 Indian ancestry women, 67,139 European ancestry women, 1,206 African American men, 2,029 Indian ancestry men, 43,100 European ancestry men, at least 4,819 European ancestry individuals

White matter hyperintensity burden
25663218

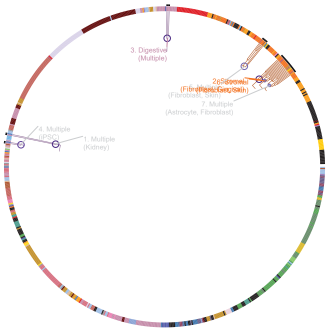


17,936 European ancestry individuals, 1,943 African American ancestry individuals, 795 Hispanic individuals, 204 Chinese ancestry individuals, 201 Malay ancestry individuals

Pulse pressure x alcohol consumption interaction (2df test)
29912962

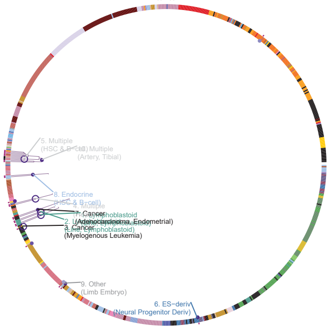


91,099 European ancestry drinkers and non−drinkers, 21,416 African American or Afro−Caribbean drinkers and non−drinkers, 12,365 Asian ancestry drinkers and non−drinkers, 8,470 Hispanic or Latin American drinkers and non−drinkers

Pulmonary function (smoking interaction)
23284291

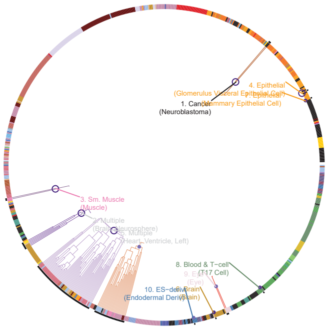


50,047 European ancestry individuals

Birth weight
23202124

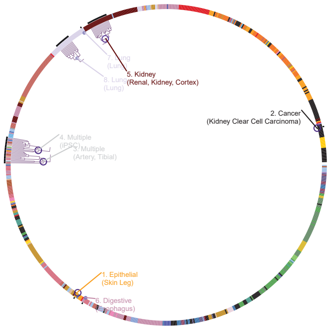


Up to 26,836 European ancestry individuals

Age at first birth
27089180

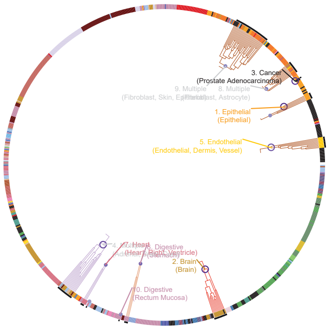


66,310 British ancestry females

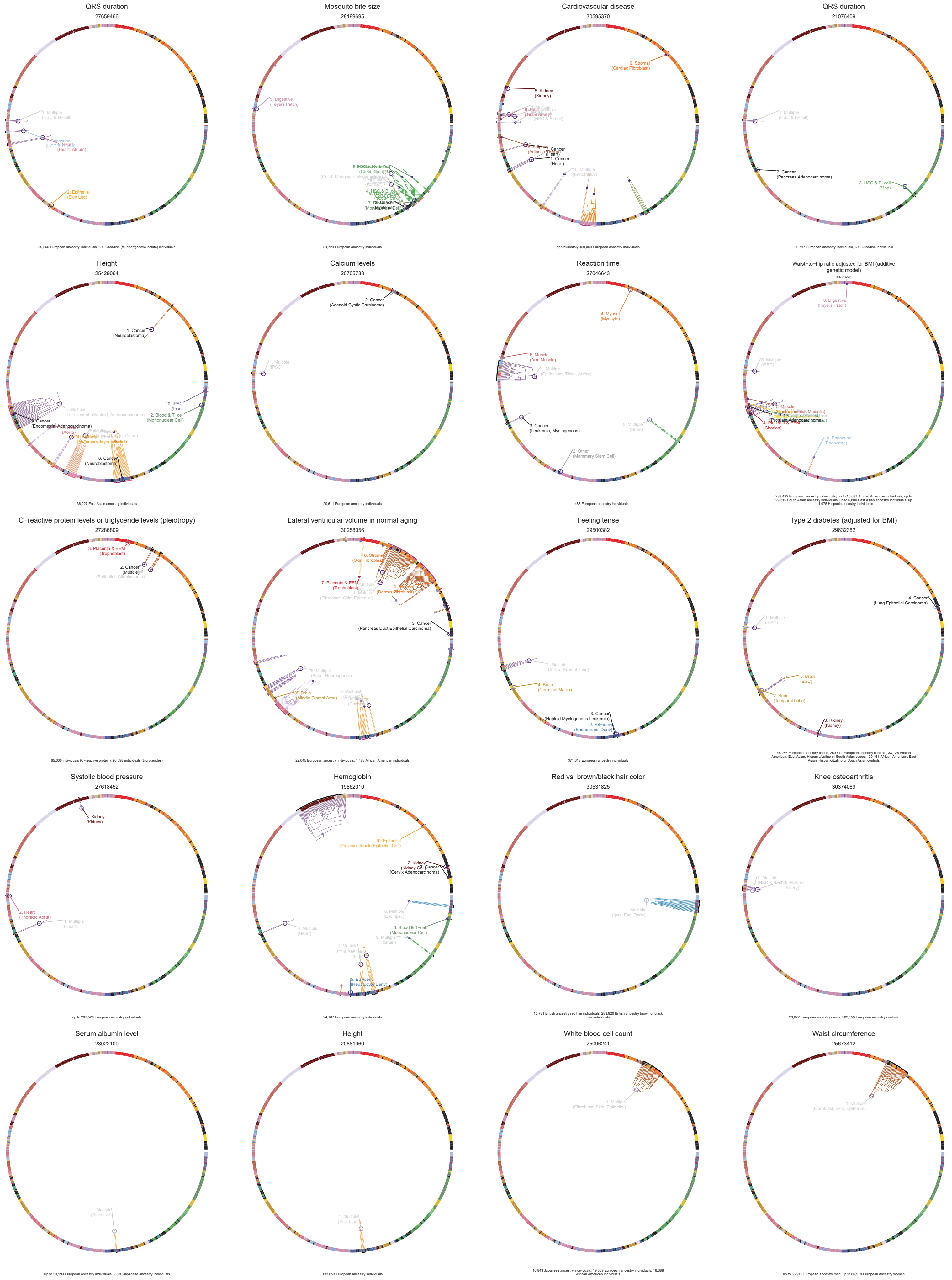

QRS duration
27659466
1. Multiple
(HSC & B−cell)
2. Epithelial
(Skin Leg)
59,565 European ancestry individuals, 690 Orcadian (founder/genetic isolate) individuals
Mosquito bite size
28199695
5. Digestive
(Peyers Patch)
84,724 European ancestry individuals
Cardiovascular disease
30595370
9. Stromal
(Cardiac Fibroblast)
5. Kidney
(Kidney)
1. Cancer
(Heart)
10. Multiple
(Endothelial)
approximately 459,000 European ancestry individuals
QRS duration
21076409
1. Multiple
(HSC & B−cell)
2. Cancer
(Pancreas Adenocarcinoma)
3. HSC & B−cell
(Mpp)
39,717 European ancestry individuals, 690 Orcadian individuals
Height
25429064
1. Cancer
(Neuroblastoma)
3. Multiple
(Line, Lymphoblastoid, Adenocarcinoma)
10. iPSC
(Ipsc)
2. Blood & T−cell
(Mononuclear Cell)
6. Cancer
(Neuroblastoma)
36,227 East Asian ancestry individuals
Calcium levels
20705733
2. Cancer
(Adenoid Cystic Carcinoma)
1. Multiple
(iPSC)
20,611 European ancestry individuals
Reaction time
27046643
4. Myosat
(Myocyte)
6. Muscle
(Arm Muscle)
1. Multiple
(Epithelium, Tibial, Artery)
3. Cancer
(Leukemia, Myelogenous)
5. Multiple
(Brain)
2. Other
(Mammary Stem Cell)
111,483 European ancestry individuals
Waist−to−hip ratio adjusted for BMI (additive genetic model)
30778226
6. Digestive
(Peyers Patch)
5. Multiple
(iPSC)
4. Placenta & EEM
(Chorion)
10. Endocrine
(Endocrine)
288,492 European ancestry individuals, up to 15,687 African American individuals, up to 29,315 South Asian ancestry individuals, up to 6,800 East Asian ancestry individuals, up to 4,075 Hispanic ancestry individuals
C−reactive protein levels or triglyceride levels (pleiotropy)
27286809
3. Placenta & EEM
(Trophoblast)
2. Cancer
(Muscle)
65,000 individuals (C−reactive protein), 96,598 individuals (triglycerides)
Lateral ventricular volume in normal aging
30258056
8. Stromal
(Skin Fibroblast)
7. Placenta & EEM
(Trophoblast)
3. Cancer
(Pancreas Duct Epithelial Carcinoma)
5. Multiple
(Brain, Neurosphere)
6. Brain
(Middle Frontal Area)
22,045 European ancestry individuals, 1,488 African American individuals
Feeling tense
29500382
1. Multiple
(Cortex, Frontal, Line)
4. Brain
(Germinal Matrix)
3. Cancer
(Haploid Myelogenous Leukemia)
2. ES−deriv
(Endodermal Deriv)
371,318 European ancestry individuals
Type 2 diabetes (adjusted for BMI)
29632382
4. Cancer
(Lung Epithelial Carcinoma)
1. Multiple
(iPSC)
5. Brain
(ESC)
2. Brain
(Temporal Lobe)
3. Kidney
(Kidney)
48,286 European ancestry cases, 250,671 European ancestry controls, 33,126 African American, East Asian, Hispanic/Latino or South Asian cases, 120,161 African American, East Asian, Hispanic/Latino or South Asian controls
Systolic blood pressure
27618452
3. Kidney
(Kidney)
2. Heart
(Thoracic Aorta)
1. Multiple
(Heart)
up to 201,529 European ancestry individuals
Hemoglobin
19862010
10. Epithelial
(Proximal Tubule Epithelial Cell)
(Cervix Adenocarcinoma)
8. Multiple
(Esc, Ipsc)
6. Blood & T−cell
(Mononuclear Cell)
9. Multiple
(Brain)
3. Multiple
(Heart)
5. ES−deriv
(Hepatocyte Deriv)
24,167 European ancestry individuals
Red vs. brown/black hair color
30531825
1. Multiple
(Ipsc, Esc, Deriv)
15,731 British ancestry red hair individuals, 283,920 British ancestry brown or black hair individuals
Knee osteoarthritis
30374069
2. Multiple
(Artery)
23,877 European ancestry cases, 562,153 European ancestry controls
Serum albumin level
23022100
1. Multiple
(Digestive)
Up to 53,190 European ancestry individuals, 9,380 Japanese ancestry individuals
Height
20881960
1. Multiple
(Esc, Ipsc)
133,653 European ancestry individuals
White blood cell count
25096241
1. Multiple
(Fibroblast, Skin, Epithelial)
16,843 Japanese ancestry individuals, 19,509 European ancestry individuals, 16,388 African American individuals
Waist circumference
25673412
1. Multiple
(Fibroblast, Skin, Epithelial)
up to 56,910 European ancestry men, up to 86,570 European ancestry women

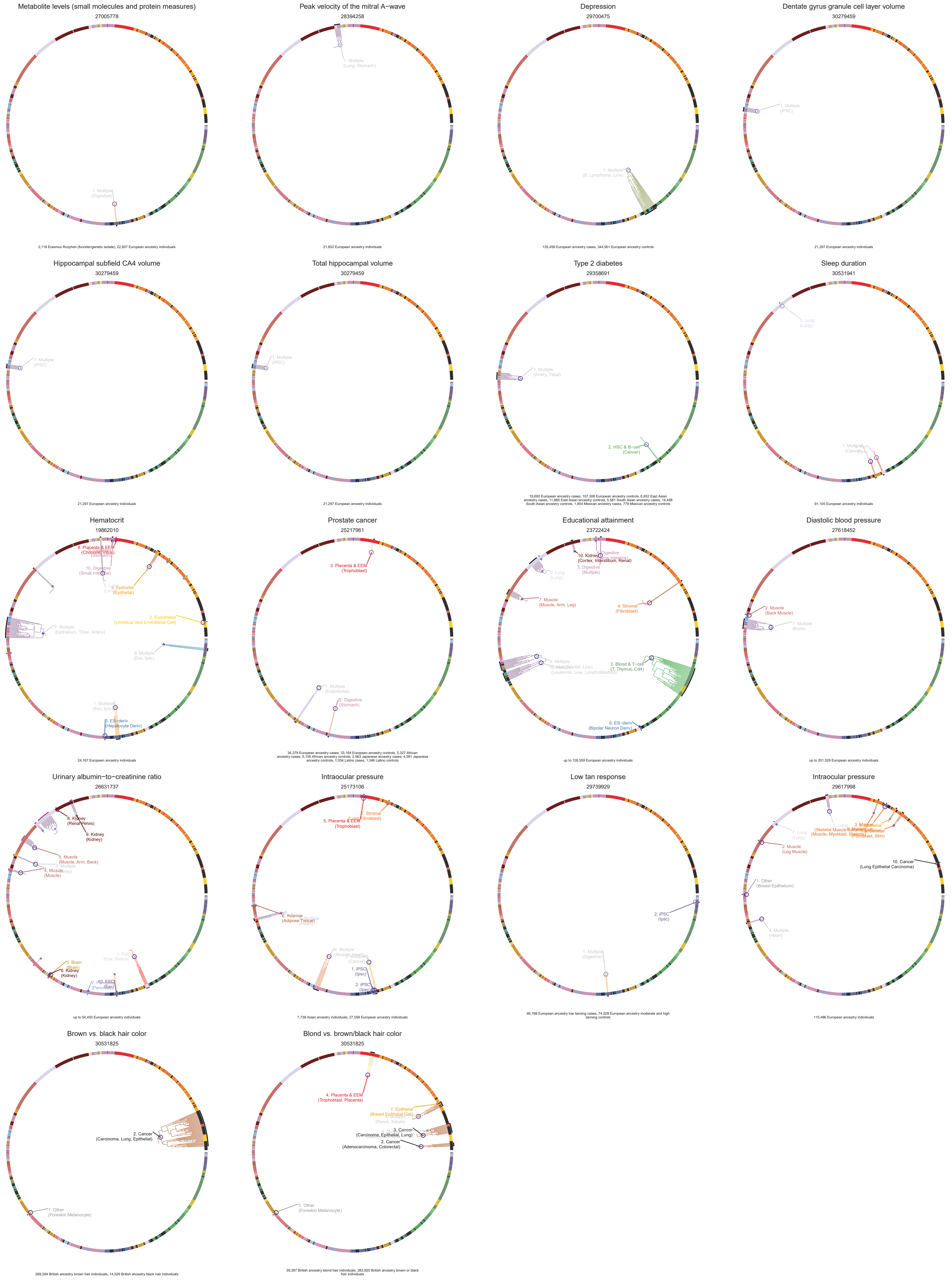
Metabolite levels (small molecules and protein measures)
27005778
2,118 Erasmus Rucphen (founder/genetic isolate), 22,807 European ancestry individuals
Peak velocity of the mitral A−wave
28394258
21,852 European ancestry individuals
Depression
29700475
135,458 European ancestry cases, 344,901 European ancestry controls
Dentate gyrus granule cell layer volume
30279459
21,297 European ancestry individuals
Hippocampal subfield CA4 volume
30279459
21,297 European ancestry individuals
Total hippocampal volume
30279459
21,297 European ancestry individuals
Type 2 diabetes
29358691
19,693 European ancestry cases, 107,308 European ancestry controls, 6,952 East Asian ancestry cases, 11,865 East Asian ancestry controls, 5,561 South Asian ancestry cases, 14,458 South Asian ancestry controls, 1,804 Mexican ancestry cases, 779 Mexican ancestry controls
Sleep duration
30531941
91,105 European ancestry individuals
Hematocrit
19862010
24,167 European ancestry individuals
Prostate cancer
25217961
34,379 European ancestry cases, 33,164 European ancestry controls, 5,327 African ancestry cases, 5,136 African ancestry controls, 2,563 Japanese ancestry cases, 4,391 Japanese ancestry controls, 1,034 Latino cases, 1,046 Latino controls
Educational attainment
23722424
up to 126,559 European ancestry individuals
Diastolic blood pressure
27618452
up to 201,529 European ancestry individuals
Urinary albumin−to−creatinine ratio
26631737
up to 54,450 European ancestry individuals
Intraocular pressure
25173106
7,738 Asian ancestry individuals, 27,558 European ancestry individuals
Low tan response
29739929
46,768 European ancestry low tanning cases, 74,528 European ancestry moderate and high tanning controls
Intraocular pressure
29617998
115,486 European ancestry individuals
Brown vs. black hair color
30531825
269,394 British ancestry brown hair individuals, 14,526 British ancestry black hair individuals
Blond vs. brown/black hair color
30531825
39,397 British ancestry blond hair individuals, 283,920 British ancestry brown or black hair individuals
